# Lessons on fruiting body morphogenesis from genomes and transcriptomes of Agaricomycetes

**DOI:** 10.1101/2021.12.09.471732

**Authors:** László G. Nagy, Peter Jan Vonk, Markus Künzler, Csenge Földi, Máté Virágh, Robin A. Ohm, Florian Hennicke, Balázs Bálint, Árpád Csernetics, Botond Hegedüs, Zhihao Hou, Xiao-Bin Liu, Shen Nan, Manish Pareek, Neha Sahu, Benedek Szathmári, Torda Varga, Hongli Wu, Xiao Yang, Zsolt Merényi

## Abstract

Fruiting bodies of mushroom-forming fungi (Agaricomycetes) are among the most complex structures produced by fungi. Unlike vegetative hyphae, fruiting bodies grow determinately and follow a genetically encoded developmental program that orchestrates tissue differentiation, growth and sexual sporulation. In spite of more than a century of research, our understanding of the molecular details of fruiting body morphogenesis is limited and a general synthesis on the genetics of this complex process is lacking. In this paper, we aim to comprehensively identify conserved genes related to fruiting body morphogenesis and distill novel functional hypotheses for functionally poorly characterized genes. As a result of this analysis, we report 921 conserved developmentally expressed gene families, only a few dozens of which have previously been reported in fruiting body development. Based on literature data, conserved expression patterns and functional annotations, we provide informed hypotheses on the potential role of these gene families in fruiting body development, yielding the most complete description of molecular processes in fruiting body morphogenesis to date. We discuss genes related to the initiation of fruiting, differentiation, growth, cell surface and cell wall, defense, transcriptional regulation as well as signal transduction. Based on these data we derive a general model of fruiting body development, which includes an early, proliferative phase that is mostly concerned with laying out the mushroom body plan (via cell division and differentiation), and a second phase of growth via cell expansion as well as meiotic events and sporulation. Altogether, our discussions cover 1480 genes of *Coprinopsis cinerea*, and their orthologs in *Agaricus bisporus, Cyclocybe aegerita, Armillaria ostoyae, Auriculariopsis ampla, Laccaria bicolor, Lentinula edodes, Lentinus tigrinus, Mycena kentingensis, Phanerochaete chrysosporium, Pleurotus ostreatus,* and *Schizophyllum commune*, providing functional hypotheses for ∼10% of genes in the genomes of these species. Although experimental evidence for the role of these genes will need to be established in the future, our data provide a roadmap for guiding functional analyses of fruiting related genes in the Agaricomycetes. We anticipate that the gene compendium presented here, combined with developments in functional genomics approaches will contribute to uncovering the genetic bases of one of the most spectacular multicellular developmental processes in fungi.

## 1. Introduction

Fungi are one of the five main extant lineages in which complex macroscopic structures emerged over the course of evolution. The best known macroscopic structures of fungi are sexual fruiting bodies, which enclose spore-producing cells into a 3-dimensional, protective environment and facilitate spore dispersal(Nagy et al. 2018). Sexual fruiting bodies are complex multicellular structures based on the definition of Knoll(Knoll 2011) and show evidence for repeated evolution across the fungal tree of life(Stajich et al. 2009; Merényi et al. 2020a). They are found in at least 8 lineages, with best known representatives belonging to the Pezizomycotina (e.g. morels, truffles), Agaricomycotina (e.g. agarics, boletes), however, spectacular examples evolved in several other lineages, such as *Endogone* spp. in the Mucoromycota(Chang et al. 2019b) or the enigmatic *Neolecta* genus in the Taphrinomycotina(Nagy 2017; Nguyen et al. 2017). Fruiting bodies reached highest complexity in the Agaricomycetes, which include developmentally integrated morphologies that follow a spatially and temporally tightly regulated developmental program(Hibbett 2004; Virágh et al. 2021). In terms of complexity level, certain Agaricomycete fruiting bodies approach the complexity of simple animals and plants(Taylor and Ellison 2010; Nagy et al. 2018).

Fruiting body morphogenesis has been a subject of intense research, however, despite tremendous efforts in this field, morphogenesis of Basidiomycota fruiting bodies is quite poorly known. In comparison, fruiting body development and its underlying genetics are considerably better explored in the Ascomycota (see e.g.(Pöggeler et al. 2006a)). This may be due to a mixture of factors, including the complexity of Basidiomycota developmental programs, the lack of easy-to-use model systems or the relative difficulty of hypothesis testing via genetic manipulation (e.g. due to low frequency of homologous recombination(De Jong et al. 2010)). Research on Basidiomycete fruiting body morphogenesis has recently been relying largely on comparative -omics techniques, which provided insights into the gene repertoires and the regulation of gene expression during fruiting body development. This complements the rich body of literature on functionally characterized genes in *Coprinopsis cinerea* (summarized in Kües(Kües 2000)), *Schizophyllum commune* and other species. There are several recent milestones in fruiting body research such as the publication of key genome sequences(Ohm et al. 2010; Stajich et al. 2010; Morin et al. 2012) or comparative transcriptomic datasets (e.g. (Morin et al. 2012; Plaza et al. 2014a; Kües and Navarro-González 2015; Almási et al. 2019; Krizsán et al. 2019; Merényi et al. 2021)). An emerging driving force of research on fruiting body morphogenesis is the mushroom industry, which produces sexual fruiting bodies for food or as sources of bioactive compounds(Royse et al. 2017; Ma et al. 2018). Fruiting body development has been a subject of several excellent reviews, surprisingly, however, the last gene-centric review of fruiting body development in mushroom-forming fungi was published in 2000(Whiteford and Thurston 2000). This paper aims to complement recent reviews on the morphogenesis of vegetative hyphae(Riquelme et al. 2018), and on the genome of *Neurospora crassa*(Borkovich et al. 2004) with knowledge on fruiting body morphogenesis in the Agaricomycetes.

The aim of this paper is to systematically catalog genes and genetic processes related to fruiting body morphogenesis in the Agaricomycetes, based on both literature reviews and a meta-analysis of published developmental transcriptomes. We provide a handbook-like inventory of gene families and their putative functions based on expression data and functional annotations borrowed from well-researched model systems. Because a negligible proportion of morphogenesis genes has so far been functionally characterized in the Agaricomycetes, it is currently impossible to build a comprehensive picture based on mechanistic studies alone, as has recently been done for hypha morphogenesis(Riquelme et al. 2018). Instead, we derive functional hypotheses for genes based on comparative expression data across multiple species. Our starting point is that genes which are developmentally regulated in the majority of species for which transcriptome data are available, should belong to the core pathways of fruiting body morphogenesis, whereas genes that show species-specific expression patterns should be responsible for sculpting species-specific morphologies or represent transcriptional or technical noise. Finally, we incorporate novel and previous observations into a new synthesis on processes active during fruiting body morphogenesis.

### 1.1. What are fruiting bodies and how do they develop

Fruiting bodies, also called basidiomes, basidiocarps or simply ‘mushrooms’ in lay language, may be some of the best known structures of fungi. They are sexual reproductive organs that facilitate spore dispersal. While in simple Basidiomycota spores are born on naked basidia, mushroom-forming fungi evolved complex 3-dimensional structures that offer tremendous advantages in spore dispersal efficiency.

Developmental events that take place in fruiting bodies have been investigated previously in several species, of which *Coprinopsis/Coprinellus* spp. and *S. commune* are the two most widely used groups of models. The development of fruiting bodies in these species has been described in previous reviews(Kües 2000; Kües and Liu 2000; Palmer and Horton 2006; Kües and Navarro-González 2015), therefore, we here only provide a short introduction, and describe the process in more detail (combining new results) in Chapter 5. Briefly, fruiting body development is initiated on sexually competent mycelia as the reprogramming of hyphal branching patterns, which results in the interlacing of hyphae to form a 3-dimensional fruiting body initial(Kües and Liu 2000; Kües and Navarro-González 2015). Rather than relying on zygotic (e.g. maternal) mRNA cargo as animals and plants do, fungal hyphae undergo transcriptional reprogramming during fruiting body initiation. This reprogramming is influenced and triggered by a mixture of - often species-specific - factors, such as starvation, light or temperature changes (reviewed by(Sakamoto 2018)). Once appropriate internal and external signals converge on the initiation of fruiting, morphogenesis starts with an undifferentiated initial (hyphal knot, aggregate), which undergoes tissue differentiation to form primordia. In subsequent late primordium and young fruiting body stages rapid growth (stipe elongation and cap expansion) and meiosis happen in a coordinated manner, which ultimately results in species-specific mature morphologies and culminates in spore formation and release.

Development of the fruiting bodies is a tightly regulated process that has a high degree of autonomy. That is, once started, the developmental program is not interrupted by even large perturbations, such as injuries, or removal of the fruiting body from the supporting colony at or after certain stages(Moore 2013). In fruiting bodies not only morphogenetic events, but also several internal processes (e.g. nutritional and sexual) are developmentally regulated. For example, in *C. cinerea*, stipe elongation starts immediately after meiosis takes place, after which spore production, maturation and cap expansion follow in a tightly choreographed chronology(Kües 2000; Moore 2013). For detailed description of the process and molecular summaries of current knowledge on fruiting body morphogenesis the reader is referred to recent reviews(Kües 2000; Kües and Navarro-González 2015; Virágh et al. 2021) and the following detailed chapters of this paper.

### 1.2. The evolution of Agaricomycete fruiting bodies: from simple to complex morphogenesis

How fruiting body complexity evolved has interested mycologists for decades and has received considerable attention recently. Therefore we provide only a brief introduction here and refer the reader to recent reviews on the topic (e.g.(Kües and Navarro-González 2015; Nagy et al. 2018; Virágh et al. 2021)). Phylogenetic studies have made tremendous progress in uncovering broad patterns of morphological transformation and trends in fruiting body evolution (e.g. (Hibbett 2004; Varga et al. 2019; Sánchez-García et al. 2020)). Fruiting body morphologies show a great diversity in the Basidiomycota, from naked lawns of basidia, to resupinate, coralloid or highly complex, or the so called pileate-stipitate fruiting bodies of well-known agarics (e.g. *C. cinerea*) (Fig. 1).

**Fig. 1.**
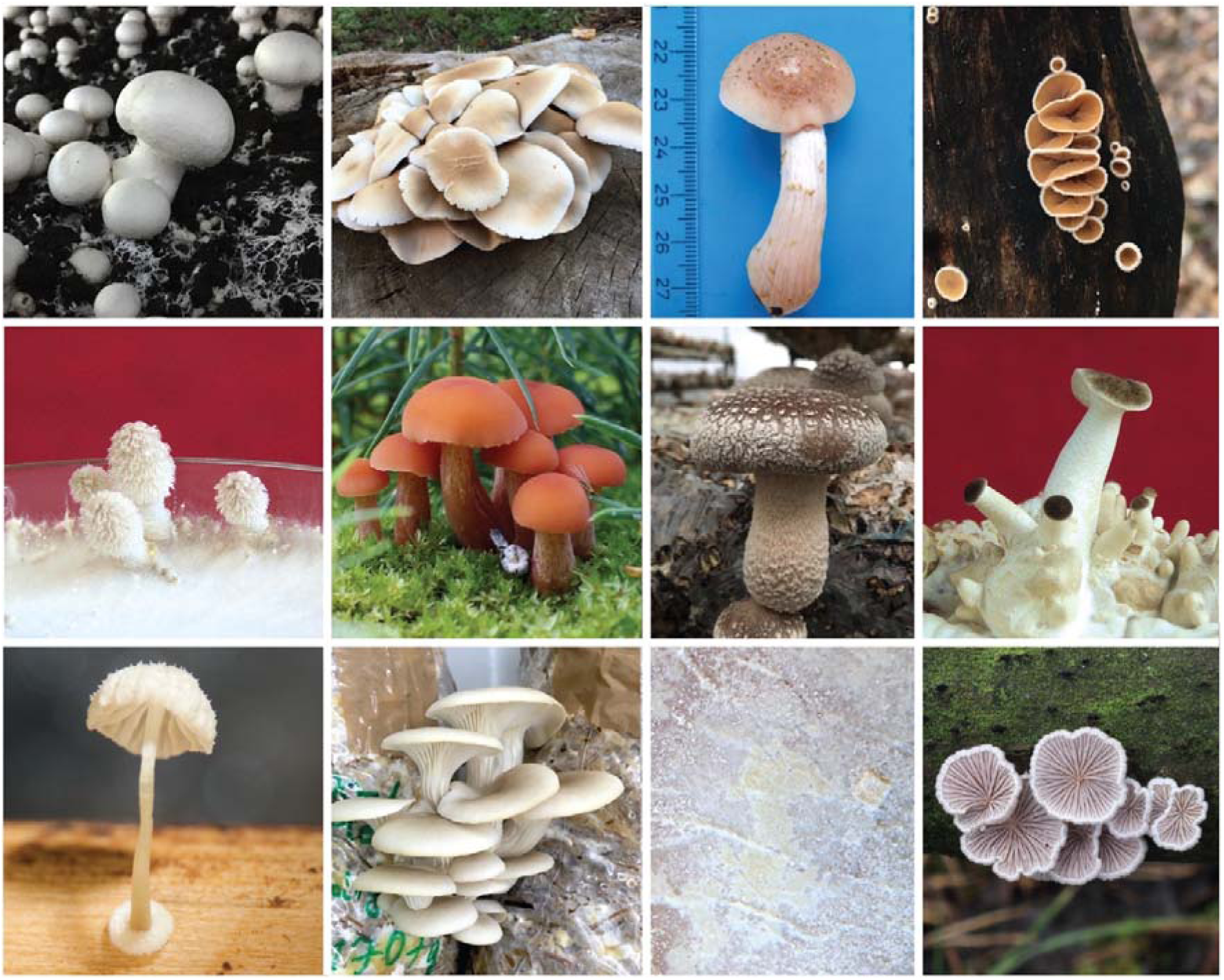
Species used in this study. Top row left to right: Agaricus bisporus, Cyclocybe aegerita, Armillaria ostoyae, Auriculariopsis ampla; Middle row: Coprinopsis cinerea, Laccaria bicolor, Lentinula edodes, Lentinus tigrinus; Bottom row: Mycena kentingensis, Phanerochaete chrysosporium, Pleurotus ostreatus, Schizophyllum commune

Agaricomycete fruiting bodies might have emerged in or around the last common ancestor of Dacrymycetes and Agaricomycetes over 400 million years ago(Varga et al. 2019; Prasanna et al. 2020), although there is some uncertainty around ancestral states. This is partly due to the presence of jelly-like telia reminiscent of fruiting bodies in the Pucciniomycotina, uncertainties in the branching order of Basidiomycota subphyla and reductive evolution in early-diverging clades (Bartheletiomycetes, Wallemiomycetes) of the Agaricomycotina subphylum (reviewed in (Prasanna et al. 2020)). Nevertheless, all Agaricomycetes share a single, fruiting body-forming ancestor, which suggests there should be a core, deeply conserved genetic toolkit of fruiting body development. Phylogenetic comparative studies have provided sound evidence that this ancestor had corticioid/resupinate/jelly like fruiting bodies (e.g. *Phanerochaete chrysosporium*) and that more complex forms derived from such morphologies repeatedly(Hibbett 2004; Varga et al. 2019). Within the Agaricomycetes, tissues that protect and enclose basidia became thicker and more complex, eventually leading to the mushroom fruiting body structures we know today. From the perspective of this paper, it is important to mention that phylogenetic studies have also established that fruiting body evolution is highly convergent, that is, many of the characteristic traits (e.g. cap, gills) evolved independently several times. Therefore, an evolutionary framework is important for understanding how fruiting bodies evolved and for establishing homologies among tissues. Gills, caps and stipes, for example, although superficially similar, are not homologous across Polyporales and Agaricales. This implies that such, convergent forms (e.g. gills of *Lentinus tigrinus* in the Polyporales and that of *C. cinerea* in the Agaricales) are not homologous across lineages and their development should not be expected to follow the same genetic principles. On the other hand, basidia, the sexual processes that happen therein and the development of 3-dimensional fruiting body tissue, are homologous across all mushroom-forming fungi (i.e. they have a single origin in the Basidiomycota), providing an reliable framework for analysis.

These examples illustrate how the diversity of fruiting bodies can complicate the identification of core components of their morphogenesis. Ideally, we are interested in developmental processes and underlying genetic components that uniformly characterize fruiting bodies of all mushroom-forming fungi, i.e. those that can be traced back to the first fruiting body-forming Agaricomycete ancestor.

Therefore, building ‘minimal models’, which encompass the minimum set of genes necessary to build a fruiting body are highly relevant. Such minimal models could be, among others, species that produce resupinate fruiting bodies (e.g. *Ph. chrysosporium*)(Krizsán et al. 2019). Recently, *Cryptococcus neoformans* (Tremellomycetes, Basidiomycota) proved to be a useful model organism, because it produces simple lawns of basidia, sometimes referred to as fruiting bodies, in which basidium differentiation, meiosis and spore formation take place(Liu et al. 2018). Therefore, *C. neoformans* ‘fruiting bodies’ encompass all processes needed for Basidiomycota sexual reproduction but lack features essential to the definition of fruiting bodies such as three-dimensional organization, tissue differentiation and others. Merenyi et al recently sorted developmental genes of *Pleurotus ostreatus* and seven other complex species into groups that were developmentally regulated both in *C. neoformans* and mushroom-forming fungi and those in which developmental expression was observed only in mushroom-forming fungi(Merényi et al. 2021). ‘Shared’ orthogroups comprised most mitosis/meiosis related genes, but also some previously thought to be specific for complex morphogenesis (e.g. fasciclins). This indicates that sexual morphogenesis, even at the very simple forms that exist in *C. neoformans*, involves a morphogenetic program. Comparing genes that are developmentally regulated in fruiting bodies to those regulated in *C. neoformans*, thus, can help constructing more precise genetic models of complex fruiting body morphogenesis.

### 1.3. Cellular and regulatory processes involved in fruiting body development

Fruiting body development is an incredibly complex process and this complexity is reflected in the underlying cellular and genetic processes. Several cellular processes are emerging as important regulatory mechanisms underlying fruiting body morphogenesis, whereas some others that proved developmentally relevant in animals and plants seem to be of limited importance in fungi (e.g. gene body methylation(A et al. 2018; Wen et al. 2019)). This chapter summarizes the most important - mostly transcription related - processes described in fruiting bodies recently. Splicing and RNA interference are discussed below under chapters 4.5.3 and 4.6, respectively.

Probably the most influential and most commonly assayed mechanisms of fruiting body development are transcriptional reprogramming and gene expression changes. Gene expression changes naturally associate with developmental processes and are informative regarding the cellular processes that take place during fruiting. *In silico* functional analyses of differentially or developmentally expressed genes have been a primary tool for mycologists to understand fruiting body functions. Developmentally dynamic gene expression patterns form the primary data source for this paper and the main part of the paper (Chapter 4) is devoted to discussing genes with conserved expression patterns. A significant unresolved question is how gene expression changes are regulated in various stages of fruiting body development. It is not known currently if transcriptional reprogramming events (e.g. during fruiting body initiation) are mediated by broad regulation of chromatin accessibility, by regulating the expression and activity of regulatory genes (e.g. transcription factors), or a mixture of these. A potentially significant regulatory mechanism is post-translational modification of transcription factors or molecular switches in intracellular signaling pathways, as demonstrated for the *Hom2* or the *Ras1* gene of *S. commune*(Knabe et al. 2013; Pelkmans et al. 2017b). Another exciting avenue of future research is the regulatory potential of selective post-translational protein modification by the ubiquitin/proteasome system. Although direct evidence for its role in fruiting body development has not yet been published, indirect evidence was provided by the remarkable expansion of genes encoding F-box, BTB/POZ and RING-type zinc finger proteins in the genomes of mushroom-forming fungi(Krizsán et al. 2019). These gene families provide the substrate-specificity of ubiquitin ligases and have been shown, in other lineages, to be involved in transcriptional regulation(Kipreos and Pagano 2000).

**Natural antisense transcripts (NATs)** are RNA transcripts that show complementarity to transcripts of protein coding genes. NATs occur ubiquitously across eukaryotes and have been detected in mushroom-forming fungi too(Ohm et al. 2010; Muraguchi et al. 2015; Merényi et al. 2021). Because NATs do not encode proteins, they often evolve very fast and are hard to link to specific functions. Nevertheless, their expression is developmentally dynamic in all species examined to date, suggesting that they may be involved in regulating development. Muraguchi et al reported that an antisense transcript of the stipe elongation related AspE gene is transcribed just before stipe elongation(Muraguchi et al. 2015). However, it should also be kept in mind that many NATs may result from random transcription initiation and may therefore be transcriptional noise in fruiting body transcriptomes(Merényi et al. 2021).

**Allele-specific expression (ASE)** refers to the imbalance of gene expression from two divergent alleles in diploid or dikaryotic organisms. Fungal dikaryons almost always harbor two allelically different monokaryons (heterokaryons), which can lead to intermediate, A- or B-dominated expression. ASE has so far been detected in fruiting body transcriptomes of *A. bisporus*(Gehrmann et al. 2018) and *P. ostreatus*(Merényi et al. 2021), but may be widespread in the Agaricomycetes. In *A. bisporus*, Gehrmann et al suggested that ASE may facilitate the division of labor between nuclei in a dikaryon(Gehrmann et al. 2018). Another study, based on *P. ostreatus*, suggested that, at macroevolutionary timescales, ASE may be a neutrally arising phenomenon that can nevertheless generate adaptive gene expression variation(Merényi et al. 2021). It was found to be mostly characteristic of evolutionarily young genes, consistent with neutral evolution. Allele-specific expression has agriculturally broad consequences in hybrid genetics of plants(Bartoš et al. 2019) and could become a similarly central question in the mushroom industry.

**RNA editing** has recently been detected in fruiting body transcriptomes of several Ascomycota species(H et al. 2016; Liu et al. 2017; Teichert et al. 2017) and subsequently reported in Basidiomycota also(Zhu et al. 2014; Wu et al. 2019). Like alternative splicing, RNA editing provides a (post-)transcriptional mechanism for diversifying transcripts encoded by a single gene, by enzymatically modifying certain bases of the primary RNA transcript. A critical analysis recently found no evidence for RNA editing in fruiting body transcriptomes of *P. ostreatus*(Merényi et al. 2021). This study raised the possibility that, because allele-specific expression and RNA editing can generate similar signatures in RNA-Seq data, ASE may have been mistaken for RNA editing in some prior Basidiomycota analyses and that more research is needed to prove a role of RNA editing in mushroom development.

Several other exciting phenomena may be involved in the morphogenesis of Basidiomycota fruiting bodies, including regulatory processes that may yet to be discovered. For example, epigenetic regulation represents a potentially widespread but largely underexplored regulatory mechanism for fruiting body morphogenesis(Vonk and Ohm 2021). Similarly, upstream open reading frames (uORFs), which are open reading frames located in the 5’ untranslated region of an mRNA, can regulate (e.g. via encoded peptides) or interfere with the transcription of the gene(HM et al. 2009). uORFs are known to be important in the asexual development of Ascomycota(Han and Adams 2001) [and have recently been detected in developmental genes of *C. cinerea* (Hegedus et al in prep)].

### 1.4. Genomic and transcriptomic resources

Genomic resources are also available for most model species of fruiting body development (e.g. *C. cinerea* (Schaeff.) Redhead, Vilgalys & Moncalvo*, S. commune* Fr.*, Cyclocybe aegerita* (V. Brig.) Vizzini)(Ohm et al. 2010; Stajich et al. 2010; Muraguchi et al. 2015; Gupta et al. 2018; Almási et al. 2019) and those used by the mushroom industry, including *Hericium erinaceus* (Bull.) Pers.(Gong et al. 2020; W et al. 2020)*, Ganoderma spp.*(Chen et al. 2012)*, Flammulina spp.*(Park et al. 2014, 2019; Chen et al. 2020)*, L. edodes* (Berk.) Pegler(Chen et al. 2016; Shim et al. 2016; Sakamoto et al. 2017a)*, Pleurotus spp*(A et al. 2018; Wang et al. 2018a; Dai et al. 2019; Lee et al. 2021)*, Agaricus spp*(Morin et al. 2012; O’Connor et al. 2019; Sonnenberg et al. 2020)*, Tricholoma matsutake* (S. Ito & S. Imai) Singer(Min et al. 2020)*, Sparassis crispa* (Wulfen) Fr.(Kiyama et al. 2018)*, Hypsizygus marmoreus* (Peck) H.E. Bigelow(Min et al. 2018), among others. More broadly, in the Agaricomycetes, draft genome sequences have been proliferating at a steady pace, due to interest in biodiversity, lignocellulose decomposition(Martinez et al. 2004, 2009a; Floudas et al. 2012; Riley et al. 2014) and mycorrhiza-formation(Martin et al. 2008; Kohler et al. 2015; Miyauchi et al. 2020). This trend has virtually eliminated genome data being the bottleneck in biological discovery in fungi. This is further supported by the spread of third generation sequencing techniques, long-read sequencing data, which allow more and more complete (telomere-to-telomere) assemblies and a range of analyses that draft genomes did not permit.

The continuous development of technologies has also influenced the identification of genes related to fruiting body morphogenesis. While early studies isolated fruiting body-specific proteins/genes using mutant analyses, hybridization techniques (among others), these were later substituted by RT-PCR and microarray studies, then by high throughput sequencing analyses, such as 5’-serial analysis of gene expression (SAGE(WW et al. 2008; Cheng et al. 2013a)) and most recently RNA-Seq. As of today, RNA-Seq datasets are available for a wide range of species and conditions. These include *C. cinerea*(Plaza et al. 2014a; Muraguchi et al. 2015; Krizsán et al. 2019; Xie et al. 2020)*, Flammulina spp*(Yan et al. 2019; Liu et al. 2020b)*, L. edodes*(Park et al. 2017; Sakamoto et al. 2017b; Song et al. 2018b; Y et al. 2018; Yoo et al. 2019; Kim et al. 2020)*, Agaricus spp*(Gehrmann et al. 2018; Lu et al. 2020; O’Connor et al. 2021)*, Pleurotus ostreatus* (Jacq.) P. Kumm.(Wen et al. 2019; Merényi et al. 2021)*, H. marmoreus*(Zhang et al. 2015b)*, C. aegerita*(Orban et al. 2021)*, Phanerochaete chrysosporium* Burds.(Krizsán et al. 2019)*, Lentinus tigrinus* (Bull.) Fr.(Krizsán et al. 2019)*, Mycena kentingensis* Y.S. Shih, Chi Y. Chen, W.W. Lin & H.W. Kao(Ke et al. 2020a)*, Rickenella mellea* (Singer & Clémençon) Lamoure(Krizsán et al. 2019)*, Armillaria ostoyae (*Romagn.) Herink(Sipos et al. 2017a)*, Pisolithus microcarpus* (Cooke & Massee) G. Cunn (Pereira et al. 2017)*, Laccaria bicolor* (Maire) P.D. Orton (Ruytinx et al in prep)*, S. commune*(Ohm et al. 2010) and *Auriculariopsis ampla* (Lév.) Maire(Almási et al. 2019). Many of these datasets build the foundation of this study, by providing windows into developmentally expressed genes in different fungal species.

A general observation we made during the analyses we present hereafter is that the resolution of the transcriptomic data is key to identifying expression dynamics and patterns. Both tissue-wise and temporal resolution of the data allowed the discovery of more developmentally relevant expression patterns. We found several cases where tissue-resolved transcriptomes provided clear signal for conserved, tissue-specific expression of a gene, whereas in non-resolved transcriptomes the same gene showed a more or less flat expression curve. This is an inherent nature of bulk RNA-Seq data: spatial expression patterns average out in samples that contain large populations of multiple cell types.

Beyond gene expression, a broad array of other types of information could be leveraged for understanding fruiting body development. Notable recent examples include assays of the overrepresentation of certain gene families in the genomes of focal species(Krizsán et al. 2019), a population genomics study combined with selection analyses and gene expression profiling in *L. edodes*(Zhang et al. 2021c) to identify genes involved in fruiting body development(Zhang et al. 2021c), or ChIP-Seq analyses for assaying promoter occupancy and thus transcriptional activity(Vonk and Ohm 2021).

## 2. Description of the approach used in this paper

In this study, we address the puzzle of how a fruiting body develops and what novel insight can be gleaned from systematic comparisons of developmental transcriptomes. We aim at providing a comprehensive listing of putatively development-related gene families in the Agaricomycetes. We provide, for each discussed functional group or gene family, information on their expression, orthology to well-established model organisms (*Saccharomyces cerevisiae, Schizosaccharomyces pombe, Aspergillus nidulans*, *Neurospora crassa*) and to key mushroom-forming fungi as tables. We also annotate several conserved pathways in *C. cinerea*, based on strict 1-to-1 orthology to known pathway members in well-researched model systems (see Methods for details).

Each chapter is structured so that it starts with a literature review of the given gene family followed by a discussion of novel findings (if any) based on our meta-analysis of transcriptomes. Our main units of investigation are **conserved developmentally expressed (CDE)** orthogroups, which are groups of single-copy genes in which the majority (>65%^1^) of the genes were developmentally expressed (see Methods and Merenyi et al 2021(Merényi et al. 2021)). For simplicity CDE orthogroups are considered to be groups of genes sharing similar functions (see limitations below) in fruiting body development.

We performed a meta-analysis of 12 developmental transcriptomes (*Agaricus bisporus*(Gehrmann et al. 2018)*, C. aegerita*(Orban et al. 2021)*, A. ostoyae*(Sipos et al. 2017a)*, A. ampla*(Almási et al. 2019)*, C. cinerea*(Krizsán et al. 2019)*, L. bicolor* (Ruytinx et al in prep)*, L. edodes*(Zhang et al. 2021d)*, L. tigrinus*(Krizsán et al. 2019)*, M. kentingensis*(Ke et al. 2020a)*, Ph. chrysosporium*(Krizsán et al. 2019)*, P. ostreatus*(Krizsán et al. 2019)*, S. commune*(Almási et al. 2019)) (Fig. 1). These species belong to two orders (Agaricales, Polyporales), which span an estimated 200 million years of evolution(Varga et al. 2019), and represent diverse ecologies and morphological adaptations, such as simple resupinate (*Ph. chrysosporium*), coralloid (*Pterula gracilis*), pileate-stipitate (e.g. *C. cinerea*) and cyphelloid (*A. ampla, S. commune*) forms. Each of these species and their ecological adaptations show several unique aspects (e.g. fruiting body ecotypes) which could be related to gene expression data. However, we are here interested in only the shared aspects of fruiting body development, and therefore use the diversity of the compared species in focusing our attention to widely conserved fruiting body genes which should help define the core building blocks required for fruiting body morphogenesis.

The comparative approach we follow is paramount to separate ‘wheat from chaff’, i.e. developmentally regulated genes that are relevant for the development of fruiting bodies in general from genes with taxonomically restricted or species-specific developmental roles. This builds on the assumption that a gene involved in core fruiting body functions should be conserved across Agaricomycetes and should show similar expression dynamics in all species. Therefore, in our approach we focus on conservation of both sequence and expression. A comparative approach is also desirable because it is now well-known that gene expression is associated with ’biological noise’, which may influence the detection of differentially expressed or developmentally regulated genes(Merényi et al. 2021) in any single species, whereas across-species comparisons effectively cancel the effects of gene expression noise.

This study, like all others, has limitations. First, it should be noted that evidence for gene function remains, in an overwhelming majority of cases, circumstantial, inferred from expression patterns or extrapolated from the functions of orthologous genes in model fungi. At the moment we rely on these types of evidence due to the scarcity of developmental genetic and in-depth functional studies in most Agaricomycetes. Despite the inferential nature of the study, many orthologs display very conserved expression patterns across Agaricomycetes species (e.g. aquaporins in stipe tissues, see below), allowing us to make confident predictions on the function of such genes. Second, a perfect comparative transcriptomic study would compare corresponding developmental stages across species. In the Agaricomycetes establishing homology relationships among individual species is complicated (see above). Furthermore, the developmental transcriptomes we used differ in the sampled time points and resolution (e.g. tissue-wise or bulk) and do not follow a unified nomenclature for developmental stages (e.g. aggregate in *S. commune* vs. hyphal knot in *C. cinerea*). Therefore, we here focused broadly on comparing expression dynamics and the shape of the expression curve, rather than attempting to find 1-to-1 correspondence between stages of different species. Finally, in the present study, we focus on genes with developmentally dynamic expression. However, not all morphogenetically relevant genes show expression dynamics; constantly expressed genes can often be important for development, for example, if regulated post-transcriptionally (e.g. alternative splicing isoforms(Gehrmann et al. 2016; Krizsán et al. 2019)) or post-translationally (e.g. phosphorylation, ubiquitylation etc.(Knabe et al. 2013; Pelkmans et al. 2017a)).

## 3. Methods

### Bioinformatic analyses of RNA-Seq data

The raw RNA-Seq data of 12 previously published Basidiomycota species were reanalysed following Merenyi et al(Merényi et al. 2021). The following reference genomes were used: *C. cinerea* (AmutBmut pab1-1 v1.0(Muraguchi et al. 2015))*, A. ostoyae* (C18/9(Sipos et al. 2017b)), *A. ampla* (NL-1724 v1.0(Almási et al. 2019)), *S. commune* (H4-8a and H4-8b(Ohm et al. 2010)), *L. bicolor* (v2.0(Martin et al. 2008))*, L. edodes* (Le(Bin) 0899 ss11 v1.0(Zhang et al. 2021c))*, L. tigrinus* (RLP-9953-sp(Wu et al. 2018) ), *Ph. chrysosporium* (RP-78 v2.2(Ohm et al. 2014)), *P. ostreatus* (PC15 v2.0(Riley et al. 2014))*, M. kentingensis*(Ke et al. 2020a)*, A. bisporus* (var. bisporus H97(Morin et al. 2012))*, C. aegerita* (AAE-3(Gupta et al. 2018)). To remove adaptors, ambiguous nucleotides and any low quality read ends, reads were trimmed using bbduk.sh and overlapping read pairs were merged with bbmerge.sh (part of BBMap/BBTools; http://sourceforge.net/projects/bbmap/) with the following parameters: qtrim=rl trimq=25 minlen=40. A two-pass STAR alignment(Veeneman et al. 2016) was performed against reference genomes with the same parameters as in our previous study(Krizsán et al. 2019) except that the maximal intron length was reduced to 3000 nt. Reads were counted and summarized for each exon with FeatureCounts(Liao et al. 2014) taking into account the mode of sequencing (paired-end, single-end). Since the *Cyclocybe* transcriptome was obtained with the Quantseq method, reads were counted only in the -100 - +400 region of the 3’ ends of the genes. Read count data were normalized using EdgeR(Robinson et al. 2010). Expression levels were calculated as fragments per kilobase of transcript per million mapped reads (FPKM).

Developmentally regulated genes were identified as before(Krizsán et al. 2019; Merényi et al. 2021), based on a stricter four-fold or a more permissive two-fold change cutoff between adjacent developmental stages or tissue types. Genes showing a 4-fold upregulation in the first primordium stage relative to vegetative mycelium were termed ‘FB-init’, and considered regulated during fruiting body initiation. It should be noted that the data we analyze is noisy, based on different chemistries, different labs and protocols, etc., which introduces a certain amount of data loss in our analyses. Nevertheless, the transition to fruiting body development is such a dramatic change that we expect similar gene expression changes to be discoverable across species, despite the heterogeneity of the data. This is supported by previous comparative transcriptomic studies(Plaza et al. 2014b; Krizsán et al. 2019; Merényi et al. 2020b, 2021) as well as the biologically meaningful conclusions obtained here.

### Orthology based on reciprocal best hits

Proteins of each species were searched against the proteomes of other species using the reciprocal best hit (RBH) module of MMSeqs2(Steinegger and Söding 2017). To remove spurious RBHs, a bidirectional 50% coverage and at least 1e-6 e-value were required. Proteins were clustered into single copy clusters using a connected component clustering algorithm in the igraph package([CSL STYLE ERROR: reference with no printed form.]), based on reciprocal best hits. This step clustered 74.3% of the proteins in the examined species. The other 25.7% of proteins form clusters with duplication in any or more species. For these 1:1 orthogroups were circumscribed using the ‘ortholog coding’ algorithm of the COMPARE pipeline(Nagy et al. 2014). In the case of terminal duplications, orthogroup membership was decided based on connectivity; the protein which showed more hits to other members of the orthogroup was selected while the other(s) moved to separate orthogroup(s). Complete orthology relationships of genes across all 12 species are given in this paper are presented in Supplementary Table 1.

This approach is suitable for identifying moderately to very conserved gene groups, whereas for highly volatile genes (i.e those that show frequent duplication/loss), a more liberal approach building on orthologous groups (like(Krizsán et al. 2019)) is more appropriate. For fast-evolving genes, approaches looking at sequence conservation might not work. Therefore, in the case of certain gene families, such as defense genes or cell wall remodeling CAZymes, we relied on primary gene family classification instead of reciprocal best hit based orthogroups. In the case of CAZymes, we followed the cazy.org classification(Levasseur et al. 2013).

### Identification of CDE orthogroups

The central unit of discussion in this paper is referred to as ‘conserved developmentally expressed (CDE) orthogroup’, which refers to groups of single-copy genes (one gene per species) in which the majority of genes is developmentally regulated either at fold change >2 or >4. As a principal rule, we considered an orthogroup to be a CDE orthogroup if >65% of the genes in it were developmentally regulated, and at least 8 species were represented. Deviations from this rule were allowed in unique cases (e.g. taxonomically restricted but important gene family) and are transparently presented in tables and supplementary tables under each of the following chapters.

## 4. Results: developmentally expressed gene classes in the Agaricomycetes

In this paper we synthesized literature reviews with a meta-analysis of developmental transcriptomes to identify conserved gene families involved in fruiting body development. For the latter we reanalysed data for 12 species and identified developmentally regulated genes, i.e. those that showed >2 or >4-fold changes between successive developmental stages or adjacent tissue types (Supplementary Table 2). Statistics on developmentally expressed genes are provided in Supplementary Table 2/b. Numbers of developmentally regulated genes were similar to those reported by previous studies,(Almási et al. 2019; Krizsán et al. 2019; Merényi et al. 2020b, 2021) therefore, we here do not discuss these in detail, rather focus on those that are conserved across species.

We organized developmentally regulated genes into strict 1-to-1 orthogroups following the logic of Merenyi et al(Merényi et al. 2021). Orthogroups in which >65% of the genes were developmentally regulated were designated as conserved developmentally expressed (CDE) orthogroups at fold change values of >2 or >4. We identified 921 CDE orthogroups across the twelve Agaricomycetes species; these form the basis of the discussion presented in the rest of this paper. Statistics on the number of species which had developmentally regulated genes in these orthogroups are shown on Fig. 2/A. The transcriptome data of *A. bisporus* was an outlier in several aspects. We currently do not know if this is due to biological differences of this species or technical reasons, nevertheless this transcriptome was assigned lower weight and discussed rarely in the following chapters. By relying on conserved developmentally expressed orthogroups, we here take shared expression dynamics as a proxy for conservation of function. It should be mentioned that this approach may be, in several cases, only a crude approximation of real conservation of function, but facilitates comparative discussion of gene families and their speculated functions in fruiting body morphogenesis.

**Fig. 2.**
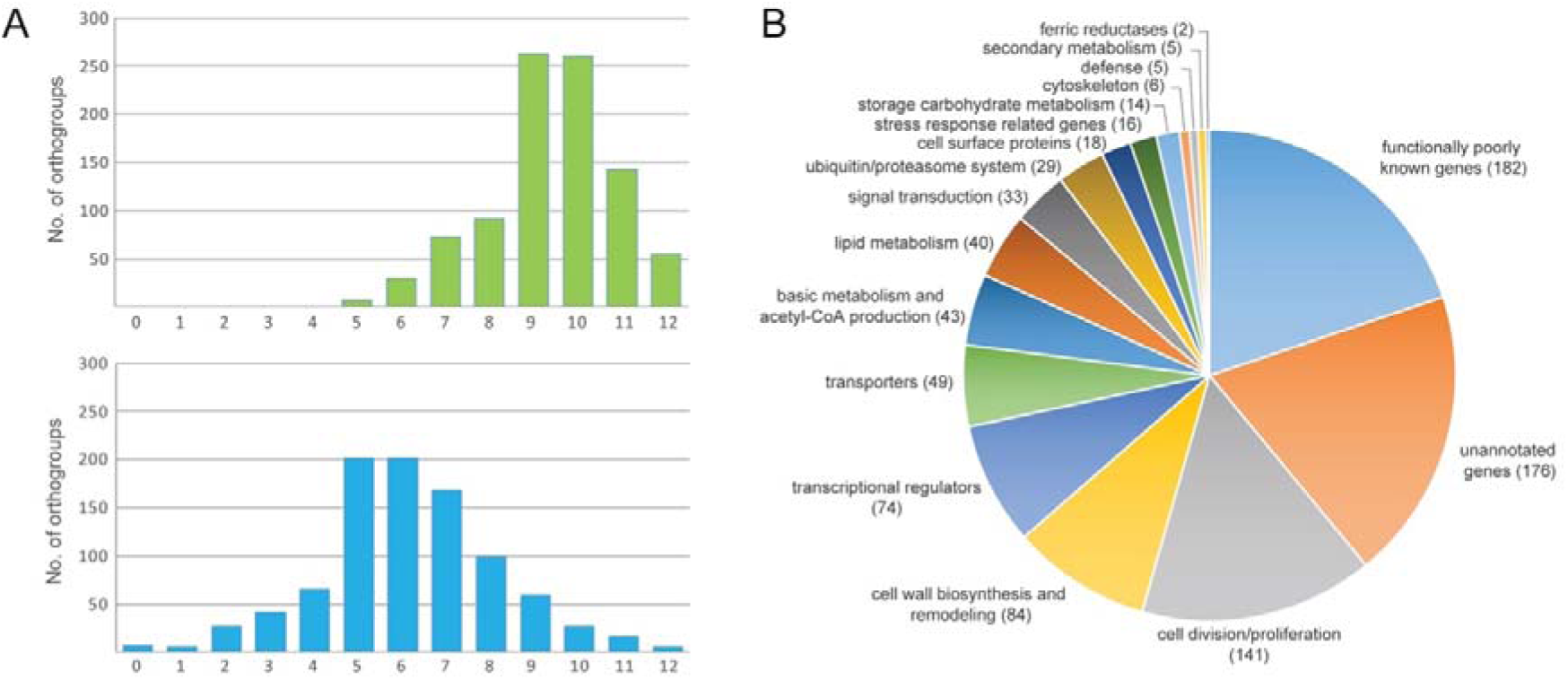
The distribution of conserved developmentally regulated orthogroups across species. A The number of orthogroups as a function of the number of species in which the orthogroup is developmentally regulated at fold change >2 (top) and fold change >4 (bottom). Horizontal axis represent the number of species. B Numbers of genes in each of the functional categories discussed in the paper.

We functionally characterized CDE orthogroups and arranged them into broader groups based on function inferred by orthology to genes in well-known model systems (*S. cerevisiae, Schizosaccharomyces pombe, A. nidulans, N. crassa*) and/or their Gene ontology and InterPro annotations. We defined 17 broader functional groups, cell division/proliferation, defense, transcriptional regulators, signal transduction, cell wall biosynthesis and remodeling, cell surface proteins, secondary metabolism, cytoskeleton, basic metabolism and acetyl-CoA production, lipid metabolism, storage carbohydrate metabolism, ubiquitin/proteasome system, transporters, functionally poorly known genes, unannotated genes, stress response related genes and ferric reductases (Fig. 2/B). The main categories were further subdivided into finer groups to facilitate discussions of putative functions. We categorized developmental genes manually to the best of our knowledge based on inferred gene function, though the subdivision is probably not impeccable. For example, overlaps exist between chromatin remodeling and DNA replication related regulatory factors or between the latter and cytoskeleton related genes (e.g. kinesins involved in moving chromatids during mitosis).

The two largest categories were functionally poorly characterized and unannotated genes, these contained 182 and 176 CDE orthogroups, respectively (Fig. 2/B). The first, ‘functionally poorly characterized genes’ contains orthogroups which we could not confidently link to any fruiting body functions. It is possible that this category could be subdivided and with further manual annotation more orthogroups could be explained in the context of fruiting body development. On the other hand, unannotated genes are those that contain no existing Pfam or InterPro domain signatures and lack all kinds of functional annotation. The largest functional group for which a function could be determined was related to cell proliferation (DNA replication, DNA repair, mitosis, meiosis) followed by cell wall biosynthesis and remodeling. Transcriptional regulators (including transcription factors, chromatin-related and RNA-binding regulatory proteins), transporters and Acetyl-CoA production related genes were also represented by a considerable number of genes. Certain functional categories, such as ‘defense’ and ‘secondary metabolism’ contained only a small number of CDE orthogroups. In these cases, these low numbers reflect the lack of conservation of gene sequences, rather than the lack of a role of these gene families in fruiting bodies.

Our discussion is guided here by conservation of developmentally regulated families. However, there are gene families that do not form conserved orthogroups, but are nevertheless very important for fruiting body development. These can be either multigene families that undergo frequent duplication and thus orthology relationships are intricate (e.g. hydrophobins), or fast-evolving families in which divergence quickly erases orthology relationships (e.g. F-Box proteins, defense-related proteins). Therefore, in addition to gene families/pathways identified through CDE orthogroups, we discuss broader, fast-evolving gene families based on annotations at the gene family level only (i.e. without orthogroups) and performed manual annotations to identify broader containing gene families or cellular pathways. Accordingly, in addition to the 921 CDE orthogroups, we discuss further 558 gene groups which belong to broad functionalities involved in fruiting body development. For some processes we were able to reconstruct (nearly) complete pathways, such as in the case of mitosis/meiosis related genes or fatty acid biosynthesis, whereas in other cases we only document the widespread developmental regulation and expression profile conservation of a gene family, but without information on the containing pathway of broader cellular function. For some gene groups, such as cytoskeletal genes, GPCRs or MAP kinase pathways, we provide annotations despite their limited expression dynamics during fruiting body development. Such gene families may be important in fruiting body development despite their low expression dynamics, therefore, we decided to annotate them in order to facilitate comparative discussion across mushroom-forming fungi and more thoroughly investigated Ascomycota model systems.

In the following chapters we discuss each of the more important gene groups in detail, with emphasis on expression profiles, putative function and role in fruiting body development. Besides well-known ones, we intentionally focus on lesser-known gene families with widespread developmental expression, to potentially distill new insights into fruiting body development. We provide catalogues and references to recent reviews for well-researched gene families as well (e.g. hydrophobins or laccases).

### 4.1. Cell division, proliferation and growth

#### 4.1.1. Meiotic and mitotic gene expression show two distinct expression peaks

Mitosis and meiosis are key to the development and growth of multicellular organisms. Mushroom fruiting bodies are no exception: mitosis leads to cell proliferation early in fruiting body development, whereas meiosis happens in basidia, which are localized to gills, to produce spores.

Transcriptomic data provided clear evidence for expression peaks of genes related to mitotic or meiotic cell division, as well as associated processes in DNA replication, repair, chromosome dynamics/movement etc., during fruiting body development. The enrichment of meiotic/mitotic genes during fruiting body development and sporulation was noted in *C. cinerea*(Burns et al. 2010; Muraguchi et al. 2015; X et al. 2015; Krizsán et al. 2019), *Pisolithus microcarpus*(Pereira et al. 2017)*, Tricholoma matsutake*(Tang et al. 2020) and *L. edodes*(Song et al. 2018b), *Agaricus blazei*(Lu et al. 2020) as well as in a six-species comparison published by Krizsan et al(Krizsán et al. 2019), among others. Similar signal for meiotic gene upregulation has been detected in Ascomycota fruiting bodies as well(Rodenburg et al. 2018). *C. cinerea* has been used as a model system of Basidiomycota meiosis, in particular in so called white cap mutants that have meiosis-associated defects(Lu et al. 2003). These works revealed many conserved and some unique aspects of the meiotic process as well as resulted in meiosis being one of the best understood processes in this species(Seitz et al. 1996; NY et al. 1997; L et al. 1999; Gerecke and Zolan 2000; Nara et al. 2001; Cummings et al. 2002; Lu et al. 2003; SH et al. 2004; Muraguchi et al. 2008a; AM et al. 2009).

Probably the most detailed such study was performed by Burns et al(Burns et al. 2010), who investigated by microarray the transcriptional events that happen in *C. cinerea* gills during a time course that encompasses the meiotic events. They found that 2,721 genes exhibited changing probe intensity during the six time points around meiosis. Using dikaryotic mycelium as reference, they identified 886 genes that were expressed in gill tissue only; these genes included core meiotic components. Like in other species, genes were induced in successive waves during the meiotic process; these waves were arranged into 9 clusters based on expression trajectories. The clusters separated genes into functional groups, e.g. those required for prophase I or sporulation in early and late in the time series, respectively. Early genes further included ones related to DNA replication, cytoskeleton organization and regulation. Late induced genes in their study are mostly related to spore formation and show overlaps with genes related to ascospore formation in *S. cerevisiae*(S et al. 1998a; Primig et al. 2000) and *Sch. pombe*(Mata et al. 2002). Ribosomal protein coding gene expression was found to be high up to karyogamy, at which point it starts to decrease and does not lift up again. This might reflect the shutting down of protein synthesis as the fruiting body enters the sporulating phase.

Freitas Pereira et al examined sporogenesis in *Pisolithus*, an ectomycorrhizal fungus in the Boletales, and found that meiotic genes (annotated by KEGG) showed dynamics throughout the development of basidiospore-containing peridioles(Pereira et al. 2017). In this species spores are produced in peridioles, lentil-shaped to globular compartments within the fruiting body. Several genes that were differentially expressed in peridioles of different maturation stages were annotated as meiotic/cell cycle or cell division related, most of which showed an expression peak in unconsolidated, young, and mature peridioles. These observations are consistent with microscopic observations on the sporogenesis of *P. microsporus*(AN and MD 2010).

We annotated 180 conserved orthogroups as involved in cell proliferation, such as mitosis, meiosis, DNA replication, DNA repair or the cell cycle, based on expression conservation 141 of these qualified as CDE orthogroup (Table 1). These included previously functionally characterized genes such as *ku70*, *rad50* and *Mre11* of *C. cinerea*(AM et al. 2009; Nakazawa et al. 2011), DMC1 (=Lenedo1_1211634) of *L. edodes*(Sakamoto et al. 2009)*, ku80* of *S. commune*(De Jong et al. 2010) and *Msh4* of *P. ostreatus*(Lavrijssen et al. 2020). Most of these genes showed developmental regulation and similar expression profiles in most of the species. This is not surprising, given that both mitosis and meiosis have well-documented roles in sculpting fruiting bodies. Nevertheless, to our knowledge, these genes represent the most comprehensive, although surely not complete, list of cell proliferation related genes to date. We tentatively subdivided CDE orthogroups into ones related to DNA replication, DNA repair, meiosis as well as both mitosis and meiosis; these contained 87, 26, 13, 54 genes, respectively (Table 1). These include members of several well-characterized protein complexes described from S. cerevisiae, such as Ku, SMC, MCM, Ndc80, Gins and cohesin complexes (Table 1). Orthologs of *C. cinerea* arp9, which was described as a putative actin-related protein involved in chromatin remodeling(Nakazawa et al. 2016), were also found to follow similar expression profiles, although at moderate dynamics (Table 1).

**Table 1.**
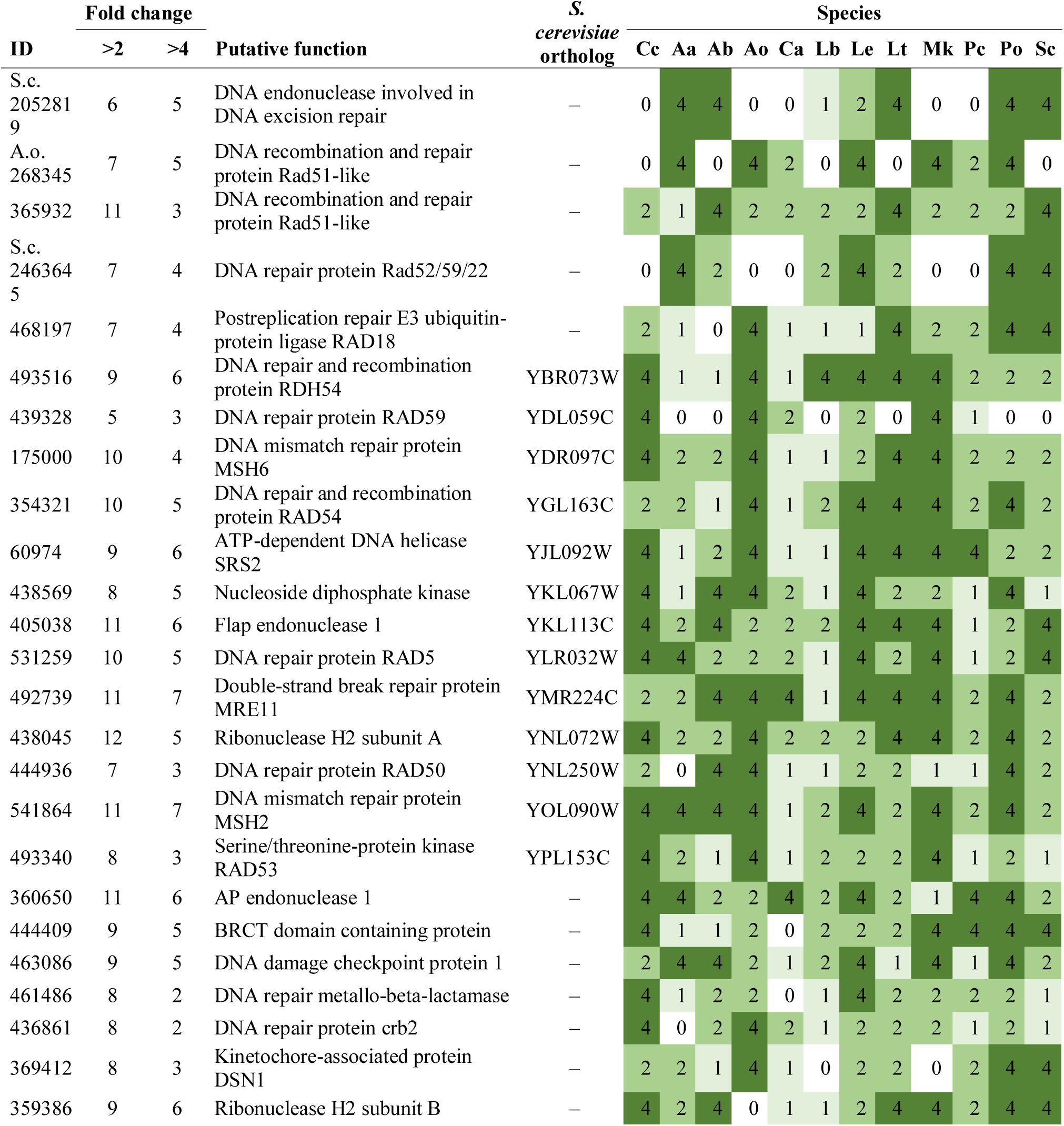
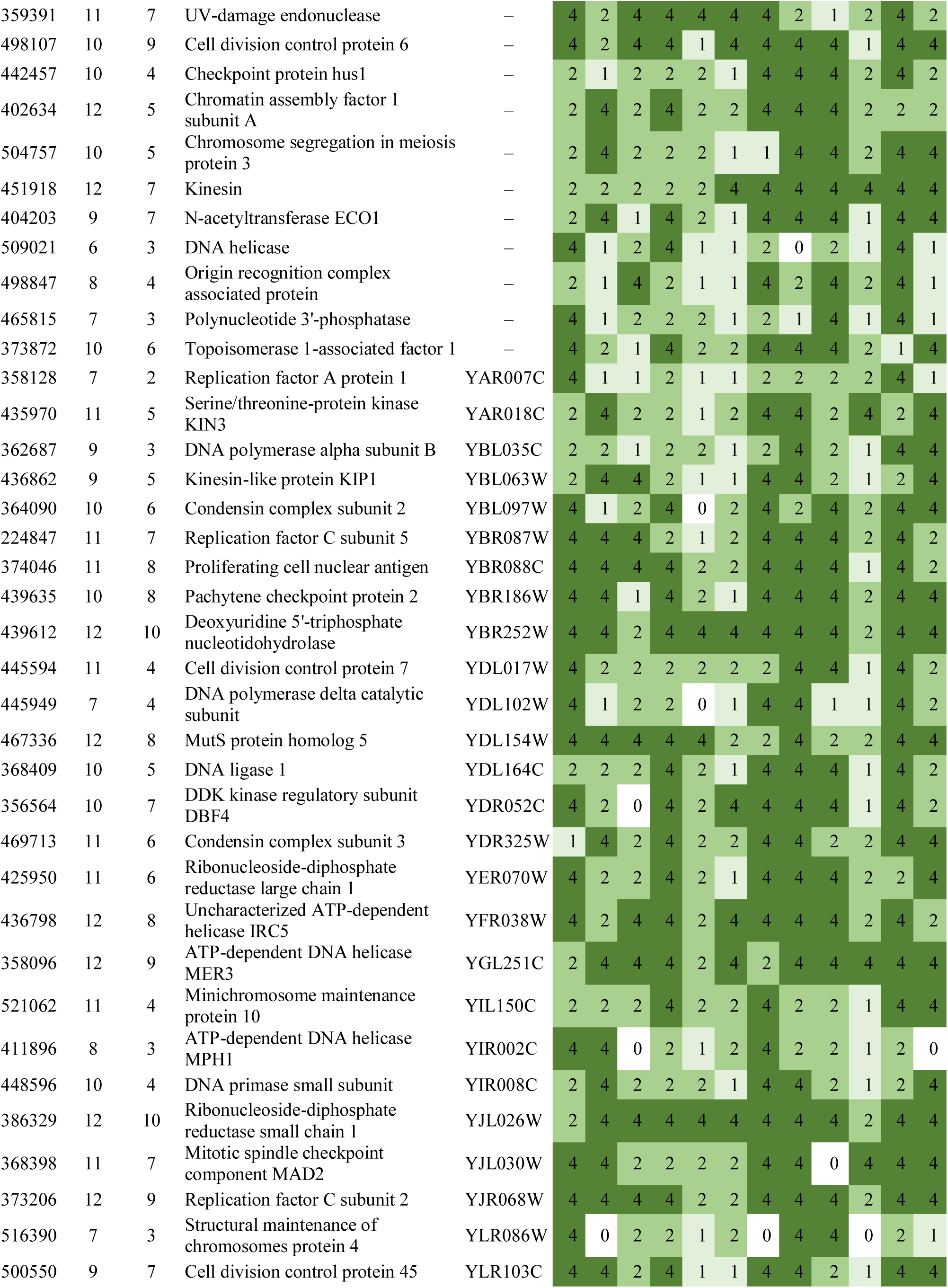
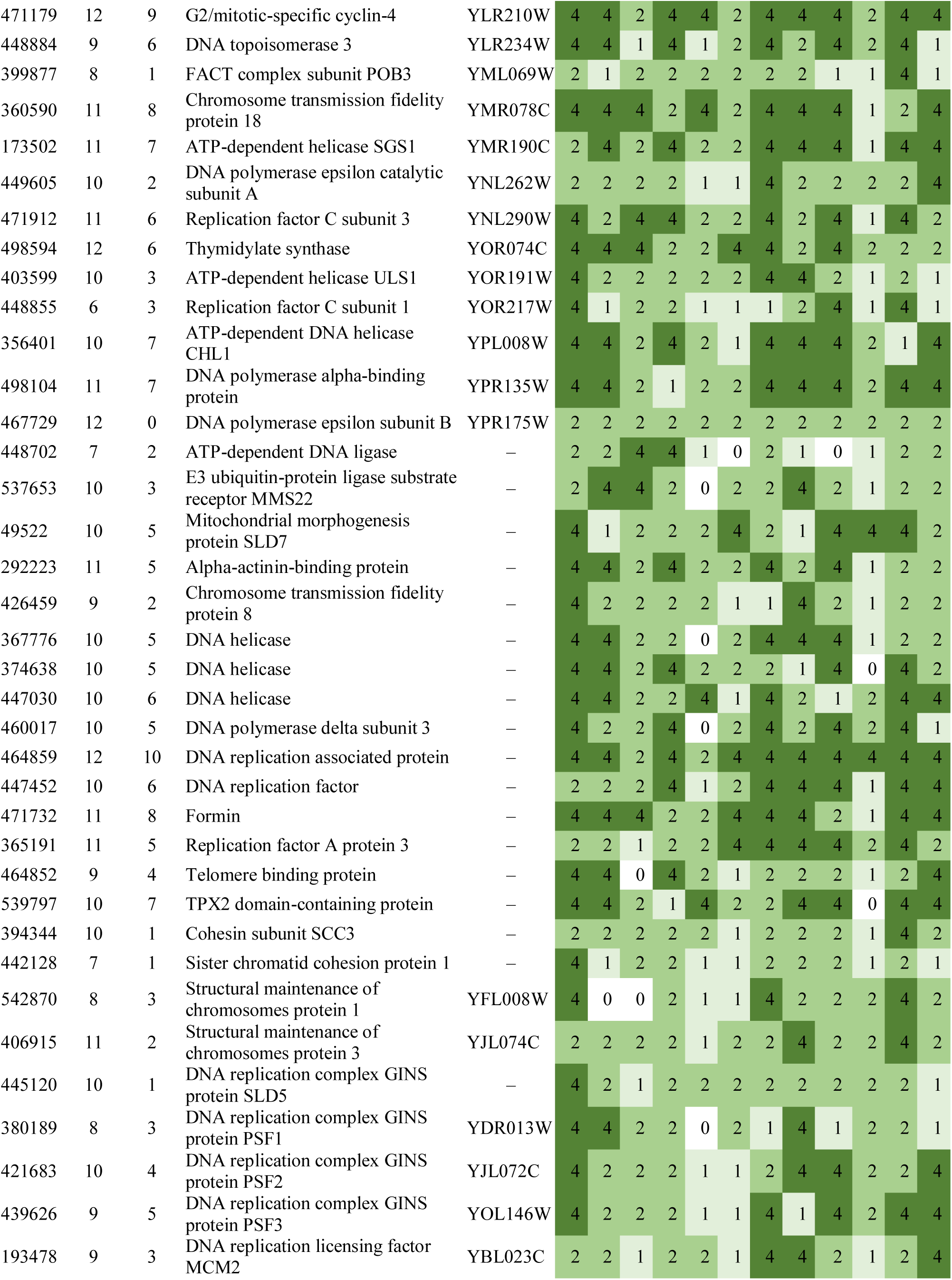
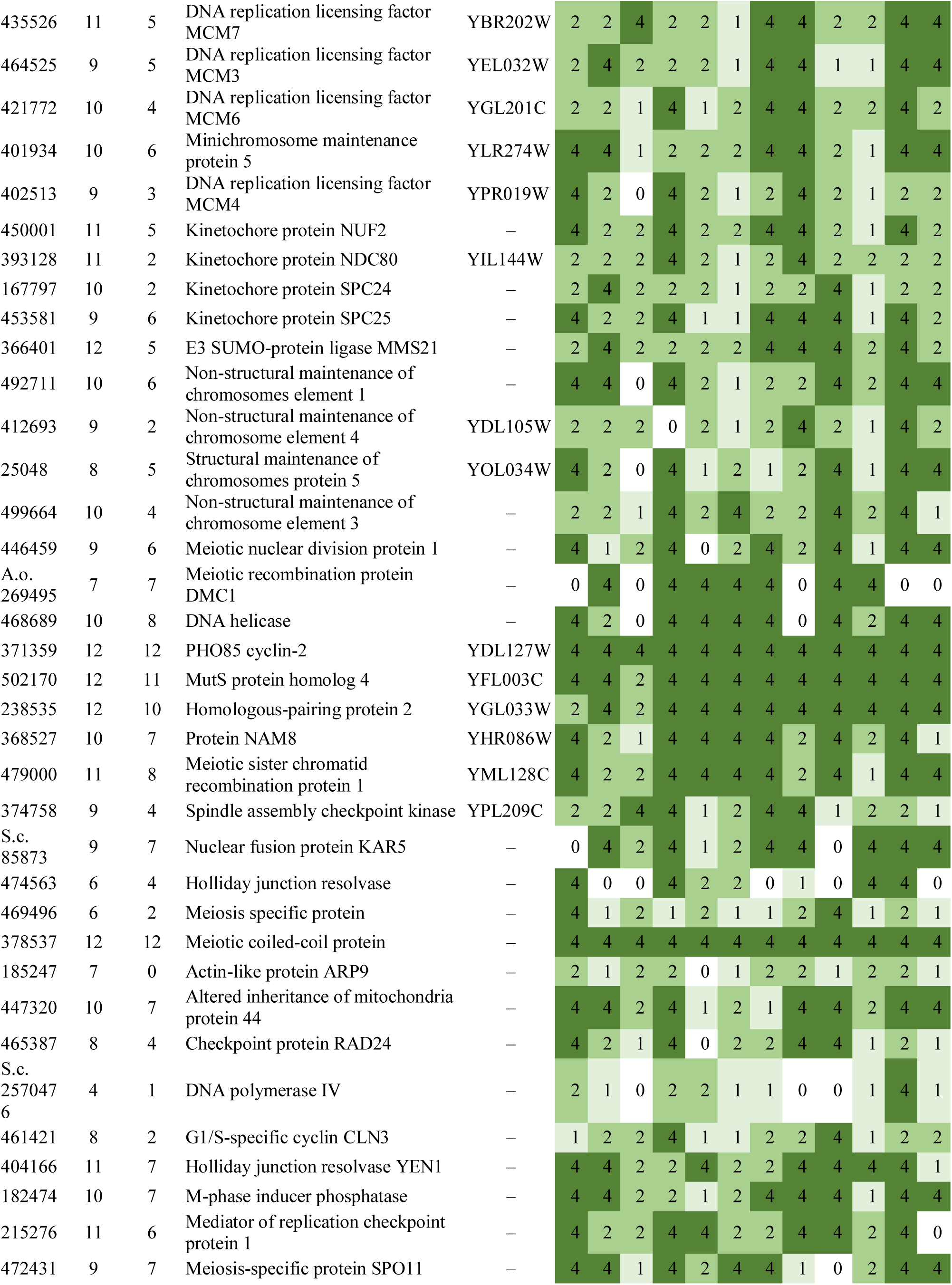
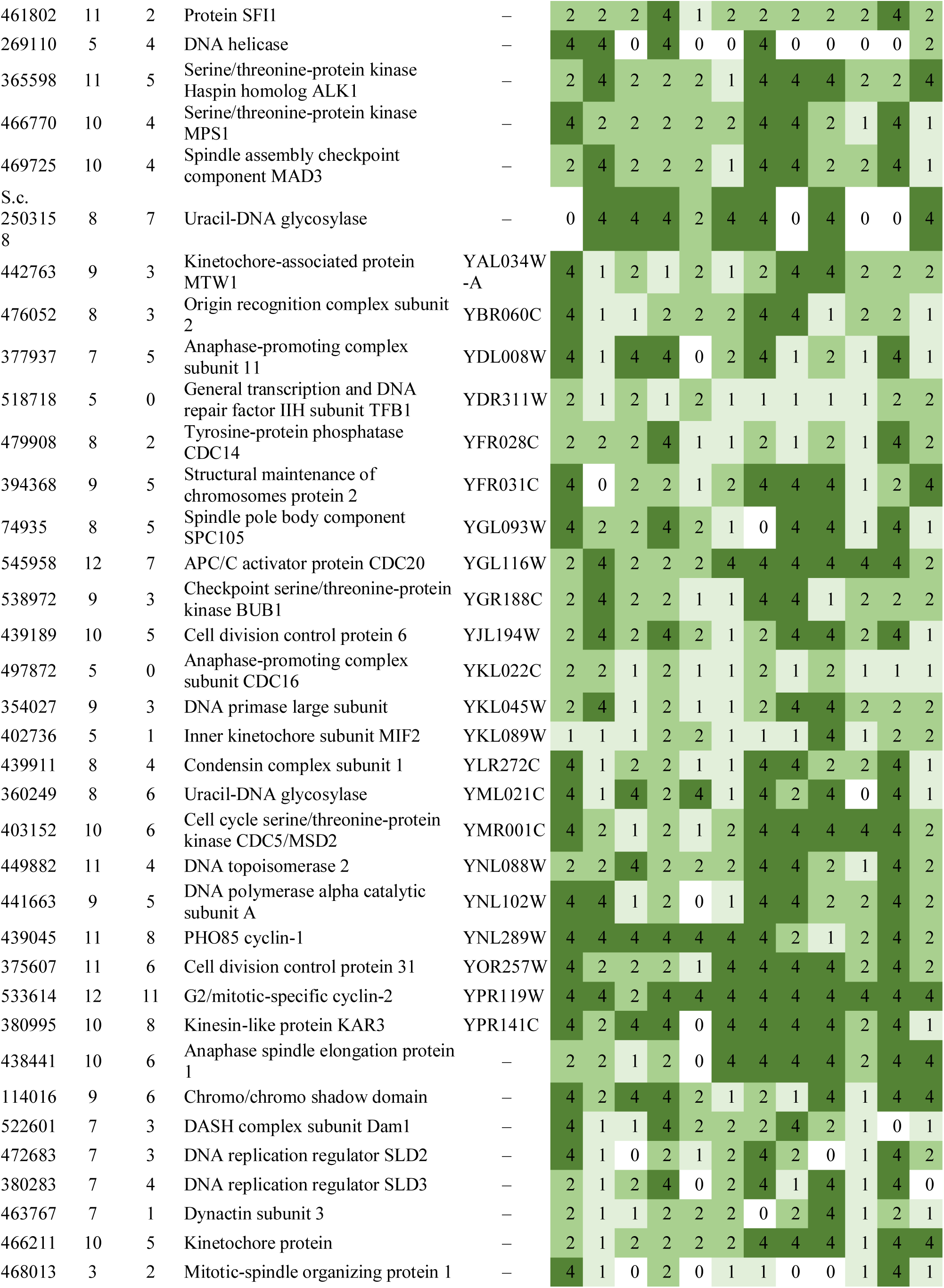
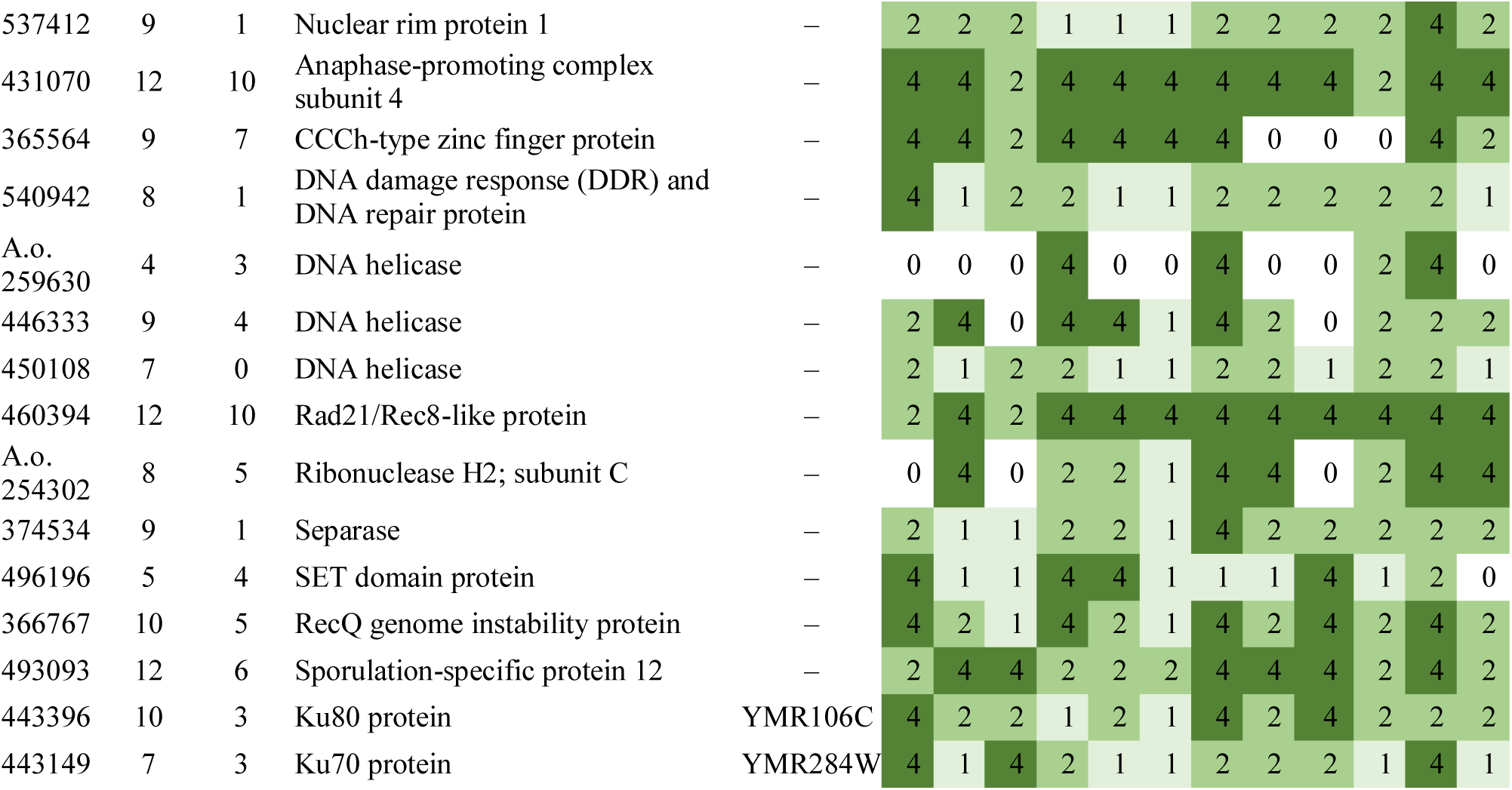
Summary of developmental expression dynamics of CDE orthogroups of cell division-related genes across 12 species. Protein ID of a representative protein is given follows by the number of species in which the orthogroup is developmentally regulated at fold change 2 and 4 (FC>2 and FC>4, respectively). Putative function and ortholog in *S. cerevisiae* (if any) are also given. Abbreviations: 0-gene absent, 1-gene present but not developmentally regulated, 2 - developmentally regulated at fold change >2, 4- developmentally regulated at fold change >4. Species names are abbreviated as: Cc – *C. cinerea*, Aa – *A. ampla*, Ab – *A. bisporus*, Ao – *A. ostoyae*, Ca – *C. aegerita*, Lb – *L. bicolor*, Le – *L. edodes*, Lt – *L. tigrinus*, Mk – *M. kentingensis*, Pc – *Ph. chrysosporium*, Po – *P. ostreatus*, Sc – *S. commune*.

These genes showed characteristic expression profiles in most species. The expression of meiotic genes shows a distinct peak in gill tissues (Fig. 3, Supplementary Fig. 1) or in stages that contain meiotic tissue. Meiotic gene expression precedes early phases of sporulation and was clearly discernible in *C. neoformans* or *P. ostreatus* in previous transcriptomic studies(Liu et al. 2018; Merényi et al. 2021). Because the onset of meiosis is tightly regulated in most species (among mushrooms best known in *C. cinerea*(Kües 2000)) meiotic gene expression also provides a landmark to calibrate developmental chronologies among species where this may be hard to do because of morphological divergence. It should be noted that some species develop in ways so that we do not expect a clear peak. For example, cell proliferation and spore production is not synchronized in *S. commune*, where it appears that sporulation progresses from the basal, older parts and progresses towards the edges of the fruiting body(Kües and Liu 2000). In *S. commune*, this may be explained by the highly derived morphology of the fruiting bodies, consisting of an assemblage of cyphelloid fruiting bodies. Therefore, we would not expect one clear peak of meiotic gene expression, unless we sample different zones of the fruiting body separately.

**Fig. 3.**
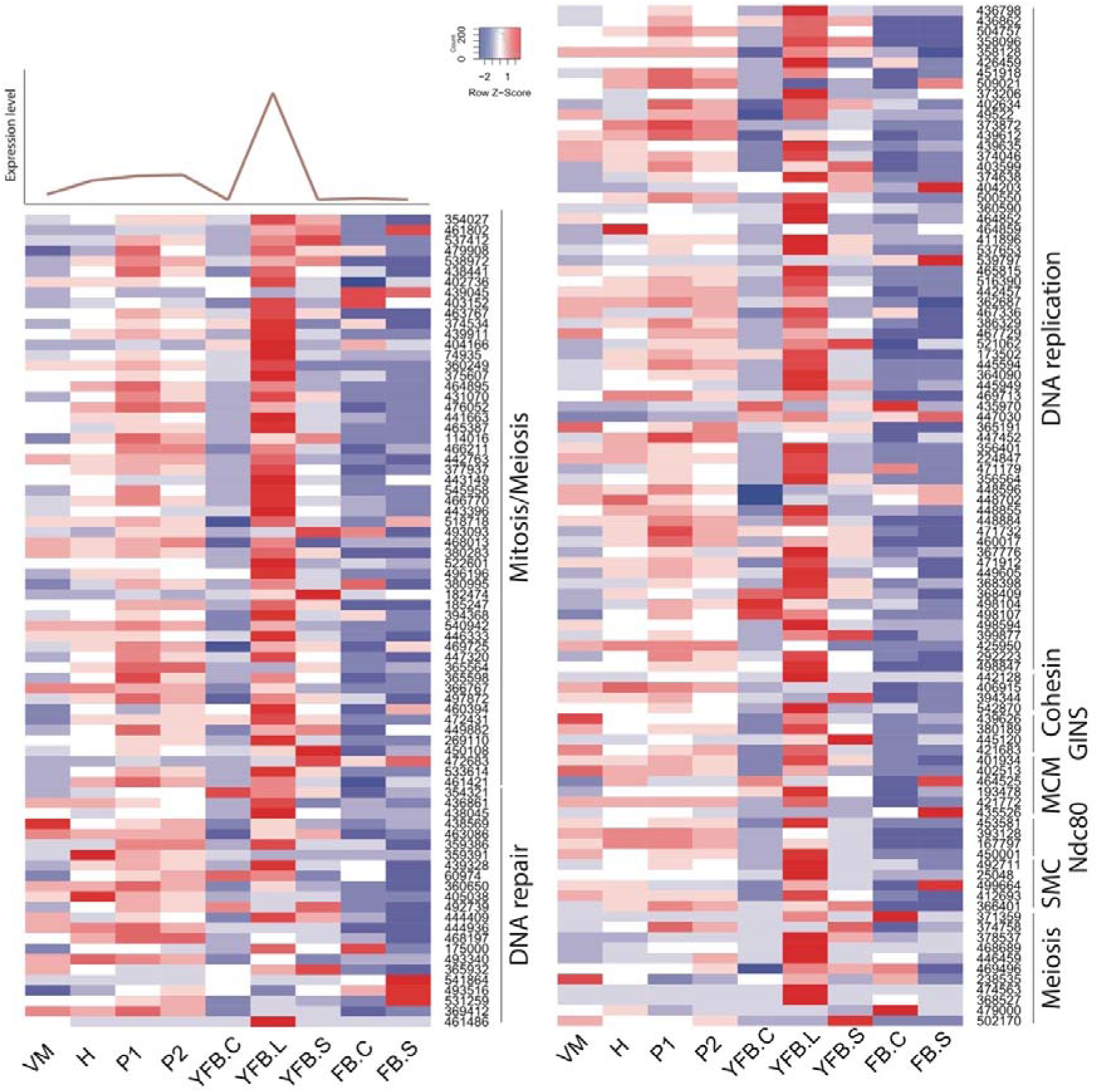
Expression heatmap of DNA replication, repair, mitosis and meiosis related genes in *C. cinerea*. A model expression trajectory, illustrating the two peaks characteristic of most examined species, is shown above the left panel. Well-delimited complexes mentioned in the paper are shown separately. Genes are denoted by Protein IDs. Blue and red colors represent low and high expression, respectively. Developmental stages are abbreviated as follows: VM – vegetative mycelium, H – hyphal knot, P1 – stage 1 primordium, P2 – stage 2 primordium, YFB.C – young fruiting body cap, YFB.L – young fruiting body gills, YFB.S – young fruiting body stipe, FB.C – mature fruiting body cap (including gills), FB.S – mature fruiting body stipe.

On the other hand, mitotic, DNA replication and repair genes showed a clear peak associated with primordium stages, and their expression leveled off in mature fruiting bodies (Fig. 3, Supplementary Fig. 1). This probably reflects intense cell proliferation in primordia, which establishes the main tissues and developmental modules of the mushroom. The expression of these genes later decreases to low levels in young fruiting body stipes and caps (also high in meiotic tissues where one round of mitosis happens) and in mature fruiting bodies. This peak was also detected in previous studies on *Agaricus blazei*(Lu et al. 2020) and *P. ostreatus*(Merényi et al. 2021) and probably relates to the biphasic development of fruiting bodies, where the first phase is concerned with cell proliferation and tissue formation whereas the second with growth by cell expansion without much change in cell numbers. Our data suggest that this pattern should be conserved across agarics, but potentially missing or less clear in species that follow different developmental patterns (e.g. polypores).

*Cryptococcus neoformans* recently proved to be a particularly useful model for teasing apart morphogenetic processes(Merényi et al. 2021). This species forms simple fruiting bodies composed of lawns of aerial hyphae bearing basidia. Liu et al pointed out in *Cryptococcus*, that genes co-induced with meiotic genes might be related to basidium morphogenesis, a process that greatly overlaps with meiosis(Liu et al. 2018). Based on transcriptome data it may be hard to separate such genes from meiotic genes, because of the temporal and spatial overlap of their expected expression. Nevertheless, one of the genes reported by Liu et al, *C. neoformans* Csa1 (*C. cinerea* protein ID: 471238, see Table TRA below), which has been implicated in basidium development (and to a lesser extent meiosis(Liu et al. 2018)), is developmentally expressed in 10/10 species in our data and shows expression peaks that overlap with that of meiotic genes.

An orthogroup that includes the serine-threonine kinases *Sch. pombe* Ran1 and *S. cerevisiae* Sks1 (represented by *C. cinerea* 456276, Table 1) showed marked peaks in gills of mature fruiting bodies in all agaricoid species (except *C. cinerea*) and in mature fruiting bodies in species with no separable gills or caps (*S. commune, A. ampla*). Ran1 is a negative regulator of *Sch. pombe* meiosis (with Mei2 as target(Caligiuri et al. 1997)), whereas Sks1 is involved in the adaptation to low glucose concentrations and pseudohyphal growth in yeast. Despite this apparent strong contradiction, the characteristic expression peaks in meiotic tissues and developmental stages in mushroom-forming fungi suggest roles in sporulation-related processes, possibly the repression of meiotic processes during spore morphogenesis.

In summary, mitotic and meiotic (including associated processes such as DNA replication and repair) gene expression showed conserved patterns across the examined mushroom-forming fungi. This probably reflects, on one hand, intense cell proliferation during primordium development and, on the other hand, meiosis that precedes sporulation in basidia. These gene expression patterns are easily recognizable in transcriptomic data and provide landmarks for calibrating developmental series. The downregulation of mitosis related genes in the second half of fruiting body development coincides with ceasing cell proliferation and the transition of the mushroom to growth by cell expansion.

#### 4.1.2. Ribosomal genes

Ribosomes produce proteins, the necessary building blocks for living cells. They are composed of four RNA subunits and up to 80 different proteins. The amount of ribosomes, and thus ribosomal protein expression, in a cell is proportional to its protein synthesis demand(Kraakman et al. 1993; Jorgensen et al. 2002). Therefore, ribosomal protein expression has been used as a proxy for the proliferative activity of a cell or a tissue and we expect it can be similarly informative in mushrooms too.

We observed a preponderance of ribosomal protein genes among CDE orthogroups, especially at FC>2 and to a lesser extent at FC>4. To look globally at structural proteins of ribosomes, we identified ribosomal protein genes based on previously reported ribosomal compositions, excluding mitochondrial ribosomal proteins and ribosome biogenesis-related genes. The 74 orthogroups identified this way (Table RIB) showed very characteristic expression dynamics, albeit at low fold change values, in most species. Ribosomal genes were highly expressed in the early developmental stages of most species followed by gradual shutting down in young and mature fruiting bodies. A second, sharp expression peak was observed in gill tissues of *P. ostreatus, A. ostoyae, A. bisporus* and *C. cinerea (*Fig. 4, Supplementary Fig. 2). Like in the case of mitotic and meiotic genes, we hypothesize that these two peaks correspond to an early, proliferative stage of development and the protein synthesis demand of spore production/meiosis. Ribosomal genes seem to be co-expressed with genes involved in cell division and proliferation (meiotic, DNA replication and repair, see above) in both primordia and in the gills. This could signify the need for increased protein production for cell proliferation in primordia and for meiosis and spore production, although other processes happening in gills requiring a large amount of protein cannot be ruled out. The characteristic expression of ribosomal genes is a conspicuous pattern and was detected in partial or complete form in several transcriptomic papers in both Basidiomycota(Cheng et al. 2013b; Zhou et al. 2014; Zhang et al. 2015b; Song et al. 2018b; Krizsán et al. 2019; Liu et al. 2020a; Lu et al. 2020) and Ascomycota(Rodenburg et al. 2018; Tong et al. 2020).

**Fig. 4.**
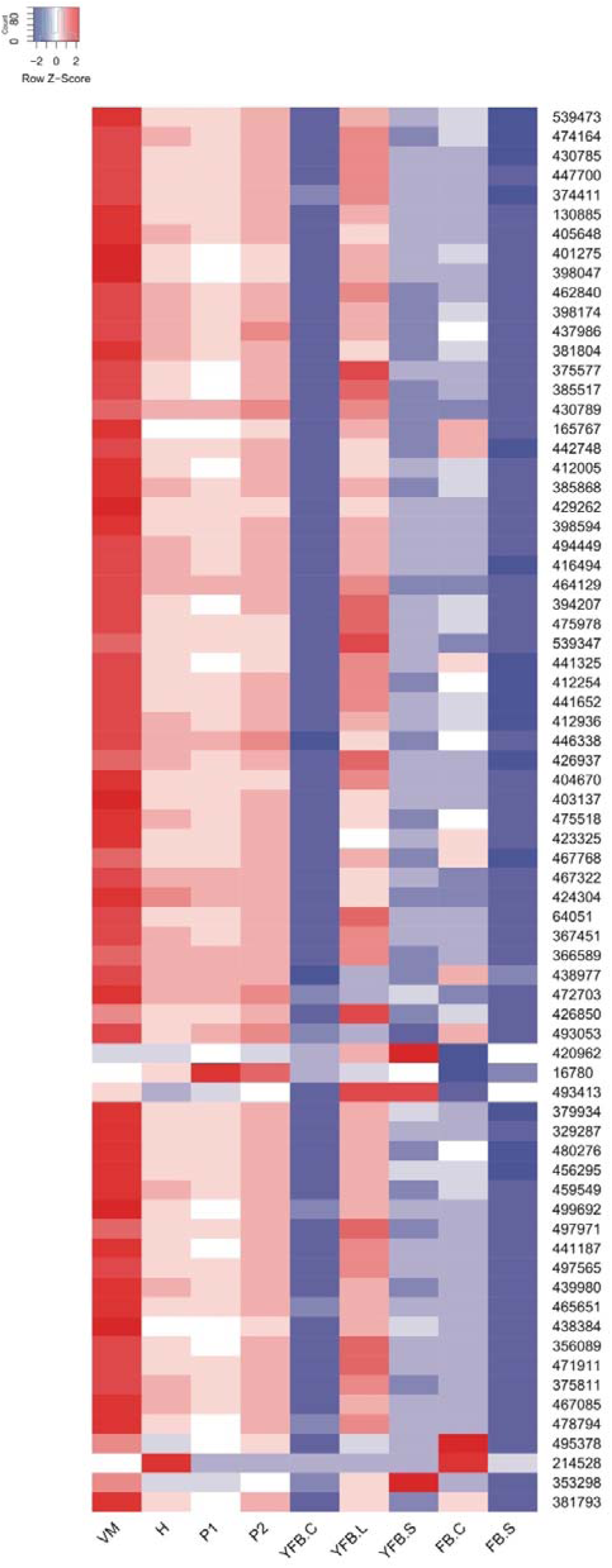
Expression heatmap of ribosomal protein encoding genes in *C. cinerea*. Genes are denoted by Protein IDs. Blue and red colors represent low and high expression, respectively. Developmental stages are abbreviated as follows: VM – vegetative mycelium, H – hyphal knot, P1 – stage 1 primordium, P2 – stage 2 primordium, YFB.C – young fruiting body cap, YFB.L – young fruiting body gills, YFB.S – young fruiting body stipe, FB.C – mature fruiting body cap (including gills), FB.S – mature fruiting body stipe.

The vegetative mycelium generally showed high ribosomal protein-coding gene expression, though not in all species, which we hypothesize depended on whether samples for RNA-Seq were harvested from older, inactive or actively growing (e.g. colony edge) parts of the mycelium.

A ribosome-bound Hsp70 chaperone protein formed a CDE orthogroup developmentally regulated in 9 species (FC>2) (*C. cinerea* protein ID: 438977) (Table 2). This gene was also detected previously as upregulated in primordia of *C. cinerea*(Cheng et al. 2013b). It is orthologous to *S. cerevisiae* Ssb2, a Hsp70-family chaperone that is bound by the ribosome and escorts the folding of nascent polypeptides(Peisker et al. 2010).

**Table 2.** Summary of developmental expression dynamics of CDE orthogroups of ribosomal genes across 12 species. Protein ID of a representative protein is given follows by the number of species in which the orthogroup is developmentally regulated at fold change 2 and 4 (FC>2 and FC>4, respectively). Putative function and ortholog in *S. cerevisiae* (if any) are also given. Abbreviations: 0-gene absent, 1-gene present but not developmentally regulated, 2 - developmentally regulated at fold change >2, 4- developmentally regulated at fold change >4. Species names are abbreviated as: Cc – *C. cinerea*, Aa – *A. ampla*, Ab – *A. bisporus*, Ao – *A. ostoyae*, Ca – *C. aegerita*, Lb – *L. bicolor*, Le – *L. edodes*, Lt – *L. tigrinus*, Mk – *M. kentingensis*, Pc – *Ph. chrysosporium*, Po – *P. ostreatus*, Sc – *S. commune*.

| ID | Fold change |  | Putative function | S.<br>*cerevisiae*<br>ortholog | Species |  |  |  |  |  |  |  |  |  |  |  |
| --- | --- | --- | --- | --- | --- | --- | --- | --- | --- | --- | --- | --- | --- | --- | --- | --- |
|  | >2 | >4 |  |  | Cc | Aa | Ab | Ao | Ca | Lb | Le | Lt | Mk | Pc | Po | Sc |
| 423325 | 10 | 3 | 60S ribosomal protein L28e | – | 4 | 2 | 2 | 2 | 2 | 4 | 2 | 2 | 4 | 1 | 2 | 1 |
| 405648 | 10 | 2 | 60S ribosomal protein L19-B | YBL027W | 4 | 2 | 2 | 2 | 2 | 4 | 2 | 2 | 2 | 1 | 2 | 1 |
| 412936 | 10 | 2 | 40S ribosomal protein S8-A | YBL072C | 4 | 2 | 2 | 2 | 1 | 2 | 2 | 2 | 2 | 2 | 4 | 1 |
| 478794 | 11 | 3 | 60S ribosomal protein L23-A | YBL087C | 4 | 2 | 2 | 2 | 2 | 4 | 2 | 2 | 4 | 2 | 2 | 1 |
| 366589 | 9 | 5 | 60S ribosomal protein L4-A | YBR031W | 4 | 2 | 4 | 2 | 0 | 4 | 4 | 2 | 4 | 1 | 2 | 1 |
| 497971 | 7 | 5 | 40S ribosomal protein S11-B | YBR048W | 4 | 2 | 4 | 0 | 2 | 4 | 0 | 1 | 4 | 1 | 4 | 1 |
| 464129 | 8 | 4 | 40S ribosomal protein S6-B | YBR181C | 4 | 2 | 4 | 2 | 1 | 4 | 2 | 1 | 4 | 1 | 2 | 1 |
| 430785 | 8 | 2 | 60S ribosomal protein L21-A | YBR191W | 4 | 2 | 2 | 2 | 2 | 1 | 4 | 1 | 0 | 2 | 2 | 1 |
| 467322 | 9 | 5 | 40S ribosomal protein S14-A | YCR031C | 4 | 2 | 4 | 2 | 2 | 4 | 2 | 1 | 4 | 1 | 4 | 1 |
| 493413 | 9 | 1 | Putatively involved intra-nuclear transport of pre-60S ribosomal subunits | YCR072C | 4 | 2 | 2 | 2 | 1 | 2 | 2 | 2 | 2 | 1 | 2 | 1 |
| 403137 | 11 | 4 | 60S ribosomal protein L31-A | YDL075W | 4 | 2 | 4 | 2 | 2 | 4 | 2 | 2 | 4 | 2 | 2 | 1 |
| 398174 | 9 | 3 | 40S ribosomal protein S16-B | YDL083C | 4 | 2 | 2 | 2 | 2 | 2 | 2 | 1 | 4 | 1 | 4 | 1 |
| 412254 | 8 | 2 | 60S ribosomal protein L35-B | YDL136W | 4 | 2 | 2 | 2 | 2 | 4 | 2 | 1 | 0 | 1 | 2 | 1 |
| 493053 | 10 | 4 | Putatively involved in pseudouridylation of rRNA | YDL208W | 4 | 2 | 2 | 4 | 2 | 2 | 2 | 2 | 4 | 1 | 4 | 1 |
| 64051 | 9 | 4 | 40S ribosomal protein S13 | YDR064W | 4 | 2 | 4 | 2 | 1 | 4 | 2 | 2 | 2 | 1 | 4 | 1 |
| 381793 | 10 | 4 | 60S acidic ribosomal protein P2-beta | YDR382W | 4 | 2 | 2 | 2 | 2 | 4 | 2 | 2 | 4 | 1 | 4 | 1 |
| 459549 | 8 | 2 | 60S ribosomal protein L12-B | YDR418W | 4 | 2 | 2 | 2 | 1 | 4 | 2 | 1 | 2 | 1 | 2 | 0 |
| 404670 | 11 | 4 | 40S ribosomal protein S17-B | YDR447C | 4 | 2 | 4 | 2 | 2 | 4 | 2 | 1 | 4 | 2 | 2 | 2 |
| 442748 | 8 | 2 | 40S ribosomal protein S18-A | YDR450W | 4 | 2 | 2 | 2 | 0 | 4 | 2 | 1 | 0 | 1 | 2 | 2 |
| 499692 | 10 | 5 | 60S ribosomal protein L27-B | YDR471W | 4 | 2 | 2 | 2 | 2 | 4 | 4 | 1 | 4 | 2 | 4 | 1 |
| 429262 | 9 | 4 | 60S ribosomal protein L34-A | YER056C-A | 4 | 2 | 2 | 2 | 2 | 4 | 4 | 1 | 0 | 2 | 4 | 1 |
| 439980 | 9 | 3 | 40S ribosomal protein S24-A | YER074W | 4 | 0 | 0 | 2 | 2 | 4 | 2 | 2 | 2 | 1 | 4 | 2 |
| 329287 | 10 | 5 | 40S ribosomal protein S26-B | YER131W | 4 | 2 | 4 | 2 | 2 | 4 | 2 | 2 | 4 | 1 | 4 | 1 |
| 426937 | 9 | 2 | 60S ribosomal protein L2-A | YFR031C-A | 4 | 2 | 2 | 2 | 2 | 4 | 2 | 1 | 0 | 2 | 2 | 1 |
| S.c.<br>274266<br>7 | 10 | 4 | 60S ribosomal protein L7A | YGL076C | 0 | 2 | 4 | 2 | 2 | 4 | 4 | 2 | 4 | 2 | 2 | 1 |
| 441325 | 10 | 3 | 60S ribosomal protein L28 | YGL103W | 4 | 2 | 2 | 2 | 2 | 4 | 2 | 1 | 4 | 2 | 2 | 1 |
| 475518 | 10 | 4 | 40S ribosomal protein S2 | YGL123W | 4 | 2 | 4 | 2 | 2 | 4 | 2 | 2 | 4 | 1 | 2 | 1 |
| 437986 | 9 | 4 | 60S ribosomal protein L1-B | YGL135W | 4 | 2 | 2 | 2 | 2 | 4 | 2 | 1 | 4 | 1 | 4 | 1 |
| 480276 | 8 | 3 | 40S ribosomal protein S25-A | YGR027C | 4 | 2 | 4 | 2 | 2 | 4 | 2 | 1 | 0 | 1 | 2 | 1 |
| 475978 | 10 | 3 | 60S ribosomal protein L26-B | YGR034W | 4 | 2 | 2 | 0 | 2 | 4 | 4 | 2 | 2 | 2 | 2 | 1 |
| 130885 | 9 | 3 | 60S ribosomal protein L11-B | YGR085C | 4 | 2 | 4 | 2 | 2 | 4 | 2 | 2 | 0 | 1 | 2 | 1 |
| 495378 | 5 | 0 | 40S ribosomal protein S23-A | YGR118W | 2 | 1 | 2 | 2 | 1 | 2 | 1 | 1 | 0 | 2 | 1 | 1 |
| 379934 | 9 | 3 | 60S ribosomal protein L24-B | YGR148C | 4 | 2 | 2 | 2 | 2 | 4 | 2 | 0 | 4 | 1 | 2 | 1 |
| 494449 | 10 | 4 | 40S ribosomal protein S0-A | YGR214W | 4 | 2 | 4 | 2 | 2 | 4 | 2 | 2 | 4 | 1 | 2 | 1 |
| 467085 | 10 | 3 | 60S ribosomal protein L14-B | YHL001W | 4 | 2 | 2 | 2 | 2 | 4 | 2 | 2 | 4 | 1 | 2 | 1 |
| 165767 | 9 | 3 | 40S ribosomal protein S20 | YHL015W | 4 | 2 | 2 | 2 | 2 | 4 | 2 | 2 | 0 | 1 | 4 | 1 |
| 438384 | 6 | 1 | 40S ribosomal protein S27-B | YHR021C | 4 | 1 | 1 | 2 | 0 | 2 | 2 | 1 | 2 | 1 | 2 | 1 |
| 214528 | 6 | 6 | Protein involved in rRNA-processing | YHR087W | 4 | 1 | 4 | 4 | 1 | 1 | 4 | 4 | 1 | 1 | 4 | 0 |
| 441187 | 9 | 4 | 60S ribosomal protein L42-B | YHR141C | 4 | 2 | 1 | 2 | 2 | 4 | 2 | 1 | 4 | 2 | 4 | 1 |
| 441652 | 7 | 3 | 40S ribosomal protein S4-B | YHR203C | 4 | 2 | 0 | 2 | 0 | 4 | 2 | 1 | 4 | 1 | 2 | 1 |
| 497565 | 8 | 2 | 60S ribosomal protein L39 | YJL189W | 4 | 1 | 2 | 2 | 0 | 4 | 2 | 2 | 0 | 2 | 2 | 1 |
| 375577 | 9 | 5 | 40S ribosomal protein S22-A | YJL190C | 4 | 2 | 4 | 2 | 2 | 4 | 4 | 1 | 4 | 1 | 2 | 1 |
| 385517 | 8 | 4 | 60S ribosomal protein L43-B | YJR094W-A | 4 | 2 | 0 | 0 | 0 | 4 | 2 | 2 | 4 | 2 | 4 | 1 |
| 446338 | 10 | 3 | 40S ribosomal protein S5 | YJR123W | 4 | 2 | 4 | 2 | 2 | 4 | 2 | 1 | 2 | 2 | 2 | 1 |
| 472703 | 8 | 2 | 60S ribosomal protein L8-B | YLL045C | 2 | 2 | 4 | 2 | 2 | 2 | 2 | 1 | 0 | 1 | 4 | 1 |
| 374411 | 7 | 2 | 60S ribosomal protein L22-A | YLR061W | 2 | 2 | 4 | 2 | 2 | 1 | 2 | 1 | 0 | 1 | 4 | 1 |
| 471911 | 7 | 4 | Ubiquitin-40S ribosomal protein S31 | YLR167W | 4 | 0 | 4 | 2 | 1 | 4 | 2 | 1 | 4 | 1 | 2 | 1 |
| 447700 | 7 | 3 | 60S ribosomal protein L37-A | YLR185W | 4 | 0 | 2 | 2 | 0 | 4 | 2 | 1 | 4 | 1 | 2 | 1 |
| 353298 | 7 | 2 | 40S ribosomal protein S28-B | YLR264W | 4 | 2 | 2 | 2 | 0 | 4 | 2 | 1 | 0 | 2 | 0 | 0 |
| 398047 | 7 | 3 | 60S ribosomal protein L38 | YLR325C | 4 | 2 | 2 | 0 | 0 | 4 | 2 | 1 | 0 | 2 | 4 | 1 |
| 367451 | 7 | 3 | 60S acidic ribosomal protein P0 | YLR340W | 4 | 2 | 1 | 2 | 1 | 2 | 4 | 1 | 2 | 1 | 4 | 1 |
| 398594 | 9 | 3 | 40S ribosomal protein S29-A | YLR388W | 4 | 2 | 2 | 2 | 0 | 4 | 2 | 2 | 0 | 2 | 4 | 1 |
| 385868 | 10 | 4 | 60S ribosomal protein L6-B | YLR448W | 4 | 2 | 4 | 2 | 2 | 4 | 2 | 2 | 0 | 2 | 4 | 1 |
| 474164 | 9 | 5 | 40S ribosomal protein S1-B | YML063W | 4 | 2 | 4 | 2 | 2 | 4 | 4 | 1 | 2 | 1 | 4 | 1 |
| 424304 | 8 | 4 | 60S ribosomal protein L13-B | YMR142C | 4 | 2 | 0 | 2 | 2 | 4 | 0 | 4 | 2 | 1 | 4 | 1 |
| 456295 | 8 | 2 | 60S ribosomal protein L36-A | YMR194W | 4 | 2 | 2 | 2 | 0 | 4 | 2 | 2 | 0 | 1 | 2 | 1 |
| 375811 | 8 | 3 | 60S ribosomal protein L20-A | YMR242C | 4 | 2 | 4 | 2 | 0 | 4 | 2 | 1 | 2 | 1 | 2 | 1 |
| 412005 | 9 | 3 | 60S ribosomal protein L9-B | YNL067W | 4 | 2 | 2 | 2 | 2 | 4 | 2 | 1 | 2 | 1 | 4 | 1 |
| 416494 | 5 | 2 | 40S ribosomal protein S3 | YNL178W | 4 | 2 | 0 | 0 | 1 | 4 | 2 | 1 | 2 | 1 | 0 | 1 |
| 438977 | 9 | 0 | Ribosome-bound, Hsp70-type chaperone | YNL209W | 2 | 2 | 2 | 2 | 2 | 1 | 2 | 2 | 2 | 1 | 2 | 1 |
| 462840 | 8 | 4 | 60S ribosomal protein L18-B | YNL301C | 4 | 2 | 4 | 2 | 1 | 4 | 2 | 1 | 2 | 1 | 4 | 1 |
| 21836 | 6 | 0 | Putatively involved in nucleolar processing of pre-18S ribosomal RNA | YNL308C | 2 | 1 | 2 | 2 | 2 | 1 | 1 | 1 | 2 | 1 | 2 | 1 |
| 381804 | 8 | 2 | 40S ribosomal protein S15 | YOL040C | 4 | 2 | 2 | 2 | 0 | 4 | 2 | 1 | 2 | 1 | 2 | 1 |
| 465651 | 10 | 4 | 40S ribosomal protein S19-A | YOL121C | 4 | 2 | 4 | 2 | 2 | 4 | 2 | 2 | 4 | 1 | 2 | 1 |
| 426850 | 7 | 4 | 60S ribosomal protein L3 | YOR063W | 4 | 2 | 2 | 0 | 2 | 4 | 0 | 1 | 4 | 1 | 4 | 1 |
| 401275 | 9 | 6 | 40S ribosomal protein S12 | YOR369C | 4 | 2 | 1 | 4 | 0 | 4 | 4 | 2 | 4 | 2 | 4 | 1 |
| 430789 | 10 | 4 | 40S ribosomal protein S9-A | YPL081W | 4 | 2 | 4 | 2 | 2 | 4 | 2 | 2 | 4 | 1 | 2 | 1 |
| 467768 | 8 | 5 | 60S ribosomal protein L5 | YPL131W | 4 | 2 | 4 | 2 | 2 | 4 | 4 | 1 | 0 | 1 | 4 | 1 |
| 356089 | 10 | 4 | 60S ribosomal protein L33-A | YPL143W | 4 | 2 | 2 | 0 | 2 | 4 | 2 | 2 | 4 | 2 | 4 | 1 |
| 539347 | 5 | 3 | 60S ribosomal protein L29 | YFR032C-A | 4 | 2 | 0 | 0 | 0 | 4 | 4 | 1 | 0 | 2 | 0 | 1 |
| 394207 | 7 | 3 | 60S ribosomal protein L30 | YGL030W | 4 | 2 | 0 | 2 | 0 | 4 | 2 | 1 | 4 | 1 | 2 | 1 |
| 539473 | 7 | 2 | 60S ribosomal protein L16-A | YIL133C | 4 | 2 | 2 | 0 | 2 | 4 | 2 | 1 | 0 | 1 | 2 | 1 |

In summary, a characteristic expression trajectory of ribosomal genes is characteristic of most mushroom-forming fungi, which likely reflects high protein demands of intensely growing tissue in primordial stages and gills and supports a bi-phasic model of fruiting body development (see below).

#### 4.1.3. Growth by cell expansion and turgor manipulation - a biphasic model of fruiting body development

A key difference in fruiting bodies and hyphae is that in the latter growth and expansion takes place at the tip, whereas in the former, cell expansion can vary between apical and isotropic. Already de Bary established, by studying *Mycena vulgaris*, that mushroom fruiting bodies grow mostly by cell expansion not by cell proliferation(Bary et al. 1887). This appears to be a rather widespread feature of fruiting bodies and shows broad similarity to growth by cell expansion in plant fruits(Krizsán et al. 2019). More precisely, fruiting body development seems to be divided into a first proliferative stage, during which cell division, differentiation as well as tissue formation takes place, and a growth phase, which is primarily concerned with cell expansion to achieve the mature shapes and sizes of the fruiting bodies(MOORE et al. 1979; Kües 2000; G et al. 2013; Kües and Navarro-González 2015; Krizsán et al. 2019; Liu et al. 2021). Of the better investigated species, this two-phased development is probably most spectacular in *C. cinerea*, which is adapted to very rapidly lifting cap above ground and expanding it for spore dispersal(Kües 2000).

It should be noted that the two-phase development is not fully universal among fungi and does not always mean a complete separation of proliferative and expansive phases. For example, Craig et al(Craig et al. 1977) showed that the top regions of *A. bisporus* stipes elongated up to 9-fold, whereas cell lengths increased up to 2.2-fold, and from these data derived that cell division also occurs to achieve this elongation rate. In *S. commune*, it appears that outer parts of the fruiting body are still growing (by cell proliferation) while older parts are already sporulating(Kues and Liu 2000).

Whether for the entire fruiting body or only parts of it, growth by cell expansion seems to be a widespread trait during the second half of development of mushrooms. How cell expansion is achieved, in general, is poorly known, but it probably requires water supply, turgor manipulation (by osmolytes), cell wall elasticity and new cell wall material deposition. Water supply may be realized by the regulated expression of aquaporin genes (see below). On the other hand, the identity of the osmolytes has not been resolved yet. In the Ascomycota ions, ion channels, glycerol and sugars were reported as main players of turgor manipulation(Fischer et al. 2004). In Basidiomycota fruiting bodies, sugars (mannitol, trehalose) offer themselves as straightforward candidates, but Moore([CSL STYLE ERROR: reference with no printed form.]) speculated that high molecular weight sugars are unlikely to be the osmolytes that drive stipe expansion. Trehalose and polyols have been excluded as osmolytes(Kües 2000; F et al. 2005), whereas potassium oxaloacetate has been hypothesized, which would be consistent with the upregulation of oxalate metabolism (see below), but not confirmed(Knoll; Kües 2000). On the other hand, ions would either need to be transported to the fruiting body from the mycelium, or released locally.

Factors modulating cell wall elasticity and remodeling are better known. During fruiting body growth, cell wall deposition occurs throughout the cell surface, and not only in the tip region as in hyphae, which was termed diffuse extension(Kües 2000). In *Agaricus bisporus* larger proportion of β-1,6-linked glucose side branches in a β-1,3-glucan backbone was observed in the rapidly elongating top segment of the stipe(Mol and Wessels 1990). This causes hydrogen-bonds among glucan chains to be weaker in rapidly elongating stipe compared with mature stipe and vegetative hyphae. It has been hypothesized that these weak bonds are disrupted and reformed by turgor pressure rather than hydrolysing enzymes(Mol et al. 1990). The architecture of cell wall also regulates stipe elongation; in primordial stages chitin microfibrils are arranged isotropically which is remodeled to helical arrangement during development(Kamada et al. 1991). The turgor pressure acting on the cell wall and insertion of new chitin microfibrils allow elongation and in that dynamic rearranges of microfibrils chitin synthases, chitinases and glucanases play the major role in *C. cinerea*(Kamada et al. 1991). Niu et al observed differences in structure of chitin microfibrils in elongating and mature stipe regions(Niu et al. 2015). Extension of the cell wall happens by insertion of newly synthesized microfibrils. In contrast, studies in *Flammulina* and *Coprinopsis* suggested that stipe cell wall extension is mediated by endogenous expansin-like proteins, not turgor pressure or cell wall glycoside hydrolases(Fang et al. 2014; W et al. 2014). Plant expansins mediate Turgor-mediated cell expansion(Baccelli 2015) and several other morphogenetic processes (e.g. organ abscission, fruit softening(Sampedro and Cosgrove 2005)). The auxin-driven expansion of plant cells without changes in cell number is known as acid growth(Rayle and Cleland 1992; Arsuffi and Braybrook 2018), because low pH is required by cell wall modifying proteins (e.g. expansins) to loosen the cell wall. In fungi, it is not known if an analogous process exists, but the acidifying effect of external CO_2_ has been hypothesized as a factor in stipe cell elongation(Liu et al. 2021) and there is evidence for expansins participating in the process (see below). Accordingly, high external CO_2_ during mushroom cultivation was speculated to cause malformed, long-stipe fruiting bodies by acidifying the extracellular space and activating cell wall modifying proteins(Liu et al. 2021).

### 4.2. Acetyl-CoA production by central carbon metabolism

We found evidence for widespread developmental expression of multiple acetyl-coenzyme A (Ac-CoA) producing enzyme families. Ac-CoA is a key intermediate in central carbon metabolism as well as other processes (biosynthesis of lipids and certain amino acids(Chen et al. 2015)). Under normal carbon metabolic conditions, the generated Ac-CoA, which comes from either glycolysis or fatty acid beta-oxidation enters the TCA cycle, where it is oxidized to CO_2_ for energy (ATP) production. However, Ac-CoA is required also for the production of biological membranes(Strijbis and Distel 2010; Chen et al. 2015), and our results point in the direction of multiple basic metabolic pathways converging on excess Ac-CoA production for membrane biosynthesis. Combined with gene expression profiles of these genes, we hypothesize that excess Ac-CoA is being channeled into ergosterol and fatty acid production during fruiting body development, as detailed below.

For this work we started from Ac-CoA related CDE orthogroups detected in the transcriptomes of the examined species and developmental stages, which belonged to several pathways. We reasoned that if there is an increased demand for Ac-CoA at certain developmental stages of fruiting bodies, then this should be reflected in the regulation of Ac-CoA-producing pathways. Thus, we annotated the TCA cycle, the glyoxal cycle, the fatty acid beta-oxidation, ethanol utilization, glycolytic as well as pyruvate decarboxylation pathways of *C. cinerea*, based on protein orthology to corresponding pathways of *S. cerevisiae* (Fig. 5, Supplementary Fig. 3, Table 3). These led to a model we put forth on Fig. 5, with the additional note that it reflects one possible synthesis of the observations into a mechanistic model compatible with what we know about fruiting body development, but that other interpretation may also be viable.

**Fig. 5.**
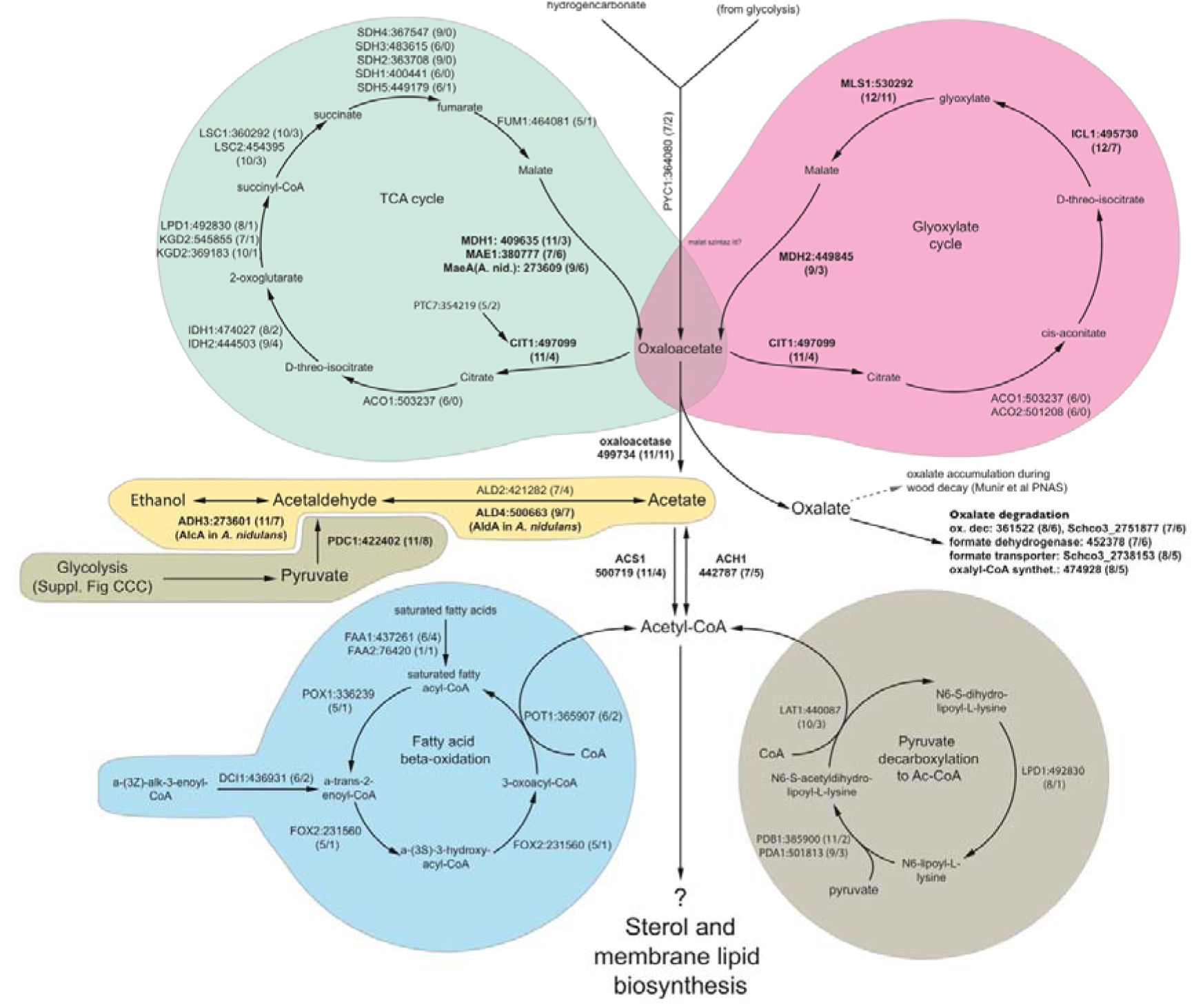
Basic metabolic pathways of *C. cinerea.* Genes in the pathways are denoted by the *S. cerevisiae* gene name, followed by the protein ID of the *C. cinerea* ortholog and, in parentheses the number of species in which the gene was developmentally regulated at fold change >2/>4. Pathways are colored differently for clarity.

**Table 3.**
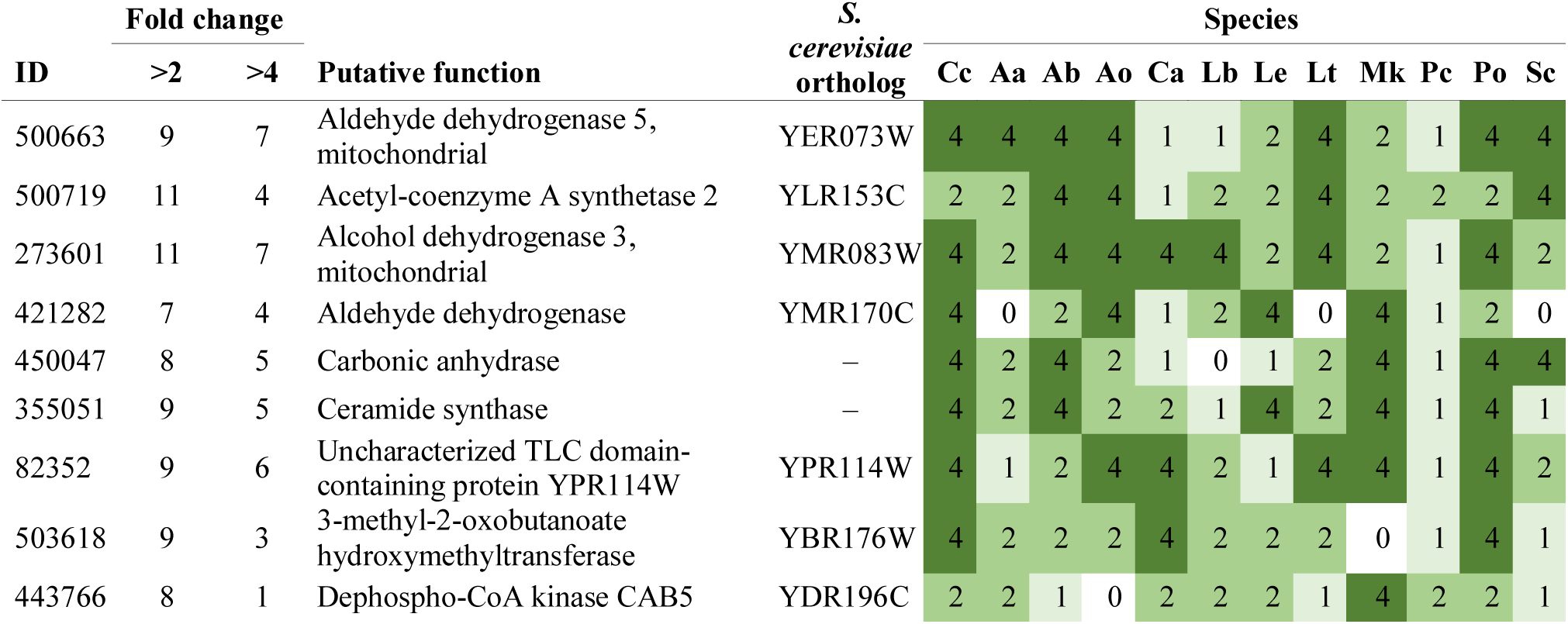
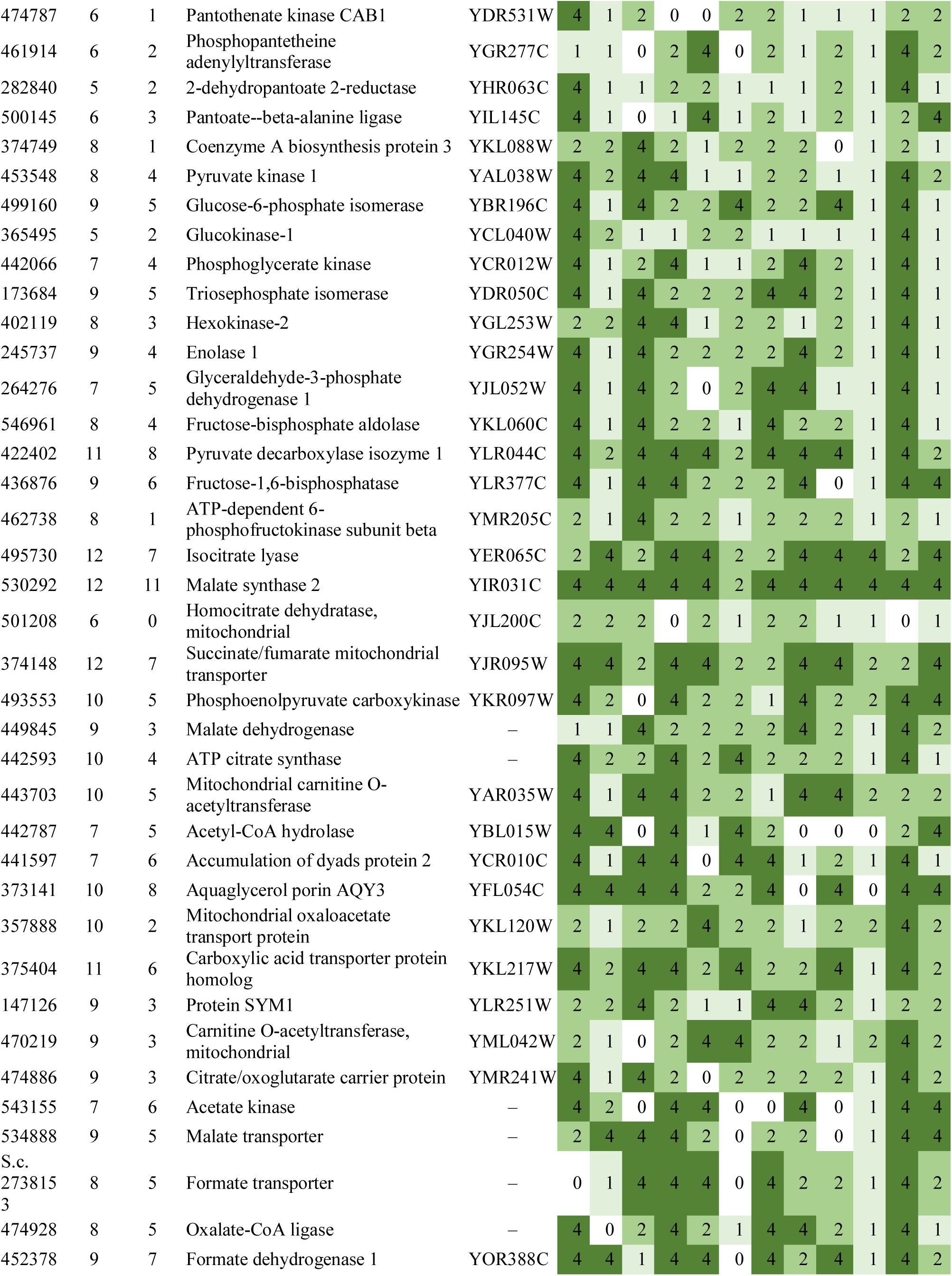
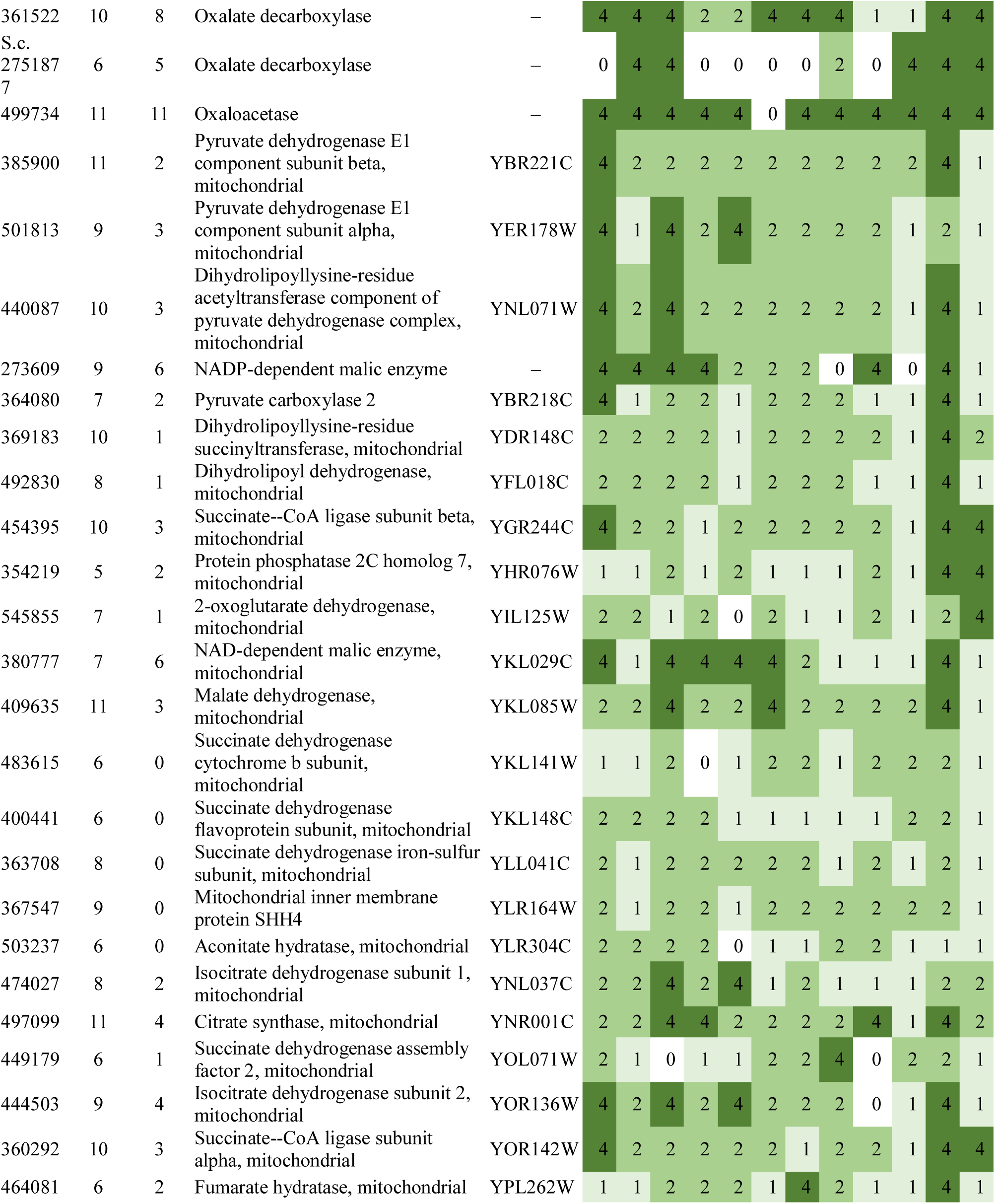
Summary of developmental expression dynamics of CDE orthogroups of genes related to Acetyl-CoA production and basic metabolism across 12 species. Protein ID of a representative protein is given follows by the number of species in which the orthogroup is developmentally regulated at fold change 2 and 4 (FC>2 and FC>4, respectively). Putative function and ortholog in *S. cerevisiae* (if any) are also given. Abbreviations: 0-gene absent, 1-gene present but not developmentally regulated, 2 - developmentally regulated at fold change >2, 4- developmentally regulated at fold change >4. Species names are abbreviated as: Cc – *C. cinerea*, Aa – *A. ampla*, Ab – *A. bisporus*, Ao – *A. ostoyae*, Ca – *C. aegerita*, Lb – *L. bicolor*, Le – *L. edodes*, Lt – *L. tigrinus*, Mk – *M. kentingensis*, Pc – *Ph. chrysosporium*, Po – *P. ostreatus*, Sc – *S. commune*.

First of all, we found that oxaloacetase (oxaloacetate lyase) genes were developmentally expressed in 11 species (FC>4, *C. cinerea* 499734). Oxaloacetases produce oxalate and acetate from oxaloacetate, which is an intermediate of both the TCA and the Glyoxylate cycles. In *S. cerevisiae* acetate is converted to Ac-CoA by acetyl-CoA synthases (Acs1), orthologs of which were developmentally regulated in 11 and 4 species at fold change of 2 and 4, respectively (represented by *C. cinerea* 500719). We also found a CDE orthogroup comprising putative orthologs of the *S. cerevisiae* Ac-CoA hydrolase Ach1 (represented by *C. cinerea* 442787, FC>2/4: 7/5). Ach1 is a relatively poorly known gene, which was recently shown to be responsible for the conversion of Ac-CoA to acetate, which can then be easily transferred from the mitochondrion to the cytoplasm(Chen et al. 2015). Cytoplasmic acetate is then converted to cytoplasmic Ac-CoA, a starting material for lipid synthesis, suggesting that Ach1 mediated Ac-CoA supply for lipid synthesis. Remarkably, Ach1 orthologs are present in only 8 species and completely missing in *A. bisporus* (missing even in telomere-to-telomere assemblies(ASM et al. 2020)), *M. kentingensis, Ph. chrysosporium, L. tigrinus, Pt. gracilis*. This patchiness might indicate that the role of these genes is dispensable and probably others can take over their roles.

Acetate is also an intermediate of the ethanol utilization pathway, in which two enzymes form a linear pathway that terminates in the production of Acetyl-CoA(M et al. 2003). Aldehyde dehydrogenases orthologous to AldA of *A. nidulans* and Ald4 of *S. cerevisiae* formed a CDE orthogroup (*C. cinerea* protein ID: 500663, 9/7); the encoded enzymes also contribute to acetate production and, in *C. cinerea*, showed peak expression in young fruiting body caps. Ald4/AldA converts the toxic compound acetaldehyde to acetate(B et al. 2001). Although the ethanol utilization pathway is mostly discussed in the context of extracellular ethanol (a nonfermentable carbon source) uptake and degradation, as well as anaerobic fermentation, acetaldehyde can also be converted to ethanol (Fig. 5), if it is present in excess amounts in the cells. This may explain the developmental regulation of orthologs of *S. cerevisiae* Adh3 and *A. nidulans* AlcA/AlcC (represented by *C. cinerea* 500663), which were developmentally expressed in 11/7 species (FC>2/4, Table 3). Enzymes of the alcohol utilization pathway were recently detected in proteomics data of fruiting bodies of *Guyanagaster necrorhiza*, where their presence was interpreted as evidence for anaerobic fermentation within the fruiting body, as part of an intriguing mutualism with N_2_ fixing bacteria(Koch et al. 2021).

The last step of glycolysis is catalyzed by the pyruvate decarboxylase Pdc1 in *S. cerevisiae,* which converts pyruvate to acetaldehyde. Pdc1 generates cytoplasmic Ac-CoA, which is essential for the biosynthesis of lipids(Chen et al. 2015). Pdc1 orthologs (represented by *C. cinerea* 422402) were developmentally expressed in 11/8 species (FC>2/4), whereas genes encoding other enzymes in glycolysis showed much smaller dynamics (Supplementary Fig. 3). Together, these observations suggest a widespread upregulation of acetaldehyde and from that either acetate or ethanol producing enzyme genes in fruiting bodies.

Upstream of oxaloacetases, enzymes responsible for oxaloacetate production in the TCA (yeast Mdh1, *C. cinerea* 500663, 11/3) and glyoxylate cycles (yeast Mdh2, *C. cinerea* 449845, 9/3) were also quite widely developmentally regulated. In the glyoxylate cycle we found orthologs of *S. cerevisiae* Mls1 (malate synthase, *C. cinerea* 530292) and Icl1 (isocitrate lyase, *C. cinerea* 495730) to form CDE orthogroups developmentally regulated in 12/11 and 12/7 species (FC>2/4), respectively. This suggests that the glyoxylate cycle is strongly induced in fruiting bodies. On the other hand, members of the TCA cycle showed lower developmental dynamics (Fig. 5).

What happens to the oxalate that is produced by oxaloacetases? It is possible that the resulting oxalate is a waste product, but it may also be used somehow by the fungus, e.g. as a developmental signal molecule(Palmieri et al. 2019) or as an electron donor, as has been observed in wood-decay(Hastrup et al. 2012; Zhu et al. 2016; Grąz et al. 2017), where it acts as an electron donor for oxidative enzymes. One hypothesis for the role of oxalate is that oxidative enzymes, which might participate in the modification of the fungal cell wall during development (see above), utilize it as an electron donor, as do enzymes involved in wood-decay. On the other hand, if oxalate is a by-product, excess amounts may need to be broken down actively. We found evidence for the upregulation of oxalate catabolism in the compared species. The two oxalate degradation pathway in fungi comprises oxalate decarboxylase, which converts oxalate to formate and CO_2_ and formate dehydrogenase, which breaks formate down to CO_2_(Mäkelä et al. 2014). We detected two CDE orthogroups of oxalate decarboxylases (*C. cinerea* protein ID: 361522, 8/6, Schco3_2751877, 7/6) and one of formate dehydrogenases (*C. cinerea* protein ID: 452378, 7/6, orthologous to Fdh1 of *S. cerevisiae*)(Table 3), which were upregulated in fruiting bodies widely, suggesting that fruiting bodies utilize enzymatic mechanisms for the neutralization of oxalate. An alternative route of oxalate degradation in *S. cerevisiae* is catalyzed by PCS60, an oxalyl-CoA synthetase(Foster and Nakata 2014). Orthologs of this gene were found in only a subset of species (represented by *C. cinerea* 474928), but were developmentally regulated in most of them. In addition, a CDE orthogroup of formate transporters, orthologous to the *Postia placenta* gene reported in Martinez et al(Martinez et al. 2009b), was developmentally expressed in 8/5 species (FC>2/4, represented by Schco3_2738153, missing in *C. cinerea*).

Building on the evidence for intense Ac-CoA production, we annotated and analyzed the Coenzyme-A producing pathway as well as two other sources of Ac-CoA, the fatty acid beta-oxidation pathway and the cellular cycle for pyruvate decarboxylation to Ac-CoA. These pathways were reconstructed and annotated based on corresponding pathways in *S. cerevisiae*(JM et al. 2012). Members of these pathways did not show remarkable developmental dynamics (Table 3).

Several genes connected to acetate metabolism and transport also formed CDE orthogroups. Orthologs of *S. cerevisiae* Ady2, an acetate transporter(S et al. 2004), are widely developmentally regulated (represented by *C. cinerea* 441597, 7/6), and were strongly induced in young fruiting body caps of *C. cinerea*. A CDE orthogroup of putative monocarboxylate (lactate, acetate, succinate, etc.) transporters (*C. cinerea* protein ID: 375404 [7/5])( Table 3) orthologous to *S. cerevisiae* Jen1(M et al. 1999) were also highly induced in YFB-caps in *C. cinerea*. We also identified putative orthologs of *S. cerevisiae* Yat1 (represented by *C. cinerea* 443703, 10/5), a carnitine acetyltransferase that shuttles Ac-CoA across the mitochondrial membrane. A CDE orthogroup of aquaglyceroporins orthologous to *S. cerevisiae* Fps1, which is involved in the uptake of acetate (*C. cinerea* protein ID: 373141 7/6), were developmentally regulated in 7/6 species. This gene encodes a protein that has been associated with either glycerol influx/efflux(Luyten et al. 1995) or acetate uptake(Mollapour and Piper 2007; Watcharawipas et al. 2018), which leaves some uncertainty on which role it fulfills in fruiting bodies. Orthologues of *S. cerevisiae* Sym1 also formed a CDE orthogroup, although only at a fold change cutoff of 2 (*C. cinerea* protein ID: 147126, 9/3) (Table 3). This gene is said to be involved in ethanol utilization and stress response, but mechanistic details are not known(Reinhold et al. 2012; Gilberti et al. 2018). Similarly, orthologs of yeast Sfc1, a succinate/fumarate antiporter that transports C4 succinate (which is produced from acetate(Bojunga et al. 1998)), to the mitochondria, are developmentally regulated in several species (represented by *C. cinerea* 374148, 12/7). The Sfc1 ortholog is also strongly induced late in the development of *C. neoformans*(Liu et al. 2018). Finally, a CDE orthogroup of putative malate symporters orthologous to *Sch. pombe* Mae1, which is involved in the uptake of extracellular malic acid was detected as developmentally regulated in 9/5 species (FC>2/4, represented by *C. cinerea* 534888, Table 3). Whether these genes are related to the above discussed observation of acetate/Ac-CoA related gene upregulation, or how cell surface transporters may be related remains to be determined. The induction of these genes does not happen in the same or similar developmental stage in all species (although there is consistency within species, see the upregulation of several genes in YFB-caps of *C. cinerea*), nevertheless, their induction suggests differential acetate concentrations and an important role for acetate during in fruiting body development.

CO_2_ produced by the TCA cycle may be fixed into oxaloacetate by the conversion of CO_2_ to hydrogencarbonate by carbonic anhydrases, then conversion of HCO_3_^-^ and pyruvate into oxaloacetate by pyruvate carboxylases(Elleuche and Pöggeler 2010) (Pyc1 in *S. cerevisiae*). We found that several carbonic anhydrases (2-9 genes in Agaricomycetes genomes) are developmentally regulated in mushroom forming fungi and they form at least one CDE orthogroup (represented by *C. cinerea* 450047)(Table 3). *C. cinerea* 450047 shows an upregulation in young fruiting bodies, coincident with the induction of Ac-CoA producing enzymes. This observation raises the possibility that CO_2_ produced by the TCA cycle in fruiting bodies may be fixed into oxaloacetate, which can further be converted to Ac-CoA (Fig. 5). Of note, carbonic anhydrases are involved in the regulation of sexual development in the Ascomycota(Elleuche and Pöggeler 2010).

Taken together, we observed developmental regulation of key enzymes in several pathways that seem to converge on increased acetyl-CoA production, either directly, or indirectly (through acetate or oxaloacetate). Several connected pathways, including oxalate degradation, the ethanol degradation pathway, the terminal reaction of glycolysis as well as evidence from the TCA and glyoxylate cycles were pointing in this direction. The expression of these genes tended to be upregulated late during development and showed good consistency within, but not necessarily across, species. A possible interpretation of these data is that increased Ac-CoA production is needed for growth by cell expansion late in development, as a starting material for lipid and sterol biosynthesis. This is underpinned by the fact that biosynthetic pathways of ergosterol and other membrane components are also upregulated during late developmental stages (e.g. the young fruiting body stage in *C. cinerea*), probably for membrane assembly needed for rapid cell growth. This interpretation would be consistent with the two-phased development of several mushroom-forming species, which consists of a first stage of cell division and differentiation and a second, during which cell numbers hardly change but their volumes increase considerably via Turgor-mediated expansion. Such a reprogramming of the fungus’s basic metabolism from using Ac-CoA for energy production (when citrate is in the mitochondrion) to using it for lipid metabolism (when citrate is in the cytoplasm) seems particularly well-suited for fruiting body formation, which requires tight regulation of lipid metabolism. It should be mentioned that Ac-CoA is also required for melanin production(Lee et al. 2019), although it is unlikely to be the driving force of our observation, because pigment production is not conserved across the examined species. A previously reported reprogramming of the glyoxylate cycle in *Fomitopsis palustris* for the production of oxalate(Munir et al. 2001) seems related, however, in that case, the main product in demand was hypothesized to be oxalate that facilitates wood-decay, whereas in our case we hypothesize the product in demand is acetate and finally Ac-CoA. Both variants of the metabolic pathways nevertheless pass through oxaloacetate(Munir et al. 2001). The excess Ac-CoA production may rely on, or be connected to carbohydrate reserves that are deposited in the subhymenial part of the cap of the mushroom(Ji and Moore 1993; Muraguchi et al. 2015) during primordium development. However, there may be alternative interpretations and the topic is certainly of interest for detailed, further research, that integrates more functional data beyond gene expression observations.

### 4.3. Lipid metabolism

Lipid metabolism is pivotal to very basic organismal processes such as growth, differentiation, cell division, energy storage and cell membrane composition, to name a few. Therefore, in depth understanding of gene expression changes associated with lipid metabolism, catabolism and transport can be expected to provide novel insights into fruiting body development. Indeed, our meta-analysis of Agaricomycete transcriptomes revealed several CDE orthogroups related to lipid metabolism. We discuss these below, complemented with annotations of major related pathways (but no attempt is made to comprehensively annotate all pathways) and processes. We base our annotations on data from well-studied model organisms, such as *S. cerevisiae, A. nidulans* and *Sch. pombe*, with orthology relationships to Agaricomycetes genes established on the basis of 1-to-1 reciprocal best Blast hits.

#### 4.3.1. Yeast beta oxidation

We reconstructed the beta-oxidation pathway (Table 4) in *C. cinerea* based on the yeast beta-oxidation pathway(van Roermund et al. 2003) (Fig. 5). The expression of genes comprising the beta-oxidation pathway in general does not appear to be widely dynamic in fruiting bodies. We found orthologs of yeast CTA1, a catalase A, to form a CDE orthogroup. However, this catalase in yeast is only a marginal component of the beta-oxidation pathway, and may be involved in a wide range of other functions (e.g. signaling). The more widespread developmental expression of ACS1 orthologs is remarkable (11/4 species at FC=2/4), for a possible connection with sterol biosynthesis (see above). ACS1 produces Acetyl-CoA, which is the starting point for ergosterol and membrane lipid biosynthesis.

**Table 4.** Summary of developmental expression dynamics of CDE orthogroups of fatty acid beta- oxidation related genes across 12 species. Protein ID of a representative protein is given follows by the number of species in which the orthogroup is developmentally regulated at fold change 2 and 4 (FC>2 and FC>4, respectively). Putative function and ortholog in *S. cerevisiae* (if any) are also given. Abbreviations: 0-gene absent, 1-gene present but not developmentally regulated, 2 - developmentally regulated at fold change >2, 4- developmentally regulated at fold change >4. Species names are abbreviated as: Cc – *C. cinerea*, Aa – *A. ampla*, Ab – *A. bisporus*, Ao – *A. ostoyae*, Ca – *C. aegerita*, Lb – *L. bicolor*, Le – *L. edodes*, Lt – *L. tigrinus*, Mk – *M. kentingensis*, Pc – *Ph. chrysosporium*, Po – *P. ostreatus*, Sc – *S. commune*.

| ID | Fold change |  | Putative function | S.<br>*cerevisiae*<br>ortholog | Species |  |  |  |  |  |  |  |  |  |  |  |
| --- | --- | --- | --- | --- | --- | --- | --- | --- | --- | --- | --- | --- | --- | --- | --- | --- |
|  | >2 | >4 |  |  | Cc | Aa | Ab | Ao | Ca | Lb | Le | Lt | Mk | Pc | Po | Sc |
| 437115 | 7 | 4 | General amino acid permease AGP2 | YBR132C | 4 | 2 | 4 | 1 | 4 | 2 | 4 | 0 | 1 | 0 | 1 | 2 |
| 336239 | 6 | 1 | Acyl-coenzyme A oxidase | YGL205W | 1 | 1 | 1 | 2 | 2 | 1 | 2 | 2 | 1 | 1 | 4 | 2 |
| 438072 | 10 | 5 | Acyl-CoA-binding protein | YGR037C | 4 | 2 | 4 | 4 | 2 | 2 | 2 | 4 | 2 | 1 | 4 | 1 |
| 365907 | 6 | 2 | 3-ketoacyl-CoA thiolase, peroxisomal | YIL160C | 4 | 1 | 2 | 2 | 4 | 2 | 0 | 1 | 2 | 1 | 1 | 1 |
| 231560 | 5 | 1 | Peroxisomal hydratase-dehydrogenase-epimerase | YKR009C | 2 | 1 | 1 | 4 | 2 | 1 | 2 | 1 | 2 | 1 | 1 | 1 |
| 354296 | 6 | 2 | Aspartate aminotransferase | YLR027C | 2 | 2 | 4 | 0 | 0 | 2 | 1 | 1 | 1 | 1 | 4 | 2 |
| 433737 | 8 | 3 | Peroxisomal 2,4-dienoyl-CoA reductase SPS19 [(3E)-enoyl-CoA-producing] | YNL202W | 4 | 1 | 4 | 2 | 2 | 4 | 1 | 1 | 2 | 1 | 2 | 2 |
| 374386 | 7 | 1 | Peroxisomal membrane protein PMP27 | YOL147C | 2 | 1 | 2 | 2 | 2 | 2 | 1 | 1 | 2 | 1 | 4 | 1 |
| 436931 | 6 | 2 | Delta(3,5)-Delta(2,4)-dienoyl-CoA isomerase | YOR180C | 2 | 1 | 2 | 1 | 0 | 4 | 1 | 2 | 0 | 1 | 4 | 2 |
| 401814 | 5 | 2 | Peroxisomal long-chain fatty acid import protein 2 | YPL147W | 4 | 1 | 1 | 2 | 4 | 1 | 1 | 1 | 2 | 1 | 2 | 1 |
| 395825 | 8 | 3 | Peroxisomal adenine nucleotide transporter 1 | YPR128C | 4 | 1 | 2 | 2 | 1 | 4 | 2 | 2 | 2 | 1 | 4 | 1 |
| 437261 | 6 | 4 | Long-chain-fatty-acid--CoA ligase 1 | YOR317W | 2 | 1 | 4 | 4 | 2 | 1 | 4 | 1 | 0 | 1 | 4 | 1 |

#### 4.3.2. Ergosterol biosynthesis

We reconstructed the ergosterol biosynthesis pathway based on orthology to known pathway member genes in *S. cerevisiae*(JM et al. 2012) (Table 5). ERG pathway genes of Agaricomycetes are generally higher expressed in fruiting bodies than in vegetative mycelia (VM/P1>1 in 20 out of 21 genes in *C. cinerea*), with expression peaking in young fruiting bodies of *C. cinerea* (Fig. 6, Table 5). Expression peaks at similar stages in other species too, in primordium 3/young fruiting body in *P. ostreatus* and young fruiting bodies/mature fruiting bodies stages in *A. ostoyae* (Supplementary Fig. 4). Similar pictures were obtained for other species. It should be noted that although the ergosterol biosynthesis pathway shows clear expression trends, the dynamics of expression is relatively low, mostly between 2<FC<4. Therefore, ergosterol biosynthesis genes mostly do not form CDE orthogroups, nevertheless, the expression patterns are distinctive so they warrant discussion here.

**Fig. 6.**
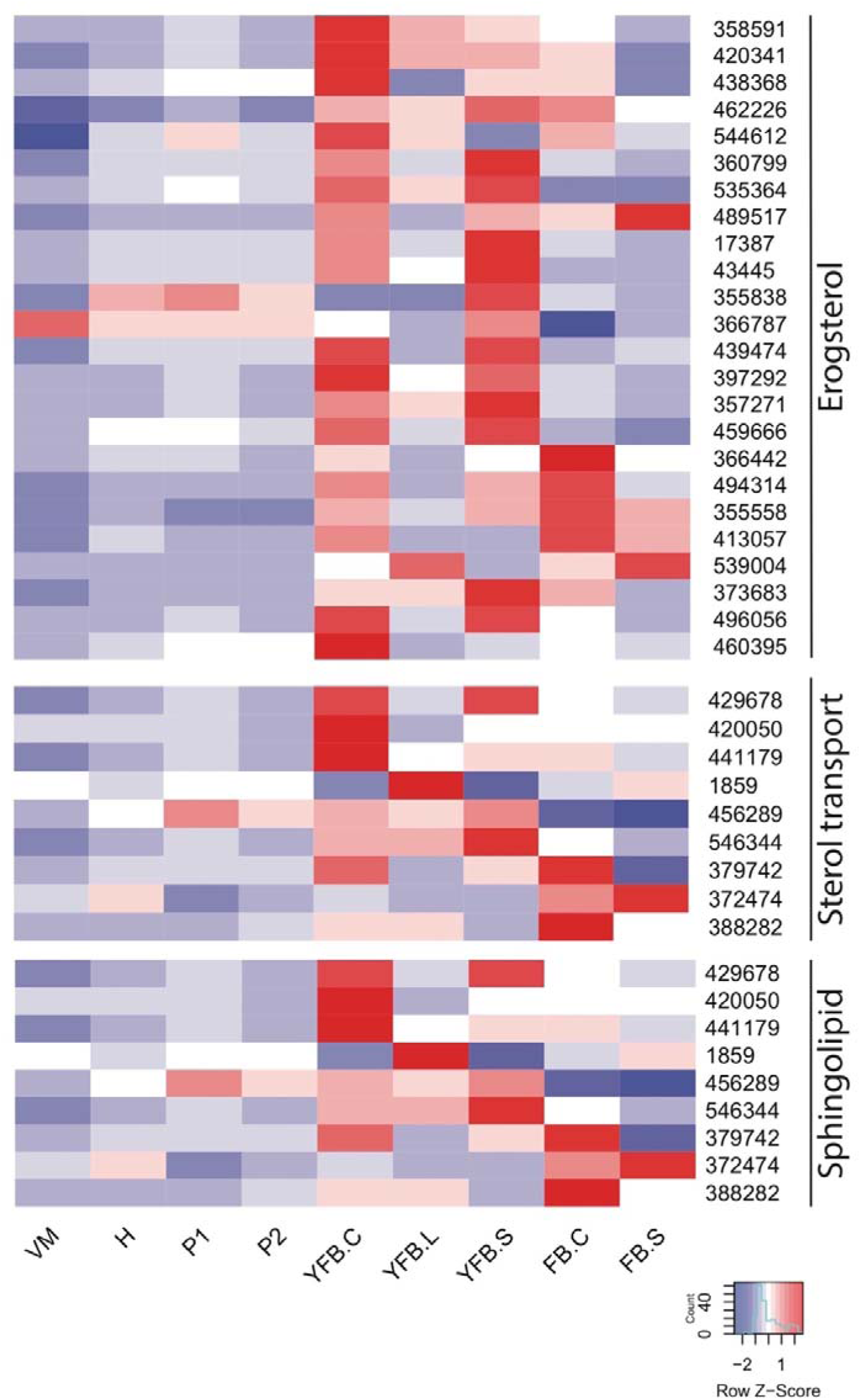
Expression heatmap of ergosterol and sphingolipid biosynthesis related genes in *C. cinerea*. Genes are denoted by Protein IDs. Blue and red colors represent low and high expression, respectively. Developmental stages are abbreviated as follows: VM – vegetative mycelium, H – hyphal knot, P1 – stage 1 primordium, P2 – stage 2 primordium, YFB.C – young fruiting body cap, YFB.L – young fruiting body gills, YFB.S – young fruiting body stipe, FB.C – mature fruiting body cap (including gills), FB.S – mature fruiting body stipe.

**Table 5.** Summary of developmental expression dynamics of CDE orthogroups of ergosterol and sphingolipid biosynthesis related genes across 12 species. Protein ID of a representative protein is given follows by the number of species in which the orthogroup is developmentally regulated at fold change 2 and 4 (FC>2 and FC>4, respectively). Putative function and ortholog in *S. cerevisiae* (if any) are also given. Abbreviations: 0-gene absent, 1-gene present but not developmentally regulated, 2 - developmentally regulated at fold change >2, 4- developmentally regulated at fold change >4. Species names are abbreviated as: Cc – *C. cinerea*, Aa – *A. ampla*, Ab – *A. bisporus*, Ao – *A. ostoyae*, Ca – *C. aegerita*, Lb – *L. bicolor*, Le – *L. edodes*, Lt – *L. tigrinus*, Mk – *M. kentingensis*, Pc – *Ph. chrysosporium*, Po – *P. ostreatus*, Sc – *S. commune*.

| ID | Fold change |  | Putative function | S.<br>*cerevisiae*<br>ortholog | Species |  |  |  |  |  |  |  |  |  |  |  |
| --- | --- | --- | --- | --- | --- | --- | --- | --- | --- | --- | --- | --- | --- | --- | --- | --- |
|  | >2 | >4 |  |  | Cc | Aa | Ab | Ao | Ca | Lb | Le | Lt | Mk | Pc | Po | Sc |
| 488426 | 9 | 3 | Transcription factor | – | 4 | 1 | 0 | 4 | 2 | 2 | 4 | 2 | 2 | 2 | 2 | 1 |
| Late pathway |  |  |  |  |  |  |  |  |  |  |  |  |  |  |  |  |
| 366787 | 9 | 4 | Ergosterol biosynthetic protein 28 | YER044C | 2 | 1 | 2 | 2 | 4 | 2 | 4 | 4 | 2 | 1 | 4 | 1 |
| 366442 | 11 | 6 | Sterol-4-alpha-carboxylate 3-dehydrogenase, decarboxylating | YGL001C | 4 | 2 | 4 | 4 | 2 | 4 | 2 | 4 | 2 | 2 | 4 | 1 |
| 544612 | 10 | 7 | Delta(24(24(1)))-sterol reductase | YGL012W | 4 | 2 | 4 | 2 | 0 | 4 | 4 | 4 | 2 | 4 | 4 | 1 |
| 496056 | 10 | 7 | Methylsterol monooxygenase | YGR060W | 4 | 1 | 4 | 2 | 4 | 4 | 2 | 4 | 2 | 4 | 4 | 1 |
| 489517 | 9 | 6 | Squalene monooxygenase | YGR175C | 4 | 1 | 2 | 2 | 0 | 4 | 4 | 4 | 4 | 2 | 4 | 1 |
| 397292 | 11 | 7 | Lanosterol 14-alpha demethylase | YHR007C | 4 | 2 | 4 | 2 | 2 | 4 | 4 | 4 | 2 | 4 | 4 | 1 |
| 413057 | 6 | 3 | NADPH--cytochrome P450 reductase | YHR042W | 4 | 1 | 4 | 2 | 4 | 1 | 1 | 2 | 1 | 1 | 2 | 1 |
| 420341 | 8 | 5 | Lanosterol synthase | YHR072W | 4 | 2 | 4 | 2 | 0 | 1 | 4 | 4 | 1 | 2 | 4 | 1 |
| 494314 | 7 | 4 | Delta(7)-sterol 5(6)-desaturase | YLR056W | 4 | 2 | 4 | 1 | 2 | 2 | 4 | 1 | 1 | 4 | 1 | 1 |
| 462226 | 8 | 4 | 3-keto-steroid reductase | YLR100W | 4 | 1 | 4 | 2 | 0 | 1 | 4 | 2 | 2 | 1 | 4 | 2 |
| 373683 | 10 | 7 | Sterol 24-C-methyltransferase | YML008C | 4 | 1 | 4 | 2 | 4 | 4 | 2 | 4 | 4 | 4 | 2 | 1 |
| 357271 | 10 | 6 | Cytochrome P450 61 | YMR015C | 4 | 0 | 4 | 2 | 2 | 4 | 2 | 4 | 2 | 4 | 4 | 1 |
| 17387 | 11 | 7 | C-8 sterol isomerase | YMR202W | 4 | 2 | 4 | 2 | 2 | 4 | 2 | 4 | 4 | 4 | 4 | 1 |
| 535364 | 10 | 3 | Delta(14)-sterol reductase | YNL280C | 4 | 2 | 2 | 2 | 1 | 2 | 4 | 2 | 2 | 2 | 4 | 1 |
| Mevalonate pathway |  |  |  |  |  |  |  |  |  |  |  |  |  |  |  |  |
| 358591 | 7 | 2 | Cystathionine beta-lyase | YGL184C | 4 | 4 | 1 | 1 | 1 | 1 | 2 | 2 | 2 | 2 | 2 | 1 |
| 43445 | 9 | 6 | Squalene synthase | YHR190W | 4 | 2 | 4 | 4 | 0 | 4 | 2 | 4 | 2 | 1 | 4 | 1 |
| 459666 | 9 | 5 | Farnesyl pyrophosphate synthase | YJL167W | 4 | 2 | 2 | 1 | 2 | 2 | 4 | 4 | 4 | 1 | 4 | 1 |
| 360799 | 8 | 2 | 3-hydroxy-3-methylglutaryl-coenzyme A reductase 2 | YLR450W | 4 | 1 | 2 | 2 | 2 | 2 | 1 | 2 | 1 | 2 | 4 | 1 |
| 439474 | 12 | 5 | Hydroxymethylglutaryl-CoA synthase | YML126C | 4 | 2 | 4 | 2 | 4 | 4 | 2 | 4 | 2 | 2 | 2 | 2 |
| 355838 | 12 | 10 | Phosphomevalonate kinase | YMR220W | 4 | 4 | 4 | 4 | 4 | 4 | 4 | 4 | 2 | 4 | 4 | 2 |
| 438368 | 8 | 2 | Diphosphomevalonate decarboxylase | YNR043W | 2 | 2 | 2 | 1 | 2 | 1 | 2 | 2 | 1 | 4 | 4 | 1 |
| 355558 | 7 | 0 | Acetyl-CoA acetyltransferase | YPL028W | 2 | 2 | 1 | 2 | 1 | 2 | 1 | 2 | 1 | 2 | 2 | 1 |
| 460395 | 6 | 2 | Isopentenyl-diphosphate Delta-isomerase | YPL117C | 4 | 1 | 1 | 2 | 1 | 1 | 2 | 4 | 1 | 2 | 2 | 1 |
| 539004 | 12 | 7 | 3-oxo-5-alpha-steroid 4-dehydrogenase | – | 4 | 4 | 2 | 4 | 2 | 4 | 4 | 2 | 2 | 2 | 4 | 4 |
| Sphingolipid biosynthesis |  |  |  |  |  |  |  |  |  |  |  |  |  |  |  |  |
| 388282 | 9 | 5 | Alkaline ceramidase YPC1 | YBR183W | 4 | 2 | 4 | 4 | 2 | 2 | 1 | 4 | 2 | 1 | 4 | 1 |
| 429678 | 6 | 1 | Serine palmitoyltransferase 2 | YDR062W | 4 | 1 | 2 | 1 | 2 | 2 | 1 | 2 | 1 | 1 | 2 | 1 |
| 372474 | 7 | 1 | Sphingosine-1-phosphate lyase | YDR294C | 2 | 1 | 2 | 4 | 2 | 1 | 2 | 2 | 1 | 1 | 2 | 1 |
| 379742 | 7 | 0 | Sphingolipid C4-hydroxylase SUR2 | YDR297W | 2 | 1 | 1 | 2 | 0 | 2 | 2 | 1 | 2 | 1 | 2 | 2 |
| 456289 | 8 | 0 | Ceramide synthase LAG1 | YHL003C | 1 | 2 | 0 | 2 | 2 | 1 | 2 | 2 | 2 | 1 | 2 | 2 |
| 420050 | 6 | 3 | Dihydrosphingosine 1-phosphate phosphatase LCB3 | YJL134W | 4 | 1 | 1 | 4 | 1 | 2 | 2 | 1 | 1 | 1 | 4 | 2 |
| 441179 | 7 | 2 | Ceramide very long chain fatty acid hydroxylase SCS7 | YMR272C | 4 | 1 | 2 | 2 | 2 | 4 | 2 | 1 | 1 | 1 | 2 | 1 |
| 546344 | 6 | 3 | Serine palmitoyltransferase 1 | YMR296C | 4 | 1 | 2 | 1 | 4 | 1 | 2 | 1 | 2 | 1 | 4 | 1 |
| 1859 | 6 | 2 | Sphingoid long chain base kinase 4 | YOR171C | 4 | 1 | 1 | 1 | 1 | 2 | 4 | 2 | 2 | 1 | 2 | 1 |

In these species expression peaks seem to coincide with intense growth by cell expansion in fruiting bodies. Above we showed that mitotic, ribosomal and histone gene expression level off in late primordium stages (P2 in *C. cinerea*, see Fig. 3), after which growth by cell expansion takes place. This might create an increased demand for membrane constituents during rapid cell expansion/growth by turgor, which is what we might see in the expression patterns of genes in the ergosterol biosynthesis pathway (Fig. 6, Supplementary Fig. 4). This hypothesis would also explain why there is no clear peak in *C. neoformans* and *Pt. gracilis*, where sudden growth by cell expansion is lacking.

Similarly, a homolog of the Sterol-Regulatory Element Binding Proteins (SREBP(CM and PJ 2010)), similar to the *S. cerevisiae* transcription factor HMS1 shows a very similar expression pattern: in *C. cinerea* (*C. cinerea* protein ID: 488426) expression peak in YFB-S and YFB-C (Fig. 6), whereas in *P. ostreatus* it peaks in mature fruiting body tissues (PleosPC15_2_174910). This suggests, based on expression data, that these putative SREBPs might be regulating ergosterol biosynthesis genes also in the Agaricomycetes.

A connection with the above-mentioned Ac-CoA producing basic metabolic pathways, where certain, but not all, components were found widely upregulated, is possible here. These pathways produce Acetyl-CoA, which is the starting point of ergosterol biosynthesis. This raises the possibility that the upregulation of the ethanol utilization pathway components (see above, Fig. 5) and certain components of the glyoxylate cycle reflect increased Ac-CoA production which is required as input to ergosterol biosynthesis and eventually membrane assembly. A competing hypothesis was put forth in *P. microcarpus*, where upregulation of glyoxylate shunt genes was observed and suggested that it underlies the production of storage materials for spores(Pereira et al. 2017). Using our more resolved transcriptome data and based on the observation of ergosterol pathway expression dynamics, we propose that the connection to the glyoxylate shunt might represent a link to membrane constituent synthesis, although more data would be needed to verify this hypothesis.

The upregulation of the ergosterol synthesis pathway was observed in *F. velutipes*, where most ergosterol biosynthesis genes were upregulated in the young fruiting body stage(Wang et al. 2019) (approximately in agreement with our definition of young fruiting bodies) and showing intense growth. Sterol metabolism genes were also found to be more abundant in fruiting bodies of *H. erinaceus* compared to vegetative mycelia(Zeng et al. 2018). It was reported that orthologs of the lanosterol-alpha-demethylase Erg11 of *S. cerevisiae*, a component of the ergosterol biosynthesis pathway (corresponding *C. cinerea* ortholog: 397292), was upregulated in fruiting bodies of *Antrodia cinnamomea*(Lu et al. 2014) and *P. ostreatu*s(Sunagawa and Magae 2005). This is consistent with the observations made in this study.

We also checked the sphingolipid biosynthesis pathway based on orthology to pathway members in *S. cerevisiae*. Sphingolipids, along with sterols are important in membrane rafts and microdomains and form key components of fungal cell membranes. Although some yeast genes did not have clear orthologs in Agaricomycetes, genes involved in sphingolipid biosynthesis showed a similar expression profile in *C. cinerea* (Fig. 6, Table 5). This may underscore our hypothesis on the link between growth by cell expansion and membrane assembly.

A characteristic CDE orthogroup of putative C14-sterol reductase genes was developmentally expressed in 12/7 species (represented by *C. cinerea* 539004)(Table 5, Fig. 6). These proteins are not part of the ergosterol biosynthesis pathway, but show a high similarity to *A. nidulans* and *C. albicans* membrane phospholipid metabolism proteins AN2040 and Erg24, respectively, which are involved in ergosterol metabolism. Genes in this orthogroup show peak expression in late development, coincident with rapid expansion of fruiting bodies. In several species they were also upregulated in primordia relative to vegetative mycelium samples. These expression patterns resemble that of ergosterol biosynthesis genes, and suggest that this orthogroup may be linked to ergosterol metabolism.

#### 4.3.3. Sterol transport

We detected several CDE orthogroups that were annotated as related to ergosterol transport and showed similar upregulation in late development as ergosterol biosynthesis genes did. We therefore inventoried sterol transport related genes of the families Niemann-Pick type C protein family (*S. cerevisiae* Npc2), the yeast Ncr1 family, the nonspecific sterol transport protein (Scp2) family, StAR-related lipid-transfer (START) proteins as well as members of the Oxysterol Binding Protein Homologue (OSH) protein family(Schulz and Prinz 2007).

Of these, the Ncr1, Npc2, Scp2 and START gene families are single-copy families in mushroom-forming fungi, whereas the OSH family is represented by 5 members in all mushroom-forming fungi (except *S. commune* which has 6, Table 6). Npc2 (ortholog *C. cinerea* 475807), Scp2 (ortholog *C. cinerea* 478741) and START proteins as well as one member of the OSH family were (*C. cinerea* protein ID: 427232)(Table 6) upregulated during the second half of development, at or after the young fruiting body stage. This is similar to the expression patterns seen in ergosterol biosynthesis genes. One member of the OSH family orthogroup (*C. cinerea* protein ID: 378276) showed a clearly stipe-specific expression, with peaks in stipes of young fruiting bodies or fruiting bodies of *C. cinerea, A. ostoyae, M. kentingensis, A. bisporus and P. ostreatus,* albeit mostly with low fold change values (2<FC<4). Interestingly, the corresponding gene was cap-upregulated in *L. bicolor* (Lacbi2_456536), whereas in simpler species (*S. commune, A. ampla*), it showed diverse expression peaks. Ncr1 orthologs (represented by *C. cinerea* 494709) showed consistently higher expression in fruiting bodies than in vegetative mycelia. Fold changes were generally low in these gene families, but nevertheless the patterns are phylogenetically very consistent.

**Table 6.** Summary of developmental expression dynamics of CDE orthogroups of sterol transport related genes across 12 species. Protein ID of a representative protein is given follows by the number of species in which the orthogroup is developmentally regulated at fold change 2 and 4 (FC>2 and FC>4, respectively). Putative function and ortholog in *S. cerevisiae* (if any) are also given. Abbreviations: 0-gene absent, 1-gene present but not developmentally regulated, 2 - developmentally regulated at fold change >2, 4- developmentally regulated at fold change >4. Species names are abbreviated as: Cc – *C. cinerea*, Aa – *A. ampla*, Ab – *A. bisporus*, Ao – *A. ostoyae*, Ca – *C. aegerita*, Lb – *L. bicolor*, Le – *L. edodes*, Lt – *L. tigrinus*, Mk – *M. kentingensis*, Pc – *Ph. chrysosporium*, Po – *P. ostreatus*, Sc – *S. commune*.

| ID | Fold change |  | Putative function | S.<br>*cerevisiae*<br>ortholog | Species |  |  |  |  |  |  |  |  |  |  |  |
| --- | --- | --- | --- | --- | --- | --- | --- | --- | --- | --- | --- | --- | --- | --- | --- | --- |
|  | >2 | >4 |  |  | Cc | Aa | Ab | Ao | Ca | Lb | Le | Lt | Mk | Pc | Po | Sc |
| 470387 | 7 | 6 | Prefoldin subunit 2 | – | 4 | 4 | 4 | 4 | 1 | 2 | 4 | 0 | 1 | 0 | 4 | 0 |
| 475807 | 7 | 5 | Phosphatidylglycerol/phosphatidyl inositol transfer protein | YDL046W | 2 | 1 | 4 | 4 | 0 | 4 | 1 | 1 | 4 | 1 | 4 | 2 |
| 378276 | 6 | 1 | Oxysterol-binding protein homolog 7 | YHR001W | 4 | 1 | 2 | 2 | 1 | 1 | 1 | 1 | 2 | 1 | 2 | 2 |
| 494709 | 7 | 1 | NPC intracellular sterol transporter 1-related protein 1 | YPL006W | 4 | 2 | 1 | 2 | 2 | 1 | 2 | 1 | 1 | 1 | 2 | 2 |
| 427232 | 10 | 2 | Oxysterol-binding protein homolog 4 | YPL145C | 4 | 2 | 2 | 4 | 2 | 1 | 2 | 2 | 2 | 1 | 2 | 2 |
| 478741 | 6 | 2 | SCP2 sterol transport protein family | – | 2 | 1 | 2 | 4 | 1 | 2 | 0 | 1 | 2 | 1 | 4 | 1 |

These expression patterns resemble those of ergosterol biosynthesis genes, which also showed upregulation in mid- to late development, coincident with cell expansion and growth. This suggests that ergosterol transport, in addition to biosynthesis, is an important process in the second half of the development of mushroom fruiting bodies. Like synthesis, transport of sterols might be required to fulfill the needs of rapid cell expansion, which obviously coincides with cell- and membrane surface expansion too.

Although marginally related, we mention here the CAP domain/Pry1 protein family, which includes known sterol-binding proteins. The family has been associated with diverse functionalities, including cell adhesion, pathogenicity (pathogenicity-related PR proteins belong here), morphogenesis, endocrine and paracrine functions, among others(GM et al. 2008; Schneiter and Pietro 2013). This protein family includes *S. cerevisiae* Pry1, a lipid and sterol binding protein involved in sterol export(Darwiche et al. 2017). To our best knowledge, there are no previous reports of this family in fruiting body development, whereas, their role in morphogenesis in mammals is well-established(GM et al. 2008). We detected, on average 2-5 copies of CAP proteins in the Agaricomycetes, with expansions up to 9 in *Pt. gracilis, L. bicolor* and *A. ostoyae* as well as a CDE orthogroup of Pry1-related CAP-domain proteins (represented by *C. cinerea* 470387, FC2/4: 7/6, Table 6, Fig. 6). Members of this orthogroup showed peak expression late in development, at or after the young fruiting body stage in most species. The sterol transporting ability and expression patterns suggest that this family may have a role in sterol transport during the second half of development in Agaricomycetes fruiting bodies. It should be noted that neither Polyporales species (*Ph. chrysosporium* and *L. tigrinus*) had developmentally regulated genes in this family, indicating that their role, if any, may not be universal.

#### 4.3.4. Phospholipid and fatty acid biosynthesis

The phospholipid and fatty acid biosynthesis pathways are conserved in Agaricomycetes, with most genes showing clear 1-to-1 orthologs to described pathway members of *S. cerevisiae (*Table 7). Because active cell growth and expansion requires an excess amount of membrane components (ergosterol, phospholipids), we examined pathways involved in membrane phospholipid and triacylglycerol synthesis. We reconstructed these pathways based on well-known reaction steps in *S. cerevisiae* and *Yarrowia lipolytica*(Carman and Han 2011; Fakas 2017). The first half of the pathway, which starts from glycerol-3-phosphate and leads to the production of phosphatidic acid, shows upregulation in young and mature fruiting bodies of *C. cinerea* and *A. ostoyae* (Fig. 7). Phosphatidic acid sits in a branching point of the pathway, with one branch leading to membrane phospholipids (CDP-DAG pathway), whereas the other leads to triacylglycerol (TAG), a major storage material. The CDP-DAG pathway shows a general upregulation in the second half of development (young and mature fruiting bodies) in *C. cinerea, A. ostoyae, M. kentingensis, A. bisporus* (Supplementary Fig. 5). These patterns are consistent with increased membrane lipid biosynthesis in these stages, possibly serving the needs of increasing cell volumes during rapid growth. On the other hand, TAG pathway members appear upregulated somewhat later, mostly in mature fruiting bodies in most species (Fig. 7, Supplementary Fig. 5), which coincides with spore formation and could be related to storage material synthesis in spores. In *P. ostreatus*, both the CDP-DAG and TAG pathways show remarkably uniform upregulation in primordium stages (Fig. 7), which is quite different from the pattern seen in other species. This pattern in *P. ostreatus* is also observable in fatty acid biosynthesis pathways (see below).

**Fig. 7.**
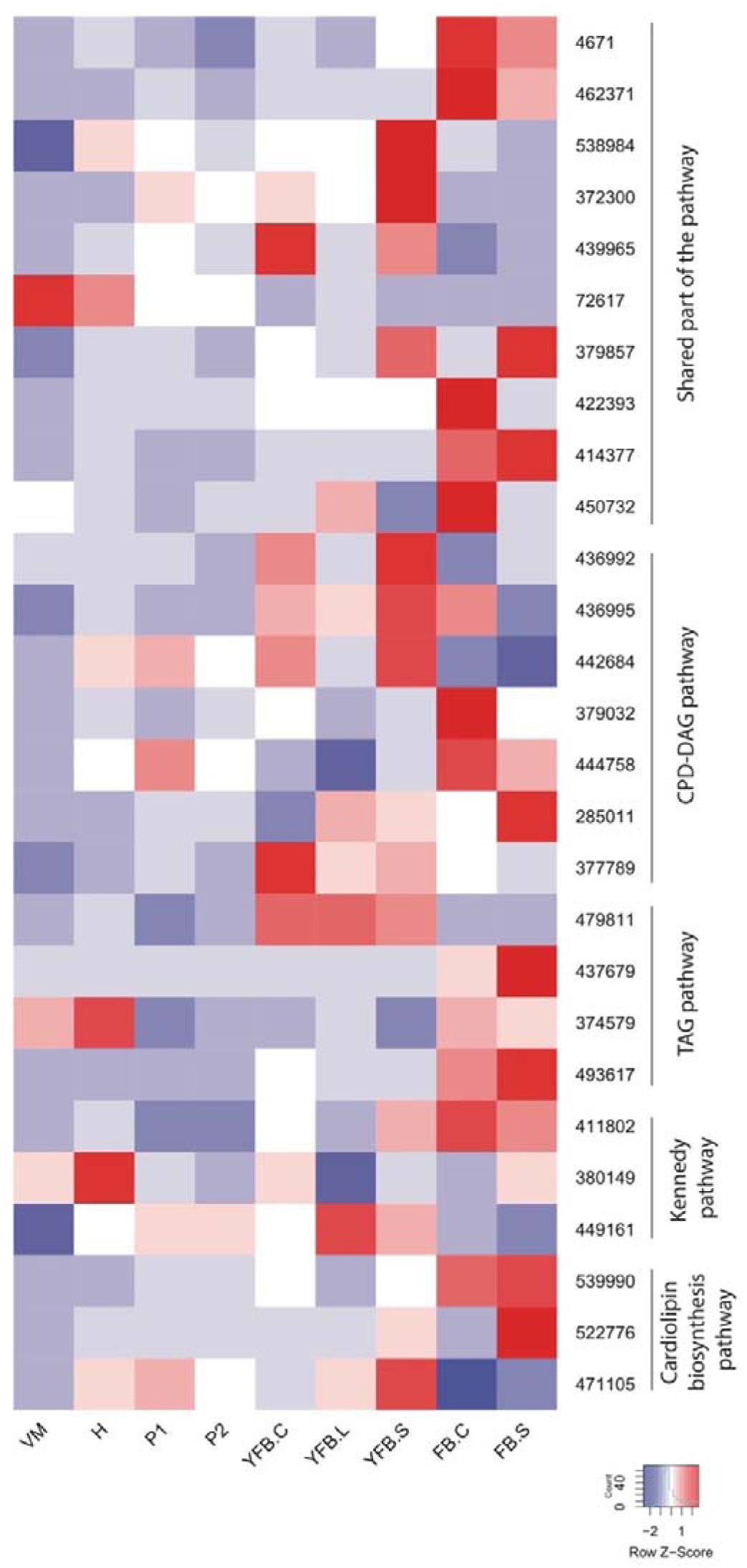
Expression heatmap of membrane phospholipid and fatty acid biosynthetic genes in *C. cinerea*. Genes are denoted by Protein IDs. Blue and red colors represent low and high expression, respectively. Developmental stages are abbreviated as follows: VM – vegetative mycelium, H – hyphal knot, P1 – stage 1 primordium, P2 – stage 2 primordium, YFB.C – young fruiting body cap, YFB.L – young fruiting body gills, YFB.S – young fruiting body stipe, FB.C – mature fruiting body cap (including gills), FB.S – mature fruiting body stipe.

**Table 7.**
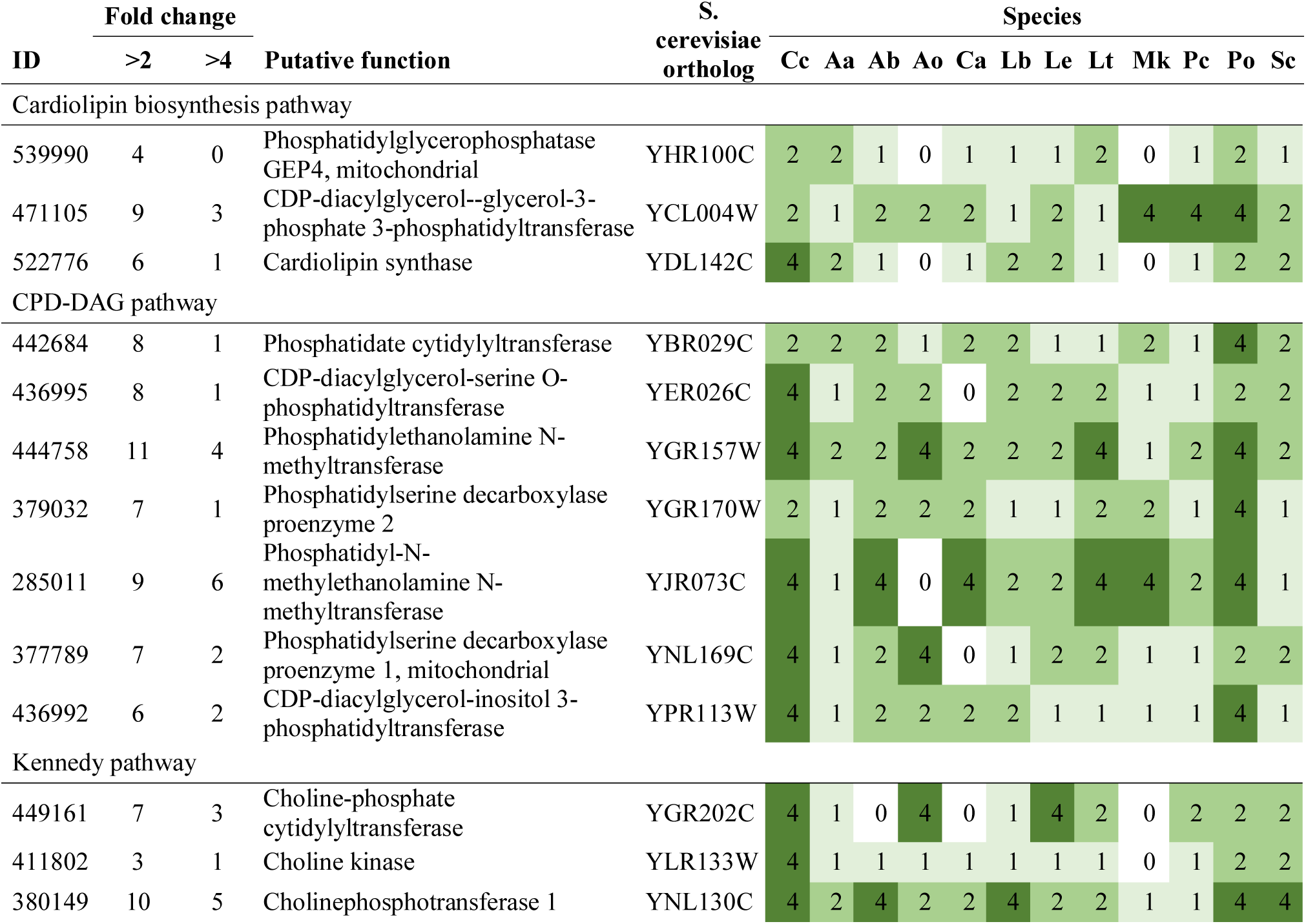
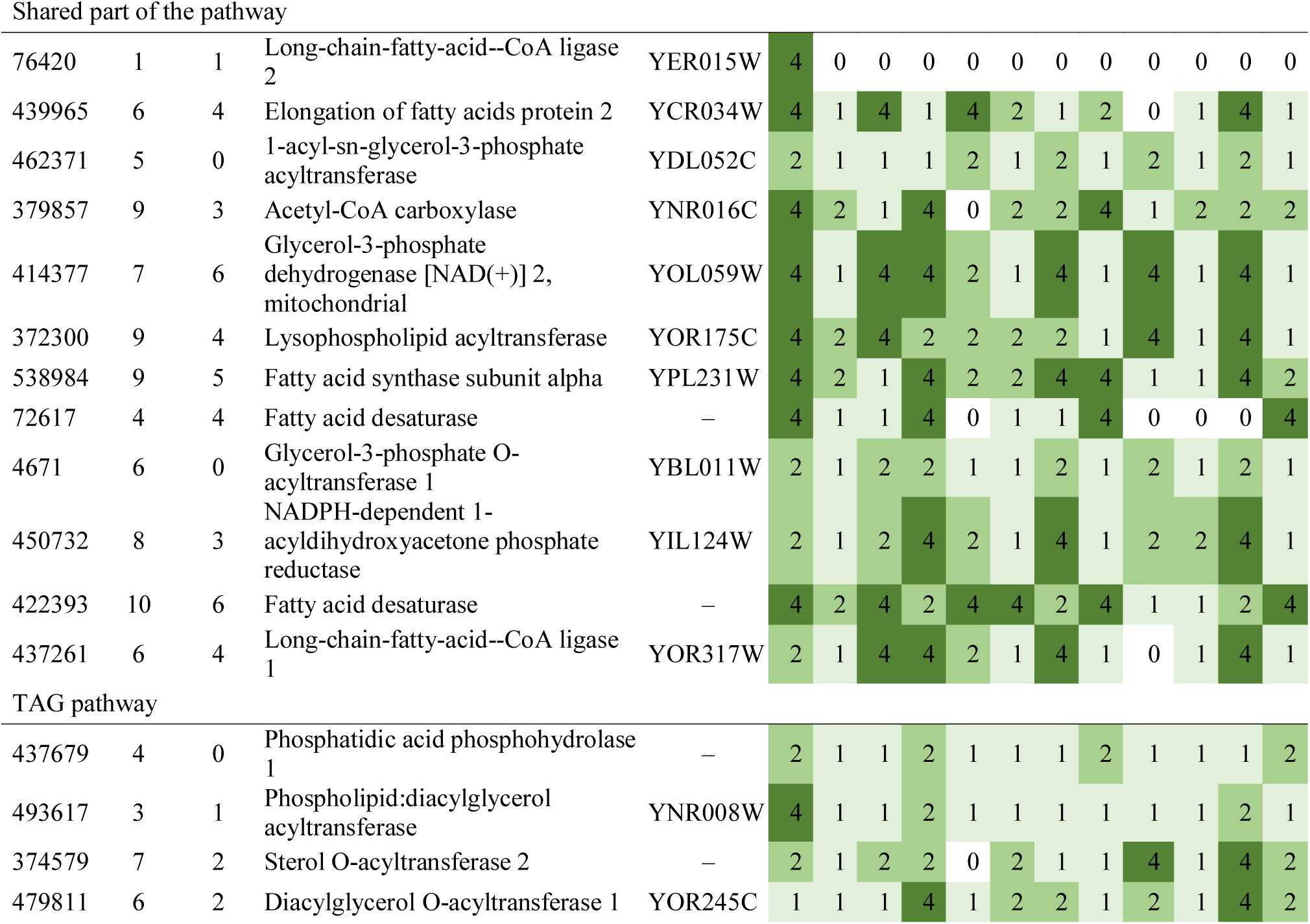
Summary of developmental expression dynamics of CDE orthogroups of membrane phoshpolipid and fatty acid biosynthesis genes genes across 12 species. Protein ID of a representative protein is given follows by the number of species in which the orthogroup is developmentally regulated at fold change 2 and 4 (FC>2 and FC>4, respectively). Putative function and ortholog in *S. cerevisiae* (if any) are also given. Abbreviations: 0-gene absent, 1-gene present but not developmentally regulated, 2 - developmentally regulated at fold change >2, 4- developmentally regulated at fold change >4. Species names are abbreviated as: Cc – *C. cinerea*, Aa – *A. ampla*, Ab – *A. bisporus*, Ao – *A. ostoyae*, Ca – *C. aegerita*, Lb – *L. bicolor*, Le – *L. edodes*, Lt – *L. tigrinus*, Mk – *M. kentingensis*, Pc – *Ph. chrysosporium*, Po – *P. ostreatus*, Sc – *S. commune*.

Similar to the CDP-DAG and TAG pathways, fatty acid biosynthesis (Table 7) shows a signal for upregulation in young fruiting body tissues, with *C. cinerea* orthologs of *S. cerevisiae* Acc1, Fas2, Elo2 and Faa1 showing peak expression in the YFB stage. Similar patterns exist in *A. ostoyae, M. kentingensis* and to some extent in *A. bisporus* (Fig. 7). In *P. ostreatus*, fatty acid biosynthesis genes are co-expressed with the other phospholipid biosynthesis genes, showing an upregulation in primordia.

Both phospholipid synthesis and fatty acid biosynthesis require Acetyl-CoA as a starting point. For the former, it is needed for acylation reactions in the conversion of lysophosphatidic acid to phosphatidic acid (last element of the shared part of the pathway) whereas in the latter, Ac-CoA is the starting point of fatty acid synthesis(Fakas 2017). This provides another link with the dynamics of Ac-CoA biosynthesis genes noted above. As in the case of ergosterol biosynthesis pathway members, phospholipid and fatty acid metabolism genes showed low expression dynamics (and hence did not form CDE orthogroups), but their expression patterns were consistent across stages and species.

The expression of mitochondrial membrane components (cardiolipin and phosphatidylglycerol) is also peaking in young fruiting bodies, mostly in stipe, but not very characteristic (Fig. 7).

#### 4.3.5. Linoleic acid producing fatty acid desaturases

Several CDE orthogroups of fatty acid desaturases (FAD) were detected among conserved developmentally expressed genes. Members of these orthogroups show high similarity to *C. albicans* Fad2 and Fad3 and *A. nidulans* OdeA, which are involved in the synthesis of alpha-linoleic acid. Linoleic acid is a major membrane component in fungi, which, as an unsaturated fatty acid, regulates membrane fluidity. It has also been speculated to be the starting point for the biosynthesis of volatiles, such as 1-octen-3-ol(A et al. 2020). Linoleic acid showed increased proportions in fruiting bodies and reproductive structures in general in both the Asco- and Basidiomycota(SHAW 1967; Goodrich-Tanrikulu et al. 1998; H and S 2003; Gessler et al. 2017; Song et al. 2018a). Linoleic acid is also a precursor of oxylipins, key developmental signaling molecules that are actively produced by reproductive cells(Combet et al. 2006; Gessler et al. 2017; Orban et al. 2021). Accordingly, in the Ascomycota, several fatty acid desaturase gene mutations were reported to asexual/sexual defects(Pöggeler et al. 2006b). The higher proportion of linoleic acid in fruiting bodies of Agaricomycetes was already noted in 1967(SHAW 1967) with several similar reports(H and S 2003; Song et al. 2018a) later. The proportion of unsaturated fatty acid (which include linoleic acid) increases during fruiting body formation(H and S 2003) in *L. edodes*. Three fatty acid desaturases, two in *L. edodes* (Le.Fad1 and Le.Fad2(H and S 2005)) and one in *C. cinerea* (*Cc.odeA*(H and S 2003, 2005; S et al. 2007b)) have been shown to be capable of producing linoleic acid. Fad genes were found to be differentially expressed in fruiting bodies of *A. bisporus* previously(Morin et al. 2012).

We found 7-16 fatty acid desaturase genes in the examined Agaricomycetes, many of which were developmentally regulated, although clear orthology of these genes to experimentally characterized Ascomycota genes is missing. Each species had at least 2-3 FAD encoding genes upregulated in primordia relative to vegetative mycelium, suggesting that their functions are key to the initiation of fruiting body development. FAD genes formed five CDE orthogroups (Table 8). One of these (*C. cinerea* protein ID: 363743) was FB-init in most species (*C. cinerea, A. ostoyae, A. ampla, M. kentingensis, P. ostreatus, S. commune, C. aegerita, L. bicolor*), showed high expression in primordium stages, then expression levels dropped significantly in later developmental stages. The CDE orthogroup containing Le.FAD1 (Lenedo1_1050661) was upregulated in primordia of most species relative to mycelium samples (*C. aegerita, A. ampla, S. commune, L. edodes. L. bicolor, L. tigrinus, M. kentingensis,* but not *A. ostoyae, P. ostreatus* and *A. bisporus,* ortholog missing in *C. cinerea)*, although only slightly (fold change = 1.5-2x, except in *L. edodes* and *Pt. gracilis* which showed >4-fold upregulation). The orthogroup encompassing *Cc.odeA* (*C. cinerea* protein ID: 422393) and Le.Fad2 (orthogroup represented by *S. commune* 2511002) was upregulated in primordia relative to vegetative mycelia of several species (*A. ampla, C. aegerita, S. commune, C. cinerea, L. edodes, L. tigrinus* and *P. ostreatus*, but not in *A. ostoyae, M. kentingensis, Pt. gracilis* and *A. bisporus*). Our expression data are consistent with previous reports of transcripts/protein abundance in case of both Le.FAD1 and Le.FAD2 in *L. edodes*(H and S 2003, 2005). The other two FAD orthogroups showed less conservation in expression (*C. cinerea* protein ID: 471478, 422976, Fig. 8), including peaks in later developmental stages, such as young fruiting body caps.

**Fig. 8.**
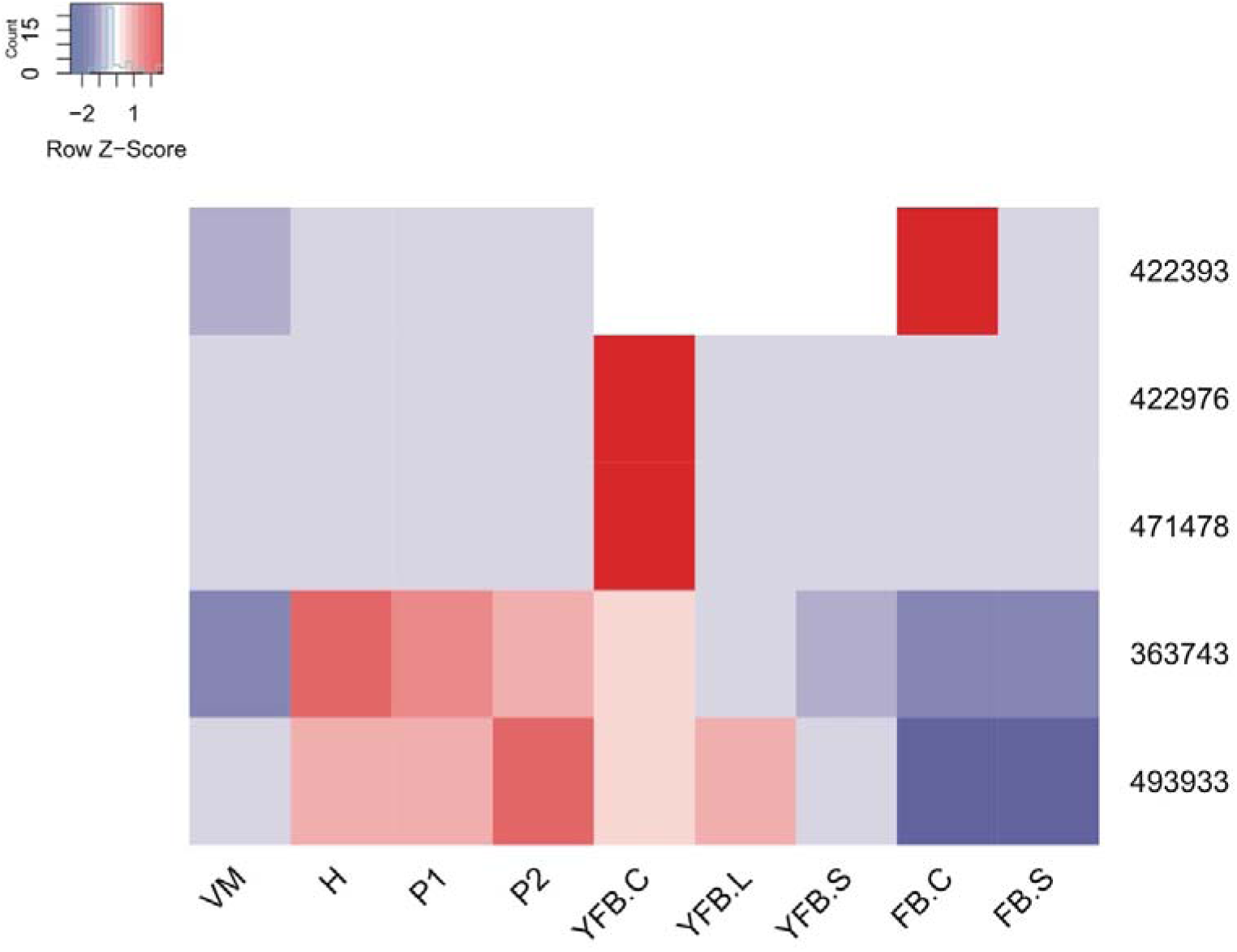
Expression heatmap of putatively linoleic acid biosynthesis related fatty acid desaturase genes in *C. cinerea*. Genes are denoted by Protein IDs. Blue and red colors represent low and high expression, respectively. Developmental stages are abbreviated as follows: VM – vegetative mycelium, H – hyphal knot, P1 – stage 1 primordium, P2 – stage 2 primordium, YFB.C – young fruiting body cap, YFB.L – young fruiting body gills, YFB.S – young fruiting body stipe, FB.C – mature fruiting body cap (including gills), FB.S – mature fruiting body stipe.

**Table 8.**
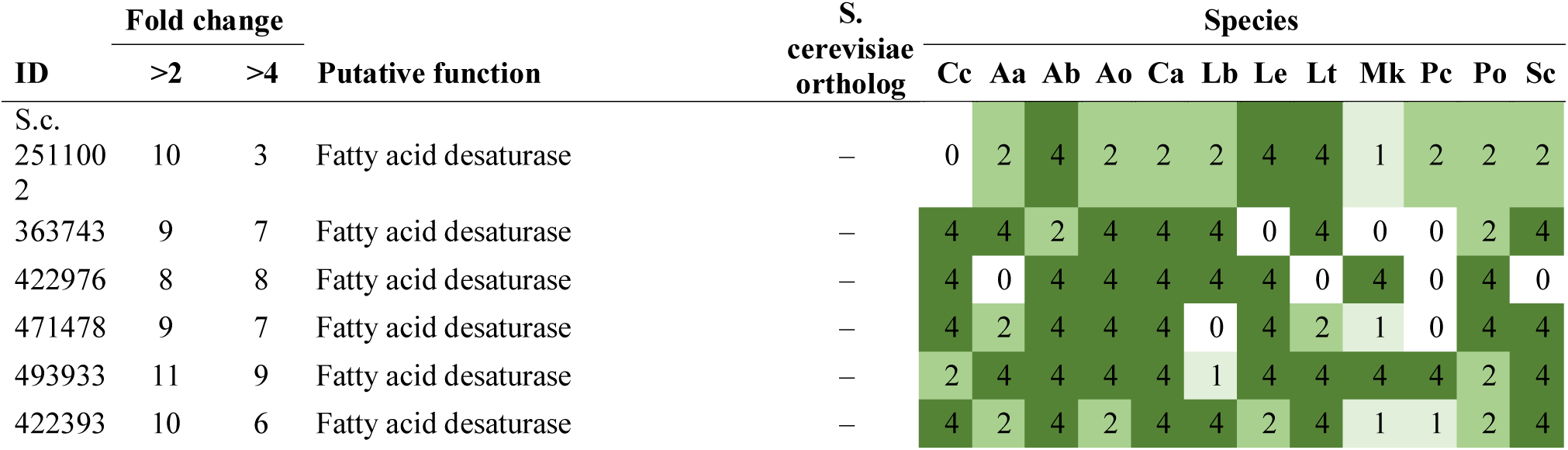
Summary of developmental expression dynamics of CDE orthogroups of linoleic acid biosynthesis related genes across 12 species. Protein ID of a representative protein is given follows by the number of species in which the orthogroup is developmentally regulated at fold change 2 and 4 (FC>2 and FC>4, respectively). Putative function and ortholog in *S. cerevisia*e (if any) are also given. Abbreviations: 0-gene absent, 1-gene present but not developmentally regulated, 2 - developmentally regulated at fold change >2, 4- developmentally regulated at fold change >4. Species names are abbreviated as: Cc – *C. cinerea*, Aa – *A. ampla*, Ab – *A. bisporus*, Ao – *A. ostoyae*, Ca – *C. aegerita*, Lb – *L. bicolor*, Le – *L. edodes*, Lt – *L. tigrinus*, Mk – *M. kentingensis*, Pc – *Ph. chrysosporium*, Po – *P. ostreatus*, Sc – *S. commune*.

The linoleic acid-producing ability of the proteins encoded by these genes suggests that the detected CDE orthogroups and expression dynamics relate to linoleic acid production. It follows from our observations of expression dynamics that linoleic acid production is upregulated in the primordia of several species. This is consistent with previous reports of higher linoleic and unsaturated fatty acid proportions in fruiting bodies of Agaricomycetes (see above). There may be at least three explanations for this observation, including (i) regulation of membrane fluidity in response to temperature changes, (ii) the same being genetically encoded in fruiting body development, or (iii) fulfilling the requirements of oxylipin production. We argue that hypothesis (i) is unlikely, however, we can’t speculate on hypotheses (ii) and (iii).

Regulation of membrane fluidity by increasing the proportion of unsaturated fatty acids is a common mechanism across organisms(H and S 2005). Therefore, a question arises whether the higher inferred proportion of linoleic acid and corresponding changes in gene expression are genetically coded traits of fruiting bodies or simply a consequence of lower temperature required by several species for fruiting. At least for the transcriptomes we produced(Sipos et al. 2017a; Almási et al. 2019; Krizsán et al. 2019) we know that vegetative mycelium samples were harvested at the same temperature as fruiting body samples for all species, which rules out the possibility that higher linoleic acid proportion is simply a response to lower fruiting temperatures. Rather, this suggests that higher linoleic acid concentration is a genetically encoded trait in fruiting bodies. Similar conclusions were reached in the context of unsaturated fatty acids in *L. edodes*(H and S 2003). From an evolutionary point of view, such a genetic adaptation could go back to the exposed nature of fruiting bodies; in the natural environments (soil, wood) the vegetative mycelium is in more stable conditions than fruiting bodies are, which might therefore need to be adapted to colder temperatures by increasing the proportion of unsaturated fatty acids in fruiting body tissues. At the same time, due to their large surfaces and continuous evaporation, fruiting bodies cool themselves (on average ∼2.5C colder than their environments(Dressaire et al. 2016)), which may explain why genetic encoding of higher linoleic acid proportions is beneficial for their development. It should be noted that linoleic acid production is also required for the synthesis of several oxylipin species(Combet et al. 2006), which might be involved in cell-to-cell communication.

#### 4.3.6. Other potentially membrane lipid related genes

We detected several CDE orthogroups which are not parts of the above-mentioned pathways, yet might be related to lipid metabolism, membrane organization or assembly. In many cases the functions of these orthogroups is not clear, but their widespread developmental expression suggests they might be interesting players in fruiting body development. Therefore, we discuss them below, with details on potential functions where available.

**Glycerophosphodiester phosphodiesterases** - A CDE orthogroup of glycerophosphodiester phosphodiesterases (represented by *C. cinerea* 20997)(Table 9) was developmentally expressed in 11 species (FC>4), which makes it one of the most widely developmentally expressed families. However, it is hard to make functional inferences as these proteins show only limited similarity to experimentally characterized fungal proteins (highest to Pgc1 of *S. cerevisiae*). *S. cerevisiae* Pgc1 (Phosphatidylglycerol phospholipase C) negatively regulates phosphatidylglycerol degradation, leading to increased concentrations in deletion mutants(Šimočková et al. 2008). The glycerophosphodiester phosphodiesterase family (IPR030395), is represented by three copies in most examined Agaricomycetes (some species have 4), of which only one is developmentally expressed in all species. These form a CDE orthogroup in which most genes are FB-init (*C. aegerita, A. ostoyae, A. ampla L. bicolor, L. tigrinus, P. ostreatus, S. commune),* whereas expression had different dynamics in *C. cinerea, L. edodes, M. kentingensis, Ph. chrysosporium*). These expression patterns suggest that the role of this orthogroup, although unknown at the moment, may be key to the initiation of fruiting body formation.

**Table 9.** Summary of developmental expression dynamics of CDE orthogroups of other lipid metabolism related genes across 12 species. Protein ID of a representative protein is given follows by the number of species in which the orthogroup is developmentally regulated at fold change 2 and 4 (FC>2 and FC>4, respectively). Putative function and ortholog in *S. cerevisiae* (if any) are also given. Abbreviations: 0-gene absent, 1-gene present but not developmentally regulated, 2 - developmentally regulated at fold change >2, 4- developmentally regulated at fold change >4. Species names are abbreviated as: Cc – *C. cinerea*, Aa – *A. ampla*, Ab – *A. bisporus*, Ao – *A. ostoyae*, Ca – *C. aegerita*, Lb – *L. bicolor*, Le – *L. edodes*, Lt – *L. tigrinus*, Mk – *M. kentingensis*, Pc – *Ph. chrysosporium*, Po – *P. ostreatus*, Sc – *S. commune*.

| ID | Fold change |  | Putative function | *S. cerevisiae* ortholog | Species |  |  |  |  |  |  |  |  |  |  |  |
| --- | --- | --- | --- | --- | --- | --- | --- | --- | --- | --- | --- | --- | --- | --- | --- | --- |
|  | >2 | >4 |  |  | Cc | Aa | Ab | Ao | Ca | Lb | Le | Lt | Mk | Pc | Po | Sc |
| 20997 | 12 | 11 | Glycerophosphodiester phosphodiesterase | – | 4 | 4 | 4 | 4 | 4 | 4 | 4 | 4 | 4 | 2 | 4 | 4 |
| 380219 | 6 | 6 | Diacylglycerol pyrophosphate phosphatases | – | 4 | 4 | 0 | 4 | 0 | 0 | 0 | 0 | 4 | 0 | 4 | 4 |
| 376602 | 9 | 9 | Caleosin | – | 4 | 4 | 1 | 0 | 4 | 4 | 0 | 4 | 4 | 4 | 4 | 4 |
| 365330 | 12 | 9 | Sphingolipid C9-methyltransferase | – | 4 | 2 | 2 | 4 | 4 | 4 | 4 | 4 | 2 | 4 | 4 | 4 |
| 395241 | 11 | 8 | Sphingolipid C9-methyltransferase | – | 4 | 2 | 4 | 4 | 2 | 4 | 4 | 4 | 4 | 2 | 4 | 1 |
| 444998 | 8 | 6 | Marvel domain protein | – | 4 | 1 | 0 | 4 | 1 | 1 | 4 | 2 | 4 | 2 | 4 | 4 |
| 462849 | 10 | 5 | Marvel domain protein | – | 4 | 1 | 0 | 2 | 2 | 2 | 4 | 2 | 4 | 4 | 2 | 4 |

We noticed that the detected glycerophosphodiester phosphodiesterases showed strongly correlated expression with members of another CDE orthogroup (represented by *C. cinerea* 380149, Fig. 7) in six species (data not shown). This contains putative orthologs of *S. cerevisiae* Cpt1, a choline/ethanolamine phosphotransferase that catalyzes the last step (formation of phosphatidylcholine) of the CPD-choline (Kennedy) pathway in *S. cerevisiae*. The strongly correlated expression in the case of six species (Pearson r >0.8) suggests that the two genes are linked and possibly form a linear pathway. However, their role in the context of fruiting body formation requires further research.

**Marvel protein family -** We also detected two CDE orthogroups (represented by *C. cinerea* 444998, 462849) (Table 9) of Marvel proteins showing low similarity to *A. nidulans* mrvA, *C. albicans* Mrv2 and *S. cerevisiae* Fhn1. The gene family seems to be generally poorly characterized. In *S. cerevisiae*, it may be involved in nonclassical protein secretion, might act as a sensor of sphingolipids or be related to membrane organization(M et al. 2010). In metazoans, marvel proteins are related to sterol-rich membrane region positioning(Sánchez-Pulido et al. 2002). Members of the first orthogroup (*C. cinerea* protein ID: 444998) were FB-init in several species *(A. ostoyae, M. kentingensis, P. ostreatus, S. commune, L. edodes)*, suggesting that they fulfill key functions in fruiting body initiation. The second orthogroup (*C. cinerea* protein ID: 462849) showed less characteristic expression patterns, with upregulation mostly late in development in most species.

**Sphingolipid C9-methyltransferases -** We found two CDE orthogroups of sphingolipid C9-methyltransferases (*C. cinerea* protein ID: 395241, 365330)(Table 9) that were FB-init in nearly all species. One of the orthogroups (*C. cinerea* protein ID: 395241) contains orthologs of Mts1 of *C. albicans,* which is involved in glucosylceramide biosynthesis. Glucosylceramides are glycosphingolipids that form structural components of the fungal cell membrane, with diverse roles (cell division, spore germination, pathogenicity, lipid rafts(Poeta et al. 2014)), although little mechanistic detail to date.

**PAP2 enzyme family -** A CDE orthogroup of diacylglycerol pyrophosphate phosphatases (*C. cinerea* protein ID: 380219)(Table 9) was FB-init in most species in which the gene was present. The encoded proteins show high similarity (but not 1-to-1 orthology) to *A. nidulans* and *C. albicans* membrane phospholipid metabolism proteins AN1671 and Dpp3, respectively. The latter is required also for farnesol biosynthesis(Nickerson et al. 2006). These genes belong to the broader *PAP2* family, which produce diacylglycerol and are key in the synthesis of phospholipids, triacylglycerol and the generation of signaling molecules with a lipid nature. The upregulation of these genes in fruiting bodies might be related to increased membrane material production associated with cell proliferation during early development.

**Caleosins -** Caleosins are well-characterized in plants and indirect data indicate they may be involved in oxylipin signaling, lipid metabolism, reproduction(Rahman et al. 2018), though exact functional knowledge seems to be missing. In fungi, caleosins are involved in sporulation, pathogenicity and lipid storage(Y et al. 2015). A previous study found that the caleosin family is present in ∼30% of fungal genomes(Rahman et al. 2018), whereas we found that it is present in most of the examined Agaricomycetes, mostly as a single-copy gene. Members of a detected caleosin CDE orthogroup (*C. cinerea* protein ID: 376602)(Table 9) were FB-init in most species and consistently had peak expression late in development, which is compatible with presumed roles in fruiting body initiation and sporulation, respectively.

### 4.4. Storage carbohydrate metabolism

Fungal colonies accumulate and mobilize various carbohydrates as reserve material and for energy storage. Previous research has revealed that there are very specific and tightly regulated accumulation and mobilization patterns of glycogen, trehalose and mannitol within fungal fruiting bodies(U et al. 2000). Storage carbohydrate metabolism and distributions in the mycelium and fruiting bodies have been studied in a number of economically important mushroom-forming species, including *F. velutipes*(Kitamoto et al. 2001)*, Pleurotus spp.*(Han et al. 2003; Zhou et al. 2016a; Zhu et al. 2019)*, Pisolithus microcarpus*(AN and MD 2010)*, A. bisporus*(WELLS et al. 1987; TM et al. 2008; Patyshakuliyeva et al. 2013)*, L. bicolor*(Deveau et al. 2008) and described to a high precision/resolution in *C. cinerea* in a series of papers by Moore et al(Kuhad et al. 1987; Ji and Moore 1993; Moore 2013), including links with cAMP levels (summarized by (U et al. 2000)). Briefly, these studies provided evidence on tightly regulated glycogen, trehalose and mannitol levels in mycelia and various tissues and developmental stages of fruiting bodies. They also revealed that the identity and quantities of major sugars in fruiting bodies differ from species to species; trehalose dominates in *L. bicolor*(Deveau et al. 2008) and *C. cinerea*(Kues 2000), mannitol in *A. bisporus*(Morton et al. 1985) and *L. edodes*(Tan and Moore), and both present in similar quantities in *P. ostreatus*(Hong Jai-Sik;Kim 1988; Zhou et al. 2016a).

More recently, genome sequences allowed the reconstruction of carbohydrate metabolism pathways based on gene orthology information to better studied model systems such as *S. cerevisiae*. Deveau et al found that most of the genes in known fungal pathways related to glycogen, trehalose and mannitol metabolism were found in the genome of the ectomycorrhizal *L. bicolor*(Deveau et al. 2008) and similar patterns can be expected in these very conserved pathways upon examination of other Agaricomycetes genomes. Transcriptomic studies further allowed previous biochemical observations to be interpreted in the light of gene expression patterns, although this has not yet been systematically performed in any species. In the following sections we review information on the metabolism of glycogen, trehalose and mannitol gleaned from transcriptomic studies of mushroom development. Many of these studies highlighted that shared patterns of gene expression related to storage carbohydrate metabolism exist among mushroom-forming species. For example, Merenyi et al showed that dynamic glycogen accumulation/breakdown is shared between *C. neoformans* basidium development and complex fruiting body morphogenesis, two extremes of Basidiomycota development along the complexity gradient(Merényi et al. 2021). On the other hand, trehalose concentrations were found to react to heat shock in fruiting bodies of *Flammulina filiformis*(Liu et al. 2016). Our discussion hereafter is based on well-described pathways of storage carbohydrate metabolism in yeast(François and Parrou 2001; JM et al. 2012).

**Glycogen metabolism** - Glycogen production and mobilization takes place in various tissues and developmental stages and follows tight regulation that may differ from species to species (reviewed in (Ji and Moore 1993; Kues 2000; Xie et al. 2020), see also (Ceccaroli et al. 2011) for Ascomycota). In the Agaricomycetes, this is best known in *C. cinerea* and *A. bisporus,* whereas mechanistically glycogen metabolism is best known in *S. cerevisiae*(François and Parrou 2001), which can be used to reconstruct corresponding pathways in mushroom-forming fungi. Glycogen synthesis encompasses nucleation by glycogenins and elongation and branching by glycogen synthases. Glycogenins (orthologs of *S. cerevisiae* Glg1) are upregulated in young fruiting body gills and/or mature fruiting body gills of most compared species (Table 10), respectively. This likely corresponds to intense glycogen synthesis during basidiospore maturation(Kües 2000). Consistent with this, orthologs of yeast Gac1, a phosphatase that positively regulates the activity of glycogen synthases, is upregulated in gills and mature fruiting bodies of multiple species. Gac1 in *S. cerevisiae* regulates glycogen synthases together with Glc7. The latter has orthologs in mushroom-forming fungi, but these are not developmentally regulated. In *S. cerevisiae*, a number of other CBM21 domain proteins (Pig1, Pig2) also participate in the regulation of glycogen synthase activity, however, these have no clear orthologs in mushroom-forming fungi. On the other hand, the glycogen branching enzyme Glc3 orthologs (represented by *C. cinerea* 365699) show more diverse expression peaks, and orthologs are upregulated mostly in stipe tissues (in *C. cinerea, L. bicolor, P. ostreatus* and *A. ostoyae*) (Fig. 9, Supplementary Fig. 6), indicating that linear glycogen chain production is transcriptionally decoupled from the addition of branched side chains to the glycogen backbone.

**Fig. 9.**
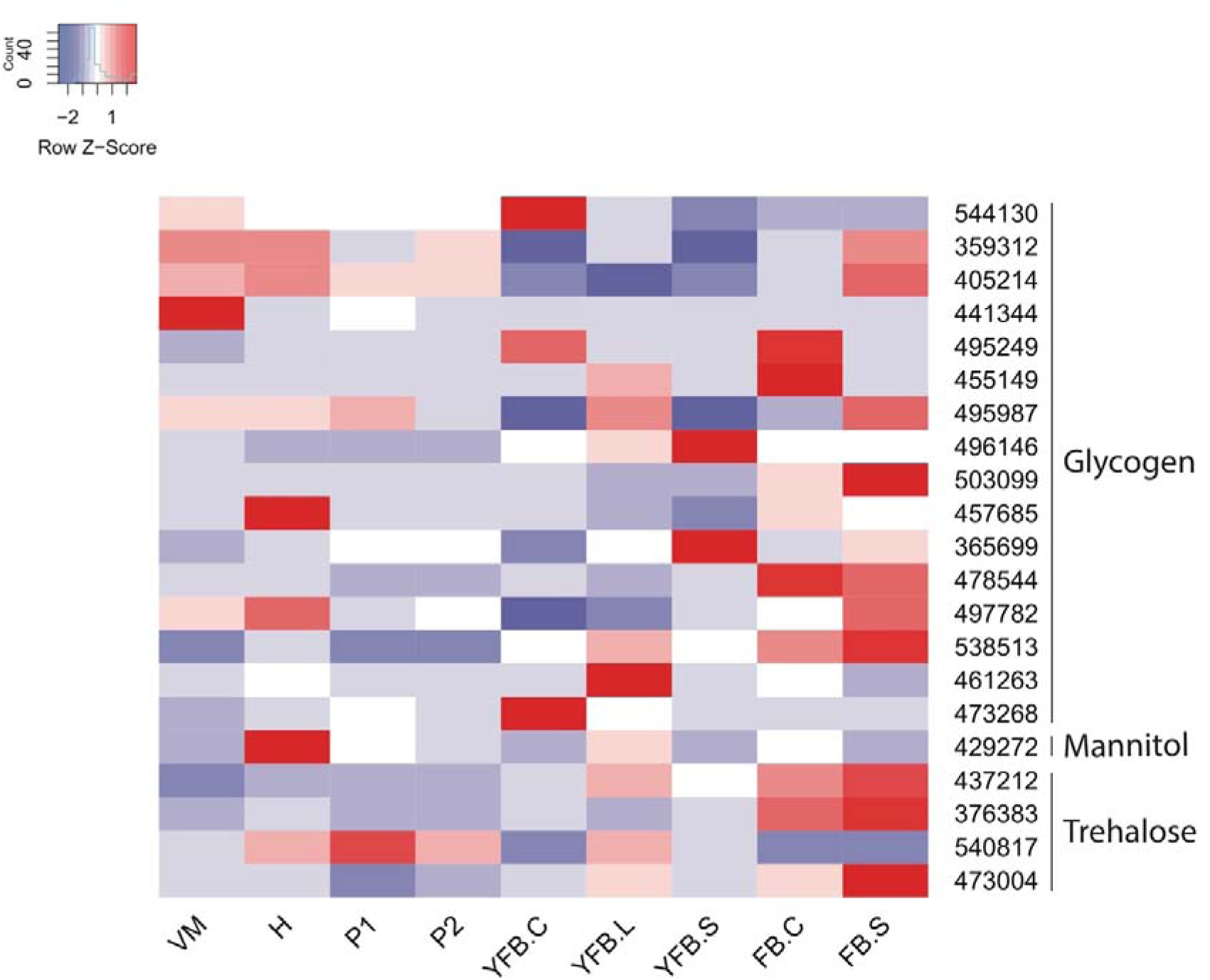
Expression heatmap of storage carbohydrate metabolism genes in *C. cinerea*. Genes are denoted by Protein IDs. Blue and red colors represent low and high expression, respectively. Developmental stages are abbreviated as follows: VM – vegetative mycelium, H – hyphal knot, P1 – stage 1 primordium, P2 – stage 2 primordium, YFB.C – young fruiting body cap, YFB.L – young fruiting body gills, YFB.S – young fruiting body stipe, FB.C – mature fruiting body cap (including gills), FB.S – mature fruiting body stipe.

**Table 10.** Summary of developmental expression dynamics of CDE orthogroups of storage carbohydrate metabolism genes across 12 species. Protein ID of a representative protein is given follows by the number of species in which the orthogroup is developmentally regulated at fold change 2 and 4 (FC>2 and FC>4, respectively). Putative function and ortholog in *S. cerevisiae* (if any) are also given. Abbreviations: 0-gene absent, 1-gene present but not developmentally regulated, 2 - developmentally regulated at fold change >2, 4- developmentally regulated at fold change >4. Species names are abbreviated as: Cc – *C. cinerea*, Aa – *A. ampla*, Ab – *A. bisporus*, Ao – *A. ostoyae*, Ca – *C. aegerita*, Lb – *L. bicolor*, Le – *L. edodes*, Lt – *L. tigrinus*, Mk – *M. kentingensis*, Pc – *Ph. chrysosporium*, Po – *P. ostreatus*, Sc – *S. commune*.

| ID | Fold change |  | Putative function | *S. cerevisiae* ortholog | Species |  |  |  |  |  |  |  |  |  |  |  |
| --- | --- | --- | --- | --- | --- | --- | --- | --- | --- | --- | --- | --- | --- | --- | --- | --- |
|  | >2 | >4 |  |  | Cc | Aa | Ab | Ao | Ca | Lb | Le | Lt | Mk | Pc | Po | Sc |
| 360850 | 9 | 8 | Alcohol dehydrogenase | – | 4 | 1 | 4 | 4 | 4 | 4 | 4 | 2 | 4 | 1 | 4 | 1 |
| 369180 | 10 | 7 | Sorbitol dehydrogenase | – | 2 | 4 | 4 | 4 | 0 | 4 | 2 | 4 | 4 | 1 | 2 | 4 |
| 455149 | 11 | 6 | CBM21 domain containing protein | – | 4 | 2 | 2 | 4 | 4 | 2 | 2 | 2 | 4 | 1 | 4 | 4 |
| 441344 | 7 | 2 | Glucoamylase | – | 2 | 4 | 0 | 2 | 1 | 0 | 0 | 2 | 4 | 2 | 2 | 1 |
| 365699 | 8 | 5 | 1,4-alpha-glucan-branching enzyme | YEL011W | 4 | 1 | 2 | 4 | 2 | 4 | 4 | 1 | 0 | 1 | 4 | 2 |
| 496146 | 2 | 0 | Serine/threonine-protein phosphatase controlling glycogen metabolism | YER133W | 2 | 1 | 1 | 0 | 1 | 2 | 1 | 1 | 1 | 1 | 1 | 1 |
| 457685 | 9 | 5 | Glycogen synthase | YFR015C | 4 | 2 | 2 | 4 | 1 | 2 | 4 | 1 | 4 | 1 | 4 | 2 |
| 473268 | 6 | 3 | Glucoamylase | YIL099W | 4 | 0 | 2 | 4 | 0 | 1 | 2 | 0 | 4 | 0 | 2 | 0 |
| 461263 | 7 | 3 | Glycogenin | YJL137C | 4 | 1 | 2 | 4 | 2 | 1 | 2 | 1 | 1 | 1 | 4 | 2 |
| 359312 | 9 | 5 | UTP-glucose-1-phosphate uridylyltransferase | YKL035W | 4 | 2 | 2 | 4 | 2 | 4 | 4 | 1 | 2 | 1 | 4 | 1 |
| 497782 | 7 | 5 | Phosphoglucumutase 2 | YMR105C | 4 | 1 | 4 | 4 | 0 | 2 | 4 | 2 | 0 | 1 | 4 | 1 |
| 538513 | 2 | 1 | Cyclin-dependent protein kinase | YPL031C | 2 | 1 | 1 | 1 | 1 | 1 | 1 | 1 | 1 | 1 | 4 | 1 |
| 495987 | 9 | 7 | Glycogen phosphorylase | YPR160W | 4 | 1 | 4 | 4 | 2 | 4 | 4 | 4 | 2 | 1 | 4 | 1 |
| 405214 | 10 | 6 | Glycogen debranching enzyme | YPR184W | 2 | 2 | 2 | 4 | 4 | 1 | 4 | 4 | 4 | 1 | 4 | 2 |
| 495249 | 10 | 7 | Glycoside hydrolase family 13<br>alpha-amylase | – | 4 | 4 | 4 | 4 | 4 | 4 | 2 | 2 | 0 | 2 | 4 | 1 |
| 544130 | 11 | 7 | CBM21 domain containing protein | – | 4 | 2 | 4 | 4 | 4 | 2 | 4 | 4 | 2 | 1 | 4 | 2 |
| 429272 | 4 | 4 | Mannitol dehydrogenase | – | 4 | 0 | 4 | 0 | 0 | 4 | 0 | 0 | 0 | 0 | 4 | 0 |
| 478544 | 8 | 2 | Alpha,alpha-trehalose-phosphate<br>synthase | YBR126C | 2 | 1 | 2 | 2 | 1 | 2 | 2 | 2 | 4 | 1 | 4 | 1 |
| 437212 | 10 | 2 | Neutral alpha-trehalase | YDR001C | 2 | 2 | 2 | 2 | 4 | 2 | 1 | 2 | 2 | 1 | 2 | 4 |
| 473004 | 8 | 4 | Trehalose-phosphatase | YDR074W | 4 | 1 | 2 | 2 | 4 | 2 | 1 | 4 | 2 | 1 | 4 | 1 |
| 503099 | 7 | 3 | Trehalose synthase complex<br>regulatory subunit TSL1 | YML100W | 4 | 1 | 2 | 2 | 1 | 4 | 1 | 2 | 2 | 1 | 4 | 1 |
| 376383 | 9 | 6 | Trehalase | – | 4 | 2 | 2 | 0 | 4 | 4 | 1 | 4 | 4 | 1 | 4 | 2 |
| 540817 | 11 | 6 | Trehalose phosphorylase | – | 4 | 4 | 2 | 4 | 4 | 1 | 2 | 2 | 2 | 2 | 4 | 4 |

Genes that encode proteins upstream of glycogen synthesis, for example orthologs of the uridinephosphoglucose pyrophosphorylase Ugp1 (*C. cinerea* protein ID: 359312) and the phosphoglucomutase PGM1 (*C. cinerea* protein ID: 497782) of *S. cerevisiae*, are developmentally regulated in multiple species (Table 10). They show some overlap in expression patterns, in particular, a tendency for upregulation late in development, but their pattern of expression is quite unique and show lower expression fold changes.

It should be noted that the glycogen synthase kinase 3 (GSK-3)(Chang et al. 2019a) gene in *C. cinerea* (*C. cinerea* protein ID: 362899), despite its name, it functionally not relevant to glycogen metabolism, it is an orthologue of *S. cerevisiae* Rim11, a pleiotropic kinase involved in cell cycle regulation.

**Glycogen mobilization** happens through three primary genes in *S. cerevisiae*(François and Parrou 2001) - all these have orthologs in mushroom-forming fungi (Table 10). Orthologs of the yeast glycogen phosphorylase Gph1 (represented by *C. cinerea* 495987) mostly show high expression in vegetative mycelia or early development, whereas those of the glycogen debranching enzyme Gdb1 are upregulated also in mature fruiting bodies in all species (Fig. 9). This probably indicates differential usage of phosphorolysis and debranching activities in different stages of development. Sga1 is a sporulation specific glucoamylase in *S. cerevisiae*. In our species orthologs (represented by *C. cinerea* 473268) of this gene do not show expression peaks in gills, the tissues in which spore development takes place. Rather, in *C. cinerea*, this gene is upregulated in young fruiting body cap tissues. How this upregulation corresponds to patterns of glycogen metabolism remains to be understood. Large-scale postmeiotic utilization of glycogen accumulated in subhymenial tissues was reported in *C. cinerea*(Kües 2000), however, subhymenial tissues were part of gill samples in previous RNA-Seq experiments, which excludes the possibility that Sga1 orthologs were involved in that process. In other species, expression patterns were different, ranging from upregulation in mature fruiting bodies in *A. ampla*, to peak expression in vegetative mycelium (*P. ostreatus* and *M. kentingensis*). We also detected a CDE orthogroup of alpha-amylases (represented by *C. cinerea* 495249), which find *A. nidulans* AmyA as best hit and are probably involved in glycogen catabolism. In *C. cinerea*, this gene showed upregulation in primordia and peak expression in young fruiting body caps, similar to other genes related to glycogen mobilization.

In addition to known members of the glycogen catabolism pathways, we identified two CBM21 domain containing CDE orthogroups that were developmentally regulated in 11/7 and 11/6 species (*C. cinerea* protein ID: 544130 and 455149) (Table 10). In most species, this gene had the highest expression in vegetative mycelia, but showed significant expression dynamics also during fruiting body morphogenesis. This seems to be a Basidiomycota-specific orthogroup, with no significantly similar proteins in model Ascomycota. The CBM21 domain mostly occurs in starch-binding proteins (glucoamylase and α-amylase(Ashikari et al. 1986)) whereas in this orthogroup, the CBM21 domain occurs alone, providing no specific clues to function. Therefore, our placement of this CDE orthogroup under this chapter should be viewed tentatively until functional studies prove or disprove the role of the family in glycogen metabolism.

Beyond these gene families, glycogen utilization patterns in fruiting bodies raise a number of exciting research avenues, such as the mode of transport within the fruiting body and the identity of transporters involved in that process.

**Trehalose metabolism** - Out of the at least five pathways known to be involved in trehalose synthesis, two have been found in Agaricomycetes(Deveau et al. 2008). The first, canonical pathway comprises the action of a protein complex formed by four proteins in *S. cerevisiae* (Tps1 - *C. cinerea* 478544, Tps2 - 473004, Tsl1 and Tps3 - no ortholog(François and Parrou 2001)). Of these, Tsl1 (*C. cinerea* protein ID: 503099) and Tps3 are paralogs that arose from a recent whole genome duplication and thus there is only one corresponding gene in filamentous fungi. The three genes showed widespread developmentally regulated expression in examined Agaricomycetes (Table 10), with upregulation mostly late in development with notable co-regulation between the three genes, as expected for members of a protein complex (Fig. 9).

A second, noncanonical pathway involving trehalose phosphorylase (TP) exists in fungi and was detected in multiple mushroom-forming species(K et al. 1998; Eis and Nidetzky 1999; Wannet et al. 1999; Deveau et al. 2008; Thammahong et al. 2017). Trehalose phosphorylase gene formed a CDE orthogroup (*C. cinerea* protein ID: 540817) in the Agaricomycetes we examined(Merényi et al. 2021) and show upregulation in stipes and in late stages of development in most species (Table 10), although deviations from this pattern exist (e.g in *C. cinerea*). For example, the gene showed peak expression in primordium stages of *A. ampla, S. commune* and *M. kentingensis*(Almási et al. 2019; Krizsán et al. 2019; Ke et al. 2020b).

For **catabolizing trehalose**, fungi exhibit two types of trehalose hydrolyzing activities, one termed acid trehalase and the other called neutral trehalase(François and Parrou 2001). Agaricomycetes species possess clear orthologs of neutral trehalases orthologous to Nth1 of *S. cerevisiae*. On the other hand, and in contrast to Deveau et al (Deveau et al. 2008), we did not find proteins in the genomes of Agaricomycetes showing significant sequence similarity (based on BLAST) to acid trehalase. A single neutral trehalase gene was found in Agaricomycetes, this formed a CDE orthogroup at FC>2 (*C. cinerea* protein ID: 437212, Table 10) and showed varied expression patterns in the species examined.

Orthologs of previously described fungal trehalose transporters were not detectable in Agaricomycetes genomes, based on sequence similarity, in agreement with a previous study on *L. bicolor*(Deveau et al. 2008). However, it should be noted that fungal trehalose transporters are poorly known and current knowledge likely misses a large proportion of genes with this activity(Thammahong et al. 2017).

**Mannitol metabolism** - Mannitol accumulates in high concentrations in fruiting bodies of certain species, possibly as an osmolyte assisting fruiting body growth(Chakraborty et al. 2004; Zhou et al. 2016b) (see also opposing evidence above). A NADP-dependent mannitol 2-dehydrogenase was reported from *A. bisporus*(JM and H 1998; S et al. 2001). In agreement, in *L. bicolor* Deveau et al reported a single mannitol 2-dehydrogenase-dependent pathway of mannitol biosynthesis(Deveau et al. 2008), which works via the direct reduction of fructose. Using the *A. bisporus* NADP-dependent mannitol 2-dehydrogenase gene(JM and H 1998) as a query, we identified an orthogroup (represented by *C. cinerea* 429272) of short-chain dehydrogenase/reductase genes that contained only *A. bisporus, L. bicolor, C. cinerea,* and *P. ostreatus*, but was missing from other species (formed multiple orthogroups probably due to duplications in *A. bisporus*). However, members of this orthogroup were developmentally regulated in all species (Table 10); they were upregulated at the initiation of fruiting and showed expression peaks in gills of *C. cinerea* and *P. ostreatus*, young and mature caps of *L. bicolor*, as well as a marked upregulation in the stipe tissues of *A. bisporus*. These observation agree with mannitol metabolism being enhanced in fruiting bodies of *L. bicolor* (Deveau et al. 2008).

In summary, the genomes of Agaricomycetes contain clear orthologs of experimentally characterized members of glycogen, trehalose and mannitol metabolism pathways. Many of the genes in these pathways are developmentally regulated in fruiting bodies, often showing conserved expression peaks in multiple species, reflecting a conservation of storage carbohydrate mobilization strategies within fruiting bodies. Interestingly, in *L. bicolor* trehalose and glycogen metabolism was upregulated in *both* fruiting bodies and mycorrhizae, providing links between fruiting bodies and ECM(Deveau et al. 2008; Martin et al. 2008).

### 4.5. Transcriptional regulators

#### 4.5.1. Transcription factors

Transcription factors are proteins that modulate gene expression by binding to specific DNA sequences and subsequently activating or inhibiting transcription. Transcription factors play a major role in the regulation of development across the tree of life, and mushroom-forming fungi are no exception. They are involved in many stages of development, ranging from the establishment of a dikaryon to various aspects of mushroom development.

The best studied are the homeodomain transcription factors of the *MatA* mating-type locus that are essential for the establishment of a fertile dikaryon, which is required for fruiting body formation in heterothallic basidiomycetes(Raudaskoski and Kothe 2010; Vonk and Ohm 2018). During mating the homeodomain transcription factors expressed from compatible *MatA* loci dimerize and activate dikaryon-specific transcriptional pathways, including the formation of clamp connections. The mechanism of mate recognition is very similar to that found in the yeast *S. cerevisiae*(Haber 2012) and is present across the fungal kingdom, although many ascomycetes do not use homeodomain transcription factors for mating(Houbraken and Dyer 2015). In some Agaricales, like *C. cinerea* and *S. commune*, the *MatA* locus has undergone expansion, improving the chances of finding a compatible mate(Freihorst et al. 2016).

Several transcription factors have been identified that act downstream of the *MatA* homeodomain transcription factors. Deletion of the transcription factors *hom2, tea1* and *wc2* in *S. commune* resulted in strains that did not show dikaryotic development beyond clamp formation(Ohm et al. 2011, 2013; Pelkmans et al. 2017a). Generally, the resulting colonies remained symmetrical and showed no sign of mushroom development. Hom2 is highly conserved in mushroom-forming fungi and features four putative phosphorylation sites. Disruption of these phosphorylation sites leads to a constitutively active Hom2 and the inhibition of vegetative growth and monokaryotic fruiting, indicating that Hom2 is post-translationally regulated(Pelkmans et al. 2017a). WC2 is part of the blue light sensing white collar complex first identified in *N. crassa*. In *N. crassa*, WC1 and WC2 interact through PAS domains and WC1 contains a LOV domain that binds a FAD chromophore when exposed to blue light. This results in conformation changes in the white collar complex and activates transcription factor domains on both WC1 and WC2(Corrochano 2019). Orthologs of WC1 (dst1 in *C. cinerea*) and WC2 (Cc.wc2 in *C. cinerea*) in *S. commune* and *C. cinerea* fulfil a similar role and deletion (*S. commune wc1* and *wc2*, *C. cinerea wc2*) or mutation (*C. cinerea wc1*) of either *wc1* or *wc2* prevents the formation of mature fruiting bodies and is called a blind phenotype(Terashima et al. 2005; Kuratani et al. 2010; Nakazawa et al. 2011; Ohm et al. 2013). In *S. commune* this results in a phenotype where no dikaryotic development is observed and in increased sensitivity to intense light. However, in *C. cinerea* a blind phenotype leads to the so called ‘dark stipe’ phenotype(Kamada et al. 2010). In this phenotype, the primordial shaft is elongated, but pileus and stipe development cannot be completed. In *S. commune*, WC1 does not contain a DNA-binding domain, unlike WC1 in *N. crassa*, making transcriptional activity dependent on WC2(Ohm et al. 2010). Orthologs of white collar complex members with the same function have been reported in *L. edodes*(Sano et al. 2009) and *P. ostreatus*(Qi et al. 2020). Furthermore, developmental expression of wc members was reported for *F. filiformis*(Liu et al. 2020b). Mutants of the gene tea1, which is downregulated in Δ*wc1,* Δ*wc2,* Δ*hom2,* Δ*bri1* and Δ*fst4* dikaryons, also did not form mushrooms(Pelkmans et al. 2017a). It should be noted that in *tea1* deletion mutants, dikaryotic colonies sporadically produced clusters of mushrooms, indicating that Tea1 is not essential for fruiting body formation and may act as a “switch” that can be bypassed. The ARID/BRIGHT DNA binding domain transcription factor Bri1 was initially also thought to play a major role in fruiting development(Ohm et al. 2011). However, later characterization of *bri1* deletion strains revealed that inactivation leads to significant growth defects that result in a delay in fruiting(Pelkmans et al. 2017a). The fungal specific transcription factor *fst4* has a very similar phenotype where no mushrooms are formed, but deletion strains of this transcription factor do form asymmetrical colonies after mating, which is the first step in mushroom development(Kües and Liu 2000; Ohm et al. 2010). In *P. ostreatus*, mutations in *gat1* resulted in a strain that did not fruit(Nakazawa et al. 2019). The missense mutation inside the zinc-finger binding domain resulted in a dominant gain of function mutation, which suggests that the mutated transcription factor has an altered DNA-binding specificity. This is in line with disruption of the ortholog *gat1* in *S. commune*, where the phenotype was not dominant and resulted in the formation of more, but smaller mushrooms(Ohm et al. 2011). A general overview of Zn(II)2Cys6 zinc cluster transcription factors, including expression patterns in *P. ostreatus* fruiting bodies, was published by Hou et al(Hou et al. 2020). In *Ganoderma lucidum*, silencing of the c_2_h_2-_type transcription factor *pacC* also resulted in the absence of fruiting bodies, as well as altered mycelial growth(Wu et al. 2016). Interestingly, PacC is conserved across the fungal kingdom and activates genes under alkaline conditions, and inhibits genes under acidic conditions(Peñalva et al. 2008). Therefore, it is unclear if the absence of fruiting can be attributed to fruiting-specific transcription or general homeostasis. In *L. edodes*, PriB was described as a primordium-upregulated transcription factor(Endo et al. 1994). Finally, silencing of *hada-1* resulted in delayed fruiting, as well as multiple growth defects, in *Hypsizygus marmoreus*(Zhang et al. 2021b).

Downstream in development, *S. commune c2h2, fst1* and *zfc7* gene disruption resulted in arrested fruiting body formation, where primordia were formed that did not develop further into mature fruiting bodies(Ohm et al. 2011; Vonk and Ohm 2021). Interestingly, overexpression of *c2h2* could increase the rate of fruiting in *A. bisporus*, indicating that *c2h2* expression is a limiting step in fruiting body development(Pelkmans et al. 2016). The transcription factors Fst1 and Zfc7 were identified due to enrichment of histone 3 lysine 4 (H3K4) dimethylation in their genes during fruiting body development after 4 days, which indicates increased genetic accessibility and transcription during this developmental stage(Vonk and Ohm 2021). Further downstream, the *C. cinerea* high-mobility group transcription factor Exp1 regulates pileus expansion and autolysis to complete fruiting(Muraguchi et al. 2008b). Notably, this transcription factor is conserved and developmentally regulated in all studied mushroom-forming fungi, except for *A. ampla, Pt. gracilis* and *S. commune*, whose fruiting bodies do not feature a pileus.

Instead of promoting fruiting body development, several transcription factors are involved in promoting vegetative growth or repressing fruiting. These include *F. velutipes lfc1* and *hmg1*, where silencing increased the rate of fruiting(Wu et al. 2020a; Meng et al. 2021). Additional transcription factors are *P. ostreatus Pofst3* and *S. commune fst3, gat1* and *hom1*(Ohm et al. 2011; Qi et al. 2019). These genes are differentially regulated in the majority of mushroom forming species, and conserved in all of them. When the genes are deleted (*S. commune*) or silenced (*P. ostreatus)* it results in the formation of more, smaller mushrooms. Finally, premature stop codons in *pcc1* in *C. cinerea* (natural variation, truncated from 561 to 210 amino acids, HMG-box not disrupted) and *P. ostreatus* (CRISPR/Cas9-assisted gene mutagenesis, truncated from 471 to 58 amino acids, HMG-box disrupted) resulted in monokaryotic fruiting and the formation of pseudoclamps(Murata et al. 1998; Boontawon et al. 2021). In *C. cinerea* these pseudoclamps were formed during monokaryotic growth, indicating a relation between the *MatA* homeodomain transcription factors and Pcc1. In *P. ostreatus* however, dikaryotic strains formed pseudoclamps, suggesting a role in clamp formation.

Many transcription factors are conserved across fungi or specifically in mushroom-forming fungi. Both *wc2* and *pacC* are conserved in both ascomycetes and basidiomycetes and except for *hmg1 (F. velutipes), lfc1 (F. velutipes), pdd1 (F. velutipes), pcc1 (C. cinerea* and *P. ostreatus)* and *zfc7 (S. commune)* all characterized transcription factors are at least conserved in mushroom-forming fungi. *Cag1* (*C. cinerea* protein ID: 466216) and *Cc.Tup1* (*C. cinerea* protein ID: 369655) of *C. cinerea* are further examples of highly conserved transcription factors(Masuda et al. 2016). They are homologs of the highly conserved general transcriptional co-repressor Tup1 of *S. cerevisiae*, which can form repressive complexes with a wide range of sequence-specific transcription factors. Deletion of Cag1 resulted in a cap-growthless phenotype, which, upon closer inspection turned out to be caused by developmental defects of gills(Masuda et al. 2016). However, the expression patterns are generally not conserved during fruiting body development(Almási et al. 2019; Krizsán et al. 2019). For example, the developmental stage with the highest expression level of the ortholog can vary greatly between species. This can indicate either that transcriptional regulation of fruiting body development is poorly conserved between species, or that transcription factors are regulated post-translationally. The *MatA* mating type genes are an example of post-translational regulation, where the transcription factors are constitutively expressed, but only become activated when heterodimers are formed between compatible homeodomain transcription factors. Similarly, in *N. crassa* WC2 is activated by structural changes in its interaction partner WC1 when blue light is detected. Furthermore, Hom2 in *S. commune* is constitutively expressed, but post-translationally phosphorylated to become inactive.

While many expression patterns of transcription factors are not conserved in mushroom-forming fungi, there are several interesting and conserved transcription factors. We identified 51 orthogroups of TFs that were developmentally regulated in a significant number of species (Table 11). Among the functionally characterized transcription factors, orthologs of dst1 (=WC1, *C. cinerea* protein ID: 369621) and Cc.wc2 (*C. cinerea* protein ID: 8610) of *C. cinerea* formed CDE orthogroups developmentally regulated in 10/4 and 10/6 (FC>2/4) species, respectively. Orthologs of dst1 and Cc.wc2 showed varied expression profiles in the examined species. An orthogroup of homeodomain transcription factors (*C. cinerea* protein ID: 497620) is developmentally regulated during fruiting body formation in all mushroom forming fungi, except for *Pt. gracilis*, which is known for its simplified fruiting body structure. The detected CDE orthogroups also included exp1 of *C. cinerea* (represented by *C. cinerea* 421729), which were developmentally regulated in 9/5 species (FC>2/4). In accordance with its reported function in *C. cinerea*(Muraguchi et al. 2008b), it showed peak expression in young fruiting body caps. However, the *A. ostoyae, L. bicolor* and *M. kentingensis* orthologs were upregulated in stipe tissues. In *P. ostreatus* we found the highest expression in cap cuticle. Furthermore, a C_2_H_2_-type transcription factor (*C. cinerea* protein ID: 494418) is conserved and expressed in all mushrooms except for *A. ampla*, *Pt. gracilis* and *S. commune*. It may therefore play a similar role to Exp1 in *C. cinerea*(Muraguchi et al. 2008b), which is also conserved and developmentally regulated in most fungi except for *A. ampla, Pt. gracilis* and *S. commune*. While many other transcription factor orthogroups are conserved in all studied mushroom-forming fungi, their expression patterns are not strongly conserved. Therefore, proteomics or functional analyses are required to further elucidate the role of transcription factors in fruiting body formation.

**Table 11.**
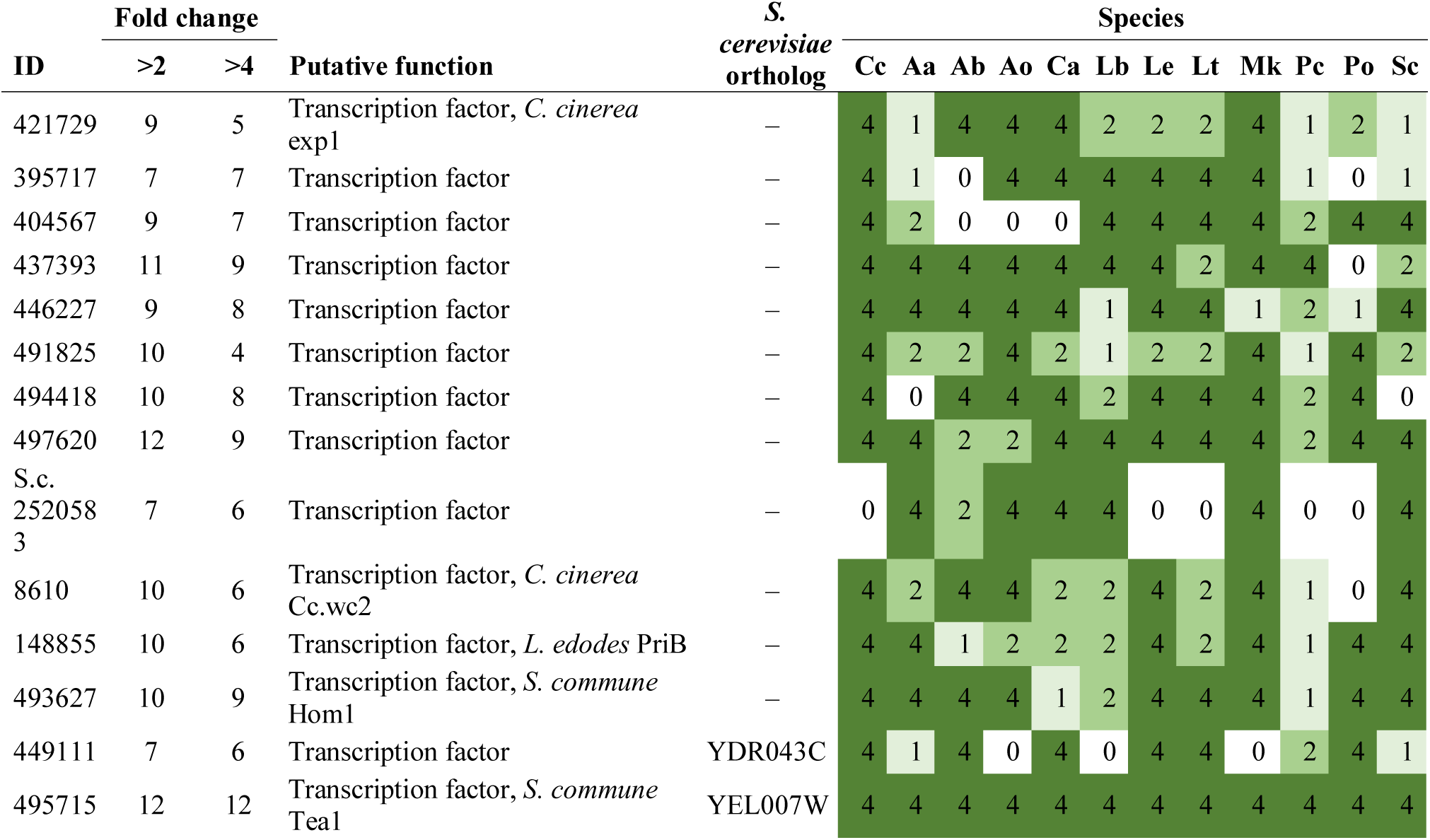
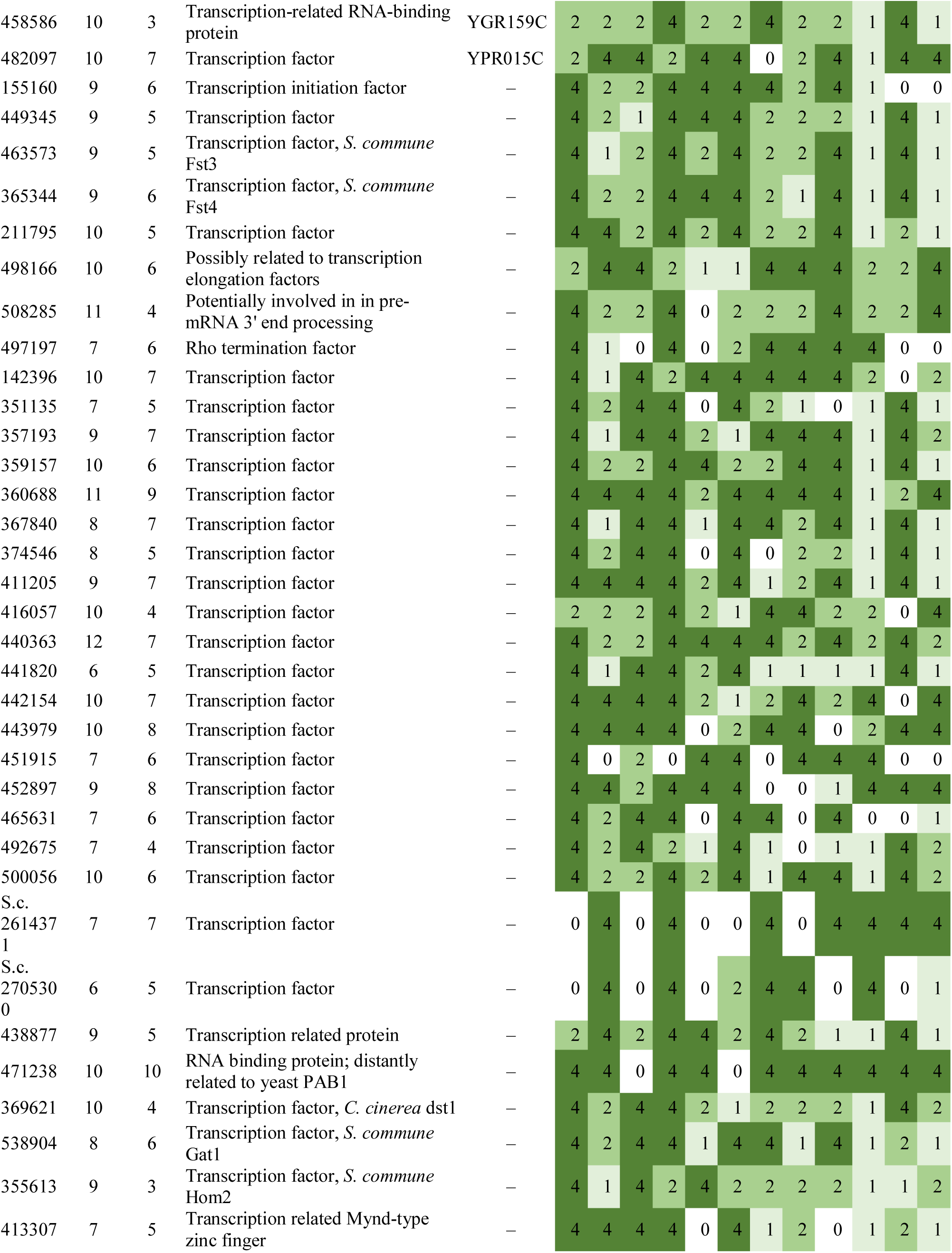
Summary of developmental expression dynamics of CDE orthogroups of transcriptional regulator genes across 12 species. Protein ID of a representative protein is given follows by the number of species in which the orthogroup is developmentally regulated at fold change 2 and 4 (FC>2 and FC>4, respectively). Putative function and ortholog in *S. cerevisiae* (if any) are also given. Abbreviations: 0-gene absent, 1-gene present but not developmentally regulated, 2 - developmentally regulated at fold change >2, 4- developmentally regulated at fold change >4. Species names are abbreviated as: Cc – *C. cinerea*, Aa – *A. ampla*, Ab – *A. bisporus*, Ao – *A. ostoyae*, Ca – *C. aegerita*, Lb – *L. bicolor*, Le – *L. edodes*, Lt – *L. tigrinus*, Mk – *M. kentingensis*, Pc – *Ph. chrysosporium*, Po – *P. ostreatus*, Sc – *S. commune*.

#### 4.5.2. RNA binding proteins

**Pumilio family** – This family comprises evolutionarily conserved sequence-specific RNA-binding proteins that bind 3’ UTR regions of specific mRNAs. This family has diverse roles in fungi, including the regulation of meiotic and sexual processes (e.g. Mcp2 of *S. pombe*)(Wilinski et al. 2017; Liu et al. 2018). Pumilio proteins were found to be active during fruiting body formation in both the Asco- and Basidiomycota(Krizsán et al. 2019; Merényi et al. 2020b). With a few exceptions, all Agaricomycetes contain six Pumilio protein encoding genes. These formed two CDE orthogroups (Table 12), one of which (*C. cinerea* protein ID: 378537) contains orthologs of the *Sch. pombe* meiotic coiled-coil protein Mcp2 and shows expression dynamics reminiscent of mitotic and meiotic genes (see Chapter 4.1.1), and another, comprising orthologs of *Sch. pombe* Puf3 and *C. neoformans* Pum1 (*C. cinerea* protein ID: 354109). Both orthogroups are developmentally regulated in a large number of species (8/7 and 7/6 at FC2/4, respectively). In *C. neoformans*, pum1 has been associated with basidium development and morphogenesis, suggesting that its orthologs may have similar roles in Agaricomycetes too.

**Table 12.**
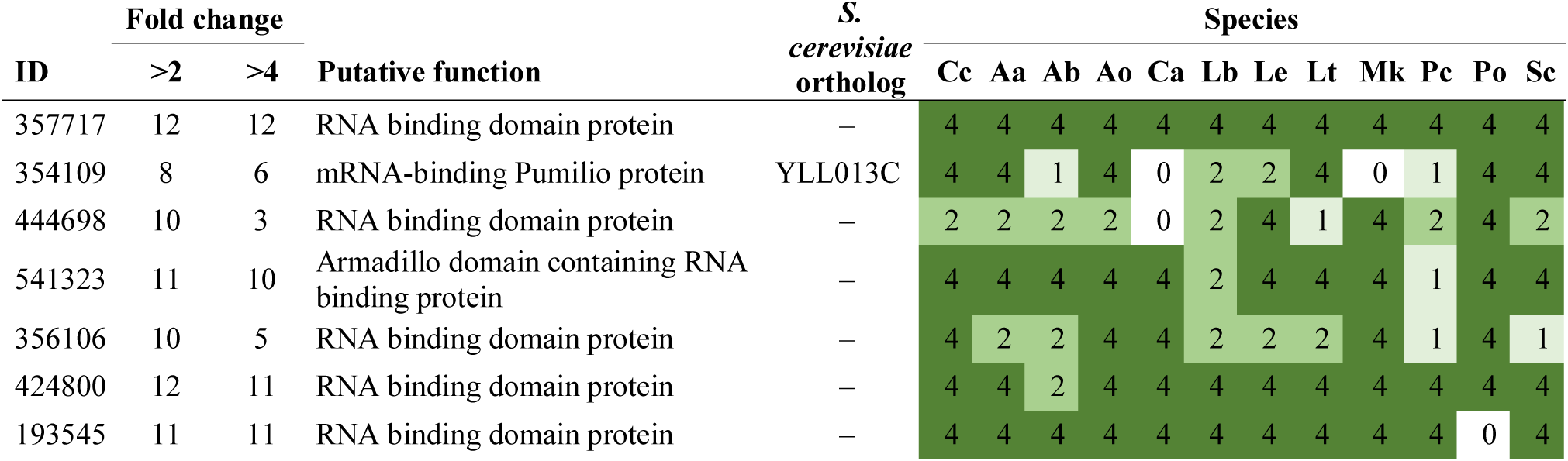
Summary of developmental expression dynamics of CDE orthogroups of RNA-binding protein encoding genes across 12 species. Protein ID of a representative protein is given follows by the number of species in which the orthogroup is developmentally regulated at fold change 2 and 4 (FC>2 and FC>4, respectively). Putative function and ortholog in *S. cerevisiae* (if any) are also given. Abbreviations: 0-gene absent, 1-gene present but not developmentally regulated, 2 - developmentally regulated at fold change >2, 4- developmentally regulated at fold change >4. Species names are abbreviated as: Cc – *C. cinerea*, Aa – *A. ampla*, Ab – *A. bisporus*, Ao – *A. ostoyae*, Ca – *C. aegerita*, Lb – *L. bicolor*, Le – *L. edodes*, Lt – *L. tigrinus*, Mk – *M. kentingensis*, Pc – *Ph. chrysosporium*, Po – *P. ostreatus*, Sc – *S. commune*.

| ID | Fold change |  | Putative function | S.<br>*cerevisiae*<br>ortholog | Species |  |  |  |  |  |  |  |  |  |  |  |
| --- | --- | --- | --- | --- | --- | --- | --- | --- | --- | --- | --- | --- | --- | --- | --- | --- |
|  | >2 | >4 |  |  | Cc | Aa | Ab | Ao | Ca | Lb | Le | Lt | Mk | Pc | Po | Sc |
| 357717 | 12 | 12 | RNA binding domain protein | – | 4 | 4 | 4 | 4 | 4 | 4 | 4 | 4 | 4 | 4 | 4 | 4 |
| 354109 | 8 | 6 | mRNA-binding Pumilio protein | YLL013C | 4 | 4 | 1 | 4 | 0 | 2 | 2 | 4 | 0 | 1 | 4 | 4 |
| 444698 | 10 | 3 | RNA binding domain protein | – | 2 | 2 | 2 | 2 | 0 | 2 | 4 | 1 | 4 | 2 | 4 | 2 |
| 541323 | 11 | 10 | Armadillo domain containing RNA binding protein | – | 4 | 4 | 4 | 4 | 4 | 2 | 4 | 4 | 4 | 1 | 4 | 4 |
| 356106 | 10 | 5 | RNA binding domain protein | – | 4 | 2 | 2 | 4 | 4 | 2 | 2 | 2 | 4 | 1 | 4 | 1 |
| 424800 | 12 | 11 | RNA binding domain protein | – | 4 | 4 | 2 | 4 | 4 | 4 | 4 | 4 | 4 | 4 | 4 | 4 |
| 193545 | 11 | 11 | RNA binding domain protein | – | 4 | 4 | 4 | 4 | 4 | 4 | 4 | 4 | 4 | 4 | 0 | 4 |

**RRM family** - We identified five very conserved CDE orthogroups of RNA recognition motif (RRM) family proteins. One of these (*C. cinerea* protein ID: 471238) were annotated as related to mitosis and meiosis and is discussed under that chapter. The RRM family comprises proteins that bind diverse RNA types and shows considerable diversity in fungi, with 68-88 genes in the Agaricomycetes. Overall, relatively few members of the family were developmentally expressed, however, the orthogroups we detected showed a conserved expression pattern. One of them (*C. cinerea* protein ID: 357717) was developmentally regulated in all 12 species (Table 12) and was upregulated in the second half of fruiting body development (young fruiting body and mature fruiting body in *C. cinerea*), in particular in gills (*A. ostoyae, P. ostreatus, M. kentingensis, C. cinerea*) where gill tissue was sampled separately, or caps (*L. bicolor, L. edodes*). It was also upregulated considerably in primordia relative to vegetative mycelia. A less characteristic, but similar and very conserved late upregulation was observed also in the second CDE orthogroup (*C. cinerea* protein ID: 356106), which contains reciprocal best hits of *Sch. pombe* Vip1.

The third CDE orthogroup (*C. cinerea* protein ID: 193545, 11/11 species at FC=2/4) contains *C. neoformans* Csa2. Members of this orthogroup showed strong induction in the first primordium stage in *C. cinerea, A. ostoyae, C. aegerita, L. bicolor, L. edodes, P. ostreatus, S. commune, A. ampla, M. kentingensis*. In *C. neoformans* it shows a strong peak 24h after the induction of sexual development, which, combined with evidence for regulation by pum1, led Liu et al to conclude it is involved in basidium development and sporulation(Liu et al. 2018). It’s an orthologue of *Sch. pombe* Mei2, which is related to meiosis. The other three basidium-related genes reported by Liu et al (Fad1, Fas1, CNA00260)(Liu et al. 2018) do not have clear orthologs in *Coprinopsis*.

The CDE orthogroup represented by *C. cinerea* 424800 (Table 12) was consistently upregulated in gills and sporulating stages (in *A. ampla, S. commune*) in all species, suggesting roles in hymenium and/or spore formation. In model Ascomycota, proteins in this orthogroup find Pab1 of both *Sch. pombe* and *S. cerevisiae* as their best Blast hits, which are related to mRNA export and the control of poly(A) tail length.

Finally, the CDE orthogroup comprising *C. cinerea* 368527 includes best reciprocal hosts of *S. cerevisiae* Nam8, a meiosis-specific splicing factor. Our expression data are consistent with a meiosis-specific role: the gene is developmentally regulated in 6/4 species (FC2/4, see above, Table 1) and shows a peak expression in meiotic tissues in all species (gills, cap+gills or mature fruiting bodies). Based on these data, we hypothesize that this gene also participates in meiosis-specific splicing events in the basidia during fruiting body formation.

#### 4.5.3. RNA interference, Small RNA

RNA interference is a conserved eukaryotic gene-silencing mechanism that acts via base pairing of small noncoding RNAs of ∼20-30 nucleotides to DNA/RNA targets. RNAi is key to developmental and differentiation processes, genome defense against viruses or transposons. There is very limited knowledge on the role of RNAi in mushroom-forming fungi (but see(Lau et al. 2018)), whereas the processes are extensively studied in the Ascomycota (in particular *N. crassa* and *Sch. pombe*(Dang et al. 2011)), where RNAi has well-documented roles in defense against viruses, transposons, meiotic silencing of unpaired DNA, translational repression and even inter-organismal pathogenic interactions(Weiberg et al. 2013).

A fully functional RNAi machinery exists in the Basidiomycota. Based on orthology to *Neurospora* proteins, Lau et al annotated dicer-like proteins as Dcl-1 (*C. cinerea* protein ID: 444647), Dcl-2 (*C. cinerea* protein ID: 366047) and Dcl-3 (*C. cinerea* protein ID: 438955)(Lau et al. 2018), whereas Stajich et al annotated the argonaute family proteins AGO (*C. cinerea* protein ID: 355234), an AGO-like protein (*C. cinerea* protein ID: 473372) and Qde-2 (*C. cinerea* protein ID: 363313)(Stajich et al. 2010) (Table 13). We identified 3-16 genes encoding argonaute and argonaute-like proteins in the examined Agaricomycetes. Furthermore, fungal RNA interference involves RNA-dependent RNA polymerases, which are required for generating double-stranded RNA or for amplifying sRNA signals(Dang et al. 2011). The genomes of the examined Agaricomycetes contain 5-13 RNA-dependent RNA polymerases; of these, the orthogroup represented by *C. cinerea* 498671 is most similar to the RDP1 protein of *S. pombe*, which was shown to be integral to RNA interference(Carmichael et al. 2004). Other RNA-dependent RNA polymerases (Table 11) might participate in alternative RNAi pathways or signal amplification(S et al. 2012b). Interestingly, we found no clear orthologs of SAD-1, a meiotic silencing protein(Borkovich et al. 2004), in the examined genomes.

**Table 13.** Summary of developmental expression dynamics of CDE orthogroups of RNA interference related genes across 12 species. Protein ID of a representative protein is given follows by the number of species in which the orthogroup is developmentally regulated at fold change 2 and 4 (FC>2 and FC>4, respectively). Putative function and ortholog in *S. cerevisiae* (if any) are also given. Abbreviations: 0-gene absent, 1-gene present but not developmentally regulated, 2 - developmentally regulated at fold change >2, 4- developmentally regulated at fold change >4. Species names are abbreviated as: Cc – *C. cinerea*, Aa – *A. ampla*, Ab – *A. bisporus*, Ao – *A. ostoyae*, Ca – *C. aegerita*, Lb – *L. bicolor*, Le – *L. edodes*, Lt – *L. tigrinus*, Mk – *M. kentingensis*, Pc – *Ph. chrysosporium*, Po – *P. ostreatus*, Sc – *S. commune*.

| ID | Fold change |  | Putative function | S.<br>*cerevisiae*<br>ortholog | Species |  |  |  |  |  |  |  |  |  |  |  |
| --- | --- | --- | --- | --- | --- | --- | --- | --- | --- | --- | --- | --- | --- | --- | --- | --- |
|  | >2 | >4 |  |  | Cc | Aa | Ab | Ao | Ca | Lb | Le | Lt | Mk | Pc | Po | Sc |
| 155983 | 6 | 2 | RNA-dependent RNA polymerase | – | 4 | 1 | 2 | 2 | 1 | 2 | 4 | 2 | 1 | 1 | 1 | 1 |
| 498671 | 4 | 1 | RNA-dependent RNA polymerase | – | 4 | 1 | 1 | 1 | 2 | 1 | 2 | 1 | 2 | 1 | 0 | 1 |
| 438955 | 4 | 2 | Dicer-like protein | – | 2 | 1 | 1 | 2 | 4 | 4 | 1 | 1 | 1 | 1 | 1 | 1 |
| 464843 | 2 | 1 | RNA-dependent RNA polymerase | YDL128W | 4 | 0 | 0 | 0 | 0 | 2 | 0 | 0 | 0 | 0 | 0 | 0 |
| 366047 | 6 | 1 | Dicer-like protein | YLR332W | 1 | 1 | 1 | 2 | 2 | 4 | 2 | 1 | 2 | 1 | 2 | 1 |
| 473372 | 8 | 0 | AGO-like protein | – | 2 | 2 | 1 | 2 | 1 | 2 | 2 | 2 | 2 | 1 | 2 | 1 |
| 495272 | 6 | 4 | Argonaute-like protein | – | 4 | 1 | 0 | 2 | 2 | 1 | 4 | 1 | 4 | 1 | 4 | 1 |
| 502116 | 9 | 6 | Argonaute-like proteins with high similarity to *A. nidulans* rsdA and *S. pombe* ago1 | – | 4 | 2 | 1 | 4 | 2 | 4 | 4 | 4 | 4 | 1 | 2 | 1 |
| 363313 | 7 | 1 | Orthologs of QDE-2 | – | 2 | 2 | 1 | 2 | 1 | 1 | 2 | 2 | 4 | 1 | 2 | 1 |
| 440860 | 5 | 1 | RNA-dependent RNA polymerase | – | 4 | 1 | 2 | 2 | 1 | 1 | 1 | 1 | 2 | 1 | 2 | 1 |
| 496192 | 5 | 1 | RNA-dependent RNA polymerase | – | 2 | 0 | 0 | 2 | 2 | 4 | 2 | 1 | 0 | 1 | 1 | 0 |
| 498410 | 5 | 1 | RNA-dependent RNA polymerase | – | 4 | 1 | 2 | 2 | 1 | 0 | 2 | 1 | 1 | 1 | 2 | 1 |
| 355234 | 1 | 1 | AGO ortholog | – | 4 | 0 | 0 | 0 | 0 | 0 | 0 | 0 | 0 | 0 | 0 | 0 |
| 368472 | 1 | 0 | Argonaute-like protein | – | 2 | 0 | 0 | 0 | 0 | 0 | 0 | 0 | 0 | 0 | 0 | 0 |
| 465906 | 1 | 0 | Argonaute-like protein | – | 2 | 0 | 0 | 0 | 0 | 0 | 0 | 0 | 0 | 0 | 0 | 0 |
| 444647 | 3 | 0 | Dicer-like protein | – | 2 | 1 | 1 | 1 | 1 | 1 | 1 | 2 | 1 | 2 | 1 | 1 |
| 124704 | 1 | 1 | RNA-dependent RNA polymerase | – | 4 | 0 | 0 | 0 | 0 | 0 | 0 | 0 | 0 | 0 | 0 | 0 |
| 444494 | 1 | 1 | RNA-dependent RNA polymerase | – | 4 | 0 | 0 | 0 | 0 | 0 | 0 | 0 | 0 | 0 | 0 | 0 |

RNAi components show some expression dynamics during fruiting body development. Genes encoding Dcl-2, Dcl-3 and AGO-like proteins were upregulated in *C. cinerea* primordia relative to vegetative mycelium(Lau et al. 2018). Similarly, AGO-like and Qde-2 were found developmentally regulated in *C. cinerea* by Krizsan et al(Krizsán et al. 2019), although Dcl-1-3 genes were not. All of the seven RNA-dependent RNA polymerases were developmentally regulated in the data of Krizsan et al. In our meta-analysis, the above-listed components did not show significant developmental expression dynamics (Table 13). The only noteworthy orthogroup (represented by *C. cinerea* 502116), was developmentally regulated in 9/6 species (FC>2/4) and contains Argonaute-like proteins that share highest similarity with *A nidulans* RsdA and *Sch. pombe* Ago1, the first of which is involved in RNA-silencing(TM et al. 2008) whereas the second has a large suite of described functions, from RNA silencing to chromosome segregation and cell cycle arrest(Carmichael et al. 2004). We found no conserved expression pattern in this orthogroup across the examined species.

The most direct evidence for the role of microRNA-like RNAs (milRNA) in mushroom development was reported by Lau et al(Lau et al. 2018), who compared small RNA transcriptome profiles of vegetative mycelium and primordium stages and detected 22 milRNA in *C. cinerea*, several of which were differentially expressed between vegetative mycelia and primordia. These results point to the potential roles of milRNA in mushroom development, although more studies are needed to understand its roles.

RNA interference has been exploited as a tool of genetic manipulation in mushroom-forming fungi, often to compensate for the lack of reliable reverse genetics tools in these organisms. Gene knockdown has been demonstrated in *S. commune*(de Jong et al. 2006)*, C. cinerea*(Namekawa et al. 2005; MA et al. 2006; Heneghan et al. 2007)*, L. bicolor*(M et al. 2009a)*, Ph. chrysosporium*(Matityahu et al. 2008)*, P. ostreatus, A. bisporus*(Costa Ana S.M.B.;Thomas 2009)*, Grifola frondosa*(Sato et al. 2015) and *L. edodes*(Nakade et al. 2011), among others, demonstrating the utility of knockdown techniques in cases/species when gene knockout was not feasible (particularly in the pre-CRISPR times).

#### 4.5.4. Chromatin regulators

Chromatin remodeling is a fundamental and general mode of epigenetic gene expression regulation. Dynamic chromatin states are generated primarily via chromatin remodeling complexes and histone modifications in fungi, in response to environmental or internal signals. Importantly, chromatin remodeling often provides the basis for broad changes in gene expression, by modulating the accessibility of DNA for transcription, as opposed to sequence-specific transcription factors, which regulate only specific genes. Epigenetic changes often associate with developmental processes; these receive increasing attention in multiple lineages, including fungi. Accordingly, studies addressing the role of chromatin remodeling in mushroom development, although quite few at the moment, revealed fundamental roles of chromatin states in fruiting-related gene transcription. For example, deleting *Cc.snf5* (*C. cinerea* protein ID: 365798, see Table 14), a member of the SWI/SNF chromatin remodeling complex, leads to the failure to initiate fruiting body development in *C. cinerea* (in addition to defects in dikaryon formation)(Ando et al. 2013). In *C. cinerea* two further genes identified in mutant analyses, *ich1*(Muraguchi and Kamada 1998) and *Cc. rmt1*(Nakazawa et al. 2010), may be related to chromatin regulation. Recent ChIP-Seq analyses of epigenetic states revealed substantial changes in transcriptional activity (with H3K4me2 as a proxy) during the transition to fruiting(Vonk and Ohm 2021), whereas histone acetyltransferase complex members were found in quantitative trait loci in *L. edodes*(Zhang et al. 2021d). The presence of SET domain methyltransferases in the Basidiomycota is also worth mentioning(Freitag 2017). Although evidence at the moment is scanty, the few available studies and extrapolation from other organisms suggests that chromatin remodeling may be a significant factor in fruiting body formation. Therefore, in the following chapters we annotate and discuss expression patterns of genes related to chromatin maintenance and remodeling based on homology to known chromatin remodeling components in *S. cerevisiae* and *N. crassa*.

**Table 14.** Summary of developmental expression dynamics of CDE orthogroups of chromatin- related genes across 12 species. Protein ID of a representative protein is given follows by the number of species in which the orthogroup is developmentally regulated at fold change 2 and 4 (FC>2 and FC>4, respectively). Putative function and ortholog in *S. cerevisiae* (if any) are also given. Abbreviations: 0-gene absent, 1-gene present but not developmentally regulated, 2 - developmentally regulated at fold change >2, 4- developmentally regulated at fold change >4. Species names are abbreviated as: Cc – *C. cinerea*, Aa – *A. ampla*, Ab – *A. bisporus*, Ao – *A. ostoyae*, Ca – *C. aegerita*, Lb – *L. bicolor*, Le – *L. edodes*, Lt – *L. tigrinus*, Mk – *M. kentingensis*, Pc – *Ph. chrysosporium*, Po – *P. ostreatus*, Sc – *S. commune*.

| ID | Fold change |  | Putative function | S.<br>*cerevisiae*<br>ortholog | Species |  |  |  |  |  |  |  |  |  |  |  |
| --- | --- | --- | --- | --- | --- | --- | --- | --- | --- | --- | --- | --- | --- | --- | --- | --- |
|  | >2 | >4 |  |  | Cc | Aa | Ab | Ao | Ca | Lb | Le | Lt | Mk | Pc | Po | Sc |
| 58294 | 10 | 0 | Actin-related protein 4 | – | 2 | 2 | 2 | 2 | 2 | 1 | 2 | 2 | 2 | 2 | 2 | 1 |
| 485296 | 9 | 0 | Protein arginine N-methyltransferase 1 | YBR034C | 2 | 1 | 2 | 2 | 2 | 2 | 2 | 2 | 2 | 1 | 2 | 1 |
| 498885 | 11 | 8 | Histone H2A.Z-specific chaperone CHZ1 | YER030W | 4 | 0 | 2 | 4 | 2 | 2 | 4 | 4 | 4 | 4 | 4 | 4 |
| 493226 | 10 | 7 | Histone isomerase | YML074C | 4 | 2 | 4 | 4 | 4 | 2 | 2 | 4 | 4 | 1 | 4 | 1 |
| 455597 | 9 | 9 | NAD-dependent histone deacetylase HST3 | YOR025W | 4 | 4 | 4 | 4 | 4 | 1 | 0 | 4 | 4 | 1 | 4 | 4 |
| 545797 | 11 | 9 | *C. cinerea* Ich1 gene | – | 4 | 4 | 4 | 4 | 1 | 2 | 4 | 4 | 4 | 4 | 2 | 4 |
| 392307 | 11 | 5 | HIRA-interacting protein 3 | – | 2 | 2 | 4 | 4 | 2 | 2 | 4 | 2 | 4 | 1 | 4 | 2 |
| 384835 | 8 | 6 | Poly(ADP-ribose) polymerase | – | 4 | 2 | 0 | 4 | 0 | 0 | 2 | 4 | 4 | 1 | 4 | 4 |
| 459336 | 9 | 5 | Poly(ADP-ribose) polymerase | – | 2 | 2 | 2 | 4 | 1 | 4 | 4 | 0 | 4 | 1 | 4 | 2 |
| 473747 | 8 | 6 | High mobility group protein | – | 4 | 1 | 4 | 4 | 2 | 2 | 0 | 4 | 4 | 1 | 4 | 0 |
| 66389 | 10 | 4 | Suppressor protein STM1 | YLR150W | 4 | 2 | 4 | 2 | 1 | 4 | 2 | 2 | 4 | 2 | 2 | 1 |
| 264120 | 2 | 0 | SET domain protein | YJL168C | 1 | 1 | 1 | 1 | 1 | 1 | 2 | 1 | 1 | 1 | 2 | 1 |
| 170659 | 1 | 0 | SET domain protein | – | 2 | 0 | 0 | 0 | 0 | 0 | 0 | 0 | 0 | 0 | 0 | 0 |
| 456684 | 3 | 0 | SET domain protein | – | 2 | 1 | 1 | 1 | 0 | 2 | 0 | 2 | 0 | 0 | 1 | 1 |
| 497003 | 1 | 0 | SET domain protein | – | 2 | 1 | 0 | 1 | 1 | 0 | 1 | 1 | 1 | 1 | 0 | 1 |
| 365798 | 2 | 0 | *C. cinerea* Snf5 family chromatin remodeling protein | – | 2 | 1 | 1 | 1 | 2 | 1 | 1 | 1 | 1 | 1 | 1 | 1 |
| 252667 | 8 | 2 | Histone acetyltransferase type B subunit 2 | YEL056W | 2 | 2 | 4 | 2 | 1 | 1 | 2 | 2 | 4 | 1 | 2 | 1 |
| 518613 | 10 | 4 | Histone acetyltransferase complex member | YLL022C | 2 | 2 | 2 | 2 | 0 | 2 | 4 | 4 | 4 | 1 | 4 | 2 |
| 420113 | 7 | 4 | Histone acetyltransferase type B catalytic subunit | YPL001W | 4 | 1 | 2 | 1 | 1 | 2 | 4 | 1 | 4 | 1 | 4 | 2 |
| 457726 | 8 | 4 | Histone acetyltransferase | – | 4 | 1 | 4 | 4 | 2 | 1 | 2 | 0 | 2 | 0 | 4 | 2 |
| 430366 | 6 | 0 | Histone deacetylase | YGL194C | 2 | 1 | 1 | 2 | 2 | 1 | 1 | 2 | 2 | 1 | 2 | 1 |
| 415916 | 4 | 0 | Histone deacetylase | YNL021W | 2 | 1 | 1 | 2 | 2 | 1 | 2 | 1 | 1 | 1 | 1 | 1 |
| 354601 | 6 | 2 | Histone deacetylase | YNL330C | 4 | 2 | 2 | 2 | 1 | 1 | 2 | 1 | 1 | 1 | 4 | 1 |
| 52602 | 5 | 3 | Histone deacetylase | YPL116W | 4 | 1 | 1 | 4 | 1 | 1 | 4 | 2 | 0 | 2 | 1 | 1 |
| 364572 | 5 | 0 | Histone deacetylase | – | 1 | 1 | 2 | 2 | 1 | 1 | 2 | 1 | 2 | 1 | 2 | 1 |
| 465141 | 8 | 2 | Histone deacetylase | – | 2 | 4 | 4 | 2 | 2 | 1 | 2 | 1 | 2 | 1 | 1 | 2 |
| 363625 | 8 | 6 | Agaricales-specific histone group | – | 4 | 0 | 0 | 0 | 4 | 4 | 2 | 4 | 4 | 2 | 4 | 0 |
| 154410 | 11 | 6 | Centromere-specific histone variant | – | 2 | 4 | 2 | 2 | 2 | 4 | 2 | 4 | 4 | 1 | 4 | 4 |
| 182362 | 11 | 5 | Histone | – | 4 | 4 | 2 | 2 | 2 | 2 | 2 | 2 | 4 | 0 | 4 | 4 |
| 473916 | 8 | 6 | Histone | – | 2 | 0 | 4 | 4 | 1 | 4 | 0 | 2 | 4 | 0 | 4 | 4 |
| 393767 | 9 | 5 | Histone H2B | – | 1 | 1 | 4 | 4 | 2 | 2 | 4 | 4 | 0 | 2 | 2 | 4 |
| 361866 | 5 | 4 | Histone H2B.2 | YBL002W | 2 | 0 | 0 | 4 | 0 | 0 | 4 | 4 | 4 | 0 | 0 | 0 |
| 418275 | 10 | 7 | Histone H2A.2 | YBL003C | 2 | 4 | 4 | 4 | 2 | 4 | 4 | 0 | 0 | 2 | 4 | 4 |
| 401831 | 9 | 6 | Histone H4 | YBR009C | 1 | 4 | 4 | 2 | 1 | 4 | 2 | 4 | 4 | 1 | 2 | 4 |
| 546355 | 4 | 1 | Histone H3 | YBR010W | 2 | 0 | 1 | 2 | 1 | 1 | 1 | 2 | 4 | 1 | 1 | 1 |
| 284515 | 11 | 2 | Histone H2A.Z | YOL012C | 2 | 2 | 4 | 2 | 2 | 2 | 2 | 2 | 4 | 2 | 2 | 1 |
| 467558 | 10 | 9 | Histone H1 | YPL127C | 4 | 4 | 4 | 4 | 4 | 4 | 4 | 4 | 1 | 2 | 0 | 4 |
| 320458 | 0 | 0 | Histone | – | 1 | 0 | 0 | 0 | 0 | 0 | 0 | 0 | 0 | 0 | 0 | 0 |
| 324852 | 0 | 0 | Histone | – | 1 | 0 | 0 | 0 | 0 | 0 | 0 | 0 | 0 | 0 | 0 | 0 |
| 380437 | 1 | 1 | Histone | – | 4 | 0 | 0 | 0 | 0 | 0 | 0 | 0 | 0 | 0 | 0 | 0 |
| 385598 | 1 | 1 | Histone | – | 4 | 0 | 0 | 0 | 0 | 0 | 0 | 0 | 0 | 0 | 0 | 0 |
| 446837 | 1 | 0 | Histone | – | 2 | 0 | 0 | 0 | 0 | 0 | 0 | 0 | 0 | 0 | 0 | 0 |
| 492876 | 1 | 0 | Histone | – | 2 | 0 | 0 | 0 | 0 | 0 | 0 | 0 | 0 | 0 | 0 | 0 |
| 503008 | 1 | 0 | Histone | – | 2 | 0 | 0 | 0 | 0 | 0 | 0 | 0 | 0 | 0 | 0 | 0 |
| 452576 | 4 | 0 | AMP-activated protein serine/threonine kinase alpha subunit Ssp2 | YDR477W | 2 | 1 | 1 | 2 | 1 | 1 | 2 | 1 | 2 | 1 | 1 | 1 |
| 360033 | 4 | 1 | 5'-AMP-activated protein kinase subunit gamma | YGL115W | 1 | 1 | 4 | 1 | 1 | 1 | 2 | 2 | 1 | 1 | 2 | 1 |
| 377394 | 8 | 1 | Histone-lysine N-methyltransferase, H3 lysine-79 specific | YDR440W | 2 | 1 | 1 | 4 | 2 | 1 | 2 | 2 | 2 | 1 | 2 | 2 |
| 511209 | 9 | 8 | SRA-YDG domain protein | – | 2 | 4 | 4 | 4 | 4 | 4 | 4 | 0 | 0 | 0 | 4 | 4 |

##### 4.5.4.1. Histone gene expression correlates with fruiting body initiation and sporulation

**Histones** - Histones are highly conserved proteins that assemble into octameric nucleosomes and condense DNA into chromatin in a tightly regulated manner. Shuffling of histone variants can generate nucleosome variants, which, combined with diverse posttranslational modifications of histones (acetylation, methylation, phosphorylation) can generate a finely diversified histone code contributing to fine-tuned regulation of chromatin states, which, in turn, impact transcriptional patterns of genes(Borkovich et al. 2004). Histone modifications are often used to make readouts of transcriptional states associated with development(Vonk and Ohm 2021). Agaricomycetes have 14-40 histone and histone-variant encoding genes. They show a very similar overall expression pattern in all species that resembles the expression of meiotic/mitotic and ribosomal genes: upregulation in primordium stages as well as meiotic tissues (gills or caps+gills) coincident with sporulation (Fig. 10, Supplementary Fig. 7). Though overall very conserved, there are slight variations to this general pattern: *A. ampla* shows a peak only in primordia, whereas in *C. cinerea* and *L. tigrinus* a meiosis/sporulation associated peak is better visible, while the primordium-peak is modest. We detected four CDE orthogroups of histones and histone variants (Table 14). These include a basidiomycetes-specific histone H2A component (*C. cinerea* protein ID: 363625), which is developmentally expressed in 8/6 species (FC2/4) and show considerable upregulation in primordia relative to vegetative mycelium in most species. Other CDE orthogroups also showed primordium- and gill-upregulation. A straightforward interpretation of the similar expression patterns of histones and cell proliferation related genes is a mutual link to DNA repair. H2A histone phosphorylation is associated with DNA repair(JA et al. 2000; Borkovich et al. 2004), providing a potential mechanistic explanation for the observed expression patterns. Somewhat contradicting this is the fact that relatives of the Mec1 kinase (*C. cinerea* protein ID: 19106), which phosphorylates H2A.Z in *S. cerevisiae*(JA et al. 2000), are not developmentally regulated in Agaricomycetes. On the other hand, histone upregulation in primordia and meiotic tissues may also be interpreted as evidence for intense transcriptional reprogramming, which has been hypothesized for the mycelium-fruiting body transition(Krizsán et al. 2019; Vonk and Ohm 2021). Alternatively, intense DNA replication may require increased supply of histones in proliferating tissues.

**Fig. 10.**
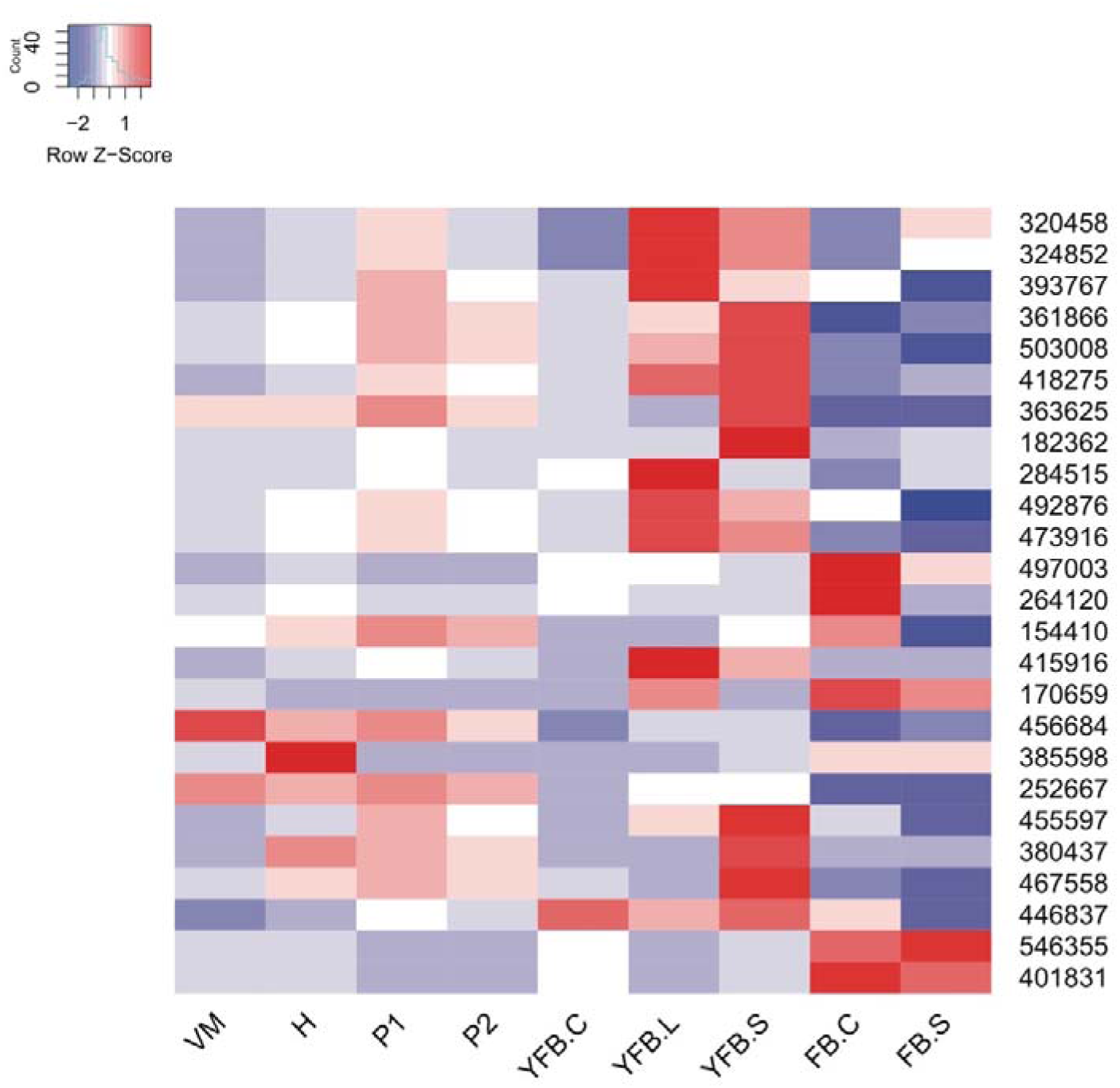
Expression heatmap of chromatin remodeling related genes (histones, histone chaperones, histone deacethylases) in *C. cinerea*. Genes are denoted by Protein IDs. Blue and red colors represent low and high expression, respectively. Developmental stages are abbreviated as follows: VM – vegetative mycelium, H – hyphal knot, P1 – stage 1 primordium, P2 – stage 2 primordium, YFB.C – young fruiting body cap, YFB.L – young fruiting body gills, YFB.S – young fruiting body stipe, FB.C – mature fruiting body cap (including gills), FB.S – mature fruiting body stipe.

**Histone chaperone** - We detected a histone chaperone CDE orthogroup containing homologs of *S. cerevisiae* Chz1 (represented by *C. cinerea* 498885)(Table 14), which forms a single-copy gene family in the Agaricomycetes. Members of this orthogroup are involved in histone replacement during transcription and showed the same expression pattern as histones (peaking in primordia and meiotic tissues), so we speculate they are involved in the same cellular processes during fruiting body initiation and sporulation. We next inventoried histone acetyltransferases, histone deacetylases, histone methyltransferases, histone ubiquitylases, histone kinases and histone ADP-ribosylases based on homology to corresponding *N. crassa* genes described by Borkovich et al(Borkovich et al. 2004) as well as diagnostic conserved domains.

**Histone acetyltransferase** (HAT) - We found one HAT gene per species in the examined Agaricomycetes (represented by *C. cinerea* 420113), in contrast to Ascomycota, that have multiple histone acetyltransferase genes(Borkovich et al. 2004). This family showed no developmental expression dynamics in our species. However, we detected several protein N-acetyltransferases, which resemble HATs, but lack orthology to known HATs and do not contain diagnostic conserved domains of HATs. They share an identical domain composition with *Sch. pombe* Ard1, a Gcn5-related N-acetyltransferase (GNAT), which acetylates diverse proteins, including transcriptional regulators. A CDE orthogroup of GNATs (*C. cinerea* protein ID: 457726) was developmentally expressed in 8/4 species (FC2/4, Table 14), however, given the lack of clear histone link we only tentatively discuss this family here, noting that functional analysis is needed to confirm their functional annotation.

**Histone deacetylases (HDACs) -** we distinguished two groups of HDACs, one related to *S. cerevisiae* Hda1 and the Sirtuin family, which were detected in 6-7 (11 in *A. ostoyae*) and 4-9 copies in the examined Agaricomycetes, respectively. Neither groups showed considerable developmental expression dynamics, except for one CDE orthogroup comprising sirtuins (represented by *C. cinerea* 455597), which was developmentally expressed in 9/9 species (FC2/4, Table 14). This family is orthologous to Hst4 of *Sch. pombe* and Hst3 of *S. cerevisiae*, which regulate DNA damage response and repair by regulating histone H3K56 acetylation(Konada et al. 2018). Consistent with their presumed role, the expression of this gene family followed closely that of mitosis/meiosis and DNA repair related genes in all species: they showed a small expression peak in primordia and a larger one in meiotic tissues coinciding with sporulation (Fig. 10).

Histone ADP-ribosylases are poly(ADP-ribose) polymerases (PARPs) and are among the least understood factors in epigenetic regulation; they attach (poly-)ADP-ribose moieties to proteins, including histones and are thought to regulate histone turnover and DNA damage response. In *N. crassa* the family is represented by a single gene, *parp*(Borkovich et al. 2004). The examined Agaricomycetes’ genomes harbor 3-9 copies of PARPs, of which two formed CDE orthogroups (*C. cinerea* protein ID: 459336, 384835), which were developmentally expressed in 8/4 and 6/5 species (FC2/4, Table 14), respectively. Their expression patterns did not follow that of histones, suggesting that their actions are independent of those.

Histone methyltransferases and histone kinases showed no noticeable expression dynamics in fruiting bodies (Table 14), suggesting that they are equally important in fruiting bodies and vegetative mycelia.

##### 4.5.4.2. Other chromatin related genes

***Ich1* gene family -** The only methyltransferase that might be connected to chromatin regulation was the gene underlying the ‘Ichijiku’ mutant, which was first identified by Muraguchi and Kamada(Muraguchi and Kamada 1998), as a mutant that produced malformed, barrel-shaped primordia, in which cap development fails. The corresponding *ich1* gene encodes a nuclear methyltransferase that contains a winged-helix DNA binding domain and was reported to be upregulated in primordia(Muraguchi et al. 2015). Our results agree and extend these observations: Ich1 orthologs form a CDE orthogroup (represented by *C. cinerea* 545797) that is developmentally expressed in 8/6 species (FC2/4)(Table 14) and shows a marked upregulation in primordia relative to vegetative mycelium in *C. cinerea, A. ostoyae* (weakly)*, A. ampla, S. commune, A. bisporus, M. kentingensis* and *P. ostreatus*. The exact mechanism of how Ich1 regulates tissue formation is not known, however, this gene represents a very interesting target for detailed studies, especially given its putative role in chromatin regulation and/or methylation. On a related note, *Cc.rmt1*, an arginine methyltransferase which was suggested to participate in histone methylation and to be required for hyphal knot and fruiting body formation(Nakazawa et al. 2010), forms an orthogroup which is developmentally regulated in nine species (at FC>2) (Table 14). The exact role of *Cc.rmt1* is not known so its role in histone methylation remains preliminary.

**Histone isomerases** - We found a CDE orthogroup that was widely developmentally expressed (represented by *C. cinerea* 493226)(Table 14). Ascomycota orthologs of these genes are histone proline isomerases in the peptidyl-prolyl cis-trans isomerase family, which contribute to nucleosome assembly and thus regulate gene expression. This orthogroup shows upregulation in gills of *C. cinerea, A. ostoyae* and *P. ostreatus* (all species in which gills were sampled separate) and in cap tissues of *L. edodes, M. kentingensis, L. tigrinus*. These expression patterns are consistent with a role in sporulation. Genes in this orthogroup also show upregulation in primordia relative to vegetative mycelia, although this is usually modest (2-3x, 10x in *P. ostreatus*).

**HMGA and AT-hook protein -** We detected a CDE orthogroup (*C. cinerea* protein ID: 473747)(Table 14) that includes primarily stipe-upregulated genes (in *C. cinerea, A. ostoyae, L. tigrinus, P. ostreatus, M. kentingensis,* but not *L. bicolor*) that shows expression peaks also in YFB cap of *C. cinerea* and FB cap of *P. ostreatus*. We found no expression dynamics in Schizophyllaceae species, which lack stipes. The orthogroup belongs to the High mobility group A family, which comprises AT-rich DNA binding proteins with an important role in chromatin organization. This family appears to be patchily distributed in the examined species, but nevertheless seems to be important for fruiting body development.

**SRA-YDG domain proteins** - We detected a CDE orthogroup (*C. cinerea* protein ID: 511209)(Table 14) of proteins containing an SRA-YDG (IPR003105) domain, which occurs in histone and chromatin binding proteins and may be associated with the cell cycle. Proteins with this domain seem to be missing in Ascomycota model species (results not shown), but show expression patterns similar to those of histones. They are upregulated in primordia and in meiotic stages (gills, caps with gills) in *C. cinerea, A. ostoyae, P. ostreatus* and *L. edodes*. In the Schizophyllaceae as well as *P. ostreatus*, the gene family is upregulated in primordia relative to vegetative mycelium samples. Based on their expression patterns, we hypothesize they are related to cell proliferation, the cell cycle or chromatin modification, but available data do not allow us to further narrow down their putative function among these categories.

### 4.6. Alternative splicing and spliceosome components

Alternative splicing is a post-transcriptional mechanism for regulating transcript identity and abundances. mRNA variants generated by splicing can increase the diversity of proteins transcribed from a single gene, but can also destine transcripts to the nonsense-mediated decay pathway and thus reduce transcript abundance posttranscriptionally(Krizsán et al. 2019). Transcriptomes allow us to describe alternative splicing complexity of organisms at an unprecedented depth. Several studies looked at alternative splicing in fungi(Zhao et al. 2013; Grützmann et al. 2014; Jin et al. 2017), including Agaricomycetes model systems(Lugones et al. 1999a; Gehrmann et al. 2016; Krizsán et al. 2019). The Basidiomycota and, in particular, the Agaricomycetes displayed a higher percentage of spliced genes than did other groups of fungi(Grützmann et al. 2014; Krizsán et al. 2019). Results of these studies highlighted that, of the several possible splicing events, intron retention (IR) is the most common in fungi while exon-skipping, which most amply contributes to generating protein isoforms, accounts for only 0.5-2.5% of fungal splicing events. Most research on splicing focuses on the latter, however, it appears that exon skipping-rich transcriptomes represent an animal innovation(Grau-Bové et al. 2018). IR events have been considered either as transcriptional noise or as a regulatory mechanism(Middleton et al. 2017). IR can introduce premature stop codons in transcripts or new functional elements for RNA binding proteins. Premature stop codons can activate nonsense-mediated decay, a surveillance mechanism that results in the degradation of the mRNA transcript. Recent results suggest that IR may not only regulate transcript levels, but can be a key driver of posttrancriptional processes(Middleton et al. 2017). There is evidence that in the Basidiomycota IR exerts its effect on transcripts independent of the nonsense-mediated decay pathway(Gonzalez-Hilarion et al. 2016).

Functionally characterized examples of developmentally relevant splicing events are known in the Ascomycota. For example, alternative splicing of the blue light receptor wc-1 gene was observed in *Phaeosphaeria* spp. and it was hypothesized that AS may control various light-dependent reactions(Chiu et al. 2010). In *Neurospora wc-1* transcription is controlled by multiple distinct cis-regulatory regions (termed different promoters), which responded differently (not, weakly or strongly) to light and resulted in alternative protein isoforms(Káldi et al. 2006). These observations, although probably just scratching the surface, implicate AS as an important contributor to the development of fruiting bodies.

Alternative transcript isoforms affect a considerable portion of genes during fruiting body development. IR-generated transcript isoforms with a dynamic expression are abundant in fruiting bodies(Gehrmann et al. 2016; Krizsán et al. 2019), affecting 20-46% of expressed genes, pointing towards potential important regulatory roles. In *S. commune*, 69% of the alternative spliced transcripts had altered predicted functional properties or subcellular localization(Gehrmann et al. 2016). Gehrmann et al reported the largest number of splicing events at the fruiting body initiation stage in *S. commune*, whereas Krizsan et al did not find substantial differences in the number of splicing events across developmental stages.

Splicing can influence transcript levels significantly during fruiting body development. A recent study found 425 genes in *S. commune* for which an alternative transcript replaced the highest expressed transcript in some stage of development. Many such events happened at fruiting body initiation (referred to as the aggregate stage), pinpointing the magnitude of transcriptome reprogramming at this developmental transition. A conceptually similar scenario, genes whose expression do not vary considerably during fruiting body development, but have splicing isoforms that vary in their expression significantly, was detected for ∼160-1300 genes in six Agaricomycetes (Krizsán et al. 2019).

Taken together, recent studies on Agaricomycetes suggest that splicing, in particular intron retention, is an important process in fruiting body development, contributing to posttranscriptional regulation of transcript levels or generates functional diversity in proteins translated from alternative transcripts. The prevalence of IR in fungal transcriptomes suggests that the main role of splicing in fungi might be posttranscriptional transcript level regulation rather than the expansion of the proteome space, although mechanistic or proteome-level studies are missing at the moment.

We here annotated spliceosome components and splicing-associated proteins in *C. cinerea* based on knowledge in *S. cerevisiae* and *Sch. pombe*(Käufer and Potashkin 2000; JM et al. 2012). We annotated 50 orthogroups associated with splicing, however, none of them showed remarkable expression dynamics in fruiting bodies (Supplementary Table 3). These included the gene described as *Le.cdc5* and *Cc.cdc5* in *L. edodes* and *C. cinerea*, respectively(Miyazaki et al. 2004), a pre-mRNA splicing factor.

### 4.7. Signal transduction systems and communication

One of the most important multicellular processes.

#### 4.7.1. Oxylipins

In fungi, oxylipins are responsible for a range of signaling functionalities and act as paracrine hormone-like substances or as so-called ‘infochemicals’(Combet et al. 2006; Holighaus and Rohlfs 2019). Ascomycota species produce diverse oxylipin compounds (reviewed in(Gessler et al. 2017)) and changes in their levels influence development. For example, oxylipins can regulate the yeast-to-hypha transition in dimorphic fungi, as well as immune evasion(Rennemeier et al. 2011) and communication with the host in pathogens, while a well-known group of oxylipins called psi (precocious sexual inducer) factors regulates sexual and asexual development in Ascomycota(Tsitsigiannis and Keller 2007). Basidiomycota oxylipins are less known. The best-known oxylipin of mushroom-forming fungi is 1-octen-3-ol, an enantiomeric compound that is mainly produced as (R)-(–)-1-octen-3-ol by widely-known species(Zawirska-Wojtasiak 2004). On the one hand, it acts as an interkingdom-signaling molecule. It may attract or repel fungivorous arthropods(Fäldt et al. 1999; Holighaus and Rohlfs 2019) or derogate insect larval development. In combination with the equally prominent octan-3-one that may attract specialist fungivores, it has also been shown to attract predators of such fungivores(Fäldt et al. 1999). Furthermore, 1-octen-3-ol has been shown to retard plant seed germination and vegetative growth(Hung et al. 2014). On the other hand, 1-octen-3-ol shows a quorum-sensing like action as it inhibits colony growth(Yin et al. 2015; Kües et al. 2018), conidiation(Fäldt et al. 1999; Yin et al. 2015), fruiting initiation(Noble et al. 2009; Eastwood et al. 2013) and germination where spores are crowded(Chitarra et al. 2004; Combet et al. 2006).

The composition of the oxylipin cocktail emitted by intact mushroom tissue changes in the course of mushroom development while measurement of the most abundant oxylipins during that process crucially depends on the sampling method: disruption of fungal tissue has been shown to trigger the release of oxylipins that are not emitted when fungal material is intact(Fäldt et al. 1999). The changing specific oxylipin bouquet emitted by a developing mushroom may not solely reflect adaptation to insect attraction for spore dispersal and repellence of fungivores. It is also possible that, e.g. octan-3-one and 1-octen-3-ol are produced by the fruiting body to inhibit on-colony spore germination(Combet et al. 2006) or colonization by mycoparasites(Berendsen et al. 2013). During fruiting body development, octan-3-one emission either positively(Combet et al. 2009) or negatively(Zhang et al. 2008; A et al. 2020; Orban et al. 2021) correlates with the amount of 1-octen-3-ol produced by (not yet) sporulating tissue samples of certain species.

The biosynthesis pathway of 1-octen-3-ol has interested mycologists for a long time(Combet et al. 2006; Kües et al. 2018; Orban et al. 2021), but remained unresolved to date. The process starts by oxygenation of polyunsaturated fatty acids (PUFAs) with linoleic acid as the predominant PUFA in Asco- and Basidiomycota(Combet et al. 2006; Brodhun and Feussner 2011). It was suggested that lipoxygenases (LOXs), linoleate diol synthases (LDSs that combine a dioxygenase – DOX – and an isomerase activity), hydroperoxide lyases (HPLs) and alcohol dehydrogenases (ADHs) participate in 1-octen-3-ol biosynthesis(Combet et al. 2006). Still, it is possible that 1-octen-3-ol can be produced by incomplete beta-oxidation or by multiple pathways(Combet et al. 2006; Brodhun and Feussner 2011). Orban et al. recently attempted to identify genes involved in oxylipin production based on correlations between 1-octen-3-ol concentrations and gene expression patterns(Orban et al. 2021) including a gene of an ene-reductase for octan-3-one formation from ADH-mediated conversion of 1-octen-3-ol.

Two main enzyme classes have been implicated in fungal oxylipin production, LOXs and LDSs(Combet et al. 2006; Orban et al. 2021). Both act on linoleic acid as the starting point, from which various compounds are produced by first an oxidation reaction, then by further modification(Combet et al. 2006). We found LOXs to be highly patchy in Agaricomycetes, whereas LDSs (basically the Ppo family of proteins) are widely conserved. Incomplete beta-oxidation has also been suggested as a possible source of oxylipins(Venter et al. 1997; Brodhun and Feussner 2011), this hypothesis is compatible with the lack of certain enzymes in many species.

**Lipoxygenases (LOXs) -** Out of the examined species, we only found lipoxygenases in *C. aegerita*, *P. ostreatus*, *L. bicolor* and *L. edodes*, which have 1-5 copies. These did not form conserved orthogroups and their expression patterns mostly did not vary substantially during development. Exceptions include the gene previously described as *PoLOX1*(Plagemann et al. 2013; Y et al. 2013) (PleosPC15_2_1068582). *PoLOX1* displays a remarkable upregulation (1880 fold, FPKM 94.000) in fruiting bodies relative to vegetative mycelium. The gene showed consistently higher expression in stipes compared to caps, which confirms the previous report on the gene(Y et al. 2013). A lipoxygenase gene (*CaeLOX4*, gene ID AAE3_04864/Genbank accession no. CAA7262624) of *C. aegerita* also showed a strong upregulation (∼150-fold) during late development (post sporulation stage) in this species, as noted by Orban et al (Orban et al. 2021). Notably, two other LOXs show strong phase-specific expression increase in fruiting stage tissue: on the one hand, *CaeLOX5* (AAE3_07753) shows an expression pattern quite congruent with the maturation-associated increase of the 1-octen-3-ol levels in the headspace of the *C. aegerita* culture. On the other hand, Cae*LOX1* (AAE3_00896) shows transcriptional upregulation before fruiting body maturation sets in. Such patterns may imply stage-specific action or (partial) functional redundancy among these genes that must, prospectively, be tested by functional characterization studies.

**Linoleate diol synthases (LDSs) -** The other well-known gene family involved in the production of oxylipins is the one of linoleate diol synthases, which makes up to PpoA-C family in *A. nidulans*. The main products of the Ppo family are the so-called psi factors, which regulate diverse developmental processes from yeast-to-hypha transition and cell aggregation (reviewed in ref(Tsitsigiannis and Keller 2007)). We identified 1-4 LDS genes in Agaricomycetes with most species having two copies. Although several genes have reciprocal best hits among Ascomycota (e.g. *C. cinerea* 423716 to PpoC of *A. nidulans*), a gene tree analysis showed that these are not true orthologs (results not shown) but likely resulted from a duplication event that predated the split of Asco- and Basidiomycota. Therefore, it is likely that Basidiomycota linoleate diol synthases do not have the psi factor producing activity that *A. nidulans* PpoA-C genes and their homologs have(Tsitsigiannis and Keller 2007).

Linoleate diol synthases grouped into three conserved orthogroups (Table 15), of which two (*C. cinerea* protein ID: 398037, 423716) were developmentally expressed in a considerable number of species. One of these (*C. cinerea* protein ID: 423716) includes *U. maydis* Ssp1 (Ustma2_2_11710), which was found to be specifically expressed in teliospores(Huber et al. 2002). Genes within these orthogroups seem to show diverse expression patterns, without noticeable conservation of expression across species. In the absence of functional information of these genes in Basidiomycota, it is impossible to speculate on their mechanistic role in fruiting body formation. They are likely to be involved in oxylipin production(Orban et al. 2021), but further studies are necessary to elucidate their functions.

**Table 15.**
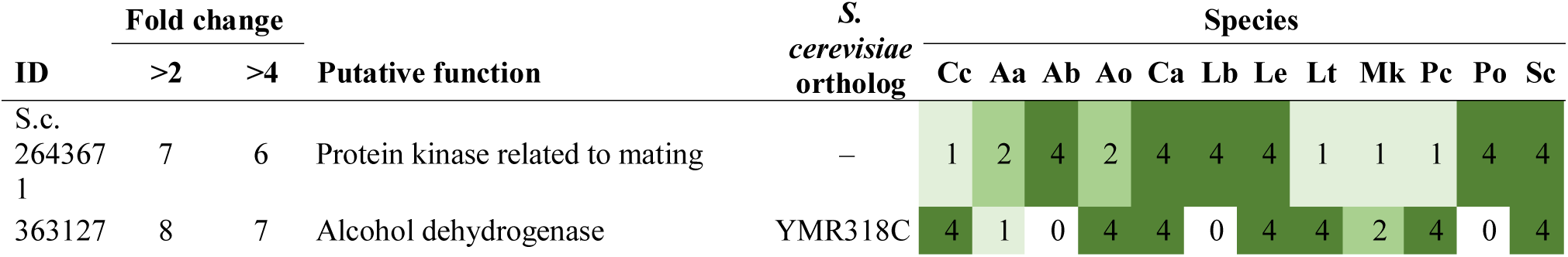
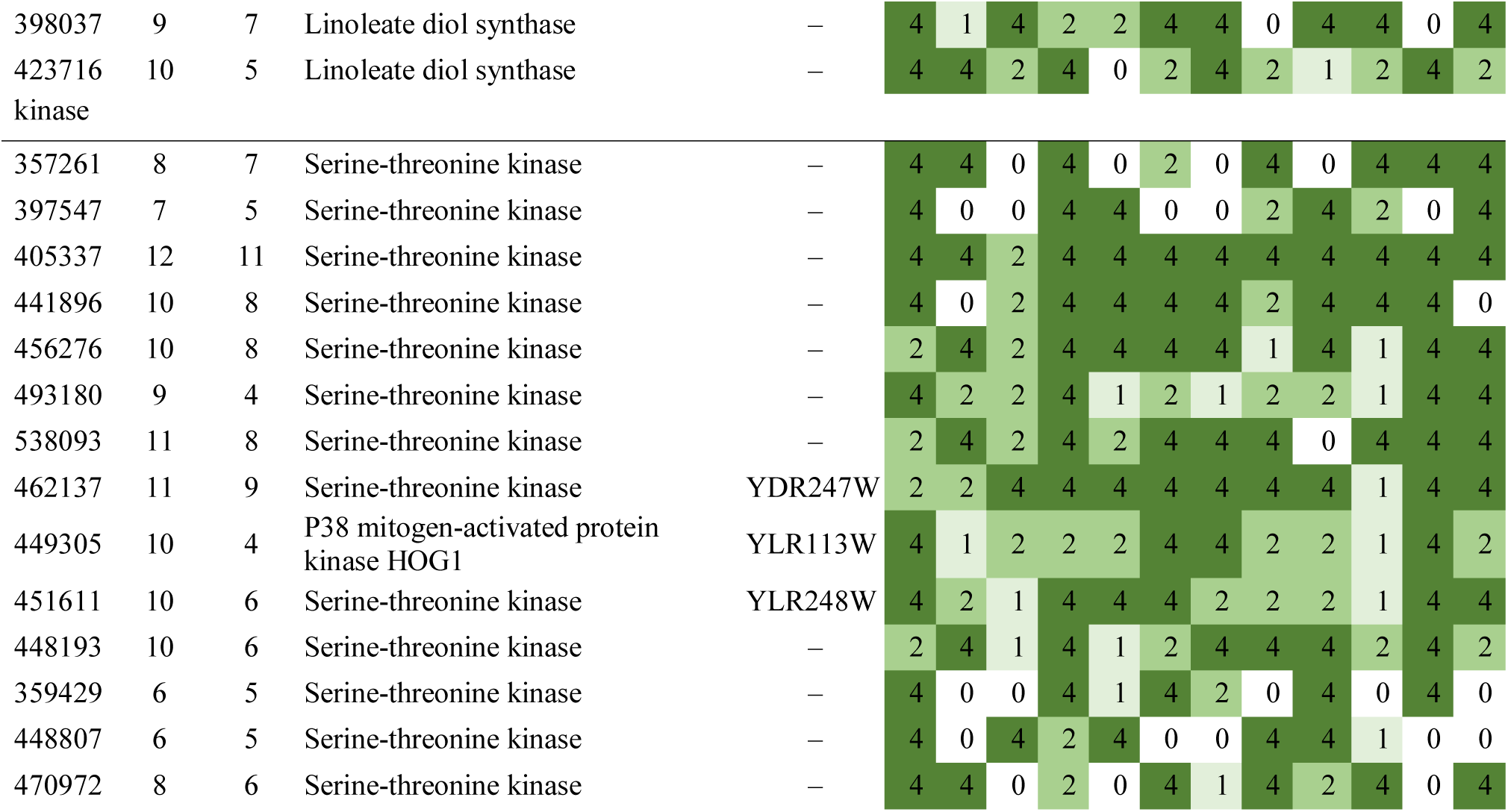
Summary of developmental expression dynamics of CDE orthogroups of signal transduction related genes across 12 species. Protein ID of a representative protein is given follows by the number of species in which the orthogroup is developmentally regulated at fold change 2 and 4 (FC>2 and FC>4, respectively). Putative function and ortholog in *S. cerevisiae* (if any) are also given. Abbreviations: 0-gene absent, 1-gene present but not developmentally regulated, 2 - developmentally regulated at fold change >2, 4- developmentally regulated at fold change >4. Species names are abbreviated as: Cc – *C. cinerea*, Aa – *A. ampla*, Ab – *A. bisporus*, Ao – *A. ostoyae*, Ca – *C. aegerita*, Lb – *L. bicolor*, Le – *L. edodes*, Lt – *L. tigrinus*, Mk – *M. kentingensis*, Pc – *Ph. chrysosporium*, Po – *P. ostreatus*, Sc – *S. commune*.

**Alcohol dehydrogenases** – the alcohol dehydrogenase family participates in a broad range of cellular processes, including oxylipin biosynthesis(Orban et al. 2021). We identified a CDE orthogroup containing *C. aegerita* Adh1-1 (represented by *C. cinerea* 363127), which showed partial conservation of expression: in most species it was upregulated during late development (*C. aegerita, A. ostoyae, L. edodes, L. tigrinus, C. cinerea*). In addition, in *S. commune, Ph. chrysosporium, A. ostoyae* and *L. edodes* it was also upregulated >4x in primordia relative to vegetative mycelium. In *M. kentingensis* and *A. ampla* the gene was not developmentally regulated, whereas *L. bicolor* and *P. ostreatus* did not have orthologs. Based on a previous report of *C. aegerita* Adh1-1(Orban et al. 2021), it is possible that this orthogroup of alcohol dehydrogenases participate in oxylipin production.

#### 4.7.2. Pheromone and pheromone receptor pathways

The structure and evolution of mating systems, including HD transcription factors (A locus), pheromone receptors and pheromone precursor genes (B locus) is among the best known aspects of Agaricomycete biology (e.g.(Kües 2000, 2015; James 2015) refs). The primary role of pheromone/receptor systems is regulating dikaryotization, i.e. the compatibility of monokaryons, nuclear movement and other downstream events. Hyphal fusion is believed to occur ubiquitously in the Basidiomycota, therefore, pheromone/receptor systems may regulate post-fusion behavior(Raudaskoski and Kothe 2010; Jones and Bennett 2011). However, pheromone/receptor systems are involved in the regulation of sexual events too. For example, mating type loci are involved in fruiting body formation in *C. cinerea*(Kues et al. 1998, 2002). Basidiomycete mating systems, pheromone and pheromone receptor genes have been reviewed recently in detail (e.g.(Raudaskoski and Kothe 2010; Jones and Bennett 2011; James 2015; Kuees 2015)). We here annotate components of the Agaricomycete mating systems, with special emphasis on copy number variation across the twelve species examined in this paper and expression dynamics during fruiting body development.

**Pheromone precursor genes** - Basidiomycetes produce mating pheromones that are structurally analogous to *S. cerevisiae* a-factor(Jones and Bennett 2011). Mating pheromones are short peptides that undergo a multi-step maturation process and regulate intercellular communication and sexual development in a wide range of fungi(Jones and Bennett 2011). Copy numbers of pheromone precursor genes vary across the Agaricomycetes and are often found in the vicinity of corresponding pheromone GPCRs(Raudaskoski and Kothe 2010). Four pheromone precursor genes were found in *L. edodes*(Wu et al. 2013)*, P. eryngii*(Kim et al. 2014)*, A. bisporus*(Foulongne-Oriol et al. 2021)*, Moniliopthora roreri*(Díaz-Valderrama and Aime 2016), five in *Ph. chrysosporium*(James et al. 2011), seven in *Pycnoporus cinnabarinus*(Levasseur et al. 2014), two in *P. touliensis*(Gao et al. 2018) and several genes (up to 20) in *C. cinerea*(Riquelme et al. 2005) and *S. commune*(Specht 1995). Synthetic pheromone administration affected PriA (a primordium-expressed gene) expression in *L. edodes*, indicating the involvement of mating pheromones in fruiting body formation(Ha et al. 2018). One of the pheromone precursor genes was expressed in glucose-limited conditions following blue light induction of fruiting in *C. cinerea*(Sakamoto et al. 2018).

In our dataset, we found up to nine pheromone-precursor encoding genes, however, we did not find any in *A. ostoyae, A. bisporus, M. kentingensis, Ph. chrysosporium* and *L. tigrinus*. The explanation for this may be that they are simply missing from the genome annotations. Pheromone precursors are short ∼50 amino acid peptides and several genome annotation protocols omit predicted proteins that are <50 or <100 amino acids long. For this same reason, it is likely that our inventories of pheromone-precursor genes in other examined species is incomplete. Nevertheless, we found pheromone precursors in *C. cinerea, A. ampla, L. bicolor, L. edodes, S. commune* and *P. ostreatus*; these genes were often developmentally regulated, implying that they may participate in developmental processes, possibly as signal molecules. While knowledge on the role of pheromones in mating and dikaryon formation is abundant, whether and how they participate in fruiting body development requires further research.

**Pheromone modification -** pheromone precursor proteins are post-translationally modified during their maturation process. This involves prenylated, carboxymethylation as well as proteolytic cleavage to remove N-terminal sequence fragments(Jones and Bennett 2011).

Post-translational prenylation (mostly farnesylation) on CAAX motifs is performed by RAM proteins in *S. cerevisiae*(Caldwell et al. 1995). Based on orthology with *S. cerevisiae* pheromone prenylating enzymes, we identified two genes as putatively involved in pheromone prenylation. The first is orthologous to *S. cerevisiae RAM1* (*C. cinerea* protein ID: 491483) and shows very modest expression dynamics (Table 16), whereas the second is an ortholog of *RAM2* (*C. cinerea* protein ID: 473005), encoding a general prenylation enzyme that also modifies other peptides, including proteins related to cell wall glucan biosynthesis. Following prenylation, the *S. cerevisiae* a-factor is cleaved by Rce1 (ortholog: *C. cinerea* 377901) and Ste24 (ortholog: *C. cinerea* 419650). Orthologs of these genes showed constant expression during fruiting body development (Table 16).

**Table 16.** Summary of developmental expression dynamics of CDE orthogroups of pheromone precursor genes across 12 species. Protein ID of a representative protein is given follows by the number of species in which the orthogroup is developmentally regulated at fold change 2 and 4 (FC>2 and FC>4, respectively). Putative function and ortholog *in S. cerevisiae* (if any) are also given. Abbreviations: 0-gene absent, 1-gene present but not developmentally regulated, 2 - developmentally regulated at fold change >2, 4- developmentally regulated at fold change >4. Species names are abbreviated as: Cc – *C. cinerea*, Aa – *A. ampla*, Ab – *A. bisporus*, Ao – *A. ostoyae*, Ca – *C. aegerita*, Lb – *L. bicolor*, Le – *L. edodes*, Lt – *L. tigrinus*, Mk – *M. kentingensis*, Pc – *Ph. chrysosporium*, Po – *P. ostreatus*, Sc – *S. commune*.

| ID | Fold change |  | Putative function | S.<br>*cerevisiae*<br>ortholog | Species |  |  |  |  |  |  |  |  |  |  |  |
| --- | --- | --- | --- | --- | --- | --- | --- | --- | --- | --- | --- | --- | --- | --- | --- | --- |
|  | >2 | >4 |  |  | Cc | Aa | Ab | Ao | Ca | Lb | Le | Lt | Mk | Pc | Po | Sc |
| 153902 | 2 | 0 | Protein-S-isoprenylcysteine O-methyltransferase | YDR410C | 2 | 1 | 0 | 1 | 2 | 1 | 1 | 1 | 1 | 1 | 1 | 1 |
| 116060 | 1 | 1 | Mating pheromone precursor | – | 4 | 0 | 0 | 0 | 0 | 0 | 0 | 0 | 0 | 0 | 0 | 0 |
| 446846 | 1 | 1 | Mating pheromone precursor | – | 4 | 0 | 0 | 0 | 0 | 0 | 0 | 0 | 0 | 0 | 0 | 0 |
| 446955 | 1 | 1 | Mating pheromone precursor | – | 4 | 0 | 0 | 0 | 0 | 0 | 0 | 0 | 0 | 0 | 0 | 0 |
| 446956 | 1 | 1 | Mating pheromone precursor | – | 4 | 0 | 0 | 0 | 0 | 0 | 0 | 0 | 0 | 0 | 0 | 0 |
| 446958 | 0 | 0 | Mating pheromone precursor | – | 1 | 0 | 0 | 0 | 0 | 0 | 0 | 0 | 0 | 0 | 0 | 0 |
| 446959 | 1 | 1 | Mating pheromone precursor | – | 4 | 0 | 0 | 0 | 0 | 0 | 0 | 0 | 0 | 0 | 0 | 0 |
| 446966 | 0 | 0 | Mating pheromone precursor | – | 1 | 0 | 0 | 0 | 0 | 0 | 0 | 0 | 0 | 0 | 0 | 0 |
| 356637 | 1 | 1 | Pheromone sensing GPCR Ste3 homolog | – | 4 | 0 | 0 | 0 | 0 | 0 | 0 | 0 | 0 | 0 | 0 | 0 |
| 394943 | 4 | 2 | Pheromone sensing GPCR Ste3 homolog | – | 4 | 0 | 0 | 4 | 2 | 1 | 0 | 0 | 0 | 0 | 2 | 0 |
| 477564 | 5 | 3 | Pheromone sensing GPCR Ste3 homolog | – | 4 | 0 | 2 | 0 | 4 | 0 | 0 | 2 | 0 | 0 | 4 | 0 |
| 377901 | 7 | 3 | CAAX prenyl protease 2 | – | 4 | 1 | 2 | 1 | 1 | 2 | 4 | 1 | 4 | 1 | 2 | 2 |
| 203704 | 6 | 3 | Pheromone sensing GPCR Ste3 homolog | – | 4 | 2 | 0 | 4 | 4 | 2 | 1 | 0 | 1 | 1 | 0 | 2 |
| 465723 | 8 | 4 | Pheromone sensing GPCR Ste3 homolog | – | 2 | 1 | 4 | 4 | 2 | 1 | 4 | 2 | 1 | 1 | 4 | 2 |
| 486965 | 5 | 3 | Pheromone sensing GPCR Ste3 homolog | – | 2 | 0 | 0 | 4 | 0 | 4 | 0 | 2 | 1 | 1 | 4 | 0 |
| 491483 | 3 | 2 | Protein farnesyltransferase subunit beta | YDL090C | 4 | 1 | 1 | 1 | 4 | 1 | 1 | 1 | 1 | 1 | 2 | 1 |
| 437675 | 5 | 0 | Pheromone-processing carboxypeptidase KEX1 | YGL203C | 2 | 1 | 1 | 1 | 2 | 2 | 2 | 1 | 1 | 1 | 2 | 1 |
| 545529 | 5 | 1 | Dipeptidyl aminopeptidase B | YHR028C | 1 | 1 | 4 | 2 | 1 | 1 | 1 | 2 | 2 | 1 | 2 | 1 |
| 419650 | 6 | 1 | CAAX prenyl protease 1 ste24 | YJR117W | 4 | 2 | 2 | 1 | 2 | 1 | 1 | 2 | 1 | 1 | 2 | 1 |
| 473005 | 8 | 2 | Protein farnesyltransferase/geranylgeranyltransferase type-1 subunit alpha | YKL019W | 4 | 2 | 1 | 4 | 2 | 2 | 2 | 2 | 1 | 1 | 2 | 1 |
| 540181 | 10 | 4 | Alpha-factor-transporting ATPase | YKL209C | 4 | 4 | 2 | 2 | 2 | 4 | 2 | 2 | 4 | 1 | 1 | 2 |
| 358206 | 8 | 2 | A-factor-processing enzyme | YLR389C | 2 | 1 | 1 | 2 | 2 | 4 | 4 | 1 | 2 | 1 | 2 | 2 |
| 448165 | 5 | 2 | Putative kexin | YNL238W | 1 | 2 | 2 | 4 | 1 | 1 | 1 | 1 | 4 | 1 | 1 | 2 |
| 106286 | 7 | 2 | Pheromone sensing GPCR Ste3 homolog | – | 4 | 1 | 2 | 4 | 2 | 0 | 2 | 2 | 2 | 0 | 0 | 1 |
| 152667 | 9 | 9 | Homolog of *S. cerevisiae* Ste14 | – | 4 | 0 | 4 | 4 | 4 | 4 | 4 | 4 | 1 | 4 | 4 | 0 |

On the other hand, we found evidence for the developmental expression of carboxymethylating enzymes. Two CDE orthogroups of protein-S-isoprenylcysteine O-methyltransferases (homologous to *S. cerevisiae* Ste14) were detected among conserved developmentally expressed genes. Ste14 mediates methylation of the isoprenylated C-terminal CAAX motif in the mating pheromone a-factor(Romano and Michaelis 2001). Whether these homologs are related to mating pheromone maturation in mushroom-forming fungi remains unclear, therefore, we only tentatively classified these as signal transduction-related genes. Two Ste14 homologs of *C. cinerea* were previously found to be expressed after blue light exposure(Sakamoto et al. 2018). We detected two CDE orthogroups of Ste14 homologs (Table 16). One of the orthogroups (represented by *C. cinerea* 152667) is present only in five species and upregulated in primordia of *C. cinerea, A. ostoyae, A. bisporus* and *P. ostreatus* but interestingly shows the reverse pattern, high in vegetative mycelium and no expression in any fruiting body stage in *M. kentingensis*. The other orthogroup (represented by *C. cinerea* 354499) was upregulated in primordia of *C. cinerea, A. ostoyae, Pt. gracilis* and *S. commune (*but not in *A. ampla* and *P. ostreatus).* More broadly, members of this gene family were developmentally regulated in several species, although they were not conserved across species. On the other hand, 1-to-1 orthologs of *S. cerevisiae* Ste14 (represented by *C. cinerea* 153902), show constant expression in all species.

As a final step of a-factor maturation, Ste24 and Axl1/Ste23 proteolytically cleave the N-terminus in two steps(Jones and Bennett 2011). The corresponding Agaricomycete orthologs did not show developmentally dynamics expression (*C. cinerea* protein ID: 419650 and 358206, respectively)(Table 16).

**Pheromone sensing GPCRs -** Seven-transmembrane domain G-protein coupled receptors have been studied in multiple Agaricomycetes(Kües 2000; Raudaskoski and Kothe 2010; Jones and Bennett 2011; Wirth et al. 2021). Pheromone-sensing GPCRs are located in the B mating type locus in Basidiomycota, together with pheromone precursor genes. Pheromone sensing GPCRs transduce signals intracellularly to a mitogen-activated protein kinase (MAPK) cascade (see next chapter). Homology relationships and copy numbers of Basidiomycete pheromone-sensing GPCRs vary significantly across species, but all Basidiomycete pheromone GPCRs are homologous to yeast Ste3, which is involved in mating (reviewed recently(Raudaskoski and Kothe 2010; Fraser et al. 2014)). For example, four pheromone-like GPCR genes have recently been annotated in *S. commune*, of which one was suggested to participate in mating, whereas the others were proposed to have a role in directing growth in vegetative mycelia(Wirth et al. 2021).

We identified 4-19 pheromone GPCR-encoding genes in the examined species. These genes in general showed low expression dynamics in fruiting bodies (Table 16), although several genes were upregulated slightly (FC<4) in stipes in certain species.

**A-factor export proteins** - We identified a CDE orthogroup (represented by *C. cinerea* 540181)(Table 16) of plasma membrane ATPases which are othologous to *S. cerevisiae* Ste6, the protein that exports farnesylated a-factor from the cell. This orthogroup showed partial conservation of expression across the examined species. In *A. ostoyae, L. bicolor, L. edodes* and *M. kentingensis* it was upregulated in stipe tissues relative to caps, whereas in *C. cinerea, S. commune, A. ampla* (FC>4) and *C. aegerita* (though only at FC>2) the gene was upregulated in primordia relative to vegetative mycelium.

#### 4.7.3. MAPK, TOR and cAMP pathways

**MAP kinase pathways -** The mitogen-activated protein kinase (MAPK) cascades are evolutionary conserved intracellular signal transduction pathways, which regulate important cellular processes(Román et al. 2007). The extracellular signal is sensed by membrane proteins or receptors, which is relayed by sequential phosphorylation and activation of three kinases (MAPKKK, MAPKK, MAPK) to transcription factors that induce or repress specific target genes. MAPK cascades usually consist of further components, like various adaptors, docking and scaffold proteins which are implicated in the spatial and temporal regulation of MAP kinase signaling(Frawley and Bayram 2020).

While in Ascomycota MAPK members have been extensively examined (e.g.(Teichert et al. 2014) ), relatively little information is available on the expression and role of MAPK pathways (except the pheromone, see above) in Agaricomycetes. In *Pleurotus tuoliensis* pheromone-dependent and starvation-dependent MAPK signaling pathway members were reported to be up-regulated during fruiting(Fu et al. 2017). Le.MAPK, a MAP kinase of *L. edodes* (Lenedo1_1058304) was reported to be developmentally regulated and one of its interacting partners was characterized(Szeto et al. 2007). In this paper, we annotated Agaricomycetes MAPK signaling pathways based on the four main MAPK pathways known in *S. cerevisiae*: the peptide pheromone pathway (Fus3), the starvation responding pathway (Kss1), the high osmolarity glycerol stress pathway (Hog1) and the cell wall integrity (Slt2/Mpk1) pathway (Table 17). We found relatively few clear orthologs in *C. cinerea* (Table 17). We performed transitive ortholog and domain-based search to find further putative *S. cerevisiae* orthologs in *C. cinerea*. MAPK pathway components do not show dynamic expression patterns during fruiting body development.

**Table 17.**
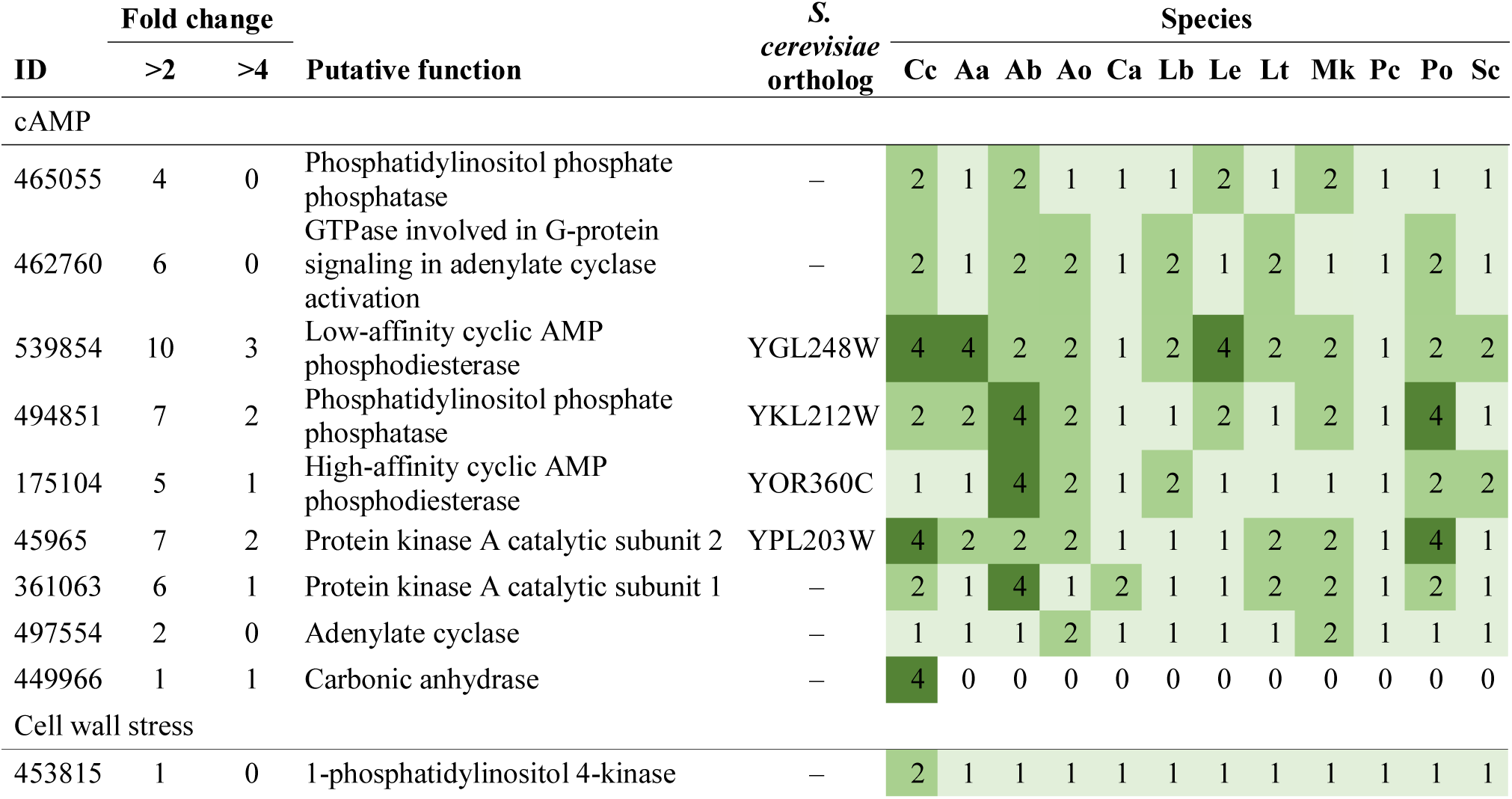
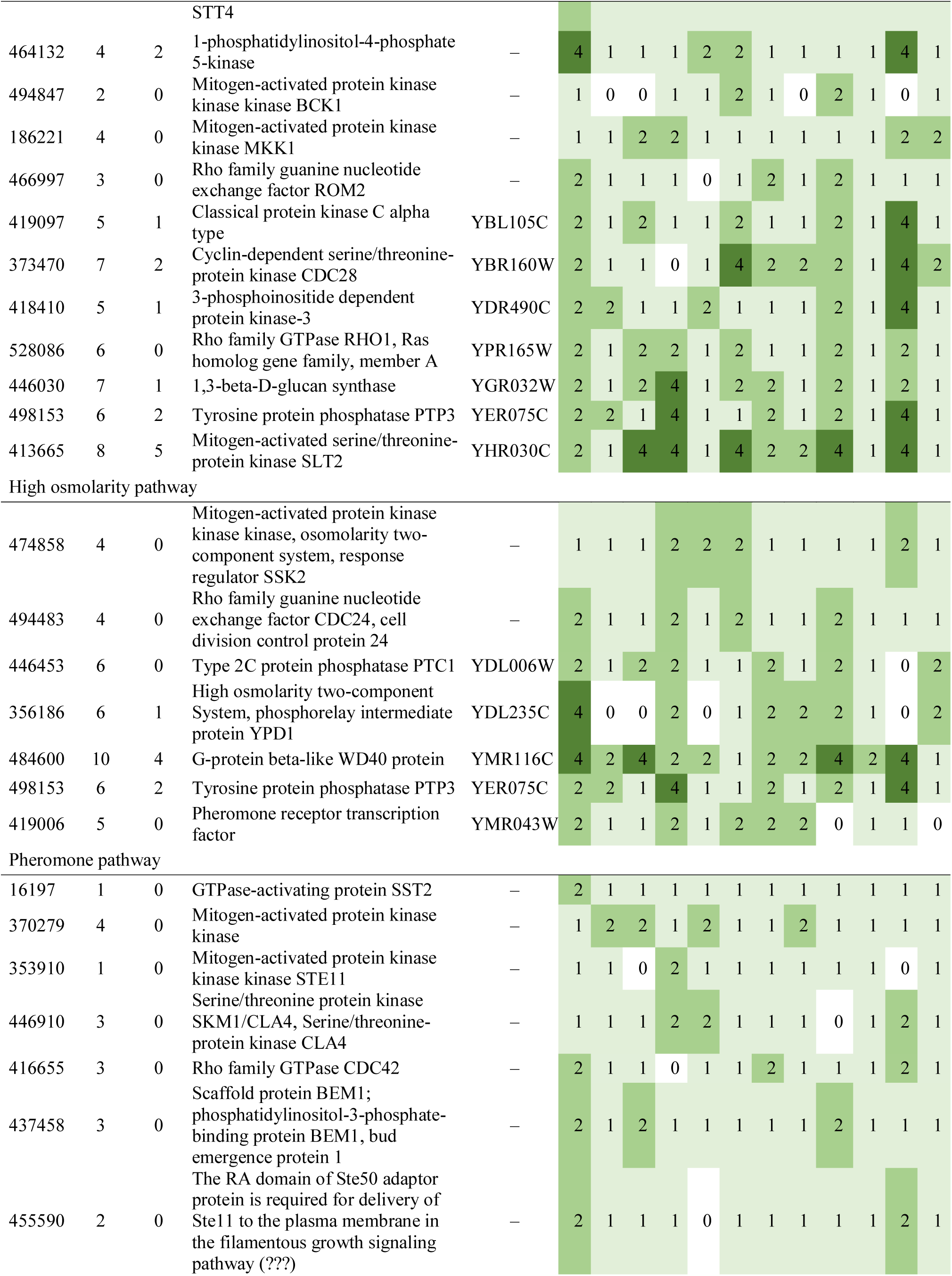
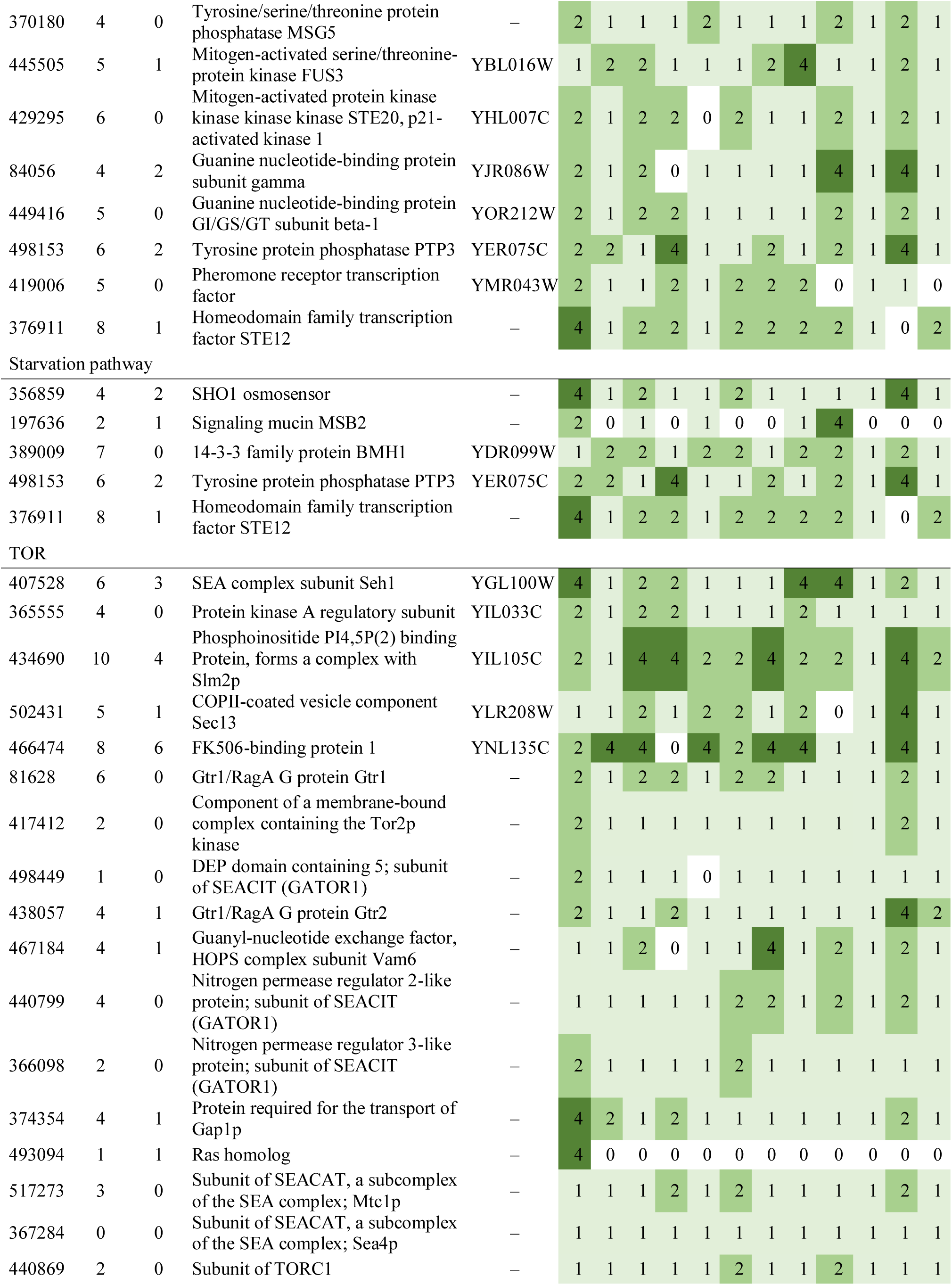
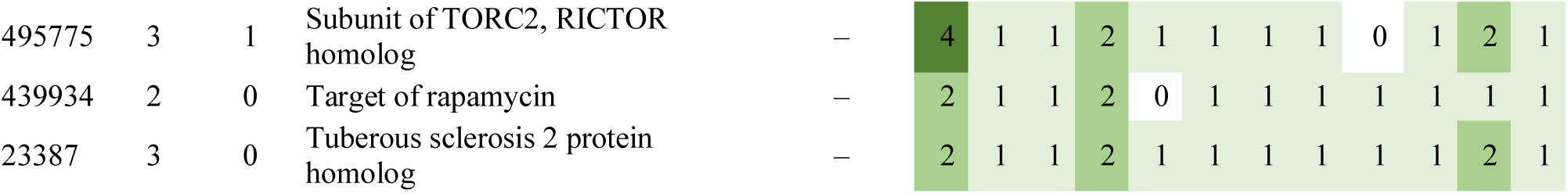
Summary of developmental expression dynamics of CDE orthogroups of conserved signal transduction pathway genes across 12 species. Protein ID of a representative protein is given follows by the number of species in which the orthogroup is developmentally regulated at fold change 2 and 4 (FC>2 and FC>4, respectively). Putative function and ortholog in *S. cerevisiae* (if any) are also given. Abbreviations: 0-gene absent, 1-gene present but not developmentally regulated, 2 - developmentally regulated at fold change >2, 4- developmentally regulated at fold change >4. Species names are abbreviated as: Cc – *C. cinerea*, Aa – *A. ampla*, Ab – *A. bisporus*, Ao – *A. ostoyae*, Ca – *C. aegerita*, Lb – *L. bicolor*, Le – *L. edodes*, Lt – *L. tigrinus*, Mk – *M. kentingensis*, Pc – *Ph. chrysosporium*, Po – *P. ostreatus*, Sc – *S. commune*.

**TOR pathway -** The target of rapamycin (TOR) is an evolutionary conserved serine/threonine kinase that is a key regulator in multiple cellular events(Tatebe and Shiozaki 2017). In fungi, the TOR pathway plays role in the response to nutrients (especially sugar and nitrogen) but also involved in stress responses, such as osmotic and oxidative stresses. TOR signaling pathways are well-studied in yeasts, but data about their functional analysis in multicellular agaricomycetes is rare. In *G. lucidum* TOR signaling was reported to play a role in the regulation of chitin and β-1,3-glucan synthesis and hence of cell wall thickness in an SLT2-MAPK dependent manner(Chen et al. 2019). Orthologs of *Ph. chrysosporium* were previously annotated(Nguyen et al. 2020). According to a 5’-SAGE study of *C. cinerea,* TOR pathway members are constantly expressed during the fruiting body formation, except Rheb (*C. cinerea* protein ID: 493094) a Ras-like small GTPase, which is able to prevent FKBP12 inhibition of TOR(Cheng et al. 2013a).

Here we annotated TOR pathway components in mushroom-forming fungi and could find orthologs for all but three yeast genes reported to participate in the pathway (Table 17). Consistent with previous reports, TOR pathway components showed modest expression dynamics during fruiting body development. Nevertheless, based on the broad role of the pathway in fungal physiology it will be interesting to investigate its connection with fruiting body morphogenesis in the future.

**cAMP pathway -** Adenylyl cyclase dependent pathway (cAMP-pathway) has diverse roles in fungi, including cell growth, metabolism, stress resistance and nutrient-sensing (especially glucose) or starvation in fungi(D’Souza and Heitman 2001). Accordingly, the cAMP pathway has been intensely researched in mushroom-forming fungi(D’Souza and Heitman 2001; Tamaki 2007). Genetic manipulation of several cAMP and connected pathway components in *S. commune* resulted in altered fruiting characteristics, either repression or complete lack of fruiting body formation(Yamagishi et al. 2002, 2004; Knabe et al. 2013; Pelkmans 2016). The cAMP pathway has also been linked to morphogenesis in *F. velutipes:* the deletion of a Gβ-like protein CPC-2 completely impaired fruiting body formation, but the addition of cAMP or it analog 8-Bromo-cAMP (also a PKA-activator) into the medium restored fruiting in *Fv.cpc2* knockdown mutants(Wu et al. 2020b). Ras and G-protein □ subunits and their regulators (e.g. *Thn1*) might also be connected to the cAMP pathway(Yamagishi et al. 2002; Palmer and Horton 2006; Knabe et al. 2013) and were repeatedly reported to affect morphogenesis of fruiting bodies (e.g.(Schubert et al. 2006)).

In general, cAMP is a known inducer of fruiting in several basidiomycetes, such as *C. macrorhizus*(Uno and Ishikawa 1973), *C. cinerea*(Swamy et al. 1985)*, S. commune*(Schwalb 1974) and *Ph. chrysosporium*(Gold and Cheng 1979). In the latter, cAMP was speculated to induce fruiting by overriding the repressive effect of glucose (i.e. mimicked starvation in the fungus(Gold and Cheng 1979)). More recent studies in *S. commune* reported a repressing effect of cAMP on fruiting, and suggested this was because cAMP is able to mimic high CO_2_ levels(Pelkmans 2016). Genes of the cAMP pathway were annotated in *S. commune* and the pde2 phosphodiesterase (Schco3_2636760), which can degrade cAMP was functionally analyzed(Pelkmans 2016). The overexpression of pde2 results in fruiting body development even at elevated CO_2_ levels. These results were interpreted as evidence for CO_2_ being sensed via the cAMP pathway(Pelkmans 2016).

We annotated members of the cAMP pathway with *C. cinerea* as reference (Table 17). Consistent with data in *S. commune*(Pelkmans 2016), we could identify most members of the cAMP pathway in the examined species. Our expression data revealed that cAMP pathway members show relatively little expression dynamics in mushroom-forming fungi (Table 17). Nevertheless, this does not preclude an important role of cAMP signaling in fruiting body formation.

#### 4.7.4. G-protein coupled receptors

G-protein coupled receptors (GPCRs) comprise the largest receptor family in fungi. GPCRs are seven transmembrane cell surface receptors, which transduce signals from a wide range of external stimuli by a G-protein heterotrimer(Xue et al. 2008). Fungal GPCRs originally divided into six classes based on their structural similarity and homology(Xue et al. 2008), however, this classification was later expanded with additional classes (reviewed in (Brown et al. 2018)).

Pheromone sensing GPCRs are among the most widely studied in connection with fruiting body development. In *Flammulina filiformis* six pheromone receptor-like genes were characterized whereby four were highly expressed in the fruiting body, while two exhibited the maximal expression level in the mycelia(Meng et al. 2020). Six pheromone-like GPCRs were characterized also in *S. commune* (*bar3, bbr2, brl1*, *brl2*, *brl3* and *brl4*); it was reported that the overexpression of *brl2, brl3* and *brl4* enhanced mating or fruiting body formation in this species(Wirth et al. 2021). We annotated 7 mating-type pheromone receptors in *C. cinerea* (Table 16). Most of the pheromone receptors are highly expressed during the fruiting, but do not have orthologs in other species. In addition, five further GPCR genes were detected, which, similarly to pheromone-sensing GPCRs mostly did not have orthologs in other species (Table 18). This may suggest that fungal species have a specialised GPCR repertoire, that is hard to examine using orthology-based approaches.

**Table 18.**
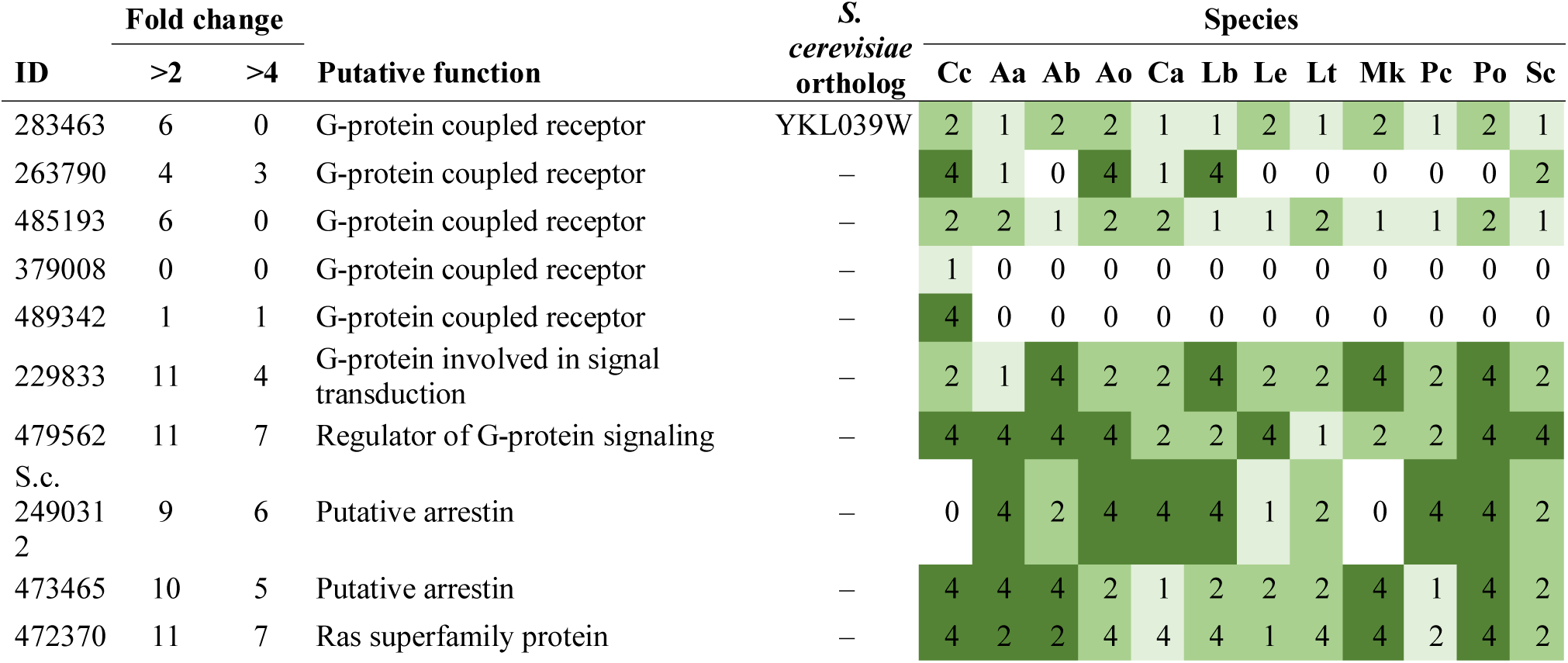
Summary of developmental expression dynamics of CDE orthogroups of G-protein coupled receptor genes across 12 species. Protein ID of a representative protein is given follows by the number of species in which the orthogroup is developmentally regulated at fold change 2 and 4 (FC>2 and FC>4, respectively). Putative function and ortholog in *S. cerevisiae* (if any) are also given. Abbreviations: 0-gene absent, 1-gene present but not developmentally regulated, 2 - developmentally regulated at fold change >2, 4- developmentally regulated at fold change >4. Species names are abbreviated as: Cc – *C. cinerea*, Aa – *A. ampla*, Ab – *A. bisporus*, Ao – *A. ostoyae*, Ca – *C. aegerita*, Lb – *L. bicolor*, Le – *L. edodes*, Lt – *L. tigrinus*, Mk – *M. kentingensis*, Pc – *Ph. chrysosporium*, Po – *P. ostreatus*, Sc – *S. commune*.

While GPCRs are not conserved, we detected three CDE orthogroups that may be related to GPCR signaling and were developmentally regulated across several species. An RGS (regulator of G-protein signaling) domain containing protein family (*C. cinerea* protein ID: 479562) was developmentally regulated in 11/7 species (FC>2/4)(Table 18). RGS proteins are involved in signal transduction by regulating G-protein activity and rapidly switching on or off G-protein coupled receptors pathways. We also found a CDE orthogroup containing putative Ran-binding proteins (*C. cinerea* protein ID: 472370), which are members of the Ras superfamily that regulates all receptor-mediated transport between the nucleus and the cytoplasm. Finally, we detected two arrestin superfamily orthogroups (*C. cinerea* protein ID: 473465, *S. commune:* Schco3_2490312) (Table 18). Arrestins are involved in signal transduction through the inactivation of G protein-coupled receptors, but some members of the superfamily have been reported in receptor endocytosis, cell cycle progression and pH sensing as well also(Herranz et al. 2005; Telzrow et al. 2019).

#### 4.7.5. Other signal transduction related families

**Kinases -** The majority of CDE orthogroups classified as signal transduction were kinases (Table 15). We found 13 conserved and developmentally expressed kinase orthogroups, however, most of these proved difficult to functionally characterize, likely due to the high rate of evolution in kinases and the lack of orthology to well-known genes in model species. One exception is the CDE orthogroup (*C. cinerea* protein ID: 451611) that contains orthologs of *S. cerevisiae* Rck2, which is a member of the high osmolarity glycerol (HOG) MAPK pathway. The HOG pathway is involved in responding to oxidative and osmotic stress. In most species, Rck2 orthologs showed early upregulation, with highest expression values in primordia, though its functional significance is not known. Rck2 orthologs were differentially expressed between dikaryons and A and B type monokaryons of *C. cinerea* as well(Stajich et al. 2010), and it was concluded that RCK2 genes in *C. cinerea* must serve different functions from that of *S. cerevisiae* orthologs. The remaining kinase orthogroups are functionally uncharacterized and may be interesting targets for functional studies, their dynamic expression patterns indicate potentially conserved and important functions in fruiting body development.

**Velvet complex** - The velvet complex has been identified as a central regulatory hub in fungal development and secondary metabolism(Bayram et al. 2008). The complex was characterized in the Ascomycota, where it consists of VeA, LaeA, VelB as well as VelC and VosA, which assemble into condition-specific (including light-responsive) complexes. It has been shown that the velvet domain, an ∼200 amino acid fold, binds DNA and resembles the structure of the metazoan transcription factors NF-κB(Ahmed et al. 2013). Proteins containing a velvet domain exist outside the Pezizomycotina as well, however, these proteins did not appear to be orthologous to members of the Velvet complex in the aspergilli(Ojeda-López et al. 2018); accordingly, how they function and whether they form complexes also in the Basidiomycota and early-diverging fungi is not known.

The first evidence on a potential role of Velvet proteins in Basidiomycota fruiting body development was provided by Plaza et al(Plaza et al. 2014b). They detected the upregulation of a velvet domain containing protein in primordia of *C. cinerea* relative to vegetative mycelia. Following these trails and combining detailed mechanistic knowledge from the velvet regulon of *A. nidulans*(Krijgsheld et al. 2013), they could show that some, albeit a minor proportion, of orthologs of the *A. nidulans* velvet-regulated genes are upregulated in fruiting bodies in *C. cinerea, L. bicolor* and *S. commune*(Plaza et al. 2014b) also. This suggested some extent of conservation of function in the Dikarya, although the lack of orthology between Asco- and Basidiomycota proteins casts some uncertainty on how these data should be interpreted. Velvet protein-encoding genes were reported to be developmentally regulated in multiple Agaricomycetes species in the comparative analyses of Krizsan et al(Krizsán et al. 2019).

Our current analysis indicated that the velvet domain occurs in 5-6 proteins (only 4 in *P. ostreatus*) encoded in the genomes of mushroom-forming fungi. Surprisingly, the yeast-like *C. neoformans* also possess 6 genes with a velvet domain, in contrast to Ascomycota yeasts, which have lost velvet genes(Nagy et al. 2014). Interestingly, Ascomycota on average have 4 genes that encode velvet-domain containing proteins, indicating an expansion in the Basidiomycota. Consistent with results of Ojeda-Lopez et al(Ojeda-López et al. 2018), we found that Basidiomycota velvet proteins lack clear orthology to Ascomycota genes. Only VelB and VosA of *A. nidulans* have reciprocal best hits in *C. cinerea* (*C. cinerea* protein ID: 365674 and 374867, respectively).

In terms of expression, observed patterns vary widely among species. For example, all six genes are developmentally regulated in *C. cinerea*, but none in *A. ostoyae* (both FC>4), indicating considerable expression plasticity through evolution. We identified three orthogroups whose members were developmentally regulated in a notable number of species (Table 19). Of these, genes in one orthogroup (represented by *C. cinerea* 385561) were mostly upregulated in stipe tissues in the pileate-stipitate species *C. cinerea, A. ostoyae, M. kentingensis*, *L. edodes* (but only in mature FBs of *L. bicolor*). The gene was not developmentally regulated in *L. tigrinus* and *P. ostreatus*, possibly indicating the distant relationship of these species from ones in which the gene is stipe-upregulated. The second most interesting CDE orthogroup (*C. cinerea* protein ID: 496341) was developmentally regulated in 10 species (FC>2), but expression peaks varied from primordia (*A. ampla, S. commune, A. ostoyae*), to young fruiting body caps (*C. cinerea*) and stipes (*M. kentingensis*).

**Table 19.**
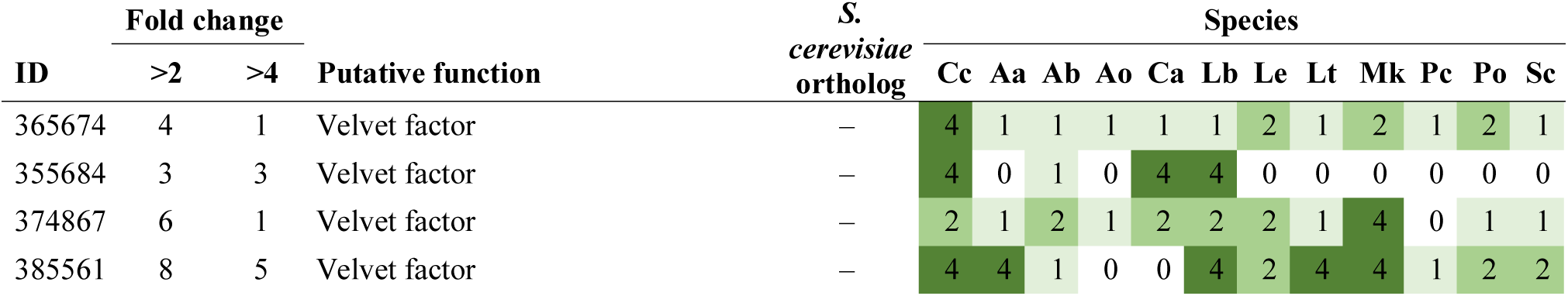
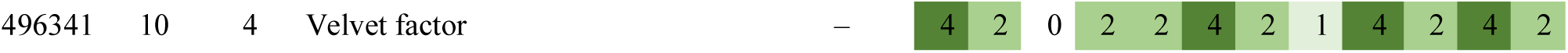
Summary of developmental expression dynamics of CDE orthogroups of velvet protein encoding genes across 12 species. Protein ID of a representative protein is given follows by the number of species in which the orthogroup is developmentally regulated at fold change 2 and 4 (FC>2 and FC>4, respectively). Putative function and ortholog in *S. cerevisiae* (if any) are also given. Abbreviations: 0-gene absent, 1-gene present but not developmentally regulated, 2 - developmentally regulated at fold change >2, 4- developmentally regulated at fold change >4. Species names are abbreviated as: Cc – *C. cinerea*, Aa – *A. ampla*, Ab – *A. bisporus*, Ao – *A. ostoyae*, Ca – *C. aegerita*, Lb – *L. bicolor*, Le – *L. edodes*, Lt – *L. tigrinus*, Mk – *M. kentingensis*, Pc – *Ph. chrysosporium*, Po – *P. ostreatus*, Sc – *S. commune*.

Based on the above data, velvet protein-encoding genes represent an interesting target for phylogenetic and functional studies of sexual fruiting bodies in the Agaricomycetes; we expect their analyses will reveal interesting insights into fruiting body development.

**STRIPAK complex -** The <u>st</u>riatin-interacting <u>p</u>hosphatase <u>a</u>nd <u>k</u>inase, STRIPAK (known as Far complex in(Kück et al. 2019)), is a widely conserved multi-protein complex that can be found from yeast to metazoans(Reschka et al. 2018). Its function in metazoans is unknown, whereas in Ascomycota model systems (e.g. *Sordaria macrospora*) it is involved in hyphal fusion and fruiting body development. It was first discovered in 2004 in fungi(S and U 2004) and its function has been increasingly clarified in a series of papers since then (see in(Reschka et al. 2018; Riquelme et al. 2018)). Mechanistically, the STRIPAK complex provides links between multiple signaling pathways in eukaryotic cells (reviewed in(Kück et al. 2019)), although many of the details of its interactions with other proteins are not known. Its role in fruiting body formation in the Basidiomycota has not yet been investigated.

We queried the *C. cinerea* genome with members of the STRIPAK complex described from *Sordaria* and *S. cerevisiae* (listed in(Reschka et al. 2018)). *C. cinerea* seems to have a fully functional STRIPAK complex, with all genes having clear orthologs, except for Far3/7 which only had weak hits. Further, the *C. cinerea* proteins were members of conserved single-copy Agaricomycetes orthogroups, which indicated that the complex is conserved in all the examined Agaricomycetes species. Members of this complex did not show significant expression dynamics (Table 20). The conservation of the complex in the Agaricomycetes suggests it might have important functions, however, its role in fruiting body development remains unclear at the moment and requires further research.

**Table 20.** Summary of developmental expression dynamics of CDE orthogroups of STRIPAK complex protein genes across 12 species. Protein ID of a representative protein is given follows by the number of species in which the orthogroup is developmentally regulated at fold change 2 and 4 (FC>2 and FC>4, respectively). Putative function and ortholog in *S. cerevisiae* (if any) are also given. Abbreviations: 0-gene absent, 1-gene present but not developmentally regulated, 2 - developmentally regulated at fold change >2, 4- developmentally regulated at fold change >4. Species names are abbreviated as: Cc – *C. cinerea*, Aa – *A. ampla*, Ab – *A. bisporus*, Ao – *A. ostoyae*, Ca – *C. aegerita*, Lb – *L. bicolor*, Le – *L. edodes*, Lt – *L. tigrinus*, Mk – *M. kentingensis*, Pc – *Ph. chrysosporium*, Po – *P. ostreatus*, Sc – *S. commune*.

| ID | Fold change |  | Putative function | S.<br>*cerevisiae*<br>ortholog | Species |  |  |  |  |  |  |  |  |  |  |  |
| --- | --- | --- | --- | --- | --- | --- | --- | --- | --- | --- | --- | --- | --- | --- | --- | --- |
|  | >2 | >4 |  |  | Cc | Aa | Ab | Ao | Ca | Lb | Le | Lt | Mk | Pc | Po | Sc |
| 51761 | 1 | 0 | Forkhead-associated domain | – | 2 | 1 | 1 | 1 | 1 | 1 | 1 | 1 | 1 | 1 | 1 | 1 |
| 439971 | 1 | 0 | Protein phosphatase PP2A regulatory subunit A | YAL016W | 1 | 1 | 1 | 1 | 1 | 1 | 1 | 1 | 1 | 1 | 2 | 1 |
| 280222 | 5 | 0 | Serine/threonine-protein kinase KIC1 | YHR102W | 1 | 1 | 1 | 2 | 2 | 2 | 1 | 2 | 1 | 1 | 2 | 1 |
| 375927 | 4 | 0 | Factor arrest protein 11 | YNL127W | 2 | 1 | 1 | 2 | 1 | 1 | 2 | 1 | 1 | 1 | 2 | 1 |
| 463106 | 5 | 2 | Serine/threonine-protein phosphatase PP2A-like PPG1 | YNR032W | 4 | 1 | 4 | 1 | 1 | 1 | 2 | 2 | 1 | 1 | 2 | 1 |
| 499433 | 8 | 2 | MOB kinase activator | – | 4 | 2 | 2 | 1 | 2 | 1 | 1 | 4 | 2 | 2 | 2 | 1 |
| 108726 | 5 | 0 | Striatin | – | 2 | 1 | 2 | 2 | 1 | 1 | 1 | 2 | 1 | 1 | 2 | 1 |

#### 4.7.6. Light sensing systems and light receptors

Fungi adapt developmental processes to light signals in time and space to ensure that sporulating cells can maximize spore dispersal efficiency(Yu and Fischer 2018). The photobiology and its connection to developmental events are quite well-known in both the Asco- and Basidiomycota. In the last decade, the molecular architectures and regulatory networks of sensing and responding to blue, red and green light have been uncovered in multiple species(Corrochano 2011; Fuller et al. 2015). In this chapter we discuss light sensing systems in Agaricomycetes and analyze their expression patterns in fruiting bodies. We note that the best known such system, the white collar complex has transcription factor activity and is discussed under chapter 4.5.1.

**Cryptochromes** - Fungal genomes include DASH-type cryptochromes, photolyase genes that function as blue light photoreceptors and are involved in DNA repair(Fuller et al. 2015). Cryptochromes are regulated by the white collar complex in *N. crassa*(Froehlich et al. 2010) and regulate fruiting body development in certain Ascomycota (e.g. *Cordyceps militaris*(F et al. 2017)), but to our best knowledge, no such information is available in the Basidiomycota. Our analysis suggests that cryptochromes are single copy genes in the Agaricomycetes, except in *L. edodes*, which has two genes. The CDE orthogroup containing Agaricomycetes cryptochromes (represented by *C. cinerea* 492962)(Table 21) was developmentally regulated in 9/7 species (FC>2/4), and showed strong induction in primordia relative to vegetative mycelium in *A. ostoyae, M. kentingensis, L. edodes, P. ostreatus, Ph. chrysosporium* and at FC>2 in *C. aegerita* and *C. cinerea*. The gene was constantly expressed in *A. ampla* and *S. commune* and was missing in *L. bicolor.* Based on these data, cryptochromes may be relevant to fruiting body formation in the Agaricomycetes.

**Table 21.**
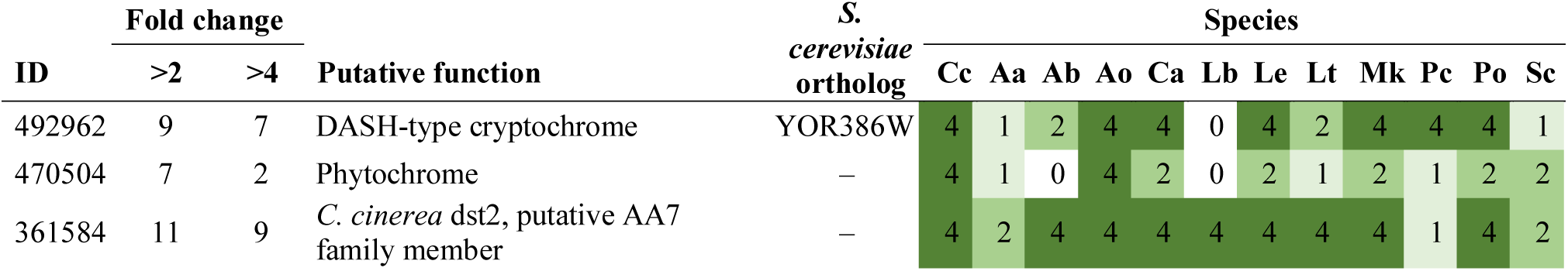
Summary of developmental expression dynamics of CDE orthogroups of light sensing related genes across 12 species. Protein ID of a representative protein is given follows by the number of species in which the orthogroup is developmentally regulated at fold change 2 and 4 (FC>2 and FC>4, respectively). Putative function and ortholog in *S. cerevisiae* (if any) are also given. Abbreviations: 0-gene absent, 1-gene present but not developmentally regulated, 2 - developmentally regulated at fold change >2, 4- developmentally regulated at fold change >4. Species names are abbreviated as: Cc – *C. cinerea*, Aa – *A. ampla*, Ab – *A. bisporus*, Ao – *A. ostoyae*, Ca – *C. aegerita*, Lb – *L. bicolor*, Le – *L. edodes*, Lt – *L. tigrinus*, Mk – *M. kentingensis*, Pc – *Ph. chrysosporium*, Po – *P. ostreatus*, Sc – *S. commune*.

**Phytochromes** are red- and far-red receptors that influence sexual and asexual sporulation in fungi. The first circumscribed fungal phytochrome was the *fphA* gene in *A. nidulans*, which represses sexual development under red light(A et al. 2005). Since then, the repression of sexual development has been proven for *N. crassa* too and several details of the process (e.g. interaction with WCC and velvet factors) have been clarified(A and J 2005; Wang et al. 2016; Yu et al. 2016). In the Basidiomycota, the few reports of phytochromes include a systematic mutant library in *C. neoformans*, which circumscribed the gene encoding *TCO3* (Lee et al. 2016), as well as observations of phytochrome upregulation in browning mycelium of *L. edodes*(Tang et al. 2013) and in rhizomorphs of *A. ostoyae*(Sipos et al. 2017b). In *U. maydis* red and blue light receptors have been shown to be required for the development of fruiting body-like structures(Sánchez-Arreguin et al. 2020).

Phytochromes seem to be single-copy genes in the Agaricomycetes examined here. They formed a conserved orthogroup (represented by *C. cinerea* 470504, Table 21), however, genes in this orthogroup did not show significant expression dynamics during fruiting body formation, except in *A. ostoyae* and *C. cinerea*. In *A. ostoyae* the expression of the phytochrome gene was ∼10x higher in stipes than caps in stage 2 primordia and young fruiting bodies. Similarly, in *C. cinerea* peak expression was observed in elongating stipes in the young fruiting body stage. However, other species did not display this expression profile, indicating that phytochrome expression is not conserved across mushroom-forming fungi. Although expression patterns did not seem to be conserved in fruiting bodies, it is possible that red light sensing phytochromes are part of a photoreceptor system that signals position in open air, which is required for sporulation and fruiting body formation for the majority of species(J et al. 2010; Fuller et al. 2015).

Fungal light sensing systems also include rhodopsins, seven-transmembrane domain proteins involved in sensing green light(Bieszke et al. 1999). We detected rhodopsins only in *L. tigrinus* and *Ph. chrysosporium*, but none of the Agaricales in our dataset.

**Genes related to blue light receptors -** Blue light is one of the most important signals for fungi and responses as well as molecular aspects of the fungal blue light response are among the best known aspects of fungal photobiology in both the Asco- and Basidiomycota. The most important receptors of blue light are the white collar complex (WCC), which comprises the photoreceptors/transcription factors WC2 and WC1. These have been discussed under the chapter on transcription factors (see above). We here only discuss putative accessory proteins similar to VIVID of *N. crassa*. These proteins harbor a single PAS domain and act as repressors of the WCC in *N. crassa* and were suggested to provide a mechanism for adapting to changing light intensities during the day(Fuller et al. 2015). Although evidence for that is missing in the Agaricomycetes, a CDE orthogroup of PAS domain proteins (represented by *C. cinerea* 500409) was developmentally regulated in 9/7 species (FC>2/4) and showed strong induction in primordia relative to vegetative mycelia in *A. ostoyae, C. aegerita, C. cinerea, L. bicolor, L. tigrinus, M, kentingensis, Ph. chrysosporium* but not in *A. ampla* and *P. ostreatus* (and missing from *L. edodes* and *S. commune*). It will be interesting to see if this or similar genes are involved in light responses in the Agaricomycetes.

**Dst2 family -** We found that orthologs of the dst2 gene of *C. cinerea* formed a CDE orthogroup (represented by *C. cinerea* 361584). Dst2 has been described as a FAD-binding and Berberine-like domain containing protein that, if mutated, results in a blind dark stipe phenotype similar to what is observed in mutations on the white collar complex(Kamada et al. 2010; Kuratani et al. 2010). Based on these observations, it has been hypothesized that dst2 represents a novel photoreceptor(Kuratani et al. 2010). The typical expression profile of these genes (seen in *A. ostoyae, C. aegerita, C. cinerea, L. edodes* and *P. ostreatus*) involved upregulation in primordia relative to mycelium and, in later stages, higher expression in stipes compared to caps. Variations to this pattern exist, such as only being FB-init in *L. bicolor* and *S. commune*, or being only stipe upregulated but not FB-init in *M. kentingensis* and *L. tigrinus*. We hypothesize that dst2 genes might be related to stipe development and, potentially, photomorphogenesis.

### 4.8. Cell wall biosynthesis and modification

The cell wall is an organelle that functions as an extracellular matrix in fungi. It provides integrity, rigidity to cells, defines cell shape, harbors extracellular proteins, represents a channel of communication with the environment, and provides the first line of physical defense, to name just a few of its major roles in fungal physiology. Structural roles are particularly important for the Agaricomycetes that form above ground fruiting bodies that should be rigid yet plastic enough to withstand environmental forces and support spore dispersal. In fruiting bodies, the cell wall must serve structural and defense roles but must also be sufficiently plastic to allow changes in shape and size during development. Although mechanisms of cell shape determination are not known in mushroom-forming fungi, regulated cell wall deposition and remodeling by hydrolytic enzymes residing in the wall itself(Verdín et al. 2019), are undoubtedly of key importance.

There is plenty of evidence for cell wall remodeling and differential cell wall compositions in fruiting bodies of various species (e.g. (Buser et al. 2010a; Almási et al. 2019; Krizsán et al. 2019; Zhou et al. 2019)), although most of that is based on transcriptome rather than functional studies. For example, stipe growth may be realized by the breaking of chitin-glucan cross-links by hydrolytic enzymes(Liu et al. 2021). Similarly, there are cross-linking (transglycosidases) and lytic (e.g. glycoside hydrolases) enzymes that rigidify and make the cell wall more plastic, respectively(Verdín et al. 2019). Mannosylation might also rigidify the cell wall and mannoproteins are often N-glycosylated, which contributes to water retention and adhesion(Lesage and Bussey 2006). In the Agaricomycetes, Buser et al reported a fruiting body-specific N-glycan from *C. cinerea*(Buser et al. 2010b). The regulated interplay of these enzymes might result in the formation of stage-, tissue-, and species-specific cell wall architectures in fruiting bodies.

In this chapter we review current knowledge on cell wall related genes in mushroom-forming fungi and analyze CDE orthogroups that are likely linked to cell wall synthesis or modification. We will not review cell wall composition in fruiting bodies, but refer to earlier reviews(Kües 2000). Cell wall functioning is quite well-researched in the Ascomycota (e.g. in *S. cerevisiae* and *N. crassa*), and to a much smaller extent in the Basidiomycota. We therefore base much of our functional hypotheses on knowledge in the Ascomycota, noting that the process may be different and/or significantly more complex in the Basidiomycota. For example, while some chitin-glucan cross linking enzyme families appear similarly important both in Asco- and Basidiomycota (e.g. GH16/GH17), there are some (e.g. GH72 and GH76) which are involved in cell wall remodeling in the Ascomycota, but apparently not in the Basidiomycota. Hereafter we discuss cell wall related CDE orthogroups broken down into synthetic and remodeling enzymes as well as by substrate (chitin/glucan).

From these analyses a few broad observations emerge, which we discuss here. Most importantly, the diversity of enzymes that show developmental regulation in fruiting bodies suggests that the fruiting body cell wall may be significantly more complex than we think, and we still know quite little about its exact structure and composition. A general pattern that also arises is that more tissue- and stage-resolved transcriptomes reveal proportionately more developmentally expressed genes, which indicates that cell wall remodeling and, consequently, cell wall architectures, likely show high tissue-specificity in fruiting bodies. Based on our results, it appears that fungal cell wall (FCW) remodeling and crosslinking genes show significantly more expression dynamics than genes encoding chitin and glucan synthases. In line with this observation, we found that intracellular components of cell wall biosynthesis (chaperones, glucan synthase complex, pathways involved in glucan monomer synthesis) also show little expression dynamics.

Comparisons across species revealed limited conservation in FCW related gene expression patterns across species, which suggests it diverges rapidly during evolution. This lends support to the quick evolution of cell wall architectures and corresponding enzyme profiles in fruiting bodies. Another observation related to this is that FCW-related genes show complex gene duplication/loss patterns, which complicates the inference of well-conserved groups of 1-to-1 orthologous genes. Thus, these genes might be more appropriately analyzed in the context of broader orthogroups (e.g. based on Orthofinder(Emms and Kelly 2015)) than based on the strict 1-to-1 orthogroups we used. However, for the sake of uniformity, we stick to the latter, with notes on overall prevalence of developmental expression in a given family.

An interesting observation is that several FCW active genes upregulated in fruiting bodies are also expressed during mycorrhiza establishment in ECM fungi such as *L. bicolor*(Martin et al. 2008; C et al. 2014). Such signals provide clues to shared morphogenetic mechanisms between ectomycorrhizal development and fruiting bodies.

Finally, an important consideration for cell wall biosynthesis/remodeling genes is whether they are upregulated in fruiting bodies is the vegetative mycelium sample that was used as control in the RNA-Seq. Non-growing inactive hyphae (where cell wall synthesis rate is low) as control of actively growing fruiting bodies can yield more gene expression differences than an actively growing mycelium sample. Although we do not think this has significantly influenced our comparisons, because we do not know the sampling conditions for all species, this possibility should be kept in mind in the interpretation of results.

We attempted to as comprehensively analyze cell wall related gene expression as possible, at least with consideration of genes that show some expression conservation at the gene family level. Nevertheless, it is certain that several important, especially species-specific genes are left out from our inventory. For example, mannosylation, a potentially interesting process contributing to cell wall functioning, is not discussed here due to the lack of orthology to experimentally characterized mannosylation components in the Ascomycota. These aspects will need to be covered in future studies.

#### 4.8.1. Cell wall biosynthesis

##### 4.8.1.1. Chitin biosynthesis (GT2)

Chitin is one of the most important, although usually not the most abundant structural polymer of the fungal cell wall. This is true for fruiting bodies as well, in which, although cell wall composition is altered relative to vegetative mycelia(M and C 1981; Mol and Wessels 1990; Kües 2000), chitin is an important component that confers rigidity. Chitin is also a scaffold for the modification of cell wall structure either by cross-linking, incorporation of glycoproteins or chemical modifications to the chitin polymers, among others(Verdín et al. 2019). Cell wall chitin assembly involves the action of the UDP-N-acetylglucosamine biosynthesis pathway (in *S. cerevisiae* Gfa1p, Gna1p, Pcm1p, Qri1p), chitin synthases (Chs1-3) which elongate the growing chitin polymer as well as chitin deacetylases, which modify the chitin on the cell surface(Shahinian and Bussey 2000; Klis et al. 2006; Verdín et al. 2019) (the latter is discussed in chapter 4.8.2.1). While yeasts have up to three chitin synthases, filamentous fungi have additional chitin synthase genes (on average up to seven(Verdín et al. 2019)), including highly processive ones that harbor a myosin motor domain and that were lost in yeasts(N et al. 2006).

We examined chitin synthesis and modification related genes based primarily on functional annotations and recent reviews(Shahinian and Bussey 2000; Klis et al. 2006; Verdín et al. 2019). Clear orthologs of the *S. cerevisiae* UDP-N-acetylglucosamine biosynthesis pathway could be identified in all examined Agaricomycetes species (Table 22). We identified 8-13 (17 in *L. bicolor*) chitin synthases of the GT2 family, 2-3 chitin synthase export chaperones as well as orthologs of *S. cerevisiae* SKT5 (activator of chitin synthase 3) in the examined Agaricomycetes. Interestingly, UDP-N-acetylglucosamine biosynthesis pathway genes showed more expression dynamics than most chitin synthase genes (Table 22). Among chitin synthases, although many formed conserved orthogroups, only a single appeared developmentally regulated in a wide array of species. Nevertheless, each species had developmentally regulated and/or FB-init (sensu Krizsan et al 2019) chitin synthases, though these did not always belong in conserved orthogroups. The most interesting orthogroup (represented by *C. cinerea* 359180) was developmentally regulated in 7/6 (at fold change 2/4) species and showed upregulation and high expression in primordia relative to vegetative mycelium in several species (*C. cinerea, A. ampla, P. ostreatus, S. commune, C. aegerita* and *L. edodes*). Comparative genomic data suggests that this orthogroup of chitin synthases is specific to mushroom-forming fungi or the Basidiomycota, suggesting that it may be a fruiting body-specific chitin synthase. A second CDE orthogroup of GT2 chitin synthases (*C. cinerea* protein ID: 519767) was also detected (Table 22).

**Table 22.** Summary of developmental expression dynamics of CDE orthogroups of chitin biosynthesis genes across 12 species. Protein ID of a representative protein is given follows by the number of species in which the orthogroup is developmentally regulated at fold change 2 and 4 (FC>2 and FC>4, respectively). Putative function and ortholog in *S. cerevisiae* (if any) are also given. Abbreviations: 0-gene absent, 1-gene present but not developmentally regulated, 2 - developmentally regulated at fold change >2, 4- developmentally regulated at fold change >4. Species names are abbreviated as: Cc – *C. cinerea*, Aa – *A. ampla*, Ab – *A. bisporus*, Ao – *A. ostoyae*, Ca – *C. aegerita*, Lb – *L. bicolor*, Le – *L. edodes*, Lt – *L. tigrinus*, Mk – *M. kentingensis*, Pc – *Ph. chrysosporium*, Po – *P. ostreatus*, Sc – *S. commune*.

| ID | Fold change |  | Putative function | S.<br>*cerevisiae*<br>ortholog | Species |  |  |  |  |  |  |  |  |  |  |  |
| --- | --- | --- | --- | --- | --- | --- | --- | --- | --- | --- | --- | --- | --- | --- | --- | --- |
|  | >2 | >4 |  |  | Cc | Aa | Ab | Ao | Ca | Lb | Le | Lt | Mk | Pc | Po | Sc |
| 371191 | 9 | 4 | Glucosamine 6-phosphate N-acetyltransferase | – | 4 | 2 | 4 | 4 | 1 | 2 | 2 | 2 | 2 | 1 | 4 | 1 |
| 538650 | 6 | 0 | Chitin synthase regulator SKT5 | YBL061C | 2 | 1 | 2 | 2 | 2 | 1 | 1 | 2 | 2 | 1 | 1 | 1 |
| 542039 | 8 | 1 | Chitin synthase 3 | YBR023C | 2 | 2 | 2 | 4 | 2 | 2 | 1 | 1 | 2 | 1 | 2 | 1 |
| 357695 | 7 | 5 | UDP-N-acetylglucosamine pyrophosphorylase | YDL103C | 4 | 2 | 4 | 4 | 4 | 1 | 1 | 2 | 1 | 1 | 4 | 1 |
| 478844 | 4 | 1 | Phosphoacetylglucosamine mutase involved in chitin precursor synthesis | YEL058W | 4 | 1 | 1 | 2 | 2 | 1 | 1 | 1 | 1 | 1 | 2 | 1 |
| 440525 | 9 | 4 | Glutamine-fructose-6-phosphate aminotransferase | YKL104C | 4 | 1 | 4 | 4 | 2 | 2 | 2 | 2 | 2 | 1 | 4 | 1 |
| 439486 | 3 | 1 | Chitin synthase 1 | YNL192W | 4 | 1 | 1 | 1 | 1 | 2 | 1 | 1 | 1 | 1 | 2 | 1 |
| 473500 | 4 | 2 | Cell wall related glycosyltransferase | YPL227C | 2 | 1 | 4 | 1 | 1 | 1 | 1 | 2 | 1 | 1 | 4 | 1 |
| 382141 | 7 | 1 | Dolichol-phosphate mannosyltransferase | YPR183W | 2 | 1 | 2 | 1 | 1 | 4 | 1 | 2 | 2 | 1 | 2 | 2 |
| 372414 | 4 | 2 | Cell wall related glycosyltransferase | – | 2 | 4 | 0 | 1 | 2 | 0 | 4 | 1 | 0 | 1 | 0 | 1 |
| 15443 | 1 | 1 | Chitin synthase | – | 4 | 0 | 0 | 0 | 0 | 0 | 0 | 0 | 0 | 0 | 0 | 0 |
| 234699 | 3 | 0 | Chitin synthase | – | 1 | 1 | 2 | 0 | 1 | 1 | 1 | 1 | 2 | 1 | 1 | 2 |
| 357091 | 5 | 1 | Chitin synthase | – | 1 | 1 | 1 | 2 | 2 | 2 | 1 | 1 | 2 | 1 | 4 | 1 |
| 359180 | 12 | 10 | Chitin synthase | – | 4 | 4 | 4 | 4 | 2 | 4 | 4 | 4 | 2 | 4 | 4 | 4 |
| 493091 | 8 | 2 | Chitin synthase | – | 2 | 1 | 4 | 2 | 2 | 2 | 2 | 1 | 2 | 1 | 4 | 1 |
| 374670 | 3 | 0 | Chitin synthase with myosin motor domain | – | 2 | 1 | 0 | 2 | 1 | 1 | 1 | 1 | 1 | 1 | 2 | 1 |
| 497826 | 5 | 0 | Chitin synthase with myosin motor domain | – | 2 | 1 | 2 | 2 | 1 | 2 | 1 | 1 | 2 | 1 | 1 | 1 |
| 519767 | 11 | 6 | Glycosyltransferase 2 family protein | – | 4 | 2 | 4 | 4 | 2 | 4 | 2 | 4 | 2 | 1 | 4 | 2 |

A chitin synthase regulator described as Csr2 from *C. neoformans*(Banks et al. 2005) and Skt5 from *S. cerevisiae*(S et al. 1993), is conserved in the Agaricomycetes (orthogroup represented by *C. cinerea* 538650), but did not show significant expression dynamics (Table 22).

##### 4.8.1.2. β*-*glucan biosynthesis

###### 4.8.1.2.1. β-1,6-glucan (Kre9/Knh1, GH16, GH63)

In addition to β-1,3-glucan, β-1,6-glucan is also found in fungal cell walls and is hypothesized to serve gluing purposes by covalently linking β-1,3-glucan to chitin and mannoproteins(Lesage and Bussey 2006). Its synthesis builds on UDP-glucose produced by the corresponding pathway in *S. cerevisiae*. We find that most members of the yeast beta- 1,6-glucan biosynthesis pathway have no clear 1-to-1 ortholog in the Agaricomycetes (Kre9, Knh1, Dfg5, Dcw1, Kre5, Kre1). Only Kre6 (*C. cinerea* protein ID: 447635), Rot1 (*C. cinerea* protein ID: 241938), Rot2 (*C. cinerea* protein ID: 359282) and CWH41 (*C. cinerea* protein ID: 486333) had clear orthologs. However, at the family level, all known members of the β-1,6-glucan synthesis machinery are represented in the Agaricomycetes (Table 23), except Kre1, which seems missing.

**Table 23.** Summary of developmental expression dynamics of CDE orthogroups of glucan biosynthesis related genes across 12 species. Protein ID of a representative protein is given follows by the number of species in which the orthogroup is developmentally regulated at fold change 2 and 4 (FC>2 and FC>4, respectively). Putative function and ortholog in *S. cerevisiae* (if any) are also given. Abbreviations: 0-gene absent, 1-gene present but not developmentally regulated, 2 - developmentally regulated at fold change >2, 4- developmentally regulated at fold change >4. Species names are abbreviated as: Cc – *C. cinerea*, Aa – *A. ampla*, Ab – *A. bisporus*, Ao – *A. ostoyae*, Ca – *C. aegerita*, Lb – *L. bicolor*, Le – *L. edodes*, Lt – *L. tigrinus*, Mk – *M. kentingensis*, Pc – *Ph. chrysosporium*, Po – *P. ostreatus*, Sc – *S. commune*.

| ID | Fold change |  | Putative function | S.<br>*cerevisiae*<br>ortholog | Species |  |  |  |  |  |  |  |  |  |  |  |
| --- | --- | --- | --- | --- | --- | --- | --- | --- | --- | --- | --- | --- | --- | --- | --- | --- |
|  | >2 | >4 |  |  | Cc | Aa | Ab | Ao | Ca | Lb | Le | Lt | Mk | Pc | Po | Sc |
| 417746 | 12 | 4 | Kre9/Knh1 family putative beta-1,6-glucan assembly protein | – | 4 | 2 | 2 | 4 | 2 | 2 | 2 | 2 | 4 | 2 | 4 | 2 |
| 181364 | 4 | 0 | Smi1 family protein regulating chitin deposition and cell wall assembly | – | 2 | 1 | 0 | 2 | 2 | 1 | 1 | 1 | 1 | 1 | 2 | 1 |
| 359282 | 9 | 2 | Glucosidase 2 subunit alpha | YBR229C | 2 | 1 | 4 | 4 | 2 | 2 | 2 | 2 | 2 | 1 | 2 | 1 |
| 486333 | 7 | 0 | Mannosyl-oligosaccharide glucosidase | YGL027C | 2 | 1 | 2 | 2 | 2 | 1 | 1 | 2 | 1 | 1 | 2 | 2 |
| 494484 | 8 | 6 | Glycoside hydrolase 63 family alpha-glucosidase | YMR196W | 4 | 1 | 4 | 4 | 2 | 1 | 4 | 1 | 4 | 1 | 4 | 2 |
| 241938 | 5 | 1 | Protein ROT1 involved in regulating cell wall 1,6-beta-glucan level | YMR200W | 4 | 1 | 2 | 2 | 1 | 1 | 1 | 1 | 2 | 1 | 1 | 2 |
| 543646 | 1 | 0 | 1,3-beta-glucanosyltransferase GAS1 | YMR307W | 2 | 0 | 0 | 0 | 0 | 0 | 0 | 0 | 0 | 0 | 0 | 0 |
| 447635 | 3 | 0 | Beta-glucan synthesis-associated protein KRE6 | YPR159W | 2 | 1 | 1 | 2 | 1 | 1 | 1 | 1 | 0 | 1 | 2 | 1 |
| S.c.<br>258440<br>5 | 7 | 6 | Beta-1,6-glucan synthase, GH16 family protein | – | 0 | 4 | 0 | 2 | 0 | 0 | 4 | 4 | 4 | 1 | 4 | 4 |
| 432992 | 6 | 1 | Big1 family protein putatively required for beta-1,6-glucan synthesis | – | 4 | 1 | 2 | 2 | 2 | 2 | 1 | 1 | 2 | 1 | 1 | 1 |
| 499280 | 8 | 6 | Glycoside hydrolase 31 family alpha-glucosidase | – | 4 | 0 | 1 | 4 | 0 | 4 | 2 | 4 | 4 | 2 | 4 | 0 |
| S.c.<br>254390<br>9 | 6 | 5 | Glycoside hydrolase 31 family alpha-glucosidase | – | 0 | 4 | 2 | 4 | 0 | 0 | 4 | 4 | 0 | 1 | 4 | 1 |
| 526754 | 10 | 3 | Glycoside hydrolase 9 family protein | – | 2 | 2 | 2 | 2 | 1 | 1 | 4 | 4 | 4 | 2 | 2 | 2 |
| 359649 | 5 | 4 | Kre9/Knh1 family putative beta-1,6-glucan assembly protein | – | 4 | 1 | 4 | 4 | 1 | 4 | 2 | 1 | 0 | 1 | 0 | 1 |
| 465604 | 10 | 3 | Kre9/Knh1 family putative beta-1,6-glucan assembly protein | – | 4 | 1 | 2 | 2 | 2 | 4 | 2 | 2 | 2 | 1 | 4 | 2 |
| 470186 | 12 | 10 | Kre9/Knh1 family putative beta-1,6-glucan assembly protein | – | 4 | 2 | 4 | 4 | 4 | 4 | 4 | 2 | 4 | 4 | 4 | 4 |
| S.c.<br>255705<br>9 | 9 | 9 | Kre9/Knh1 family putative beta-1,6-glucan assembly protein | – | 0 | 4 | 0 | 4 | 4 | 0 | 4 | 4 | 4 | 4 | 4 | 4 |
| 528086 | 6 | 0 | Rho family GTPase RHO1, Ras homolog gene family, member A | YPR165W | 2 | 1 | 2 | 2 | 1 | 2 | 1 | 1 | 2 | 1 | 2 | 1 |
| 446030 | 7 | 1 | 1,3-beta-D-glucan synthase | YGR032W | 2 | 1 | 2 | 4 | 1 | 2 | 2 | 1 | 2 | 1 | 2 | 1 |

**Kre9/Knh1 -** In yeasts and filamentous Ascomycota, the Kre9/Knh1 and GH16 families include the main glucan synthases(Lesage and Bussey 2006). Several additional well-characterized proteins in the endoplasmic reticulum are necessary for β-1,6-glucan synthesis *(e.g. Big1, Rot1, Rot2, Cwh41).* It should be noted that there is a strong overlap between genes participating in β-1,6-glucan synthesis and remodeling, therefore, separating these two processes is harder than it is in the case of chitin or β-1,3-glucan.

The Kre9/Knh1 family of cell surface proteins are involved in β-1,6-glucan assembly in the cell wall of fungi and have been characterized in a number of Ascomycota species(Brown and Bussey 1993; Dijkgraaf et al. 1996; S et al. 1998b; Costachel et al. 2005), predominantly yeasts. A member of this family has been independently reported as a developmentally regulated gene, DRMIP, of *L. edodes*(Szeto et al. 2007), which was reported to be primarily expressed in primordia. Reports of this family in fruiting body transcriptomes just started to emerge recently(Almási et al. 2019; Krizsán et al. 2019; Liu et al. 2020b), and no mechanistic study proved yet a link to cell wall in fruiting bodies. The family is overrepresented in Agaricomycetes(Krizsán et al. 2019), suggesting it might have experienced duplications specific to mushroom-forming fungi.

We identified 5-15 Kre9/Knh1 genes in mushroom-forming fungi. Of these 3-4 are developmentally regulated in each species. Notably, each species has 2-3 genes in this family that show a considerably higher expression in all fruiting body tissues than in vegetative mycelia, suggesting that these genes are involved in generating fruiting-body specific cell walls. For example, in *C. cinerea* we found three FB-init Kre9/Knh1 genes (*C. cinerea* protein ID: 470185, 359649, 465604), whereas in *L. edodes*, there was one FB-init gene, in addition to DRMIP, which was highly expressed and two-fold upregulated in all fruiting body tissues compared to vegetative mycelium. The family displayed more expression dynamics in more resolved transcriptomes (e.g. *A. ostoyae, C. cinerea*), suggesting that it has considerable tissue-specificity. In addition, each species showed for some genes an expression peak in vegetative mycelium, which is consistent with a partitioning of fruiting body and mycelium-specific roles. Taken together, the Kre9/Knh1 family is likely involved in β-1,6-glucan assembly in the cell wall, with several genes showing fruiting body-specific expression. We detected five CDE orthogroups of Kre9/Knh1 genes (represented by *C. cinerea* 359649, 465604, 470186, 417746, *S. commune* 2557059)(Table 23).

**GH16 -** The other major β-1,6-glucan synthase family is GH16, which includes both glucan synthases (e.g. Kre6 of *S. cerevisiae*, = *C. cinerea* 447635) and glucan-chitin cross-linking enzymes that participate in cell wall remodeling(Patel and Free 2019). We discuss this family in detail under the ‘Cell wall remodeling’ chapter.

Several ER-localized β-1,6-glucan synthesis related proteins have been described in yeasts and filamentous Ascomycota(Lesage and Bussey 2006; Orlean 2012; Essen et al. 2020), including Kre5, Big1, Rot1, Rot2 and Cwh41. Rot2 and Cwh41 are also involved in protein N-glycosylation(Lesage and Bussey 2006). These are conserved in the Agaricomycetes, mostly forming single-copy families with low expression dynamics during fruiting body development. The GT24 (glycoprotein α-glucosyltransferase) family includes the Kre5 gene in *S. cerevisiae* and *C. albicans.* Kre5 has an indirect role in β-1,6-glucan synthesis and may be involved in the refolding of misfolded N-glycoproteins in the ER, in particular β-1,6-glucan synthases(Lesage and Bussey 2006). GT24 is present as a single-copy family in Agaricomycetes (Table 23); their expression is approximately constant in fruiting bodies. Big1 is an ER membrane protein of unknown function that is required for β-1,6-glucan synthesis in *S. cerevisiae*(Lesage and Bussey 2006). It is a single-copy gene family (represented by *C. cinerea* 432992) in the Agaricomycetes, without developmentally relevant expression patterns (Table 23). Rot1 is a chaperone involved in protein folding (in general) located in the ER. It is required for cell wall synthesis and N-glycosylation and O-mannosylation in *S. cerevisiae*(Lesage and Bussey 2006). Rot1 orthologs are single copy genes (*C. cinerea* protein ID: 241938) in the Agaricomycetes that showed no developmental dynamics in the species we examined (Table 23). Rot2 is an ER-localized alpha-glucosidase involved in protein N-glycosylation, required for normal levels of β-1,6-glucan in the cell wall of *S. cerevisiae*(Lesage and Bussey 2006). It belongs to the GH31 family, which is present in 3-8 copies in Agaricomycetes. They form two CDE orthogroup (FC2/4: 8/6 species, represented by *C. cinerea* 499280 and FC2/4: 6/5 species, represented by *S. commune* 2543909)(Table 23), which were developmentally regulated in 6/5 and 5/4 species, however, expression patterns of these orthogroups did not show marked conservation. 1-to-1 orthologs of *S. cerevisiae* Rot2 (represented by *C. cinerea* 359282) showed very small gene expression dynamics.

The GH63 family includes alpha-glucosidases, such as Cwh41 of *S. cerevisiae,* involved in cell wall beta-1,6-glucan assembly in the ER. The family is represented by 2 copies in all but one of the examined species (*L. edodes* has 5 genes). GH63 genes form two conserved orthogroups, of which one (*C. cinerea* protein ID: 486333), containing orthologs of *S. cerevisiae* Cwh41) did not appear to be interesting, whereas members of the other (*C. cinerea* protein ID: 494484) showed marked expression peaks coinciding with spore formation in all species (Table 23). In *C. neoformans*, the gene showed upregulation at 72h, also coinciding with sporulation. Whether this GH63 orthogroup is needed for spore wall assembly or other processes that temporally overlap with sporulation cannot be determined based on these data, nevertheless, the family seems to be interesting in the context of fruiting body development.

###### 4.8.1.2.2. β-1,3-glucan (GT48)

In *S. cerevisiae* β-1,3-glucan is synthesized by the FKS (Fks1-3) and the GAS (Gas1-4 families as well as a few accessory proteins, such as Smi1 (*C. cinerea* protein ID: 181364) and Rho1 (*C. cinerea* protein ID: 528086)(Lesage and Bussey 2006; Verdín et al. 2019). Of the glucan synthases only FKS1 and Gas1 had clear 1-to-1 orthologs is the Agaricomycetes (*C. cinerea* protein ID: 446030 and 543646, respectively). The glucan synthase complex, which is formed by FKS1, Rho1, Smi1(Verdín et al. 2019), seems to be conserved in the Agaricomycetes.

Most Agaricomycetes species have two beta-1,3-glucan synthase genes of the GT48 family (except *M. kentingensis* and *L. tigrinus*, which have 5 and 7 respectively). Somewhat surprisingly, GT48 genes showed very modest expression dynamics and were mostly not developmentally regulated. Similarly, the Gas1 family of glucanosyltransferases is present in 1-2 copies in the examined species (*L. bicolor* has 7 and *M. kentingensis* has 4) and they were not developmentally regulated. Accessory members of the glucan synthase complex (Smi1, Rho1) showed constant expression in fruiting bodies. Collectively, these data suggest that β-1,3-glucan synthesis is present, but it is not dynamically regulated in fruiting bodies.

#### 4.8.2. Cell wall remodeling

##### 4.8.2.1. Chitin-active enzyme families (GH18, GH20, X325, AA9, AA14, CBM5/12, CBM50, CE4)

**GH18 -** Glycoside hydrolase family 18 contains chitinases (both endo- and exochitinases). Chitinases remodel the chitin structure by cleaving β-1,4 linkage of chitin or modify oligosaccharide(Verdín et al. 2019) (e.g. cleaving chitobiose from the cell wall). Chitinases are one of the most straightforward players in fruiting body development and accordingly have been reported in several previous genomic(Chen et al. 2012; Park et al. 2014), transcriptomic(Patyshakuliyeva et al. 2013; Sakamoto et al. 2017b; Xie et al. 2018; Krizsán et al. 2019) and mechanistic(Niu et al. 2016; Zhou et al. 2016c) studies. Despite the clear prediction of their role in morphogenesis, this has not been mechanistically shown in the Basidio- or Ascomycota(L et al. 2011), possibly because multiple paralogous genes in each species’ genome complicate functional studies.

In the Basidiomycota, Zhou et al reported chiB1 (*C. cinerea* protein ID: 368217) (Table 24), a class V exochitinase purified from *C. cinerea* and suggested to be involved in cap autolysis(Zhou et al. 2016c). Expression data agree with this, the corresponding gene shows a very strong induction in mature fruiting body caps. Similarly, ChiIII(Niu et al. 2016) of *C. cinerea* (C. cinerea protein ID: 354859) was reported to have both endo- and exochitinase activity, and to be expressed mostly in fruiting bodies, with peak expression in mature fruiting bodies (which is confirmed by our expression data). Most recently, Zhou et al assayed the role of chitinases in stipe cell wall extension(Zhou et al. 2019) and named six further chitinases in *C. cinerea*, ChiE1, (*C. cinerea* protein ID: 543586), ChiE2 (*C. cinerea* protein ID: 520359), ChiEn1 (*C. cinerea* protein ID: 91051), ChiEn2 (*C. cinerea* protein ID: 90984), ChiEn3 (*C. cinerea* protein ID: 470416), and ChiEn4 (*C. cinerea* protein ID: 358869). Chitinases have also been examined in the context of autolysis of the model mushrooms *C. cinerea* and *Coprinellus congregatus*(H and HT 2009).

**Table 24.** Summary of developmental expression dynamics of CDE orthogroups of chitin remodeling genes across 12 species. Protein ID of a representative protein is given follows by the number of species in which the orthogroup is developmentally regulated at fold change 2 and 4 (FC>2 and FC>4, respectively). Putative function and ortholog in *S. cerevisiae* (if any) are also given. Abbreviations: 0-gene absent, 1-gene present but not developmentally regulated, 2 - developmentally regulated at fold change >2, 4- developmentally regulated at fold change >4. Species names are abbreviated as: Cc – *C. cinerea*, Aa – *A. ampla*, Ab – *A. bisporus*, Ao – *A. ostoyae*, Ca – *C. aegerita*, Lb – *L. bicolor*, Le – *L. edodes*, Lt – *L. tigrinus*, Mk – *M. kentingensis*, Pc – *Ph. chrysosporium*, Po – *P. ostreatus*, Sc – *S. commune*.

| ID | Fold change |  | Putative function | S.<br>*cerevisiae*<br>ortholog | Species |  |  |  |  |  |  |  |  |  |  |  |
| --- | --- | --- | --- | --- | --- | --- | --- | --- | --- | --- | --- | --- | --- | --- | --- | --- |
|  | >2 | >4 |  |  | Cc | Aa | Ab | Ao | Ca | Lb | Le | Lt | Mk | Pc | Po | Sc |
| 360173 | 7 | 2 | CE4 chitin deacetylase | – | 4 | 1 | 2 | 4 | 0 | 1 | 2 | 2 | 0 | 1 | 2 | 2 |
| 368217 | 10 | 9 | Glycoside hydrolase 18, chitinase | – | 4 | 4 | 4 | 4 | 4 | 0 | 4 | 4 | 2 | 4 | 0 | 4 |
| S.c.<br>85210 | 9 | 7 | Glycoside hydrolase 18, chitinase | – | 0 | 4 | 4 | 2 | 4 | 0 | 4 | 4 | 4 | 2 | 4 | 1 |
| 543586 | 7 | 2 | Glycoside hydrolase 18, chitinase | YDR371W | 4 | 1 | 2 | 2 | 2 | 1 | 2 | 2 | 1 | 1 | 4 | 1 |
| 544873 | 6 | 4 | Polyamine transporter 2 | YGR138C | 4 | 1 | 0 | 4 | 0 | 2 | 4 | 2 | 0 | 1 | 4 | 1 |
| 354859 | 10 | 4 | Endochitinase | YLR286C | 4 | 2 | 1 | 2 | 4 | 2 | 2 | 4 | 4 | 2 | 2 | 1 |
| 355445 | 11 | 8 | CE4 chitin deacetylase 1 | YLR307W | 4 | 2 | 2 | 4 | 4 | 1 | 4 | 4 | 4 | 4 | 4 | 2 |
| 413423 | 11 | 7 | CBM5/12 domain protein | – | 4 | 2 | 4 | 4 | 4 | 2 | 2 | 4 | 4 | 1 | 4 | 2 |
| 375733 | 7 | 6 | CE4 chitin deacetylase | – | 4 | 1 | 4 | 4 | 2 | 4 | 1 | 1 | 4 | 1 | 4 | 1 |
| 377259 | 10 | 4 | CE4 chitin deacetylase | – | 4 | 4 | 4 | 1 | 4 | 2 | 0 | 2 | 2 | 2 | 2 | 2 |
| 379321 | 7 | 5 | CE4 chitin deacetylase | – | 4 | 0 | 4 | 0 | 4 | 2 | 2 | 0 | 4 | 0 | 4 | 0 |
| 42776 | 2 | 1 | CE4 chitin deacetylase | – | 4 | 0 | 0 | 0 | 2 | 0 | 0 | 0 | 0 | 0 | 0 | 0 |
| 440521 | 7 | 7 | CE4 chitin deacetylase | – | 4 | 4 | 4 | 4 | 4 | 4 | 0 | 0 | 0 | 0 | 4 | 1 |
| 464952 | 1 | 1 | CE4 chitin deacetylase | – | 4 | 0 | 0 | 0 | 0 | 0 | 0 | 0 | 0 | 0 | 0 | 0 |
| 502602 | 1 | 1 | CE4 chitin deacetylase | – | 4 | 0 | 0 | 0 | 0 | 0 | 0 | 0 | 0 | 0 | 0 | 0 |
| 358869 | 8 | 3 | Glycoside hydrolase 18, chitinase | – | 4 | 2 | 2 | 0 | 2 | 2 | 4 | 0 | 0 | 1 | 4 | 2 |
| 470416 | 9 | 6 | Glycoside hydrolase 18, chitinase | – | 4 | 2 | 4 | 4 | 2 | 4 | 0 | 2 | 4 | 0 | 4 | 1 |
| 488885 | 10 | 7 | Glycoside hydrolase 18, chitinase | – | 4 | 4 | 4 | 4 | 4 | 1 | 4 | 2 | 1 | 4 | 2 | 2 |
| 90984 | 1 | 1 | Glycoside hydrolase 18, chitinase | – | 4 | 0 | 0 | 0 | 0 | 0 | 0 | 0 | 0 | 0 | 0 | 0 |
| 91051 | 9 | 8 | Glycoside hydrolase 18, chitinase | – | 4 | 1 | 4 | 4 | 0 | 4 | 4 | 4 | 2 | 4 | 4 | 1 |
| 446427 | 8 | 6 | Glycoside hydrolase 20 family<br>putative β-1,6-N-acetylglucosaminidase | – | 4 | 1 | 4 | 4 | 4 | 0 | 4 | 2 | 0 | 2 | 4 | 1 |
| 464567 | 9 | 6 | Glycoside hydrolase 20 family<br>putative β-1,6-N-acetylglucosaminidase | – | 4 | 1 | 4 | 4 | 4 | 2 | 2 | 4 | 2 | 1 | 4 | 1 |
| 520359 | 10 | 7 | Glycoside hydrolase family 18<br>chitinase | – | 4 | 4 | 0 | 4 | 2 | 4 | 4 | 2 | 0 | 2 | 4 | 4 |
| 356467 | 7 | 4 | GlcN6P deaminase (Nag1 family)<br>involved in cell wall biogenesis | – | 4 | 1 | 4 | 0 | 4 | 0 | 2 | 4 | 2 | 2 | 1 | 1 |
| 454418 | 9 | 6 | X325 family LPMO-like protein | – | 1 | 2 | 4 | 1 | 2 | 4 | 4 | 4 | 4 | 1 | 4 | 2 |

GH18 genes are found in 8-12 copies in Agaricomycetes (up to 20 in Polyporales, see (Chen et al. 2012)), some of which harbor carbohydrate binding modules (CBMs), but their function has not been examined. Most chitinases are developmentally expressed in fruiting bodies. Mycelium-upregulated chitinases are also found in most species; these, although display peak expression in vegetative mycelia (often ∼10x that of fruiting bodies), often also show >4fold expression dynamics within fruiting bodies. We detected seven CDE orthogroups *(Table 24), which* included *C. cinerea* ChiB1, ChiE2, ChiEn1 and ChiEn3. In general, although, the detected chitinase orthogroups were developmentally regulated in many species, they did not show conserved expression patterns across species, which could indicate quick divergence of function or the failure of our reciprocal-best-hit based approach to detect functionally related genes (in the latter case phylogenetic analyses might help). One orthogroup (represented by *C. cinerea* 520359) shows gill-specific expression peaks in the species in which separate RNA-Seq data are available for gills (*A. ostoyae, P. ostreatus*) or late upregulation in *Pt. gracilis, C. aegerita, L. edodes, L. bicolor, Ph. chrysosporium, A. ampla* and *S. commune*, suggesting it may have something to do with processes that take place in the hymenium late in development, possibly sporulation.

**GH20 and chitin degradation products** - A chitin-connected GH family is GH20, which has β-1,6-N-acetylglucosaminidase activity and thus can cleave chitobiose released by chitinases further into N-acetylglucosamine (GlcNac). GlcNac may be taken up by the cell for repurposing of cleaved cell wall components as nutrient source(López-Mondéjar et al. 2009; de Oliveira et al. 2018) or, to trigger upregulation of chitinolytic genes in a transcriptional feedback-loop(Langner and Göhre 2015). The latter hypothesis seems more plausible for cases when the fungus feeds on chitin rather than modifies it for developmental purposes. In previous reports, GH20 genes have been identified in the post-harvest transcriptome of *L. edodes*(Sakamoto et al. 2017b). We identified 2-8 GH20 genes in the examined Agaricomycetes, some of which were developmentally regulated (including the single gene in *C. neoformans*), but did not form conserved CDE orthogroups. The role of this gene family in fruiting body development may be worth examining further.

If GlcNac is indeed taken up and metabolized in the cell, then two of the detected CDE orthogroups may provide clues to its mechanistic bases. An orthogroup of GlcNac transporters, which contains reciprocal best hits of *C. albicans* NGT1, was developmentally regulated in 5/4 species (*C. cinerea* protein ID: 464567, Table 24). *C. albicans* NGT1 encodes a specific transporter of GlcNac(FJ and JB 2007; Q et al. 2020). We speculate that this orthogroup may be involved in the uptake of GlcNac monomers generated by GH20 enzymes extracellularly.

In fungi, the GlcNac catabolic pathway is best described in *C. albicans*(Kumar et al. 2000; Yamada-Okabe et al. 2001), however, only a few of the corresponding genes are conserved in Agaricomycetes. Only the GlcN6P deaminase (Nag1, *C. cinerea* 356467 FC2/4: 7/4 species) and a GlcNac permease Nag4 (*C. cinerea* protein ID: 544873, FC2/4: 6/4)(Table 24) genes have clear reciprocal best hits in Agaricomycetes, whereas we could not identify clear orthologs of N-acetylglucosamine kinase (Nag5) or N-acetylglucosamine deacetylase (Nag2). Our annotations identified a different gene as N-acetylglucosamine deacetylase, however (*C. cinerea* protein ID: 446427), which was developmentally regulated in 8/6 species (FC=2/4). We also detected a GlcNac deacetylase orthogroup (*C. cinerea* protein ID: 446427), which may be involved in intracellular GlcNac catabolism.

In summary, the evidence presented here on GlcNac uptake and catabolism in mushroom forming-fungi is circumstantial, at best, however, provides clues for further research on this topic. GlcNac (and any other breakdown intermediate) produced during cell wall modification may be precious compounds in the fruiting body, which is otherwise dependent on distal supply of nutrients from the mycelium. This might have prompted the evolution of efficient repurposing mechanisms in fruiting bodies to mitigate its dependence on mycelial nutrient supply.

**CE4 chitin deacethylases -** Chitin deacetylases (CDA) are involved in chitosan production, a minor but key constituent of the cell wall. Chitosan, a more soluble polymer than chitin, occurs in both mycelia and fruiting bodies of basidiomycetes, although its role in fruiting body formation is thought to be limited(M and C 1981; Crestini et al. 1996; Baker et al. 2007). It is produced by the synergistic activities of chitin synthases and CDAs, where the former synthesize and the latter deacetylate chitin in various spatial or temporal patterns and to varying degrees(Sebastian et al. 2020). CDAs belong to the carbohydrate esterase 4 family (CE4) in the CAZy classification and are homologous to bacterial nodB factors that produce signal molecules in the rhizobia-legume symbiosis(Buhian and Bensmihen 2018). Deacetylation of chitin to chitosan makes it resistant to chitinases, which might provide a mechanism for regulating the extent of chitinase-mediated wall plasticity and thus cell expansion (e.g. in stipe elongation(Liu et al. 2021)). Fungal CDAs have a potential in biotechnology, as chitosan is an additive in food, cosmetic and pharmaceutical products(Morin-Crini et al. 2019). CDAs have poorly known roles in Ascomycota fruiting body development, with both *N. crassa* CDAs, nevertheless, being highly expressed during perithecium development, suggesting roles in sexual morphogenesis(Patel and Free 2019). It should be noted that CE4 also contains acetyl xylan esterases, which are able to break down the xylan backbone of hemicellulose during wood decay(X and S 2009). Therefore, some genes of the family in the Agaricomycetes may be related to wood decay too.

CDA genes have been detected in several RNA-Seq studies focusing on fruiting body development(Morin et al. 2012; Sipos et al. 2017b; Xie et al. 2018; Krizsán et al. 2019). Three chitin deacetylases have been experimentally characterized in *C. cinerea*, CDA1 (*C. cinerea* protein ID: 42776), CDA2 (*C. cinerea* protein ID: 502602), and CDA3 (*C. cinerea* protein ID: 464952)(Table 24), all of which were reported to have high expression in the elongating region of the stipe(Wang et al. 2018b; Bai et al. 2020). Another chitin deacetylase, primarily expressed in the fruiting bodies, was investigated in *F. velutipes*(M et al. 2008).

Searching the genomes of fruiting body forming fungi reveals 5-18 chitin deacetylase genes, several of which formed conserved single-copy orthogroups. The previously investigated CDA1-3 genes of *C. cinerea* formed species-specific orthogroups. The majority of CDA genes in the examined species were developmentally regulated in fruiting bodies, indicating key functions in fruiting body development. Six out of 17 *C. cinerea* CDA genes (Table 24) formed CDE orthogroups. These showed diverse expression peaks with no apparent trend in their expression. However, several genes were upregulated in stipes of young or mature fruiting bodies. One of them (*C. cinerea* protein ID: 355445) comprise proteins representing reciprocal best hits of *S. cerevisiae* CDA1, which is involved in the biosynthesis of the ascospore wall, required for spore wall rigidity. Expression of genes in this orthogroup did not peak during spore development, suggesting that this specific gene orthogroup is not involved in basidiospore wall synthesis. On the other hand, the expression of one and three genes were highest in young fruiting body gills and mature fruiting body gills and caps, respectively, which might be involved in producing chitosan in spore walls.

On a related note, we could not detect chitosanases, which degrade chitosan, in the genomes of mushroom-forming fungi, suggesting that chitosan is a terminal cell wall carbohydrate, not modified or cleaved further.

**Carbohydrate binding modules (CBMs) -** Carbohydrate binding modules, as their name suggests, bind various carbohydrates, and thus can frequently be attached to cell wall modifying enzymes. Chitin binding ability has been shown for at least 9 CBM families(Sánchez-Vallet et al. 2015). Of the most widely known chitin-binding CBMs (1, 5/12, 14, 18, 19, 50), only CBM1, 5/12, 18 and 50 families are conserved in the Agaricomycetes we examined. The CBM1 family, which primarily binds cellulose and according to some reports chitin, does not appear to be involved in fruiting body development: most genes that encode CBM1 containing proteins have virtually no expression in fruiting bodies. Some such proteins show significant expression dynamics (e.g. *C. cinerea* 363583, 499390), but are not conserved across species. The CBM18 family occurs in GH16 proteins, which have been described above.

**CBM5/12** - The CBM5/12 family comprises chitin-binding modules, which assist the action of cell wall modifying enzymes in fungal development(Hartl et al. 2011). The CBM5 family is sometimes considered a lectin due to its carbohydrate binding ability(Ismaya et al. 2020). Most CBM5/12 modules occur in combination with GH18 domains, where their chitin binding ability is believed to increase the efficiency of the chitinase(Hartl et al. 2011). We identified 3-6 CBM5/12 domains in the examined Agaricomycetes. All but one CBM5/12 modules are attached to GH18 chitinases and have thus been discussed above. The single CBM5/12 module that is not attached to a chitinase occurs on proteins that also harbor a domain of unknown function (DUF3421) and a DM9 repeat as well. Such proteins occur in single copy in Agaricomycetes genomes and comprise a CDE orthogroup (*C. cinerea* protein ID: 413423) that show high expression in vegetative mycelia and mature stipe tissues of most species (*A. ostoyae, P. ostreatus, M. kentingensis, C. cinerea, L. tigrinus, L. bicolor, L. edodes*) (Table 24). The protein is functionally uncharacterized. Both the DUF3421 and the DM9 repeat occur primarily in insects and other animals (but are found also in early-diverging fungi), but their activity is not known. Their developmental expression and the presence of a CBM5/12 domain suggests they may be linked to the chitin content in the cell wall, however, how exactly they function remains unclear.

**CBM50** - also known as LysM domain, this family binds chitin and can occur in a number of chitin-active proteins or on its own(GB et al. 2015). It has been reported to show developmentally dynamic expression in previous studies of fruiting body development(Sakamoto et al. 2017b; Sipos et al. 2017b; Krizsán et al. 2019; Huang et al. 2020). LysM domain proteins are widely known as effectors of plant pathogenic fungi, in which solitary secreted CBM50 modules can mask chitin residues and thus can dampen the immune response of plants(GB et al. 2015). Solitary CBM50 modules expressed in fruiting bodies could be involved in defense against insects(Kunzler 2015).

The examined Agaricomycetes contain 3-9 CBM50/LysM domain encoding genes (except *A. ostoyae,* which has 18, but its expansion may be related to the pathogenic nature of *Armillaria*(Sipos et al. 2017b)). The CBM50 module occurs almost exclusively on its own in the examined Agaricomycetes, only *C. aegerita* has proteins that harbored combinations of CBM50 and chitinase domain signatures. Interestingly, CBM50 encoding genes showed relatively little expression dynamics within fruiting bodies with 0-2 developmentally expressed genes per species. This, and reports on the induction of one of the CBM50 genes of *C. cinerea* in response to bacterial and nematode challenge(Kombrink et al. 2018; Tayyrov et al. 2018, 2019) could indirectly support a role in defense. However, some genes had high and constant expression during fruiting body development (e.g. *C. cinerea* 448942), suggesting chitin-linked functions.

**AA9 family (Lytic polysaccharide monooxygenases) -** The family has been reported only recently to have fruiting body specific expression patterns(Almási et al. 2019; Krizsán et al. 2019). AA9 proteins were initially considered to be endoglucanases and classified in glycoside hydrolase family 61, but turned out to be lytic polysaccharide monooxygenases (LPMOs) that cleave polysaccharide chains using a distinct oxidative mechanism(G et al. 2010).

The examined Agaricomycetes possess 7-34 AA9 encoding genes, of which 1-10 were developmentally regulated. A part of this copy number variation can be explained by lifestyle, as wood-decay species are known to have expanded, while biotrophic or mycorrhizal species have contracted AA9 repertoires(Miyauchi et al. 2020). AA9 LPMOs might be active on β-glucan or on both glucan and cellulose as well, providing them with the activity required to remodel the fungal cell wall(Krizsán et al. 2019). The identified AA9 genes hardly formed conserved orthogroups, which may be a consequence of high duplication/loss rates.

Several species showed interesting LPMO expression patterns, which were quite consistent within, but different between species. For example, in *C. cinerea* several genes had a late upregulation, suggesting that LPMOs might be involved in autolysis. On the other hand, in *Ph. chrysosporium* many genes showed high expression in vegetative mycelium and mature fruiting bodies but low in young fruiting bodies, creating U-shaped expression curves. In *L. tigrinus* several genes showed a marked peak in cap and gill samples of mature fruiting bodies. The single AA9 gene (Cryne_JEC21_1_562) of *C. neoformans* had a strong (45-fold) induction in 24h samples, coincident with basidium formation(Liu et al. 2018). In general, all species displayed several genes with high expression in vegetative mycelium, but negligible (<5 FPKM) expression in any of the fruiting body tissues, suggesting the existence of separate vegetative mycelium-specific and fruiting body-specific AA9 subgroups. Yet other AA9 genes displayed uniformly low (<5 FPKM) expression; these LPMO-s might be specific for the plant cell wall, and thus not induced during development. Nevertheless, the developmentally dynamic expression of several AA9 genes suggests that this family is involved in fungal cell wall remodeling during fruiting body development.

**AA14 family -** We found a conserved orthogroup (represented by *C. cinerea* 362412) that contained members of the recently circumscribed AA14 family of lytic xylan oxidases(Couturier et al. 2018). These proteins require copper as their native cofactor and were speculated to unlock xylan-coated cellulose fibers in the plant cell wall. We identified members of the AA14 family based on BlastP (query *Pycnoporus coccineus* AUM86167 and AUM86166): we find that blastp with e-value cutoff of 10^-10^ yielded copy numbers that match perfectly those reported by Couturier et al (Suppl Data Set 1)(Couturier et al. 2018). Fruiting body forming fungi contained 2-5 copies of the family, whereas yeast-like fungi (*U. maydis* and *C. neoformans*) contained none. Gene belonging to this family did not display characteristic expression peaks across species. However, as with other GH and AA families, their developmental expression in fruiting bodies suggests they may be active on some component of the fungal cell wall as well or have other extracellular roles.

**X325 family -** We also detected the recently described X325 family of copper-containing, LPMO-like proteins(Labourel et al. 2020). As a new gene family, the only functional knowledge available currently is that it displays a fold similar to fungal LPMOs, may be involved in copper homeostasis(Garcia-Santamarina et al. 2020), is upregulated in several mycorrhizal fungi and resides on the surface (via a GPI anchor) of the fungal cell wall(Labourel et al. 2020). A morphogenetic role for the family has been speculated. We here report that members of this family form a CDE orthogroup (represented by *C. cinerea* 454418) and are developmentally regulated in a wide range of species (9/6 at FC4/2), although expression patterns differ between species. Additional developmentally regulated X325 genes from other species were segregated into other orthogroups. The *L. bicolor* gene is slightly upregulated in primordia relative to free living mycelium (FC=1,8) and has a peak expression in young fruiting body stipes.

AA10 and AA11, two well-known chitinolytic LPMO families(Langner and Göhre 2015; Polonio et al. 2021), are missing in the examined Agaricomycetes, so they are unlikely to be contributors to fruiting body development.

##### 4.8.2.2. Glucan-linked families (GH3, GH5, GH16, GH17, GH30, GH55, GH71, GH128, GH152, Cdc43, WSC)

**Cdc43 -** We detected orthologs of *S. cerevisiae* Cdc43 (*C. cinerea* protein ID: 376392) as a CDE orthogroup in this study (Table 25). Cdc43, together with Ram2, forms the type I geranylgeranyl transferase of *S. cerevisiae*, which transfers a geranyl-geranyl moiety to proteins having a CaaX sequence on their C-terminal(AA et al. 1991). One of the targets in *S. cerevisiae* is Rho1p, a rho-type GTPase, which is the regulatory subunit of beta-1,3-glucan synthase(SB et al. 1999). Cdc43 orthologs were developmentally expressed in 11/6 (FC=2/4) (Table 25) species as well as *C. neoformans*. They showed a marked upregulation from vegetative mycelium to primordia in *A. ampla, A. ostoyae, C. aegerita, L. bicolor, L. edodes, C. cinerea, S. commune*, and more or less constant expression in *M. kentingensis, Pt. gracilis* and *P. ostreatus*. On the other hand, orthologs of Ram2 (*C. cinerea* protein ID: 473005) as well as those of the main substrate, Rho1 (*C. cinerea* protein ID: 528086) did not show strong expression dynamics. We hypothesize that the expression dynamics of Cdc43 mirrors the dynamics of cell wall glucan biosynthesis in fruiting bodies.

**Table 25.**
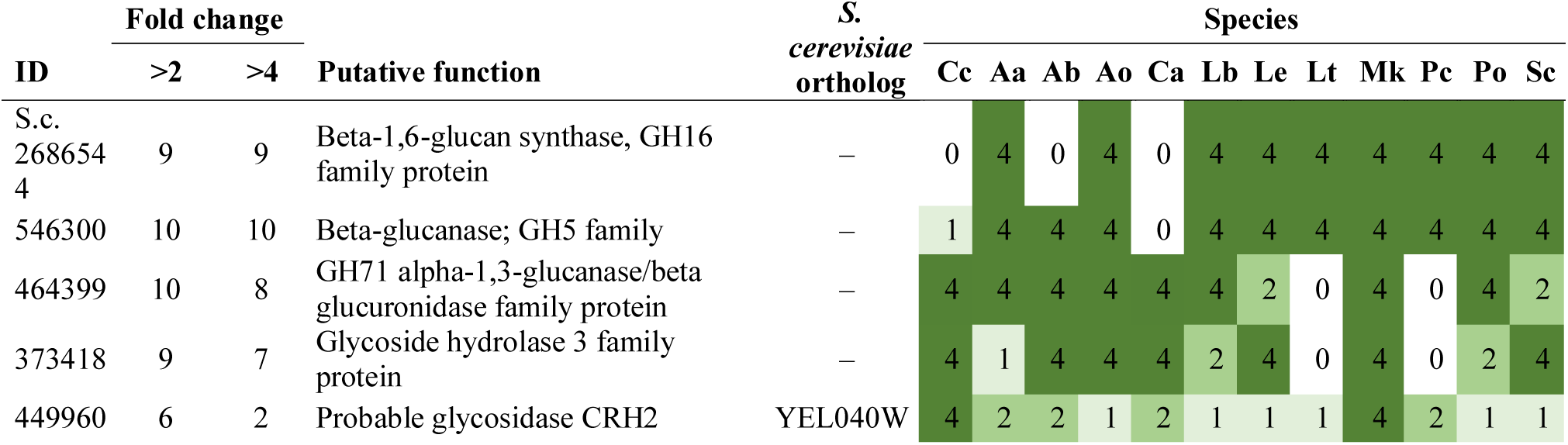
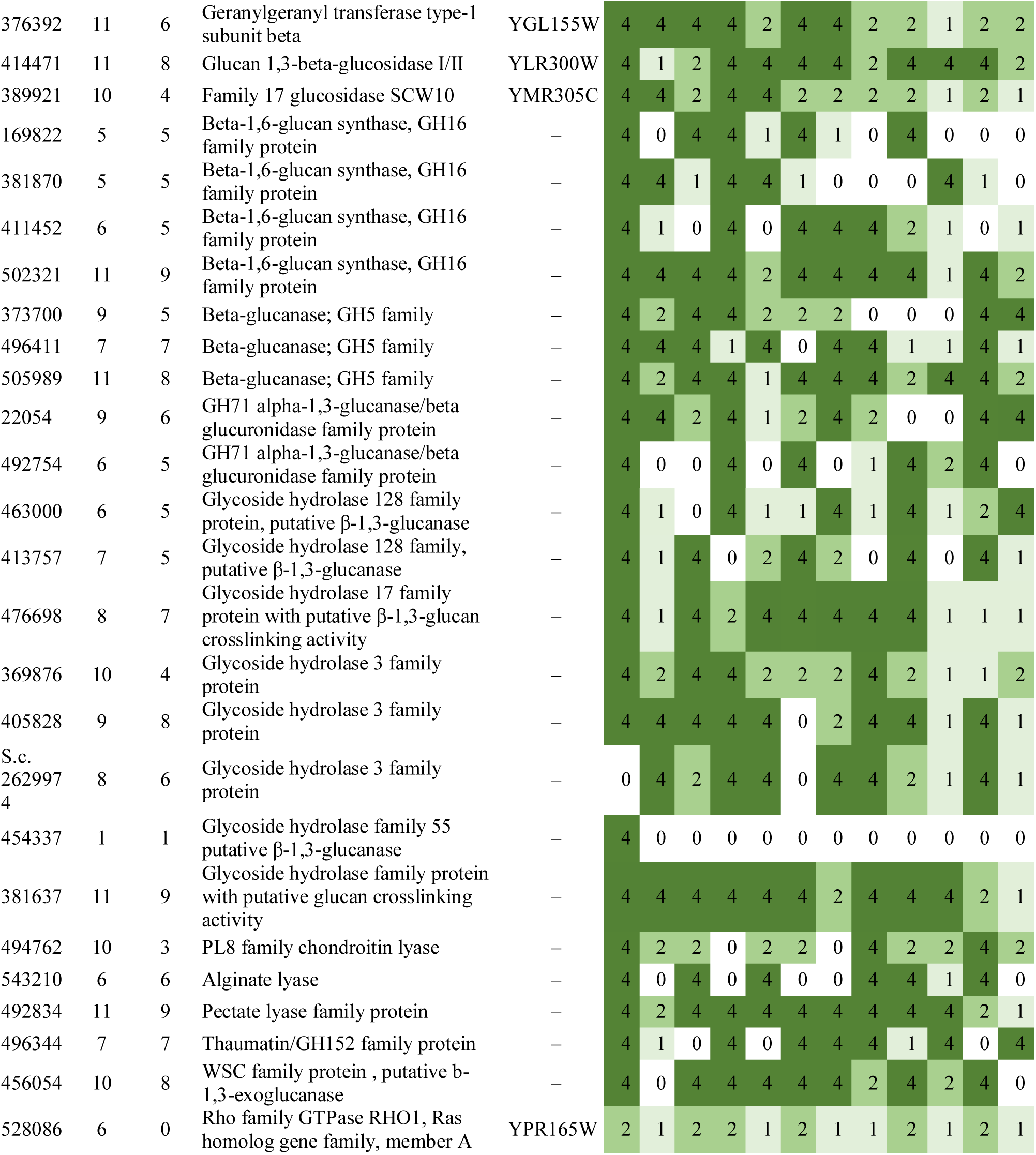
Summary of developmental expression dynamics of CDE orthogroups of glucan remodeling related genes across 12 species. Protein ID of a representative protein is given follows by the number of species in which the orthogroup is developmentally regulated at fold change 2 and 4 (FC>2 and FC>4, respectively). Putative function and ortholog in *S. cerevisiae* (if any) are also given. Abbreviations: 0-gene absent, 1-gene present but not developmentally regulated, 2 - developmentally regulated at fold change >2, 4- developmentally regulated at fold change >4. Species names are abbreviated as: Cc – *C. cinerea*, Aa – *A. ampla*, Ab – *A. bisporus*, Ao – *A. ostoyae*, Ca – *C. aegerita*, Lb – *L. bicolor*, Le – *L. edodes*, Lt – *L. tigrinus*, Mk – *M. kentingensis*, Pc – *Ph. chrysosporium*, Po – *P. ostreatus*, Sc – *S. commune*.

**GH17 -** Enzymes belonging to the GH17 family are known to crosslink β-1,3-glucan in the cell wall of yeast species and presumably *N. crassa* too(Patel and Free 2019). The family has been reported in a transcriptomic study of mushroom development(Krizsán et al. 2019) (as endo-β-1,3-glucanosyltransferase), and it was suggested to be involved in β-1,3-glucan crosslinking, although direct evidence for this is lacking at the moment. The examined Agaricomycetes possess 1-4 GH17 encoding genes. *C. cinerea* has four GH17 genes, of which only one appear to be clear orthologs of Ascomycota genes: *C. cinerea* 389921 is reciprocal best hit to SCW10 of *S. cerevisiae* a soluble cell wall protein with a poorly known function. GH17-s formed 2 CDE orthogroups (Table 25). One of the CDE orthogroups (represented by *C. cinerea* 476698) showed clear tissue-specific expression (cap- or stipe upregulation), but interestingly, this was not fully consistent across species.

**GH16 -** The GH16 family shows glycosylhydrolase and glycosyltransferase activity and contain glucan synthase and crosslinking enzymes, which are active in cell wall synthesis (e.g. Kre6 and Skn1 of *S. cerevisiae*) and remodeling. GH16s were shown to crosslink β-1,3- glucan and β-1,6-glucan to chitin fibers in *Candida* spp and *N. crassa*(Patel and Free 2019). Chitin-glucan crosslinking is key to morphogenesis in fungi(Arroyo et al. 2016). GH16s comprise the CRH (Congo Red Hypersensitive) family, three members of which (Crh1p, Crh2p and Crr1p) were originally circumscribed in *S. cerevisiae.* Of these, only Crr1 has a clear 1-to-1 ortholog in Agaricomycetes (*C. cinerea* protein ID: 449960), the other two do not. At the transcriptome level, the upregulation of GH16 genes have been reported before in a comparative study(Krizsán et al. 2019), in *A. bisporus*(Morin et al. 2012; Patyshakuliyeva et al. 2013)*, Leucocalocybe mongolica*(Duan et al. 2021)*, Pleurotus spp*(Xie et al. 2018; Ye et al. 2021) as well as during post-harvest development on *L. edodes*(Sakamoto et al. 2009, 2017b). Mlg1 is a functionally characterized member of the GH16 family in *L. edodes* assayed for its glucanase activity during postharvest senescence(Sakamoto et al. 2009).

The species in this study have 21-36 GH16 encoding genes, which is consistent with a previous report on *L. edodes*(Sakamoto et al. 2017b). About half of the GH16 genes in each species also carry a ‘Beta-glucan synthesis-associated Skn1’ domain, which signifies relationship with the Kre6/Skn1 family, which is involved in β-1,6-glucan synthesis(Garcia-Rubio et al. 2020). We found five GH16 CDE orthogroups (*C. cinerea* protein ID: 502321, 381870, 411452, 169822, *S. commune* 2686544) (Table 25), that are not orthologous to functionally characterized GH16 genes in the Ascomycota. Although most genes do not form conserved orthogroups, possibly because of volatility (frequent duplication/loss), it is noteworthy that each species has 1-3 genes which show a marked upregulation in the primordium stage and remain highly expressed in the fruiting body relative to vegetative mycelium. These might be involved in the formation of 3-dimensional fungal tissues. Another interesting observation is that eight out of 21 GH16 genes in *C. neoformans* are also developmentally regulated, with several of them showing peak expression at 24h, coincident with basidium development.

**GH30 -** Glycoside hydrolase family 30 (GH30) includes enzymes with diverse activities, which have been mostly discussed for their role as endo-xylanases in wood decay in the fungal literature(Wymelenberg et al. 2009; Katsimpouras et al. 2019). However, GH30 also includes β-1,6-glucanases, which were found to be upregulated in *L. bicolor* ectomycorrhizae(C et al. 2014), raising the possibility of morphogenetic roles. GH30 members were also upregulated during postharvest development of *L. edodes*(Sakamoto et al. 2017b). Of the 1-3 (6 in *L. bicolor*) GH30 genes in the examined Agaricomycetes, at most 1-2 are developmentally expressed in fruiting bodies and only a single orthogroup (represented by *C. cinerea* 381637) was so in several species (Table 25). Genes in this orthogroup were most often upregulated in stipe tissues relative to pileus in fruiting bodies. GH30 genes of *C. cinerea* have been proposed to facilitate stipe elongation by in the stipe base may contribute to replacing β-1,6-glucan to β-1,3-glucan crosslinking in the stipe base, which may reduce wall extensibility, leading to the cessation of growth(Liu et al. 2021). Further, the shared upregulation of GH30 genes in multiple fruiting body transcriptomes and in ectomycorrhizae provide strong evidence for its role in FCW remodeling during morphogenesis in the Basidiomycota.

**GH3 -** The GH3 family comprises enzymes with diverse activities, including exo-β-1,3-glucanase activity, and has been suggested to participate in cell wall remodeling in Ascomycetes(Patel and Free 2019). The upregulation of GH3 genes in fruiting bodies has previously been noted in a number of species(Zhou et al. 2018; Krizsán et al. 2019), raising the possibility that they participate in cell wall remodeling during fruiting body development. A GH3 family member, that was upregulated in fruiting bodies of *Auricularia heimuer*, was recently cloned and characterized(Sun et al. 2020). GH3 encoding genes are present in 7-20 copies in examined Agaricomycetes. Three CDE orthogroups belonging to the GH3 family (*C. cinerea* protein ID: 369876, 373418, *S. commune* Schco3_2629974,) were detected (Table 25), genes in these orthogroups showed diverse expression patterns in the fruiting bodies of different species. It should be mentioned that some GH3 enzymes, mostly in bacteria, possess β-N-acetylhexosaminidase activity, which allows the encoded enzymes to be involved in chitin utilization, by cleaving N-acetylglucosamine dimers into monomeric GlcNac(de Oliveira et al. 2018). Most such enzymes are bacterial, with sporadic reports of β-N-acetylhexosaminidase activity in fungi(S et al. 2014). Therefore, whether GH3 upregulation is linked to chitin utilization or glucan remodeling cannot be determined here.

**GH5 -** The GH5 family is a very diverse group that includes β-1,4-glucanases, as well as several other activities. It includes classic cellulases involved in wood decay(Floudas et al. 2012), whereas Ascomycota GH5 genes, such as Exg1, Exg2 and Exg3 of *S. pombe*, or Exg1 of *S. cerevisiae*, are well-known components of the cell wall glucan remodeling machinery(Dueñas-Santero et al. 2010). GH5s were shown to not only hydrolyze β-1,3-glucan, but also to act as transglycosylases, and thus contribute to cell wall crosslinking(Arroyo et al. 2016). Based on expression patterns, GH5s were speculated to be active on beta-glucan in fruiting body cell walls also(Y et al. 2005; C et al. 2014; Almási et al. 2019; Krizsán et al. 2019). Krizsan et al reported multiple subfamilies (GH5_7, GH5_9, GH5_15, GH5_49) with endo-β-1,4-mannanase, endo-β-1,6- and exo-β-1,3-glucanase activities to be upregulated in fruiting bodies(Krizsán et al. 2019). The family includes the exo-beta-1,3-glucanase Exg1 from *L. edodes*, which was reported to be fruiting body specific(Y et al. 2005). Our re-analysis confirms that Exg1 (Lenedo1_1176007) has consistently higher expression in stipe than in cap tissues.

GH5 is a large family with 16-20(25) copies in the examined Agaricomycetes. Consistent with their diversity, we detected six CDE orthogroups in our data (*C. cinerea* protein ID: 546300, 405828, 505989, 496411, 414471, 373700) (Table 25). One of these (*C. cinerea* protein ID: 546300), contains reciprocal best hits of *Sch. pombe* Exg3, a cytoplasmic β-1,6-glucanase with unknown cellular function. All species in which this orthogroup is developmentally regulated showed upregulation in mature fruiting bodies. This could indicate a link to sporulation, or other late developmental events. Another orthogroup (*C. cinerea* protein ID: 414471) contains 1-to-1 orthologs of *Sch. pombe* and *S. cerevisiae* Exg1, which is involved in cell expansion and sporulation, picturing potentially similar roles in Agaricomycetes too. Members of yet another orthogroup (*C. cinerea* protein ID: 373700), are consistently higher expressed in all fruiting stages (primordia to mature FB) than in vegetative mycelia in *A. ostoyae, C. aegerita, A. ampla, C. cinerea, P. ostreatus, S. commune* and to some extent in *L. bicolor* and *L. edodes* (but not in *A. bisporus* and *Pt. gracilis*).

**GH55 -** This family contains exo- or endo-β-1,3-glucanases that have been characterized functionally in fruiting bodies of two Agaricomycetes. Exg2 was found to be expressed higher in the stipe than in the cap of *L. edodes*(Sakamoto et al. 2005). The GH55 family has also been reported to be upregulated during cap expansion of *Volvariella volvacea* and the corresponding gene was named Exg2(Tao et al. 2013). We detected on average two GH55 genes in the examined Agaricomycetes (5 in *A. ostoyae*). One of these formed a CDE orthogroup (represented by *C. cinerea* 492834), which included *L. edodes* Exg2 and showed very strong induction in young fruiting body caps and stipes of *C. cinerea*, caps of *A. ostoyae* and stipes of *M. kentingensis*. Overall, these data are consistent with the suggested role of GH55 in cap expansion in *V. volvacea*. In *L. edodes*, the expression of Exg2 was higher in stipes than in cap tissues. Interestingly, the second GH55 gene of *C. cinerea* (*C. cinerea* protein ID: 454337) (Table 25) also shows this characteristic induction and high expression in young fruiting body caps and stipes. In species with simpler morphology (*S. commune, A. ampla*) the family did not show characteristic expression dynamics. A third GH55 gene (Armosto1_267056) in *A. ostoyae* showed an induction in primordia relative to vegetative mycelium.

**GH128 -** GH128 has been first circumscribed in the context of fruiting body development, based on a fruiting body expressed β-1,3-glucanase, Glu1, in *L. edodes*(Y et al. 2011). Since its discovery, some reports have mentioned upregulation of GH128 genes in fruiting bodies(Krizsán et al. 2019; Duan et al. 2021; Merényi et al. 2021), with a single report also in connection with wood decay(Mäkinen et al. 2019). On average, the GH128 family has 2-8 members in Agaricomycetes. We detected two CDE orthogroups (represented by *C. cinerea* 463000, 413757) which were expressed in the majority of species (Table 25) and several GH128 genes which were developmentally expressed in a species-specific way. In *C. cinerea*, protein 413757 represents the best hit of *L. edodes* GLU1. Available data here and in previous publications suggest that GH128 might be a primarily fruiting body-related gene family, worthy of further functional examination.

**GH71 -** Four orthogroups containing developmentally regulated GH71 alpha-1,3-glucanases and beta-glucuronidases were detected (*C. cinerea* protein ID: 22054, 464399, 456054, 492754)(Table 25). GH71 genes (also called mutanases) are involved in fungal cell wall remodeling in a wide range of species, both yeasts and filamentous fungi(Dekker et al. 2004; C et al. 2014; H et al. 2015). A well-characterized member of this family is the *Sch. pombe* endo-1,3-alpha-glucanase Agn1, which facilitates cell fission via the degradation of the septum material(Dekker et al. 2004). In previous studies of mushroom development, this family was reported only by Krizsan et al(Krizsán et al. 2019), Park et al(Park et al. 2014), and Morin et al(Morin et al. 2012). In *Pleurotus eryngii*, GH71 genes were found downregulated in primordia after blue light stimuli(Xie et al. 2018). GH71 genes are present in 3-12 copies in the examined Agaricomycetes. One of the CDE orthogroups (represented by *C. cinerea* 456054) is upregulated in primordia relative to vegetative mycelium in 7 of the 10 species (*C. cinerea, A. ostoyae, C. aegerita, L. tigrinus, L. bicolor, Ph. chrysosporium, P. ostreatus*) in which it is present.

**GH152 -** The GH152 family, also known as thaumatin-like proteins, which include both putatively defense-related antimicrobial peptides and β-1,3-glucanases. They show homology to plant thaumatin-like proteins. Because the literature reports both antimicrobial and glucanase activity for this family(Kunzler 2015), we do not attempt to subdivide them and refer to them as GH152/thaumatin proteins. The GH152/thaumatins include an experimentally characterized gene, Tlg1 in *L. edodes* (Lenedo1_561169), which was detected in a post-harvest screen of cell wall and lentinan degrading genes(Sakamoto et al. 2006). Lentinan is a glucan-like cell wall component with important medicinal properties in *L. edodes*. The expression of the gene was detected only after harvest, raising questions as to what the physiological role of the gene might be under natural circumstances. Our analysis of the data from Zhang et al did not reveal developmentally dynamic expression for Tlg1 using RNA-Seq data(Zhang et al. 2021c). Structure prediction for an *A. ostoyae* developmentally expressed protein indicated an acidic cleft in the 3D structure, characteristic of antimicrobial members of the family(Krizsán et al. 2019). A new report of antifungal activity of the thaumatin-like protein Le.TLP1 in *L. edodes* further supports putative defense roles for the GH152 family(Ma et al. 2021).

The family comprises 2-8 (19 in *Pt. gracilis*) copies in examined Agaricomycetes, many of which are poorly conserved. Therefore, while several genes are developmentally expressed in each species, only a single CDE orthogroup was detected (represented by *C. cinerea* 496344)(Table 25). The role of GH152/thaumatins in fruiting body development is unclear at the moment, it could be involved in defense against microbes, cell wall glucan remodeling, or both(Kunzler 2015).

Additional beta-glucan active families (GH1, GH6, GH12) were detected in the broader orthology-based approach of Krizsan et al(Krizsán et al. 2019), which were not detected in this study, probably because of the strict orthology definition we use.

**WSC family** - this family of lectin-like proteins comprises a widely conserved but poorly known group of fungal proteins. The family was originally described as a putative stress receptor in *S. cerevisiae*(AL et al. 1999) and as a domain in an extracellular β-1,3-exoglucanase in *Trichoderma harzianum*(Cohen-Kupiec et al. 1999), and later demonstrated to be a carbohydrate binding module, similar to CBMs(S et al. 2019). The best-characterized member in the Basidiomycota seems to be involved in beta-glucan binding/FCW remodeling during plant-fungal interaction in *Serendipita indica*(Wawra et al. 2019). Whether this is true for other WSC family members too, is not known, but is indicative of a potential role in adhesion/remodeling at the cell surface.

The examined Agaricomycetes harbored 2-12 copies of WSC-domain encoding genes. Of these, several showed strong induction in primordia and fruiting bodies in *A. ostoyae, C. aegerita, C. cinerea, L. bicolor, L. edodes, L. tigrinus, Ph. chrysosporium, P. ostreatus* and *S. commune* (but not in *A. ampla* and *M. kentingensis*). Genes with a predicted WSC domain in their encoded proteins were typically primordium-upregulated in all species. In *C. cinerea* these genes had high expression in early (proliferative) development (hyphal knots to stage 2 primordia), followed by very low expression in most genes in late developmental stages. The WSC domain can occur on its own or in combination with other enzymes (most often GH71 and glyoxal oxidase) and uncharacterized domains, indicating that it often fulfills a binding role in larger proteins. For example, WSC domain is present on GH71 proteins that form a 10/8 orthogroup (represented by *C. cinerea* 464399), which we discussed above.

##### 4.8.2.3. Miscellaneous cell wall remodeling (GH27, GH72, GH76, GH79, GH88, GT18, pectinesterases)

In addition to chitin- and glucan-active families, several further CAZyme groups showed widespread developmental regulation in fruiting bodies, for which the substrates in fruiting bodies are less clear. These make very interesting groups indicating potentially novel cell wall components, however, they are hard to place in context of fruiting body development as of now.

**GH79 and 88 -** An orthogroup containing GH79 β-glucuronidases (*C. cinerea* protein ID: 470689) (Table 26), showed developmental expression in several species, consistent with reports of Krizsan et al(Krizsán et al. 2019). Low amounts of glucuronic acid was reported in *N. crassa* cell walls(Verdín et al. 2019), but the role of GH79 enzymes in cell wall biosynthesis or remodeling are unknown. On a related note, an orthogroup of GH88 glucuronyl hydrolases was detected (*C. cinerea* protein ID: 384105)(Table 26). GH88 genes are generally linked to pectin degradation by wood-decay fungi(van den Brink and de Vries 2011); their potential role in fruiting bodies is not known.

**Table 26.**
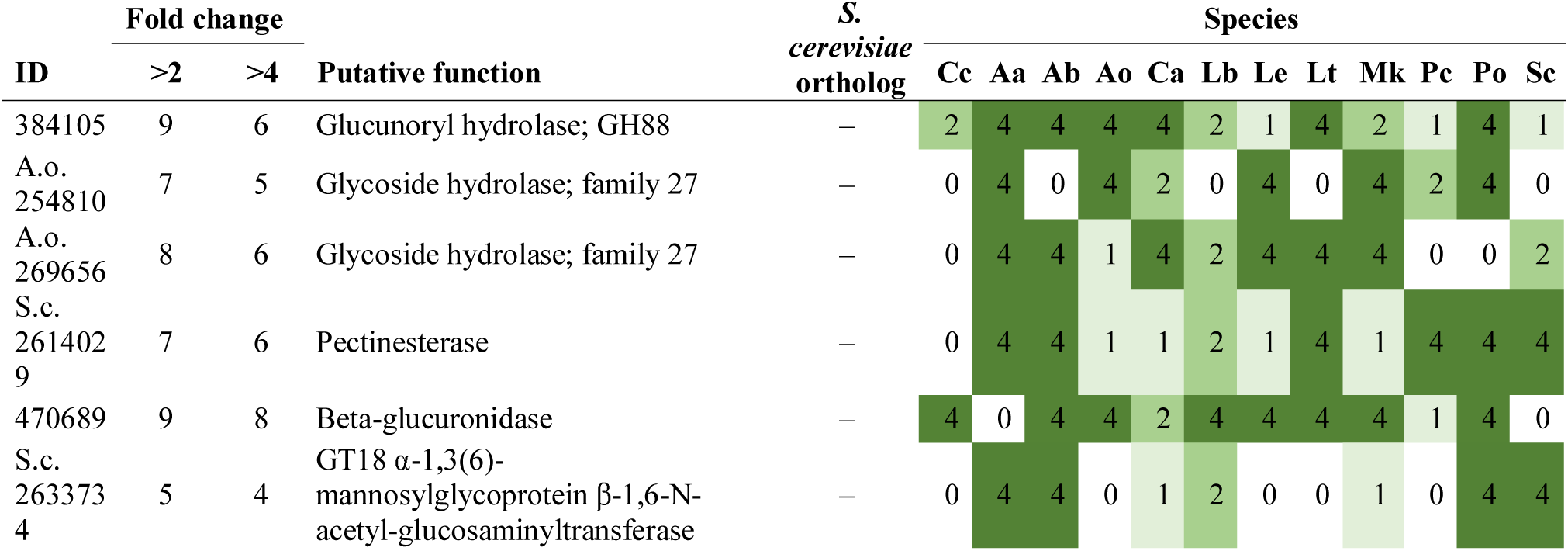
Summary of developmental expression dynamics of CDE orthogroups of miscellaneous cell wall remodeling genes across 12 species. Protein ID of a representative protein is given follows by the number of species in which the orthogroup is developmentally regulated at fold change 2 and 4 (FC>2 and FC>4, respectively). Putative function and ortholog in *S. cerevisiae* (if any) are also given. Abbreviations: 0-gene absent, 1-gene present but not developmentally regulated, 2 - developmentally regulated at fold change >2, 4- developmentally regulated at fold change >4. Species names are abbreviated as: Cc – *C. cinerea*, Aa – *A. ampla*, Ab – *A. bisporus*, Ao – *A. ostoyae*, Ca – *C. aegerita*, Lb – *L. bicolor*, Le – *L. edodes*, Lt – *L. tigrinus*, Mk – *M. kentingensis*, Pc – *Ph. chrysosporium*, Po – *P. ostreatus*, Sc – *S. commune*.

**GH27 -** Similarly, two CDE orthogroups (represented by Armosto1_254810 and Armosto1_269656) of GH27 enzymes with alpha-galactosidase or α-*N*-acetylgalactosaminidase activity were detected (Table 26). This family is discussed mostly in the context of lignocellulose degradation(Kabel et al. 2017), to our knowledge it has not been reported in fruiting body transcriptomes before. As a tendency, GH27 encoding genes were often upregulated late in the development of the containing species. Their role in fruiting body formation is not known.

**GT18 -** The GT18 family, which contains α-1,3(6)-mannosylglycoprotein β-1,6-N-acetylglucosaminyltransferases showed a patchy distribution in the examined species, being completely missing from the genomes of *C. cinerea* and *A. ostoyae*. However, it showed a strong upregulation in primordia relative to vegetative mycelium and constant high expression in fruiting bodies of *A. bisporus, A. ampla, S. commune, C. aegerita* and *P. ostreatus* (not in *M. kentingensis and L. bicolor*)(Table 26, *S. commune* protein ID: Schco3_2633734). This indicates fruiting body specific functions for this family, possibly in chitin synthesis, although its exact role remains unknown. To our best knowledge, the role of the GT18 family in modifying the fungal cell wall has not been described in any species.

**GH72 -** Contrary to our expectations, members of the GH72 family did not appear to be developmentally relevant in Agaricomycetes. This family contains cell wall transglycosylases that crosslink glycoproteins to cell wall components in the Ascomycota(Kar et al. 2019), and therefore we expected it to be developmentally expressed in Basidiomycota as well. Their expression is considerable but largely constant in Agaricomycetes fruiting bodies. This could indicate a difference between Asco- and Basidiomycota cell wall composition, or that they act during fruiting body development irrespective of their constant expression.

**GH76 -** In the Ascomycota, GH76 endo-1,6-alpha-mannosidases (Dfg5 and Dcw1 in *S. cerevisiae*) are required for normal cell wall synthesis. Their exact role may not be known with certainty, but they were hypothesized to be involved in the proper positioning of glycosylphosphatidylinositol-anchored cell wall proteins(Orlean 2012). To our best knowledge, the GH76 family has not yet been suggested to participate in fruiting body development in the Agaricomycetes. It is present 0-10 copies in the Agaricomycetes and shows predominantly mycelium-specific expression. We only found scattered expression dynamics in fruiting bodies, such as a peak in stipes of *P. ostreatus* (PleosPC15_2_1064904), but most genes are not developmentally regulated. Given that the family is missing in some species completely and the predominantly vegetative mycelium-specific expression, we hypothesize that the GH76 family does not participate, or only to a small extent, in cell wall assembly in Agaricomycete fruiting bodies.

**Pectinesterases -** We detected a CDE orthogroup (represented by Schco3_2614029) that contains genes generally linked to pectin degradation. It contains pectinesterases that are developmentally regulated in 6/5 species (FC 2/4) (Table 26). Because pectin is not a known cell wall component in Agaricomycetes, speculating about the function of these genes is hard and may require more research.

##### 4.8.2.4. Multicopper oxidases and cupredoxins

Laccases catalyze the oxidation of diverse phenolic and non-phenolic compounds, which makes them suited to diverse (extra)cellular roles, from lignin degradation, pigment production to dye decolorization or fruiting body related functions(P 2006). Their biochemical properties provide laccases with significant potential in industrial applications. Despite very intense research on laccases (mostly in the context of lignocellulose decomposition), we still have very limited information on the precise role of laccases in fruiting body development.

Several reports of upregulation of laccase, or multicopper oxidase (MCO, the broader family in which laccases belong) genes or laccase activity in fruiting bodies of Agaricomycetes(Zhao and Kwan 1999; Ohga et al. 2000; Chen et al. 2003; Wang and Ng 2005; Madhavan et al. 2014; Sakamoto et al. 2015; Sipos et al. 2017b; Nagy et al. 2018) are available (reviewed in detail by(Kües et al. 2011)). *Schizophyllum commune* was reported to have 6 MCO genes (two of which are laccases), of which 3 were reported to be upregulated in fruiting bodies(Madhavan et al. 2014) (Mco2, Mco3, Lcc1) by RT-PCR, which is consistent with later RNA-Seq data(Krizsán et al. 2019). Of these, Lcc1 showed the highest similarity to Fet3-type ferroxidases. Similarly, *P. ostreatus* laccases were reported to be expressed in a developmental stage-specific manner, with Lacc5 and Lacc12 specifically expressed in fruiting bodies and primordia, respectively, while other laccases had even or mycelium- biased expression patterns(Park et al. 2015; Jiao et al. 2018). Stage-specific MCO expression was reported in *L. bicolor* with specific laccase genes induced in fruiting bodies (Lcc2), ectomycorrhizae and free-living mycelium(Courty et al. 2009). These studies have linked laccase expression to two major functions, pigment production in either spores or fruiting bodies and fungal cell wall modification or cell-cell adhesion(Kües et al. 2011). Laccases were speculated to polymerize polyphenols in the intercellular spaces and thus contribute to the binding of neighboring hyphae to each other. The hypothesis of laccases being related to adhesion comes from Leatham and Stahmann(Leatham and Stahmann 1981), but, to our knowledge, was never experimentally verified. Although not relevant to fruiting body formation, an interesting inverse relationship between mycelial laccase expression and flushes of fruiting body development was reported in several species, possibly related to stopping mycelium growth or nutrient acquisition when nutritional competence for fruiting is achieved (reviewed in(Kües et al. 2011)).

Developmental studies indicated laccase function in fruiting body formation in *Pleurotus spp*(Das et al. 1997). and *L. edodes*(Nakade et al. 2011). In *P. florida* laccase gene silencing abolished fruiting body development and caused slower vegetative growth, whereas in *L. edodes* aerial hyphae formation was also reduced and hyphae developed thinner, non-layered cell walls upon the silencing of the Lcc1 laccase gene. Zhang et al reported faster vegetative growth and earlier fruiting in laccase (Lcc1) overexpressing strains of *H. marmoreus* compared to wild type strains(Zhang et al. 2015a), although no clear hypothesis on how overexpression of a single structural gene could result in these phenotypes was presented.

We here present annotations for multicopper oxidases, which include laccases, ascorbate oxidases and ferroxidases(Kües et al. 2011). The 11 genomes of Agaricomycetes examined here had 5-26 multicopper oxidases, a considerable proportion of which were developmentally regulated, for example, 13 out of 17 in *C. cinerea* (Table 27). We found three CDE laccase orthogroups; these comprised *C. cinerea* Lcc8, Lcc2 and their orthologs as well as reciprocal best hits of the *S. cerevisiae* ferroxidase Fet3. [Of note, we detected reciprocal best hits of the ferroxidase Fet3 in *C. cinerea*, whereas a previous report, based on phylogenetic analyses did not identify Fet3-type MCOs in this species(Kües et al. 2011); we note that this may arise from the limitations of our reciprocal best hit based search and that accurate classification of MCOs requires the examination of specific catalytic residues. Orthologs of *C. cinerea* Lcc2 (*C. cinerea* protein ID: 368135) are also involved in morphogenesis in *L. edodes (as Lcc1)*, and differentially expressed during *Cryptococcus* basidium formation(Liu et al. 2018; Merényi et al. 2021). We found that Lac4 from *Volvariella volvacea*, which was implicated in fruiting body development(Chen et al. 2003) showed highest similarity to the CDE orthogroup that comprised *L. edodes* Lcc1 (represented by *C. cinerea* 368135). In the examined Agaricomycetes, this group of genes is upregulated in caps (*P. ostreatus, C. cinerea*), stipes (*M. kentingensis*) or primordia (*A. ampla*), or shows flat expression curves (other species). The detected ferroxidase orthogroup was present in 7 species, of which it was developmentally regulated in *A. bisporus, A. ostoyae, C. cinerea, L. edodes* and *P. ostreatus* (FC>2). A Fet3-like gene was also upregulated in *S. commune*(Madhavan et al. 2014), but this gene did not group into the detected orthogroup in our analysis. Ferroxidases are membrane proteins that catalyze the oxidation of Fe^2+^ to Fe^3+^ for iron uptake and have only weak laccase activities(Larrondo et al. 2003). Finally, *C. cinerea* 440170 is the orthologs of *C. neoformans* Lac1, which has been shown to be involved in melanin biosynthesis(Lee et al. 2019). This gene shows a strong expression peak in young fruiting body gills, suggesting that it is involved in producing spore pigments in *C. cinerea*.

**Table 27.** Summary of developmental expression dynamics of CDE orthogroups of multicopper oxidase genes across 12 species. Protein ID of a representative protein is given follows by the number of species in which the orthogroup is developmentally regulated at fold change 2 and 4 (FC>2 and FC>4, respectively). Putative function and ortholog in *S. cerevisiae* (if any) are also given. Abbreviations: 0-gene absent, 1-gene present but not developmentally regulated, 2 - developmentally regulated at fold change >2, 4- developmentally regulated at fold change >4. Species names are abbreviated as: Cc – *C. cinerea*, Aa – *A. ampla*, Ab – *A. bisporus*, Ao – *A. ostoyae*, Ca – *C. aegerita*, Lb – *L. bicolor*, Le – *L. edodes*, Lt – *L. tigrinus*, Mk – *M. kentingensis*, Pc – *Ph. chrysosporium*, Po – *P. ostreatus*, Sc – *S. commune*.

| ID | Fold change |  | Putative function | S.<br>*cerevisiae*<br>ortholog | Species |  |  |  |  |  |  |  |  |  |  |  |
| --- | --- | --- | --- | --- | --- | --- | --- | --- | --- | --- | --- | --- | --- | --- | --- | --- |
|  | >2 | >4 |  |  | Cc | Aa | Ab | Ao | Ca | Lb | Le | Lt | Mk | Pc | Po | Sc |
| 408308 | 0 | 0 | *C. cinerea* laccase lcc1 | – | 1 | 0 | 0 | 0 | 0 | 0 | 0 | 0 | 0 | 0 | 0 | 0 |
| 378517 | 0 | 0 | *C. cinerea* laccase lcc15 | – | 1 | 0 | 0 | 0 | 0 | 0 | 0 | 0 | 0 | 0 | 0 | 0 |
| 443425 | 1 | 1 | *C. cinerea* laccase lcc5 | – | 4 | 0 | 0 | 0 | 0 | 0 | 0 | 0 | 0 | 0 | 0 | 0 |
| 467038 | 0 | 0 | *C. cinerea* laccase lcc6 | – | 1 | 0 | 0 | 0 | 0 | 0 | 0 | 0 | 0 | 0 | 0 | 0 |
| 89107 | 0 | 0 | *C. cinerea* laccase lcc7 | – | 1 | 0 | 0 | 0 | 0 | 0 | 0 | 0 | 0 | 0 | 0 | 0 |
| 368135 | 10 | 4 | *C. cinerea* laccase lcc15 | – | 4 | 4 | 4 | 2 | 2 | 2 | 0 | 2 | 2 | 0 | 4 | 2 |
| 353997 | 9 | 4 | Cupredoxin | – | 4 | 1 | 4 | 4 | 2 | 2 | 1 | 2 | 4 | 1 | 2 | 2 |
| 501548 | 5 | 3 | Iron transport multicopper oxidase<br>FET3 | YMR058W | 4 | 0 | 4 | 4 | 1 | 0 | 2 | 0 | 0 | 1 | 2 | 0 |
| 266892 | 2 | 1 | *C. cinerea* laccase lcc3 | – | 2 | 0 | 0 | 0 | 0 | 0 | 4 | 0 | 0 | 0 | 0 | 0 |
| 440170 | 2 | 2 | *C. cinerea* laccase lcc8 | – | 4 | 0 | 4 | 0 | 0 | 0 | 0 | 0 | 0 | 0 | 0 | 0 |
| A.o.<br>257754 | 7 | 7 | Cupredoxin | – | 0 | 0 | 1 | 4 | 4 | 1 | 4 | 4 | 4 | 4 | 4 | 0 |
| 356026 | 9 | 7 | Cupredoxin | – | 4 | 1 | 0 | 4 | 4 | 4 | 4 | 2 | 4 | 2 | 4 | 1 |
| 409847 | 5 | 3 | Cupredoxin | – | 4 | 4 | 0 | 0 | 2 | 4 | 0 | 0 | 0 | 0 | 1 | 2 |
| 441442 | 9 | 4 | Cupredoxin | – | 4 | 1 | 1 | 4 | 0 | 2 | 4 | 2 | 4 | 2 | 2 | 2 |
| 451850 | 6 | 5 | Cupredoxin | – | 4 | 2 | 4 | 4 | 1 | 0 | 0 | 0 | 4 | 1 | 4 | 0 |
| 497434 | 6 | 5 | Cupredoxin | – | 4 | 0 | 0 | 1 | 4 | 4 | 4 | 0 | 4 | 0 | 2 | 0 |

In summary, experimental and gene expression evidence on the role of multicopper oxidases in fruiting body development is abundant and -omics data reveal marked developmental stage-specific expression of laccases. Hypotheses on their role either in pigment production or cell wall crosslinking/adhesion have been put forth, however, the available data do not allow us at the moment to select between these hypotheses, until more developmental studies and gene knockout phenotypes become available.

In addition to *bona fide* MCOs, we identified four CDE orthogroup of proteins which belong to the Cupredoxin family, like laccases, but do not contain the typical multicopper oxidase domain. These might be highly diverged MCO genes or distant relatives of true MCOs with unknown functions. The five orthogroups are conserved across Agaricomycetes (*C. cinerea* protein ID: 356026, 451850, 441442, 353997, 497434) (Table 27). One of these (*C. cinerea* protein ID: 356026) is FB-init is *A. ostoyae, C. cinerea* and *P. ostreatus* (not in *A. ampla, S. commune* and *M. kentingensis*), whereas a second (*C. cinerea* protein ID: 451850) was characteristically late upregulated in most species (*A. ampla, M. kentingensis, P. ostreatus, Pt. gracilis*). Currently, it is impossible to speculate about the role of these proteins in fruiting body development.

##### 4.8.2.5. Expansins and cerato-platanins

Fungal expansins (or expansin-like proteins) show similarity to plant expansins, proteins that modify the plant cell wall but do not show hydrolytic activity towards its polymers. Rather, they exert their action by loosening the cell wall structure, hence the name loosenins(Quiroz-Castañeda et al. 2011) and swollenins(M et al. 2002a) for some members of the family. Fungal expansins with activities towards either the plant or fungal cell wall have been reported. Certain fungal expansins can bind chitin and facilitating the action of hydrolytic enzymes by separating cell wall chitin microfibrils. Two experimentally characterized Basidiomycota expansins are Loos1 from *Bjerkandera adusta*(Quiroz-Castañeda et al. 2011) and ScExlx1 from *S. commune*(Tovar-Herrera et al. 2015). Both were shown to bind chitin and facilitate the action of chitinases, therefore, it was concluded that they modify, but not hydrolyze the chitin polymer structure. Recent studies suggested that expansin-like proteins are involved in stipe cell elongation in *F. velutipes*(Q et al. 2018) and *C. cinerea*(Niu et al. 2015) and were shown to have a role in stipe elongation, by loosening the chitin polymer structure (reviewed by(Liu et al. 2021)).

Several expansins show expression patterns compatible with a role in fruiting body development in previous transcriptome-based studies(Barsottini et al. 2013; Sipos et al. 2017b; Q et al. 2018; Almási et al. 2019; Krizsán et al. 2019). Based on analyses of 201 fungal genomes, Krizsan et al reported that expansins are enriched in the genomes of Agaricomycetes(Krizsán et al. 2019). The examined mushroom-forming fungi possess 9-27 expansin genes, with *C. cinerea* having 13. Given the dual specificity of expansins, it is likely that several of these are involved in cellulose degradation. However, a significant proportion (19-85%) of expansin genes are developmentally regulated in fruiting bodies, indicating roles in morphogenesis too. Expansins are a highly volatile group of genes that did not group into conserved orthogroups in our analysis (except two CDE orthogroups, see below), therefore, comparative analysis of their expression patterns across species is hardly possible. Within *C. cinerea* expansins show diverse expression peaks, in gills, cap or stipes with some genes highest expressed in vegetative mycelia too (Fig. 11). This could indicate tissue- and stage-specific roles for expansins in fruiting body formation. For example, four genes had a strong induction in stipes (*C. cinerea* protein ID: 451456, 464389, 359952, 461698), which is consistent with previous reports of their participation in stipe elongation(Liu et al. 2021). Three *C. cinerea* genes showed significant increase in expression from the vegetative mycelium to hyphal knots (*C. cinerea* protein ID: 461698, 286849, 496836). Despite the volatility of the family, we detected three putative expansin orthogroups (represented by *C. cinerea* 496836, 467907 and 461698) which were developmentally regulated in 12/10, 11/10 and 10/5 species (at FC>2/4) (Table 28).

**Fig. 11.**
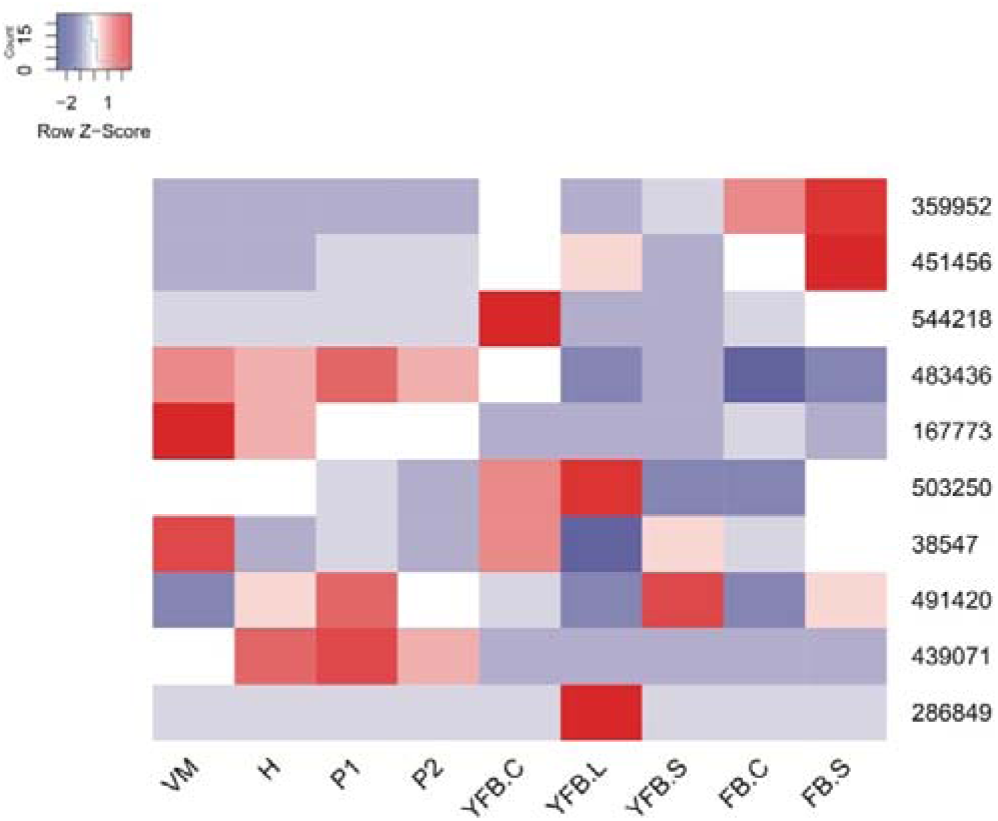
Expression heatmap of expansin-encoding genes in *C. cinerea*. Genes are denoted by Protein IDs. Blue and red colors represent low and high expression, respectively. Developmental stages are abbreviated as follows: VM – vegetative mycelium, H – hyphal knot, P1 – stage 1 primordium, P2 – stage 2 primordium, YFB.C – young fruiting body cap, YFB.L – young fruiting body gills, YFB.S – young fruiting body stipe, FB.C – mature fruiting body cap (including gills), FB.S – mature fruiting body stipe.

**Table 28.** Summary of developmental expression dynamics of CDE orthogroups of expansin genes across 12 species. Protein ID of a representative protein is given follows by the number of species in which the orthogroup is developmentally regulated at fold change 2 and 4 (FC>2 and FC>4, respectively). Putative function and ortholog in *S. cerevisiae* (if any) are also given. Abbreviations: 0-gene absent, 1-gene present but not developmentally regulated, 2 - developmentally regulated at fold change >2, 4- developmentally regulated at fold change >4. Species names are abbreviated as: Cc – *C. cinerea*, Aa – *A. ampla*, Ab – *A. bisporus*, Ao – *A. ostoyae*, Ca – *C. aegerita*, Lb – *L. bicolor*, Le – *L. edodes*, Lt – *L. tigrinus*, Mk – *M. kentingensis*, Pc – *Ph. chrysosporium*, Po – *P. ostreatus*, Sc – *S. commune*.

| ID | Fold change |  | Putative function | S.<br>*cerevisiae*<br>ortholog | Species |  |  |  |  |  |  |  |  |  |  |  |
| --- | --- | --- | --- | --- | --- | --- | --- | --- | --- | --- | --- | --- | --- | --- | --- | --- |
|  | >2 | >4 |  |  | Cc | Aa | Ab | Ao | Ca | Lb | Le | Lt | Mk | Pc | Po | Sc |
| 439071 | 6 | 5 | Cerato-platanin | – | 4 | 4 | 0 | 4 | 4 | 0 | 2 | 0 | 0 | 0 | 0 | 4 |
| 544218 | 8 | 8 | Cerato-platanin | – | 4 | 0 | 0 | 4 | 0 | 4 | 4 | 4 | 4 | 1 | 4 | 4 |
| 461698 | 10 | 5 | Expansin | – | 4 | 2 | 4 | 2 | 4 | 4 | 4 | 1 | 0 | 2 | 2 | 2 |
| 467907 | 11 | 10 | Expansin | – | 4 | 4 | 2 | 4 | 0 | 4 | 4 | 4 | 4 | 4 | 4 | 4 |
| 496836 | 12 | 10 | Expansin | – | 4 | 4 | 4 | 4 | 4 | 2 | 4 | 4 | 4 | 2 | 4 | 4 |

**Cerato-platanins** - Although initially thought to be related to hydrophobins, cerato-platanins share a common origin with expansins(Barsottini et al. 2013; Gaderer et al. 2014; Baccelli 2015; Luti et al. 2020). First described from the ascomycete *Ceratocystis platani*, they were initially and generally considered to be related to pathogenesis(Gaderer et al. 2014; Baccelli 2015; Chen et al. 2017; Luti et al. 2020), however, there is piling evidence, so far only from transcriptome studies, that they might be important in fruiting body development as well. In such studies, cerato-platanin genes showed extensive expression dynamics in fruiting bodies, suggesting an activity on the fungal cell wall(Plaza et al. 2014b; Sipos et al. 2017b; Almási et al. 2019; Krizsán et al. 2019; Liu et al. 2020b), or in defense(Kunzler 2015). Structure-based biochemistry studies in *Moniliophthora perniciosa* showed that certain cerato-platanins are able to bind chitin and are active on the fungal cell wall(Barsottini et al. 2013). Like expansins, cerato-platanins are strongly enriched in the genomes of mushroom-forming fungi(Chen et al. 2017; Krizsán et al. 2019). There is, unfortunately, little information on their mechanistic role in fruiting body development, except a the study by de Barsottini(Barsottini et al. 2013), which provides proof that cerato-platanins can be active on chitin and the fungal cell wall, providing a biochemical link to their potential role in fruiting body development.

Our analyses identified 1-8 cerato-platanins in the genomes of examined mushroom-forming fungi, which is in line with copy numbers reported recently(Chen et al. 2017). Like expansins, cerato-platanins are quite volatile and we identified only two conserved CDE orthogroups (*C. cinerea* protein ID: 439071, 544218) (Table 28), with most genes scattered in species-specific or smaller groups. The first showed opposing expression patterns in different species: in *C. cinerea* (protein ID: 544218) it was upregulated in caps relative to stipes, whereas in *A. ostoyae* and *M. kentingensis* it was upregulated in stipes relative to caps. In *P. ostreatus* and *M. kentingensis* it showed the highest expression in vegetative mycelia. In *A. ampla* and *S. commune* it showed a very strong induction (>1000-fold) in mature fruiting bodies.

Expression data across species indicates that developmentally dynamic cerato-platanin expression was observed only in certain species, while not in others. For example, no developmentally regulated cerato-platanins were found in *Ph. chrysosporium, A. bisporus*, only a single in *M. kentingensis*, but several in *A. ostoyae, C. cinerea, L. tigrinus,* etc. This indicates that developmental expression of cerato-platanins may be a species-specific, rather than a conserved feature in mushroom-forming fungi.

#### 4.8.3. Miscellaneous cell wall related families (HTP, DyP, AA3, AA5, GH76, GT8, Slt2, Cfs1, Eln3, peptidases)

We identified several CDE gene families which are potentially linked to the fungal cell wall, however, evidence is lacking on their mechanistic roles and/or they could not be confidently ascribed to either chitin or glucan, and/or are not enzymatic proteins. These are discussed in this chapter, with noting that the widely observed developmental expression makes these protein families interesting, however, their assignment to the cell wall is preliminary.

**HTPs -** Heme-thiolate peroxidases (HTPs) are fungal-specific proteins that were first described in *C. aegerita* (as haloperoxidase) and use H_2_O_2_ to modify substrates(R et al. 2004). While their enzymatic properties have been quite thoroughly studied(M et al. 2015a), their biological roles are poorly known. A remarkable expansion of the family was noted in *A. bisporus* which, combined with the upregulation of HTPs in *Agaricus* substrates led to the hypothesis that HTPs are linked to the litter decomposer lifestyle(Morin et al. 2012). In the context of fruiting bodies, HTPs were reported only in *A. ostoyae*(Sipos et al. 2017b) so far. We detected two CDE orthogroups of HTPs (represented by *C. cinerea* 479688, 442333)(Table 29) as well as several other species-specific developmentally regulated HTP genes. The examined Agaricomycetes have 2-10 genes with an expansion to 23 genes in both *M. kentingensis* and *A. bisporus* as reported earlier(Morin et al. 2012). Most of the species had at least one HTP encoding gene that is strongly upregulated (up to 50-fold) from vegetative mycelium to the first primordium stage. These observations establish HTPs as a potentially important morphogenetic gene family in fruiting body development with currently unknown function. We tentatively discuss this gene family under the chapter on cell wall modification, however, it should be mentioned that this is based on an assumption of a potential role of this family.

**Table 29.** Summary of developmental expression dynamics of CDE orthogroups of miscellaneous cell wall related genes across 12 species. Protein ID of a representative protein is given follows by the number of species in which the orthogroup is developmentally regulated at fold change 2 and 4 (FC>2 and FC>4, respectively). Putative function and ortholog in *S. cerevisiae* (if any) are also given. Abbreviations: 0-gene absent, 1-gene present but not developmentally regulated, 2 - developmentally regulated at fold change >2, 4- developmentally regulated at fold change >4. Species names are abbreviated as: Cc – *C. cinerea*, Aa – *A. ampla*, Ab – *A. bisporus*, Ao – *A. ostoyae*, Ca – *C. aegerita*, Lb – *L. bicolor*, Le – *L. edodes*, Lt – *L. tigrinus*, Mk – *M. kentingensis*, Pc – *Ph. chrysosporium*, Po – *P. ostreatus*, Sc – *S. commune*.

| ID | Fold change |  | Putative function | S.<br>*cerevisiae*<br>ortholog | Species |  |  |  |  |  |  |  |  |  |  |  |
| --- | --- | --- | --- | --- | --- | --- | --- | --- | --- | --- | --- | --- | --- | --- | --- | --- |
|  | >2 | >4 |  |  | Cc | Aa | Ab | Ao | Ca | Lb | Le | Lt | Mk | Pc | Po | Sc |
| 380322 | 6 | 4 | *C. cinerea* Eln3 | – | 4 | 0 | 4 | 4 | 1 | 4 | 2 | 0 | 2 | 0 | 0 | 0 |
| 407604 | 5 | 5 | Dye decolorising peroxidase | – | 4 | 0 | 0 | 4 | 4 | 4 | 0 | 0 | 0 | 0 | 4 | 0 |
| 447925 | 10 | 7 | *C. cinerea* Cfs1 | – | 4 | 4 | 4 | 2 | 1 | 1 | 4 | 4 | 4 | 2 | 4 | 2 |
| 469460 | 8 | 6 | Cellobiose dehydrogenase | – | 4 | 0 | 0 | 4 | 2 | 1 | 4 | 4 | 4 | 2 | 4 | 0 |
| 442333 | 11 | 7 | Chloroperoxidase | – | 4 | 2 | 4 | 4 | 4 | 2 | 4 | 2 | 4 | 1 | 4 | 2 |
| 479688 | 10 | 7 | Chloroperoxidase | – | 4 | 4 | 0 | 4 | 2 | 2 | 4 | 2 | 4 | 1 | 4 | 4 |
| 263775 | 9 | 7 | Glyoxal oxidase | – | 4 | 0 | 2 | 4 | 4 | 2 | 0 | 4 | 4 | 4 | 4 | 0 |
| 440836 | 10 | 10 | Glyoxal oxidase | – | 4 | 4 | 1 | 4 | 4 | 0 | 4 | 4 | 4 | 4 | 4 | 4 |
| 413665 | 8 | 5 | Mitogen-activated serine/threonine-protein kinase SLT2 | YHR030C | 2 | 1 | 4 | 4 | 1 | 4 | 2 | 2 | 4 | 1 | 4 | 1 |

**DyPs -** Dye decolorizing peroxidases (DyPs) are a group of fungal peroxidases that belong to the broader heme-containing peroxidase family. They are mostly discussed in the context of plant cell wall degradation by fungi(Floudas et al. 2012); to our knowledge, this is the first report of its potential role in fruiting body morphogenesis. We detected 0-5 copies (missing in *A. ampla, S. commune* and *A. bisporus*) and a patchy distribution in Agaricomycetes. However, in species that have DyP genes, these are often developmentally expressed. DyP genes show significant upregulation from vegetative mycelium to the first primordium stages in *A. ostoyae, C. cinerea, C. aegerita, L. bicolor* and *P. ostreatus*. In *M. kentingensis* all DyP genes show a marked stipe expression peak, whereas in *Pt. gracilis*, expression curves are flat. DyP genes grouped into a single CDE orthogroup (represented by *C. cinerea* 407604, Table 29). While some of the copy number variation is explained by the lignin-degrading ability of fungi(Almási et al. 2019), the developmental expression of DyP genes suggests morphogenetic roles as well, possibly in remodeling the cell wall, although this is based on assumptions distilled from the general activity of the family. It would be interesting to investigate if, similar to observations during lignin degradation, H_2_O_2_-generating enzyme gene expression is co-regulated with peroxidase expression patterns.

**AA5_1 (GLOX) -** Glyoxal oxidases (GLOX) in the AA5_1 family of auxiliary activities are H_2_O_2_ generating enzymes that facilitate the oxidative cleavage of lignin by peroxidases(Floudas et al. 2012). Surprisingly, GLOXs have also been reported to be upregulated in fruiting bodies in 6 species of Agaricomycetes(Krizsán et al. 2019) and were detected in fruiting bodies of *Auricularia polytricha* by protein mass spectrometry(Jia et al. 2017). Since lignin is obviously missing from fruiting bodies, they were speculated to contribute to cell wall remodeling(Krizsán et al. 2019). We found 1-22 GLOX genes in the examined Agaricomycetes, the fewest in the Schizophyllaceae (*A. ampla, S. commune*) possibly because of their reduced morphology or reduced lignin-degrading ability(Almási et al. 2019). Two CDE orthogroups (*C. cinerea* protein ID: 263775 and 440836 Table 29), and several species-specific developmentally regulated GLOX genes were found in our data. We found GLOX genes significantly upregulated in primordia relative to vegetative mycelium (FB-init) in *A. ostoyae, A. ampla, C. cinerea, L. tigrinus, L. bicolor, S. commune, Ph. chrysosporium, P. ostreatus and C. aegerita* (not in *M. kentingensis, L. edodes* and *A. bisporus*). Two of the three *C. neoformans* GLOX genes also showed significant upregulation during basidium development. These data indicate that GLOXs may be important morphogenetic factors in the Basidiomycota.

**AA3_1 -** cellobiose dehydrogenases in the AA3_1 family (GMC oxidoreductases) are known to donate electrons to LPMOs during lignocellulolysis in wood-decay fungi(Floudas et al. 2012). There are only two reports on the potential role of CDHs in fruiting body development, one is a comparative transcriptomics study(Krizsán et al. 2019) and the other is a report of the interaction between CDH and laccases in the formation of cinnabarinic acid, the red fruiting body pigment of *Pycnoporus cinnabarinus*(Temp and Eggert 1999). A CDE orthogroup of cellobiose dehydrogenase (CDH) genes (*C. cinerea* protein ID: 469460)(Table 29) was developmentally regulated in all species in which it was present. Additional CDH genes (2-8 genes in Agaricomycetes) were developmentally regulated in a species-specific manner in several species, although expression levels were low in many cases. The biological significance of CDH in fruiting body development is currently unknown; based on previous studies it may be related to cell wall remodeling or pigment production.

**Slt2 kinase family -** A serine-threonine kinase orthologous to the MAP kinase involved in the cell wall integrity pathway of *S. cerevisiae* (Slt2) was detected as a CDE orthogroup (*C. cinerea* protein ID: 413665). Slt2 is a terminal kinase in the pathway that phosphorylates downstream transcription factors (Rlm1 and Swi4/6 in yeast)(Levin 2005). Although the gene is developmentally regulated in several species (Table 29), expression patterns are not consistent across taxa. Given the potentially pleiotropic role of this kinase, it is hard to guess its role in fruiting bodies, but extrapolations from its role in *S. cerevisiae* would suggest links to cell wall integrity, morphogenesis or the cell cycle.

**Cfs1 family -** The orthogroup containing *C. cinerea* Cfs1 (*C. cinerea* protein ID: 447925), a cyclopropane fatty acid synthase, was developmentally regulated in 10/7 (FC>2/4) species (Table 29). Cfs1 was identified in a mutant screen of strains with developmental defects; its mutation leads to failures in the progression of hyphal knot development(Liu 2005) in *C. cinerea* (stalled at primary hyphal knot). The corresponding gene is highly induced 1h after blue light exposure(Sakamoto et al. 2018). We discuss this gene under the cell wall, although some evidence suggests it may be more closely related to plasma membrane assembly. Structurally, Cfs1 and related proteins are mycolic acid cyclopropane synthases, a family that occurs in 1-6 copies in the examined Agaricomycetes. Their function is not known, but presumably they are involved in the synthesis of long fatty acids (mycolic acid is not known to occur in fungi). In most examined species (*A. ampla, M. kentingensis, C. cinerea, P. ostreatus, S. commune*), the gene is upregulated in primordia relative to vegetative mycelium, consistent with its phenotypic role in hyphal knot development.

**Eln3 family -** This family is based on the *Eln3* gene of *C. cinerea*, which was found in a REMI mutagenesis screen. The mutant strain displayed defects in stipe elongation and mutant stipe cells were disorganized and there were large gaps among stipe cells, whereas in the wild type strain stipe cells align closely with each other(Liu 2005). It has been hypothesized that *Eln3* is involved in cell-cell connections, possibly adhesion.

We found that *C. cinerea Eln3* (*C. cinerea* protein ID: 380322) formed a small CDE orthogroup present in 7 species and was developmentally regulated in 6 species (Table 29). In these species the gene was predominantly FB-init and stipe upregulated, in agreement with a previously reported role in stipe elongation(Arima et al. 2004). It should be noted that, in many species, upregulation in primordia is easy to confuse with stipe-specific upregulation, because primordia often comprise predominantly stipe tissue. In general, we found 7-8 genes the examined Agaricomycetes (only 4 in *A. bisporus*) with identical domain composition to *Eln3*. This does not guarantee identical function, nevertheless is indicative of the diversity of this family in mushrooms. These genes were often FB-init and showed stipe or cap-specific upregulation depending on the gene and the species. For example, in *C. cinerea* we found that 5 out of 6 genes were stipe-upregulated. In *A. ostoyae* we found four cap- and two stipe-upregulated, whereas in *L. bicolor* two cap- and one stipe-upregulated gene. The widespread developmental regulation of *Eln3* and its homologs suggest morphogenetic functions, possibly in association with the cell wall. Based on its domain composition, we hypothesize it represents a cell wall active, possibly new CAZyme family not yet recognized by Cazy.org.

**GT8 -** The GT8 family has 1-7 (most frequently 5) copies in the Agaricomycetes. The GT8 family includes both N-acetylglucosaminyltransferases linked to the modification of N-linked glycans in the Golgi, and glycogenins involved in glycogen metabolism. Buser et al hypothesized that fruiting body-specific N-glycans generated by GT8 enzymes may contribute to adhesion in *C. cinerea*(Buser et al. 2010a). GT8 genes formed one CDE orthogroup, represented by *C. cinerea* 461263 FC2/4: 7/3 species), which is closest to *S. cerevisiae* Glg1 glycogenin (discussed under carbohydrate metabolism). Of the other GT8 genes *C. cinerea* (*C. cinerea protein ID:* 453097, =*CcGnt1)*(Buser et al. 2010a)*, A. ostoyae, L. tigrinus, P. ostreatus, A. ampla* and *S. commune* had one FB-init gene each, however, these did not form CDE orthogroups. CcGnt2 (*C. cinerea* protein ID: 412828) was not expressed. Other species did not have developmentally regulated genes. Due to the variability of the family, it is hard to determine whether these are closer to glycogenins or N-acetylglucosaminyltransferases.

**N-glycosylation -** Mannosylation and N-glycosylation of proteins in the ER is a key process for cell wall synthesis. Berends et al provided a detailed view of the N-glycosylation machinery in *S. commune, C. cinerea* and *L. bicolor* based on orthology to the *S. cerevisiae* N-glycosylation pathway members and expression data in *S. commune* fruiting bodies and monokaryotic mycelium(Berends et al. 2009). They conclude that Agaricomycetes have the gene repertoire to produce oligomannose structures only, in contrast to some Ascomycota which can produce very complex modifications to proteins. Our expression results broadly agree with theirs in that we detected very low dynamics of corresponding genes (not shown) in the fruiting body transcriptomes analyzed here.

### 4.9. Cell surface and cell wall proteins

Secreted cell surface proteins fulfill diverse functions in the life of fungi, ranging from signaling, interactions with the environment, adhesion, cell wall modifications, but also communication with hosts in mutualistic or pathogenic interactions(Pellegrin et al. 2015; Essen et al. 2020). The fungal cell surface proteome and secretome profile of each species in a given environment is unique, making constituents of fungal communities recognizable by their extracellular protein profiles(Campbell et al. 2015). Secreted proteins have been the subject of intense research, mostly in the context of interactions between fungi and other organisms, adhesion (e.g. via GPI-anchored adhesins) of fungal cells to various surfaces(Essen et al. 2020; M et al. 2020b). Fruiting bodies, as environmentally exposed, 3-dimensional multicellular structures, can be expected to display unique cell-surface attributes. In fact, cell surface proteins include some of the best-known fruiting body-associated proteins (e.g. hydrophobins, lectins, see below) or proteins related to defense (see these in a separate chapter). Conceivably, adhesion and/or cell-to-cell communication, two of the most important multicellular functions(Knoll 2011), could be realized via the action of secreted cell surface proteins in fruiting bodies, to just name a few of the diverse ways mushroom-forming fungi might make use of their cell surface protein repertoire. A significant open question in fruiting body biology is the nature and role of the extracellular matrix and the biochemical and communication events that take place in the interhyphal space. Cell surface proteins are undoubtedly important players in these processes, and therefore represent interesting targets of future research.

While some cell surface proteins are the best examined fruiting body proteins (e.g. hydrophobins), the following inventory of conserved developmentally expressed genes shows that there is plenty of new insight to be learned about the cell surface proteome of Agaricomycetes fruiting bodies. In this chapter we discuss cell surface protein families in the context of CDE orthogroups detected in our meta-analysis of fruiting body transcriptomes.

#### 4.9.1. Hydrophobins

Hydrophobins are some of the best-known proteins in fruiting bodies. They confer hydrophobicity to hyphae, which they achieve by self-assembling into a water-repellent rodlet layer on the cell surface. Hydrophobins were hypothesized to line air channels in fruiting bodies(Lugones et al. 1999b), but also contribute to water sealing, which is important in fruiting bodies as they rely on water transported to them from the vegetative mycelium(Nehls and Dietz 2014). The family has extensively literature, in general(Wösten 2003; Linder et al. 2005; Bayry et al. 2012), in relation to fruiting body development(Ohm et al. 2010; Kües and Navarro-González 2015; Kim et al. 2016; Sammer et al. 2016; Y et al. 2019) or on potential uses in bioengineering(Wösten and Scholtmeijer 2015). We therefore concentrate on novel aspects detected in our data that have not or only rarely been reported before.

Basidiomycota hydrophobins mostly belong to class I hydrophobins(Sammer et al. 2016). On average 11-44 hydrophobin genes (only 6 in *M. kentingensis*) were detected in the examined Agaricomycetes, a significant proportion of which were developmentally regulated. They can be arranged into distinct groups based on expression patterns. Previous studies reported a near complete turnover of hydrophobin genes from vegetative mycelium to fruiting body, indicating that two distinct sets of hydrophobins exist(Ma et al. 2007; WW et al. 2008; Cheng et al. 2013a). We also observed that mycelium-expressed and fruiting body-expressed hydrophobins form distinct groups. For example, the numbers of hydrophobins showing mycelium- vs fruiting body specific expression was 6/5, 1/9, 10/4, 23/3 and 5/20 (at fold change 4/2) in *Ph. chrysosporium, S. commune, P. ostreatus, A. ostoyae* and *C. cinerea*, respectively. *A. ostoyae* further has two rhizomorph-specifically expressed hydrophobins, indicating morphogenesis-related diversification of the family. This is in line with observations in *F. velutipes,* where most hydrophobin genes were found to be primordium-upregulated(Kim et al. 2016). We also detected hydrophobin genes expressed during early development, but not later, indicating functions linked to primordium development, which are turned off during later development. These observations further substantiate the functional diversification, probably driven by morphogenesis and complexity.

Despite the large diversity and developmental expression of hydrophobins, only 2 CDE orthogroups were detected (*C. cinerea* protein ID: 99661, 419467) (Table 30), which were present in only a subset (11 and 8 species, respectively) of the examined species. This is due to the frequent duplication of hydrophobin genes(Mgbeahuruike et al. 2017), which quickly erases clear orthology relationships. CDE orthogroups did not show conserved expression patterns. Interestingly, *M. kentingensis* has only 6 hydrophobin genes, none of which were developmentally expressed in our reanalysis of data from Ke et al(Ke et al. 2020a). A possible explanation for this is that this species produces hygrophanous, rather than hydrophobic fruiting bodies, which absorb rather than repel water. The relatively low number of developmentally regulated hydrophobin genes in *A. ostoyae* may also be explained by the hygrophanous caps of this species. Hygrophanicity is a widespread trait in the Agaricales and might explain some of the differences in hydrophobin diversity and expression.

**Table 30.** Summary of developmental expression dynamics of CDE orthogroups of hydrophobin genes across 12 species. Protein ID of a representative protein is given follows by the number of species in which the orthogroup is developmentally regulated at fold change 2 and 4 (FC>2 and FC>4, respectively). Putative function and ortholog in *S. cerevisiae* (if any) are also given. Abbreviations: 0-gene absent, 1-gene present but not developmentally regulated, 2 - developmentally regulated at fold change >2, 4- developmentally regulated at fold change >4. Species names are abbreviated as: Cc – *C. cinerea*, Aa – *A. ampla*, Ab – *A. bisporus*, Ao – *A. ostoyae*, Ca – *C. aegerita*, Lb – *L. bicolor*, Le – *L. edodes*, Lt – *L. tigrinus*, Mk – *M. kentingensis*, Pc – *Ph. chrysosporium*, Po – *P. ostreatus*, Sc – *S. commune*.

| ID | Fold change |  | Putative function | S.<br>*cerevisiae*<br>ortholog | Species |  |  |  |  |  |  |  |  |  |  |  |
| --- | --- | --- | --- | --- | --- | --- | --- | --- | --- | --- | --- | --- | --- | --- | --- | --- |
|  | >2 | >4 |  |  | Cc | Aa | Ab | Ao | Ca | Lb | Le | Lt | Mk | Pc | Po | Sc |
| 99661 | 7 | 7 | Hydrophobin | – | 1 | 4 | 0 | 0 | 4 | 1 | 4 | 4 | 0 | 4 | 4 | 4 |
| 419467 | 6 | 6 | Hydrophobin | – | 4 | 4 | 4 | 4 | 0 | 4 | 1 | 0 | 0 | 1 | 0 | 4 |

Taken together, hydrophobins, as already established by previous research, are one of the best-known fruiting body-related structural cell surface proteins in the Agaricomycetes, with almost all species expressing developmental stage-specific suites of hydrophobin genes.

#### 4.9.2. Lectins

Lectins will be also discussed in Chapter 10 on defense-related proteins, however, a role in defense has been proven for only a subset of the diverse lectin repertoires of mushroom-forming fungi; the remaining non-defense, or functionally unknown lectins are discussed in this chapter (even though not all lectins may be secreted or their secretion status is unclear at the moment, (Boulianne et al. 2000)). Lectins are proteins that specifically recognize and bind various carbohydrate moieties, mostly on the cell surface. This ability makes them suitable for multiple biological roles, from bacterial agglutination and cytotoxicity against mushroom grazers(Bleuler-MartÍnez et al. 2011; Kunzler 2015) to carbohydrate binding modules that fulfill roles related to the fungal cell wall (e.g. remodeling), adhesion, cell-to-cell communication or other functions in the extracellular matrix. Their carbohydrate binding ability makes them interesting in a wide range of biomedical fields(Hassan et al. 2015), including cancer research, which led to the broader lectin family rapidly expanding. Mushroom fruiting bodies remain one of the richest sources of lectins for applied research. However, regarding their physiological role for the fungi, boundaries between defense-related and other lectins are far from clear at the moment(Bleuler-MartÍnez et al. 2011). Due to interest in their practical applications, there is often more information on the structure of mushroom lectins than on their roles in the fungus’ life. Fruiting bodies express a diverse set of lectins (reviewed by(A et al. 2013; Kobayashi et al. 2014; Y and H 2014)) and several of the well-known lectin families were first isolated from mushrooms. For example, the fungal fruit body lectin family was established based on the *Xerocomus chrysenteron* XCL protein, which was suggested to have insecticidal activity(C et al. 2004). Besides interest in their biotechnological applications, lectins have been in the focus of research on fruiting bodies. Early reports identified specific hyphal knot and primordium upregulated galectins (Cgl1 and Cgl2) in *C. cinerea*(Boulianne et al. 2000; Bertossa et al. 2004), which were initially postulated to be cell-cell adhesive or communication related molecules that may bind basidiolipids(Walser et al. 2005). Simultaneous silencing of the *cgl1* and *cgl2* genes did, however, not disturb fruiting, arguing against his hypothesis(MA et al. 2006). The lectins were rather shown to inhibit the development of insect and nematode larvae, in nematodes via binding to a specific N-glycan epitope, suggesting a function in defense against predators(Butschi et al. 2010; Bleuler-MartÍnez et al. 2011). Both genes were found upregulated in primordia, relative to vegetative mycelium in recent RNA-Seq based studies (cgl1 = *C. cinerea* 473274, cgl2 = *C. cinerea* 488611). Later, Wälti et al described cgl3 (=*C. cinerea* 539794), which resembles galectins, but binds chitooligosaccharides and had no effect on larval development of insects and nematodes(Wälti et al. 2008). Beta-glucan binding lectins of *Serepindita indica* may be involved in fungal cell wall remodeling at the plant-fungal interface(Wawra et al. 2019). Several other recent reports of individual lectins from a diverse set of medicinal or edible mushrooms is available but their detailed discussion is beyond the scope of this paper. Lectins are also involved in Ascomycota fruiting body (perithecium) formation and interestingly, the gene recently described in *Podospora anserina* showed highest similarity to *Xerocomus* lectins(Nowrousian and Cebula 2005). Several lectin families are overrepresented in mushroom-forming fungi(Krizsán et al. 2019), suggesting key roles in their life cycle.

‘Lectomes’ evolve fast (like other defense genes) both in terms of sequence and gene content. Most lectin families a phylogenetically patchily distributed in mushroom forming fungi, with the probable sole exception of ricin-B lectins, which occur in all examined species. It has been suggested that the fast gene turnover can contribute to the expression of defense gene cocktails by the fungus, which is advantageous and that horizontal gene transfer from bacteria can provide a mechanistic explanation for the phylogenetic patchiness of several lectin families(Moran et al. 2012; Kunzler 2015).

We here provide an overview of lectin copy numbers in the examined Agaricomycetes in the context of 8 main families, but note that more recent classifications(Lebreton et al. 2021) significantly reorganized lectin families and may provide a more nuanced view on lectin repertoires. Lectin copy numbers and expression patterns were summarized in the context of 6 Agaricomycetes species by Krizsan et al(Krizsán et al. 2019) with lectins showing the RicinB fold (which includes protease inhibitors too, see above and(J et al. 2016)) being most diverse and present in all species, whereas other families are more patchy. Nevertheless, all lectin families had developmentally regulated genes, often showing strong induction in primordium stages.

As noted earlier (e.g. (Kunzler 2015)), lectins are poorly conserved and hence mostly did not form conserved orthogroups in our analysis. Only two CDE orthogroups, all belonging to the RicinB family were detected (*C. cinerea* protein ID: 495335, 353931)(Table 31). Both orthogroups contained secreted proteins. Expression patterns were poorly conserved within these orthogroups, but we note that each had FB-init genes in several species, indicating that these secreted Ricin-B family members are involved in the initiation of fruiting body development.

**Table 31.**
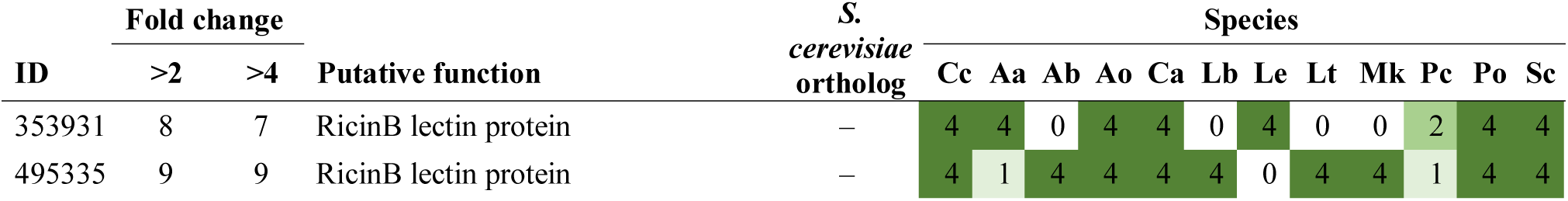
Summary of developmental expression dynamics of genes belonging to conserved lectin orthogroups across 12 species. Protein ID of a representative protein is given follows by the number of species in which the orthogroup is developmentally regulated at fold change 2 and 4 (FC>2 and FC>4, respectively). Putative function and ortholog in *S. cerevisiae* (if any) are also given. Abbreviations: 0-gene absent, 1-gene present but not developmentally regulated, 2 - developmentally regulated at fold change >2, 4- developmentally regulated at fold change >4. Species names are abbreviated as: Cc – *C. cinerea*, Aa – *A. ampla*, Ab – *A. bisporus*, Ao – *A. ostoyae*, Ca – *C. aegerita*, Lb – *L. bicolor*, Le – *L. edodes*, Lt – *L. tigrinus*, Mk – *M. kentingensis*, Pc – *Ph. chrysosporium*, Po – *P. ostreatus*, Sc – *S. commune*.

#### 4.9.3. Other cell surface proteins (HsbA, Con6, PriA, AA7, CFEM, wax synthases, fasciclins, peptidases, intradiol ring cleavage dioxygenase)

**The HsbA family** comprises poorly known cell wall galactomannoproteins that includes HsbA from *Aspergillus oryzae*(S et al. 2006b), a *Magnaporthe* homolog(Soanes et al. 2012) as well as genes named Mp1 in *Talaromyces marneffei*(Cao et al. 1998) and in *Aspergillus fumigatus*(KY et al. 2001). Detailed functional information is not available for either of these genes. The family contains GPI-anchored cell surface proteins that can bind to various surfaces and was circumscribed in the context of appressorium attachment and lignocellulose degradation in Ascomycetes. The common denominator between these circumstances may be the need for adhesion of hyphae to various surfaces, enabling secreted lytic enzymes to exert their action. In the context of fruiting bodies, the family was first reported in fruiting bodies of *A. ostoyae*(Sipos et al. 2017b). In our current analysis, we detected 0-3 members of the family in Agaricomycetes (missing from *L. tigrinus, Ph. chrysosporium, A. ampla. S. commune*), all of which were developmentally expressed. The family did not form CDE orthogroups. Expression of members of this family showed significant dynamics (>100-fold), mostly with gill- or cap-specific expression peaks late during development. Functional analysis of this gene family in mushroom-forming fungi would be very interesting. If, as speculated above, the family is related to adhesion, then it may provide a missing link to adhesion in complex multicellularity in mushrooms.

**The Con6 protein family** include small hydrophilic peptides that were first discovered as conidium-specific proteins in *N. crassa* and later shown in *A. nidulans* (ConF and ConJ) to be responsible for desiccation resistance of asexual spores(Suzuki et al. 2013). The family also includes the meiotically expressed *Sch. pombe* gene *cum1*. The Con6 family is conserved in Agaricomycetes, with 1-7 copies in the examined genomes, most of which are developmentally regulated. They showed two major expression patterns in all species: (i) strong induction in mature gills or combined gill+cap samples and/or (ii) upregulation in primordia relative to vegetative mycelia (Fig. 12, Supplementary Fig. 8). For some genes both for others only one of these expression patterns were observable. The family seems poorly conserved, nevertheless, they formed three CDE orthogroups (represented by *C. cinerea* 470234, 186809, *S. commune* 2624349)(Table 32). The gill-specific upregulation, combined with the conidium-specific expression of the gene family in filamentous Ascomycota, suggests that Con6 proteins may be involved in producing a spore-specific protein coating in these species. The orthogroup represented by *C. cinerea* 186809 showed upregulation in primordia of most species and has low expression in vegetative mycelia and later developmental stages. The strong upregulation of several Con6-encoding genes in primordia relative to vegetative mycelia in virtually all species suggests that it has general roles in fruiting bodies as well, possibly in regulating hydrophobicity/hydrophilicity of the cell surface, although this requires further research.

**Fig. 12.**
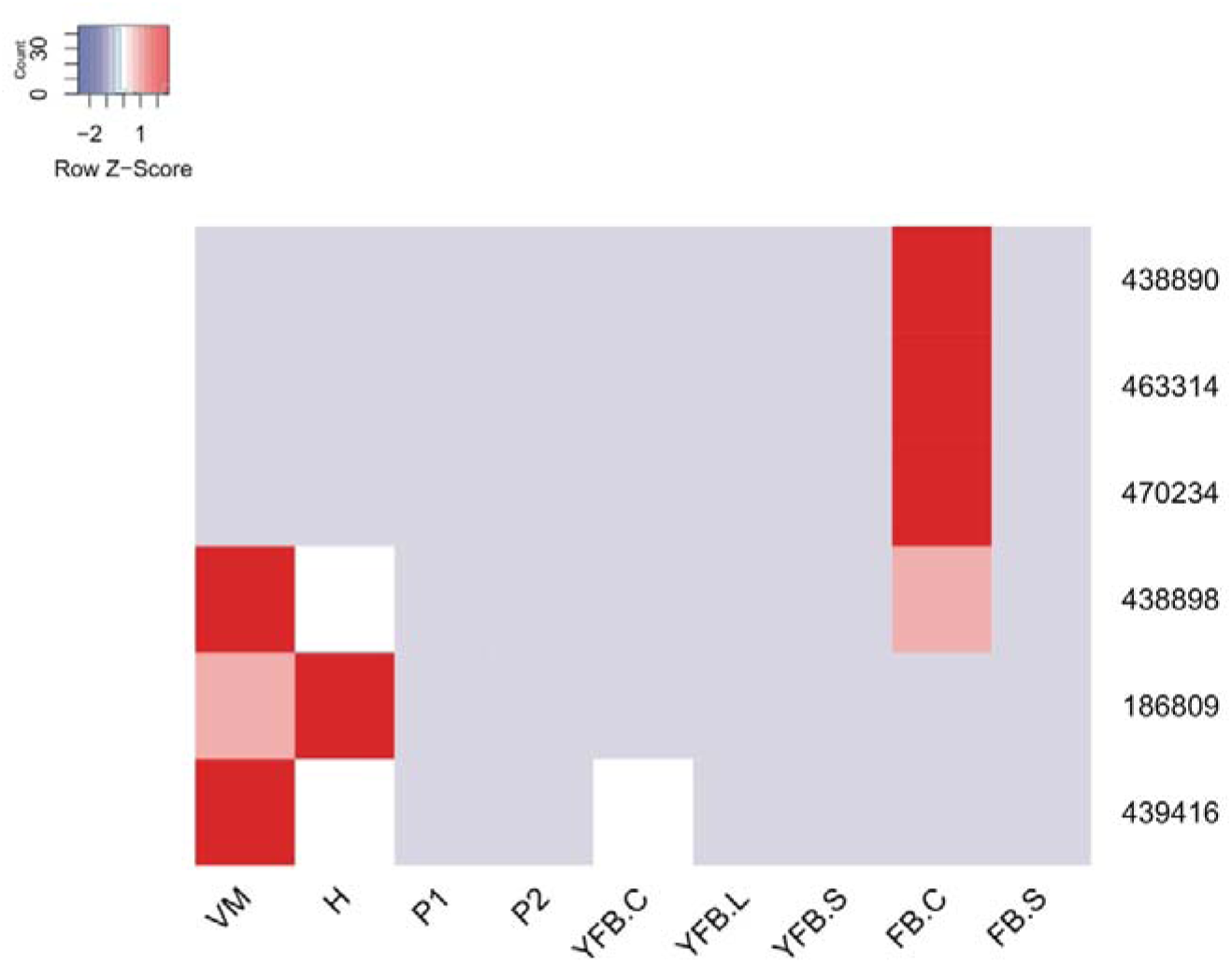
Expression heatmap of Con6 family cell surface protein-encoding genes in *C. cinerea*. Genes are denoted by Protein IDs. Blue and red colors represent low and high expression, respectively. Developmental stages are abbreviated as follows: VM – vegetative mycelium, H – hyphal knot, P1 – stage 1 primordium, P2 – stage 2 primordium, YFB.C – young fruiting body cap, YFB.L – young fruiting body gills, YFB.S – young fruiting body stipe, FB.C – mature fruiting body cap (including gills), FB.S – mature fruiting body stipe.

**Table 32.**
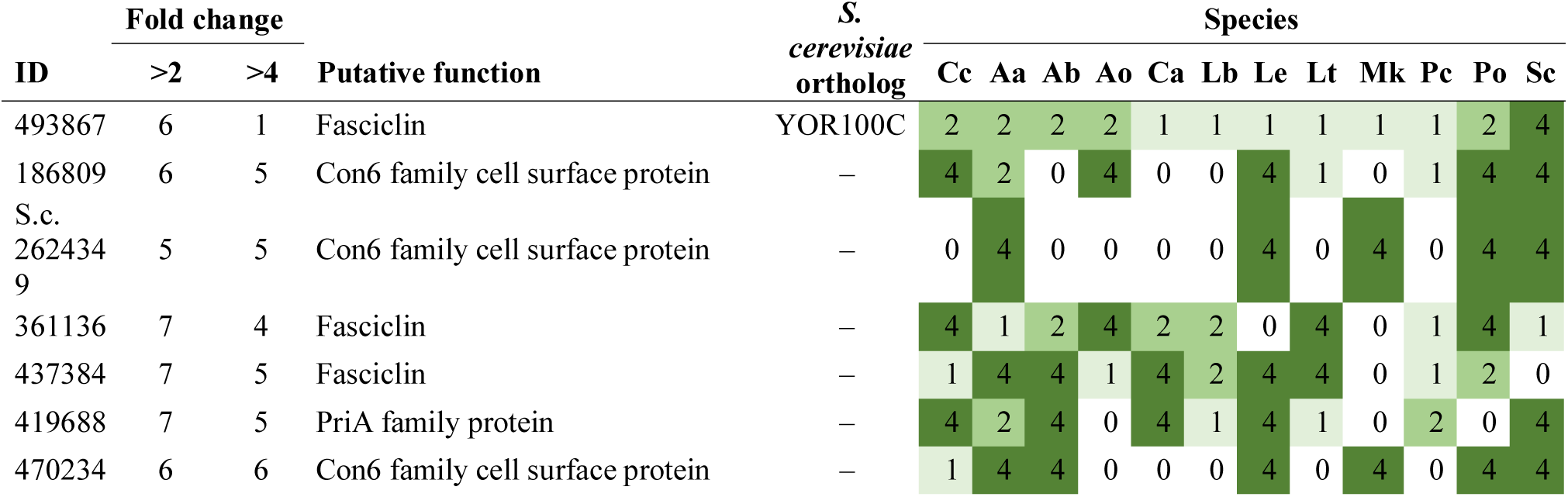
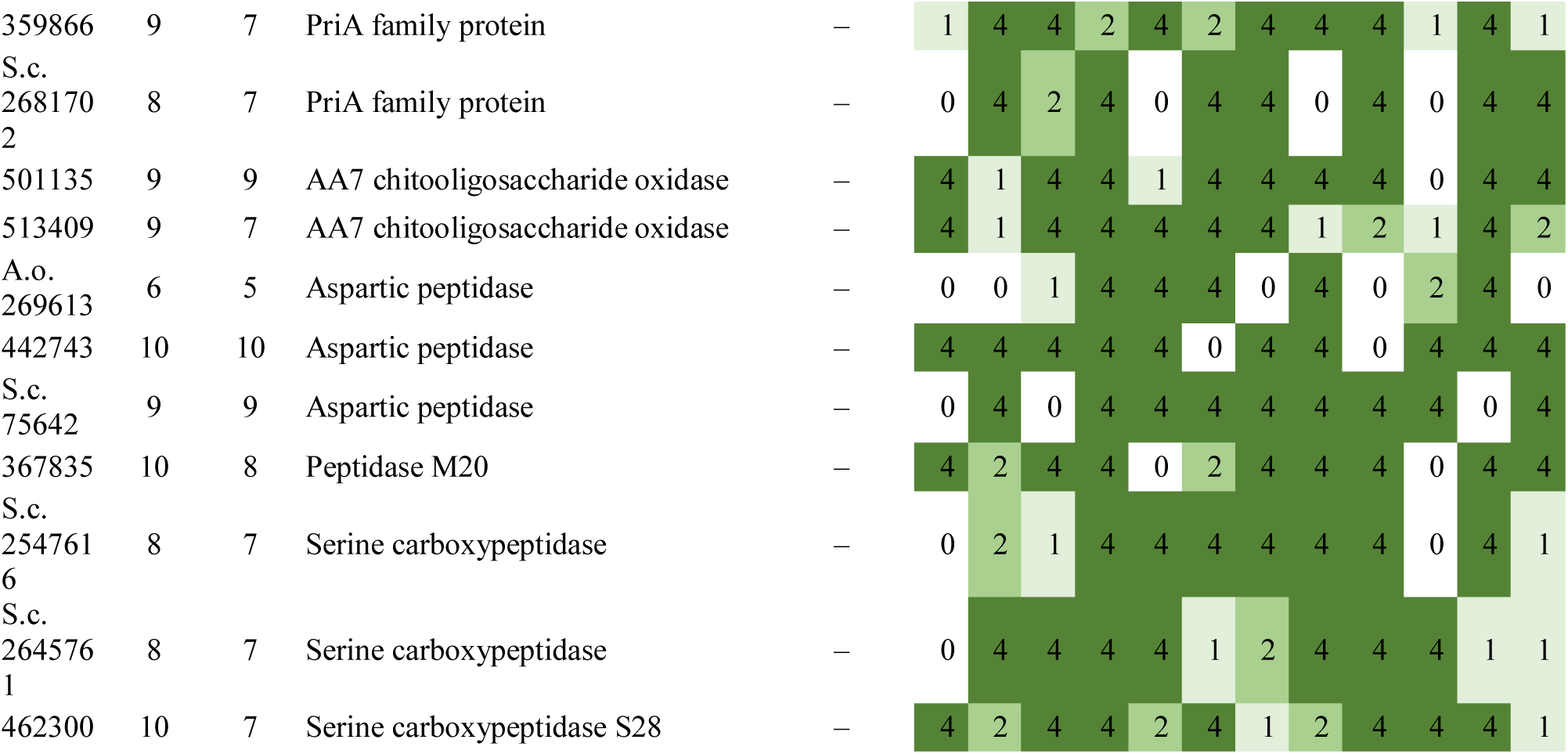
Summary of developmental expression dynamics of CDE orthogroups of cell surface protein encoding genes across 12 species. Protein ID of a representative protein is given follows by the number of species in which the orthogroup is developmentally regulated at fold change 2 and 4 (FC>2 and FC>4, respectively). Putative function and ortholog in *S. cerevisiae* (if any) are also given. Abbreviations: 0-gene absent, 1-gene present but not developmentally regulated, 2 - developmentally regulated at fold change >2, 4- developmentally regulated at fold change >4. Species names are abbreviated as: Cc – *C. cinerea*, Aa – *A. ampla*, Ab – *A. bisporus*, Ao – *A. ostoyae*, Ca – *C. aegerita*, Lb – *L. bicolor*, Le – *L. edodes*, Lt – *L. tigrinus*, Mk – *M. kentingensis*, Pc – *Ph. chrysosporium*, Po – *P. ostreatus*, Sc – *S. commune*.

**Wax synthases** are long-chain alcohol O-fatty-acyltransferases that synthesize wax esters, a typical constituent of the cuticula of plants or the sebum of the animals(Li et al. 2008). To our best knowledge, only Krizsan et al(Krizsán et al. 2019) indicated developmental expression for this family previously.

We detected 4-16 wax synthases in the examined Agaricomycetes, the fewest in *A. bisporus*, but found no CDE orthogroups, suggesting a high duplication rate in the family. However, each species (except *Pt. gracilis* and *A. bisporus*) had 1-2 FB-init wax synthase genes, which suggests fruiting body specific roles for these genes. We observed considerable (>100-fold) upregulation compared to vegetative mycelia in many cases (Fig. 13, Supplementary Fig. 9). Given the widespread upregulation of wax synthases in mushroom-forming fungi, it is possible that they produce a wax-like layer on the surface of fruiting body cells. On plants, wax-coating of leaves serves as a defense system that inhibits penetration of pathogenic fungi and water-sealing(Lewandowska et al. 2020). Products of fungal wax synthases, although unknown at the moment, could have similar properties in fruiting bodies, worth investigating in the future.

**Fig. 13.**
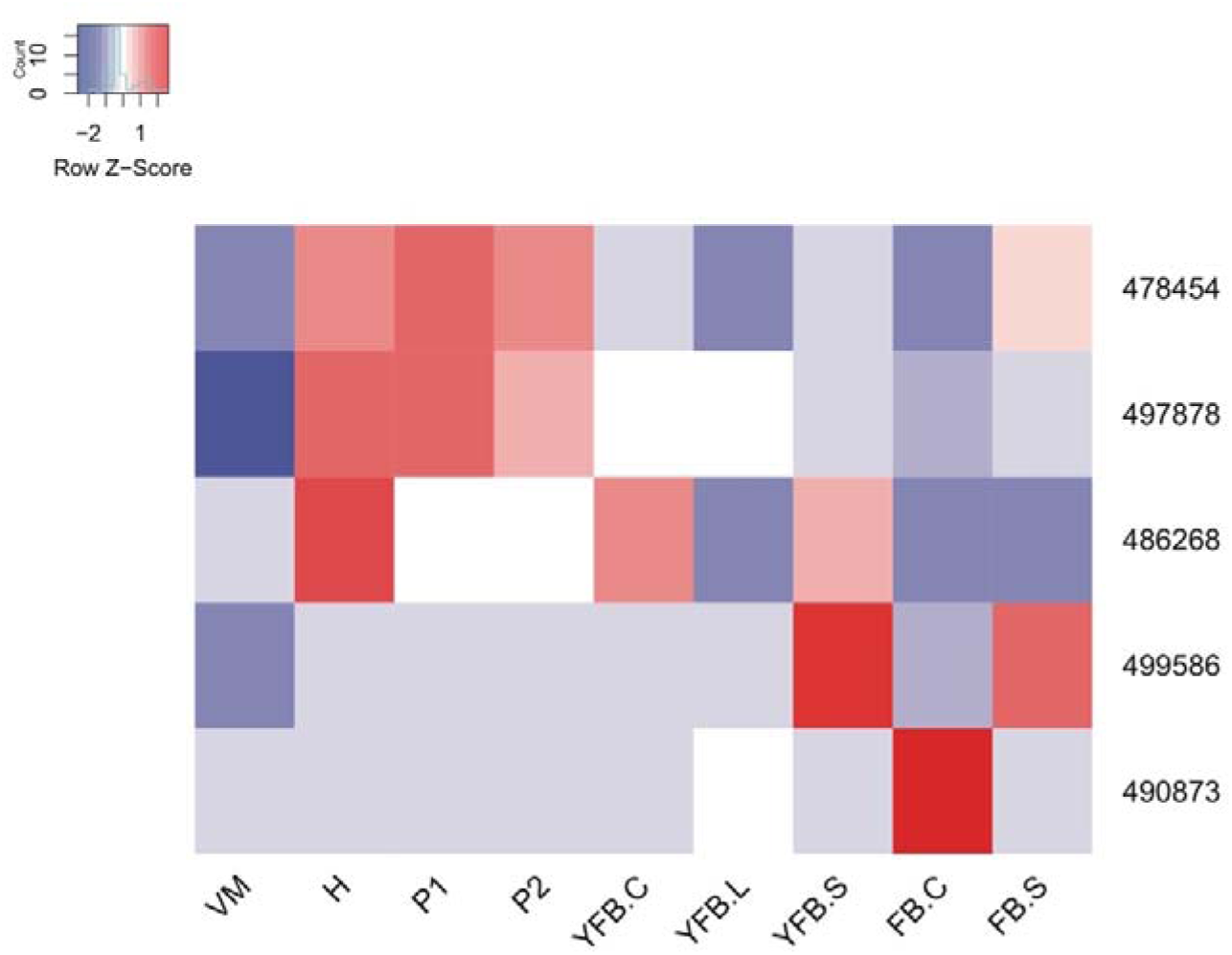
Expression heatmap of putative wax synthase encoding genes in *C. cinerea*. Genes are denoted by Protein IDs. Blue and red colors represent low and high expression, respectively. Developmental stages are abbreviated as follows: VM – vegetative mycelium, H – hyphal knot, P1 – stage 1 primordium, P2 – stage 2 primordium, YFB.C – young fruiting body cap, YFB.L – young fruiting body gills, YFB.S – young fruiting body stipe, FB.C – mature fruiting body cap (including gills), FB.S – mature fruiting body stipe.

**PriA family** - this family comprises secreted cell surface proteins first described as a primordium-induced protein in *L. edodes* (hence its name)(S et al. 1992). The gene has been quite extensively studied in *L. edodes*(T and K 2000; T et al. 2004a), where a link to zinc homeostasis has been suggested. It was also detected as developmentally regulated in *F. velutipes*(Liu et al. 2020b)*, P. eryngii*(Fu et al. 2017), *L. edodes*(S et al. 1992) and in six species by Krizsan et al(Krizsán et al. 2019). The family also includes the virulence-related secreted adhesin Cfl1 of *C. neoformans*, which is highly induced during sexual development and has been thoroughly researched in *C. neoformans*(Wang et al. 2012, 2013). Cfl1 has been suggested to act as a morphogen that induces neighboring cells to phenocopy the producing cells’ morphology(Wang et al. 2013). Cfl1 has 4 homologs in *C. neoformans*, which have stage-specific expression including Dha1, which is primarily expressed in basidia(Gyawali et al. 2017). The family is present in 0-4 copies in the Agaricomycetes and formed two CDE orthogroups. The first of these (*C. cinerea* protein ID: 359866)(Table 32) showed a very conserved, mycelium biased expression; it was highly expressed in vegetative mycelia and the first primordium stage, then virtually unexpressed in all later stages in *A. ampla S. commune, M. kentingensis* and *P. ostreatus*. In *A. ostoyae* expression dropped immediately after the mycelium sample. The second CDE orthogroup (represented by *C. cinerea* 419688) contains best hits of *C. neoformans* Dha1 and showed a peak expression in gills and caps of *C. cinerea, A. bisporus, C. aegerita*, *L. edodes* and late stages of *S. commune* and *A. ampla.* At the same time, these genes were significantly upregulated (up to 50-100x) in primordia relative to vegetative mycelium. In *L. edodes*, our RNA-Seq based observations are consistent with the first reports of the PriA gene as a primordium-upregulated one(S et al. 1992). The third orthogroup (represented by *S. commune* 2681702), was present in only 8 species, and was FB-init in *A. ampla, A. ostoyae, P. ostreatus, M. kentingensis* and *S. commune* (Table 32). In summary, the PriA family appears to be a very interesting family from the perspective of fruiting body development. In *C. neoformans* adhesive and signaling roles have been suggested for the family, although exact mechanisms of function are still unknown.

**Aspartic peptidase A1** - This family is a versatile group of endopeptidases that has undergone a characteristic expansion in fungi(Revuelta et al. 2014). They were reported to be involved in pathogenesis, nutrition, cell wall maintenance and a range of other functions. The best researched family of A1 aspartic peptidases are yapsins, which are involved in cell wall assembly and remodeling, although their function is not clear; limited evidence suggests they proteolytically process secreted (cell wall) proteins(Gagnon-Arsenault et al. 2006; Gow et al. 2017; M et al. 2021). Based on functional knowledge on yapsins, and the abundance of GPI-anchored A1 aspartic peptidases in fungi, we discuss this family in the cell surface section, despite almost certainly not all of them are related to the cell wall. The family has been reported to be upregulated in fruiting bodies by several studies previously(Rahmad et al. 2014; Song et al. 2018b; Krizsán et al. 2019). Aspartic peptidase A1 family was found as 15-60 copies in Agaricomycetes genomes. We detected 3 CDE orthogroups and several species-specifically developmentally expressed genes (Table 32), though these did not display strong expression conservation in the examined species. Nevertheless, the developmental expression of several members of this family, combined with evidence on yeast yapsins(Gagnon-Arsenault et al. 2006) and the secreted nature of these proteins suggest a cell wall related role for aspartic peptidases which may be worth investigating. Of note, we detected further CDE orthogroups of various peptidase families (M20, M28, M16, S10), which may also be involved in fruiting body development (Table 32), but our functional knowledge at the moment is too limited to speculate about their roles. Other peptidase families that were reported in mushroom-forming fungi include the metalloprotease M43 family, which was reported from *P. ostreatus*(Joh et al. 2004) and has a strong (>700x) induction upon fruiting in the data published by Merenyi et al(Merényi et al. 2021).

**AA7 family** - One of the CDE orthogroups (represented by *C. cinerea* 501135)(Table 32) were top hits of *Fusarium graminearum* ChitO, the only experimentally examined member of the family known to us(Heuts et al. 2007), which oxidises chitooligosaccharide species. AA7 comprises oligosaccharide oxidising flavo-enzymes which also contain a Berberine-like domain and produce H_2_O_2_ during their action. They have been speculated to be involved in the detoxification of lignocellulose degradation intermediates and in the production of various secondary metabolites(Levasseur et al. 2013). Based on the activity of their homologs and their expression in fruiting bodies, it is likely that these AA7 family members are active on chitin, although their exact roles remain obscure for now.

**CFEM domain proteins** - We detected 2 CDE orthogroups of CFEM domain containing proteins (Table 32). Homologous CFEM domain proteins may be involved in defense, pathogenicity, host-pathogen interactions or iron uptake, as is the rbt5 family of *Candida spp*.(Gaurav Bairwa et al. 2017) In *C. cinerea* certain mycelium expressed CFEM domain proteins were reported to be involved in defense(Plaza et al. 2014b). The detected CDE orthogroups comprise cell surface proteins with unknown functions, though they might be involved in defense, however, evidence for that is missing at the moment.

**Fasciclins** are adhesive cell surface proteins that facilitate the attachment of cells to various surfaces. The few fungal mentions include a report of a role in the adhesion of *Magnaporthe oryzae* to host surfaces(Liu et al. 2009) as well as a primordium-upregulated gene in *L. edodes* (*Le.flp1*)(Y et al. 2007). Recently, fasciclins were hypothesized to have been co-opted for fruiting body development independently in both the Agarico- and Pezizomycotina(Merényi et al. 2020b). We identified 2-8 fasciclin-encoding genes in the examined Agaricomycetes. All examined species have developmentally expressed fascicling genes, although these formed only small CDE orthogroups (Table 32). Several genes showed an upregulation in primordia relative to vegetative mycelium (as reported in *L. edodes*), suggesting that this family has widespread roles in mushroom formation, possibly by facilitating adhesion. We found that the sequence deposited by(Y et al. 2007) (Accession number: AB287432) as well as ascomycete homologs to which they analyzed are cupredoxins, not fasciclins, leaving doubts as to the identity of Le.flp1. Nevertheless, this does not undermine the observation that fasciclin encoding genes are widespread developmentally expressed genes in mushroom-forming fungi, probably with roles in fruiting body development.

**Intradiol ring cleavage dioxygenases** catalyze a ring-opening step in the degradation of aromatic compounds, and were hypothesized to be involved in xenobiotic detoxification or the utilization of aromatic compounds as carbon sources(Semana and Powlowski 2019). However, the physiological role of intradiol ring cleavage dioxygenases in fungi is not known. The only previous mention of a potential relationship to fruiting bodies reported developmental regulation in 4 out of 6 species(Krizsán et al. 2019). We identified 2-7 copies (28 in *A. ostoyae*) in the examined Agaricomycetes, with 1-3 developmentally expressed genes in most of the species that, however, did not form conserved orthogroups. Members of the family showed a significant upregulation in primordia relative to vegetative mycelium in *A. ampla, S. commune, C. cinerea, A. ostoyae* as well as a strong upregulation (>100fold) during late development of *C. neoformans*. Two genes show a very distinct upregulation in young fruiting body caps of *C. cinerea*. No developmentally expressed genes were found in *Ph. chrysosporium* and *M. kentingensis.* In summary, intradiol ring cleavage dioxygenases might be important genes during fruiting body development of several species; they might degrade aromatic compounds during fruiting, however, the identity of those compounds are unknown at the moment.

### 4.10. Defense

Agaricomycete fruiting bodies (mushrooms) are, due to the concentration of biomass, attractive targets of predators, parasites and pathogens. Accordingly, these structures are known to be subject to grazing by slugs, to infestations with larvae of phorid and sciarid flies and to infections by bacteria and molds(Wood et al. 2004; Krivosheina 2008; Lakkireddy et al. 2020; M et al. 2020a). Since mushrooms are the sexual reproduction organs and a considerable investment in terms of resources and energy, Agaricomycetes have evolved several strategies for the protection and defense of these structures(Künzler 2018). Such strategies include physical barriers(Varga et al. 2021) and chemical defense i.e. the production of molecules (toxins) impairing the growth, development and/or viability of antagonists(P 2008; FY and NP 2014; Stadler and Hoffmeister 2015).

Using saprobic Agaricomycete species, whose sexual cycles can be completed under laboratory conditions, it could be demonstrated that the production of these toxins occurs axenically i.e. in the absence of the antagonist(Plaza et al. 2014b; Tayyrov et al. 2019). This axenic toxin production is spatiotemporally regulated in that the toxin-encoding genes are expressed solely during fruiting body formation and in the fruiting body tissue. Some of these genes belong to the most highly upregulated genes of the entire genome during fruiting body formation. This expression pattern results in an efficient, constitutive protection of the fruiting body because at least some toxins are already in place in case the structure is attacked by an antagonist. It is unclear whether the chemical defense of the Agaricomycete fruiting body can be fortified by the induction of toxin production in response to environmental factors e.g. the presence of antagonists. Antagonist-specific induction of antibacterial and nematotoxic proteins upon challenge with bacteria and fungivorous nematodes, respectively, has recently been reported for the vegetative mycelium of C. cinerea(Kombrink et al. 2018; Tayyrov et al. 2019).

Toxins of Agaricomycete fruiting bodies include small molecules (secondary metabolites)(Kües et al. 2018), peptides (ribosomally or non-ribosomally synthesized) and proteins, all of which act by binding to and (in case of toxins with enzymatic activity) converting specific target molecules in the antagonists. In the following, we will summarize the current knowledge about the structure, function, biosynthesis, regulation and distribution of Agaricomycete fruiting body toxins. In accordance with the focus of this review on genomics, we restrict our summary to compounds where the biosynthetic genes have been identified. We also make references to available gene expression data, but it must be noted that most toxin-encoding genes evolve fast and thus they do not easily conform to orthology-based categorizations that are central to the rest of this paper.

**Small molecules** - The structurally related alkaloids ibotenic acid and muscimol of the fly agaric *Amanita muscaria* are two well-known examples of defensive secondary metabolites of Agaricomycete fruiting bodies(Michelot and Melendez-Howell 2003). The toxicity of these compounds toward flies gave rise to the name of the producer fungus. Their bioactivity is due to their structural similarity to the neurotransmitters glutamate and GABA and their binding to respective receptors in the brains of both invertebrates and vertebrates. The recent discovery of the biosynthetic gene cluster for the compounds revealed that their biosynthesis initiates from glutamate and makes it possible to identify additional *Amanita* species producing these compounds on a genetic basis(Obermaier and Müller 2020). Another psychotropic alkaloid produced by Agaricomycete fruiting bodies is psilocybin produced by several *Psilocybe* species(J et al. 2017; Torrens-Spence et al. 2018). This tryptophan derivative is an agonist of serotonergic receptors and is currently tested for treatment of depressions(Tullis 2021). The transcriptional analysis of the recently identified biosynthetic genes confirms its fruiting body-specific production(R et al. 2020). The blueing reaction of psilocybin-producing fruiting bodies upon injury has recently been demonstrated to be due to enzyme-mediated oxidative oligomerization of psilocybin suggesting that at least part of the defense function of this compound may be based on optical deterrence(Lenz et al. 2020). Wound-activation by injury (oxidation) has been reported for other secondary metabolites of Agaricomycete fruiting bodies(Stadler and Sterner 1998; P 2008) but the genetic basis of these compounds and reactions is often not known.

**Defense peptides** - The bicyclic octapeptide α-amanitin produced by some *Amanita*, *Galerina*, *Conocybe* and *Lepiota* species is probably the most famous toxin of Agaricomycete fruiting bodies(Walton 2018). The thermal stability and the very low lethal dose (0.1 mg/kg body weight) of the peptide have led to the common names ’death cap’ (*Amanita phalloides*) or ’destroying angel’ (*Amanita bisporigera*) of α-amanitin producers. Upon ingestion, the peptide is released from the fruiting body tissue and able to passively enter the intestinal epithelial cells where it binds and inactivates the essential enzyme RNA polymerase II in the nucleus(X et al. 2018). The fungus protects itself from the toxin by expressing an insensitive variant of the target enzyme(PA et al. 1978). In 2007, Walton and coworkers demonstrated that α-amanitin and its relative phallacidin are derived from ribosomally synthesized precursor peptides with common flanking sequences (MSDIN) by the action of a dual function prolyloligopeptidase-macrocyclase (POPB)(Hallen et al. 2007; H et al. 2012). Genome surveys of above Agaricomycete species revealed a large family of peptide precursor genes differing in the core peptide sequence and, thus, coding for α-amanitin- and phallacidin-related, but structurally different peptides, referred to as cycloamanides, whose function is unknown(Pulman et al. 2016; H et al. 2018; He et al. 2020; Zhou et al. 2021). Localization of POPB expression confirmed the fruiting body-specific expression of α-amanitin and phallacidin(Luo et al. 2010; CH et al. 2018). Above described secondary metabolites and peptides have received considerable attention, mainly due to their activity towards humans. During the last two decades of research on fungal defense, it has become evident, however, that a large part of the chemical defense of Agaricomycete fruiting bodies, in particular against invertebrate predators and parasites, is based on protein toxins(M et al. 2002b; J et al. 2016; Tayyrov et al. 2018). These toxins have received less attention because their activity is usually lost upon boiling and is specific for a particular group of invertebrates. In addition, based on their fruiting body-specific expression, some of these proteins have been implicated in fruiting body formation rather than defense (see the example of *C. cinerea* galectins CGL1 and CGL2 mentioned above), which has delayed their recognition and functional analysis as toxins(S et al. 2007a; A et al. 2019a). Most of these protein toxins are stored intracellularly and released upon ingestion of the fruiting body tissue by fungivores(Kunzler 2015; Künzler 2018). Once released, they interact with the digestive tract of the fungivore. Characterized classes of protein toxins from Agaricomycete fruiting bodies include lectins(Goldstein and Winter 2007; Bleuler-MartÍnez et al. 2011; A et al. 2013), protease inhibitors(S et al. 2015; Sabotič et al. 2019), pore-forming toxins (PFTs)(Anderluh and Maček 2003; JM et al. 2010; Panevska et al. 2020), ribotoxins(Tayyrov et al. 2019; Ragucci et al. 2021) and biotin-binding proteins(Y et al. 2009; S et al. 2012a).

**Defense related lectins** - Fruiting body lectins have long been used as tools in glycobiology without addressing their biological function(Y and H 2014). In the meantime, it has been established that these proteins bind to specific glycoepitopes on the epithelial cells of the digestive tract of the fungivore which can ultimately lead to disruption of the structure and function of this tissue(Stutz et al. 2015). The mechanism of lectin-mediated toxicity is not clear yet but, in case of lectins without additional functional domain (hololectins), the bioactivity appears to depend on the crosslinking of protein-bound glycoepitopes on the epithelial cell surface(S et al. 2017), whereas in case of lectins with additional functional domains (chimerolectins), the bioactivity appears to depend on binding to lipid-bound glycoepitopes and endocytosis(T et al. 2011; Juillot et al. 2016). Fruiting body lectins can adopt various folds and carbohydrate specificities amongst which the RicinB-like (β-trefoil) fold and fucoside-binding, respectively, are most common. A prototypic example for such a fruiting body lectin is the hololectin CCL2 (*C. cinerea* protein ID: 408852) from the model Agaricomycete *Coprinopsis cinerea* which is present in other Agaricomycete species, adopts a RicinB-like fold, forms a dimer, binds to a specific fucoside present on nematode N-glycan cores and is toxic towards bacterivorous and fungivorous nematodes(Schubert et al. 2012; S et al. 2017; Tayyrov et al. 2018). A summary of diverse entomo- and nematotoxic lectin have been described from fruiting bodies has been given by Sabotic et al (J et al. 2016). Our data confirm previous reports on the upregulation (>150-fold) of CCL1 (*C. cinerea* protein ID: 456849) and CCL2 (*C. cinerea* protein ID: 408852) in primordia (Table 33).

**Table 33.** Summary of developmental expression dynamics of CDE orthogroups of defense-related genes across 12 species. Protein ID of a representative protein is given follows by the number of species in which the orthogroup is developmentally regulated at fold change 2 and 4 (FC>2 and FC>4, respectively). Putative function and ortholog in *S. cerevisiae* (if any) are also given. Abbreviations: 0-gene absent, 1-gene present but not developmentally regulated, 2 - developmentally regulated at fold change >2, 4- developmentally regulated at fold change >4. Species names are abbreviated as: Cc – *C. cinerea*, Aa – *A. ampla*, Ab – *A. bisporus*, Ao – *A. ostoyae*, Ca – *C. aegerita*, Lb – *L. bicolor*, Le – *L. edodes*, Lt – *L. tigrinus*, Mk – *M. kentingensis*, Pc – *Ph. chrysosporium*, Po – *P. ostreatus*, Sc – *S. commune*.

| ID | Fold change |  | Putative function | S.<br>*cerevisiae*<br>ortholog | Species |  |  |  |  |  |  |  |  |  |  |  |
| --- | --- | --- | --- | --- | --- | --- | --- | --- | --- | --- | --- | --- | --- | --- | --- | --- |
|  | >2 | >4 |  |  | Cc | Aa | Ab | Ao | Ca | Lb | Le | Lt | Mk | Pc | Po | Sc |
| 185637 | 9 | 5 | CFEM domain containing protein | – | 4 | 1 | 4 | 4 | 2 | 2 | 2 | 2 | 4 | 1 | 4 | 1 |
| 439390 | 9 | 6 | CFEM domain containing protein | – | 4 | 4 | 4 | 4 | 2 | 2 | 2 | 4 | 0 | 1 | 1 | 4 |
| 485770 | 7 | 5 | RicinB lectin protein | – | 4 | 2 | 4 | 0 | 0 | 1 | 4 | 2 | 0 | 0 | 4 | 4 |

The RicinB-like (beta-trefoil) fold is also adopted by protease-inhibitors isolated from Agaricomycete fruiting bodies(J et al. 2016). These proteins were shown to inhibit serine and cysteine proteases of various insect larvae. A prototypic example for this type of fruiting body toxin is Clitocypin, an insecticidal cysteine-protease inhibitor isolated from the Agaricomycete *Clitocybe nebularis*(J et al. 2011; Šmid et al. 2015). Of the examined species, the genomes of *C. aegerita* and *L. bicolor* encode 1 and 20 putative clitocypins, respectively. Our data identified a putatively defense-related RicinB protein orthogroup (represented by *C. cinerea* 485770)(Table 33). Another example is the six-bladed beta-propeller lectin Tectonin2 from L. bicolor whose production is induced in fruiting bodies and ectomycorrhizae(Martin et al. 2008). This lectin forms tetramers harboring 24 binding sites for O-methylated glycoepitopes and exhibits toxicity towards nematodes and agglutination of bacteria displaying respective glycoepitopes on their cell surfaces(Wohlschlager et al. 2014; Sommer et al. 2018). We detected tectonin homologs in the genomes of *L. bicolor* and *C. aegerita*.

**Pore-forming toxins** - A common and probably very ancient class of protein toxins of Agaricomycete fruiting bodies are pore-forming toxins (PFTs)(Peraro and van der Goot 2015). These toxins are divided into α- and β-PFTs depending on whether the predominant secondary structure motif is an α-helix or β-sheet. Within the producing cell, these proteins are usually monomeric and soluble, but upon release and binding to a target cell membrane, they undergo a dramatic conformational change. This change in conformation leads to oligomerization and insertion of the protein into the membrane. The result of this complex process is a proteinaceous pore that leads to lysis of the target cell. Among PFTs occurring in Agaricomycete fruiting bodies, members of the aerolysin and <u>M</u>embrane <u>A</u>ttack <u>C</u>omplex <u>P</u>er<u>F</u>orin/<u>C</u>holesterol <u>D</u>ependent <u>C</u>ytolysin (MACPF/CDC) β-PFT superfamilies have been characterized most extensively(Reboul et al. 2016; Cirauqui et al. 2017). Examples for aerolysins are the hemolytic *Laetiporus sulphureus* lectin (LSL)(JM et al. 2010), a homologous and recently identified nematotoxic protein, CC1G_11805 (*C. cinerea* protein ID: 546868) and its two paralogs, from *C. cinerea*(Plaza et al. 2014b) and the hemolytic flammutoxin from *F. velutipes*(T et al. 2004b). Pri1, a primordium-expressed gene in *C. aegerita* (Agrae_CAA7262647) also belongs to pore-forming toxins(Fernandez Espinar and Labarère 1997). The MACPF/CDC superfamily is represented by the insecticidal pleurotolysin complex, consisting of PlyA and PlyB, from the edible mushroom *P. ostreatus*(K et al. 2014; M et al. 2015b; N et al. 2015; A et al. 2019a). The main representatives of the two β-PFT superfamilies differ in the specificity determinant of the target membrane and the dimensions of the formed membrane pore. Whereas LSL likely binds to lactose-containing glycoepitopes (as part of glycolipids or GPI-anchored glycoproteins) on the target membrane via an N-terminal RicinB-type(β-trefoil) lectin domain and forms homomeric transmembrane β-barrels (pores) of seven or eight protomers with an inner diameter of ∼4 nm(I et al. 2011; Szczesny et al. 2011), pleurotolysin PlyA binds to sphingophospholipids (sphingomyelins) of cholesterol-rich membranes and directs PlyB to the formation of transmembrane β-barrels (pores) consisting of 13 heteromeric protomers and an inner diameter of ∼ 8 nm(Novak et al. 2020).

**Ribotoxins** are ribonucleases cleaving a single phosphodiester bond in the universally conserved sarcin/ricin loop (SRL) of 23S or 28S rRNA in intact prokaryotic or eukaryotic ribosomes, respectively(M et al. 2017b). Upon this action, the ribosome is unable to interact with translation elongation factors and translation is stalled. The SRL is named after α-sarcin, one of the best characterized ribotoxins secreted by the ascomycetous mold *Aspergillus giganteus*(M et al. 2017b). Other fungal ribotoxins have been isolated from ascomycetous entomopathogens and are generally highly active against insects(A et al. 2009; M et al. 2017a). Recently, the first basidiomycetous ribotoxin, ageritin (Agt1), from fruiting bodies of the edible mushroom *C. aegerita* was isolated(Landi et al. 2017). Heterologous expression of the respective cDNA in *Escherichia coli* and phylogenetic analysis showed that the protein is widespread but patchy among Agaricomycetes and displays entomotoxic activity similar to its ascomycetous counterparts(A et al. 2019b). In contrast to its ascomycetous counterparts, however, ageritin does not contain a canonical signal peptide for secretion suggesting that the protein, as most protein toxins of Agaricomycete fruiting bodies, is stored intracellularly(Baglivo et al. 2020). In order to avoid damage to the ribosomes of the producing fungal cell, the protein might be sequestered into a specific subcellular compartment which might involve N-terminal truncation of the protein(Ragucci et al. 2021). The agt1 gene (*C. aegerita* CAA7259527) was induced considerably in primordia relative to vegetative mycelium in *C. aegerita*, based on data produced by Orban et al (Orban et al. 2021).

**Tamavadins** - Finally, defense proteins of some Agaricomycete fruiting bodies, similar to animals and bacteria, include proteins that are able to avidly bind and thus sequester biotin at equimolar concentrations(Michael Green 1990). The prototype of these proteins, avidin, is thought to protect the eggs of birds from bacterial infections, but the protein and its relatives also exhibit broad and strong insecticidal activity(JT et al. 2010). A family of avidin-related biotin-binding proteins, tamavidin (called Tam1 and Tam2), was isolated from the edible mushroom *Pleurotus cornucopiae* (Tamogitake mushroom) and shown to possess very strong entomotoxic and nematotoxic activity(Y et al. 2009; S et al. 2012a). Amongst the fruiting body protein toxins tested against two fungivorous nematode species, tamavidin was the most effective one(Tayyrov et al. 2018). It is unclear how the producing cells shield their own biotin against this cytoplasmic protein.

Taken together, this data shows that the chemical armory of Agaricomycete fruiting bodies is very diverse, in terms of compound classes, molecular targets and antagonist specificity of the toxins, but three common themes can be recognized: (1) Toxin-encoding genes are phylogenetically patchy, with each family present in only a subset of species (2) protein toxins provide a large part of the chemical defense of these structures; and (3) intracellularly stored toxins directed against nematodes and arthropods are predominant. Latter finding may reflect the use of Agaricomycete fruiting bodies as habitat by these organisms(Ruess and Lussenhop 2005; Krivosheina 2008). It has even been postulated that some mycophagous Drosophila flies exploit fruiting body nematotoxins to purge themselves from insect-parasitic nematodes(JA et al. 2017).

### 4.11. Secondary metabolism and natural product biosynthesis

Secondary metabolites are one of the most important classes of molecules in chemical warfare and developmental signaling in fungi(Spiteller 2015; Keller 2019; Gressler et al. 2021). Based on recent genome surveys it appears that the fungal secondary metabolome is extremely diverse and is a largely untapped resource of natural products and bioactive molecules(Keller 2019). This may be particularly true for the Agaricomycetes, despite several lines of evidence for a large diversity of secondary metabolites produced by Agaricomycetes(Spiteller 2015; Gressler et al. 2021). From the perspective of fruiting body formation, it is important to note that secondary metabolites can serve defense or repellent purposes in fruiting bodies (see chapter on Defense) but can also influence developmental processes. For example, the deletion of a polyketide synthase gene resulted in malformed perithecia of *Sordaria macrospora*(D and M 2014). The lack of coprinoferrin production in *C. cinerea* resulted in the loss of fruiting body formation(Tsunematsu et al. 2019).

Due to differences in basic genome organization between Asco- and Basidiomycota, the latter possess fewer secondary metabolism gene clusters, yet, according to a recent study some developmentally regulated genes formed ‘clusters’ in the genomes of 8 species and some of these ‘clusters’ overlapped with predicted biosynthetic gene clusters that might encode secondary metabolism genes(Merényi et al. 2021).

The number of functionally characterized secondary metabolism genes and linked natural products of Agaricomycetes is rapidly increasing. For example, *ArmB* and orsellinic acid in *Armillaria* spp(Lackner et al. 2013), *LpaA* and laetiporic acid in *Laetiporus* sp(Seibold et al. 2020), or *St.pks1* and strobilurin in *Strobilurus tenacellus*(Nofiani et al. 2018) as well as numerous terpene synthases have been circumscribed in the past decade (reviewed in more detail by(Gressler et al. 2021)). Strobilurin derivatives are available commercially and are the world’s most sold fungicides(DW et al. 2002). Siderophore-type secondary metabolites, coprinoferrin, ferrichrome A and basidioferrin in *C. cinerea, Omphalotus olearius* and *Ceriporiopsis subvermispora*, respectively, and their putative synthesizing genes were also identified. In *C. cinerea cfp1* (*C. cinerea* protein ID: *354233*) was reported to be responsible for coprinoferrin production(Tsunematsu et al. 2019), whereas in *O. olearius* a cluster composed of *fso1, omo1* and *ato1*(K et al. 2005). Coprinoferrin proved necessary for fruiting body development, as the cfp1 mutant did not, only in the presence of exogenously administered coprinoferrin produce fruiting bodies. Contrary to expectations based on Ascomycota knowledge, deleting a homolog (*C. cinerea* protein ID: 357031) of *A. nidulans LaeA* did not compromise coprinoferrin production(Tsunematsu et al. 2019). Despite recent advances in understanding Agaricomycetes secondary metabolism, given the number of putative secondary metabolism genes encoded by fungal genomes, we are likely hardly scratching the surface of natural product diversity in mushroom-forming fungi.

Since many secondary metabolites specifically occur in fruiting bodies of Agaricomycetes, it is expected that their biosynthetic gene clusters may be developmentally regulated. However, secondary metabolism related genes evolve fast, which does not make them easily detectable in the orthology-based framework we use here. Nevertheless, we identified several secondary metabolism related genes that showed shared and conserved developmental expression profiles in the examined Agaricomycetes, although the natural products they produce are unknown in most cases.

**Polyketide synthases** (PKS) are large (often >1500 amino acid) multi-domain enzymes that produce diverse secondary metabolites. They are widely distributed in fungi and are subject to intense research, mostly in the Ascomycota. Polyketide synthases and the few examples of known natural products of the Basidiomycota have recently been reviewed(Gressler et al. 2021). Basidiomycete PKSs mostly belong to type 1, which is divided into highly-, partially- and non-reducing PKSs. Hybrid nonribosomal peptide synthase-PKS genes have been shown to underlie fungal bioluminescence(Ke et al. 2020a; Gressler et al. 2021).

We identified orthologs of previously described siderophore biosynthesis related genes in the examined Agaricomycetes. Members of the biosynthetic cluster described from *O. olearius*(K et al. 2005) seem to be conserved, with fso1 being orthologous to *C. cinerea* cfp1 (*C. cinerea* protein ID: 354233), whereas L-ornithine N(5)-monooxygenase omo1 and the N-acyltransferase ato1 seem single-copy in Agaricomycetes genomes (represented by *C. cinerea* 492586 and 161175). These genes were not developmentally regulated and had highest expression in vegetative mycelia. We further identified a single CDE orthogroup of polyketide synthases (represented by *C. cinerea* 446258, Table 34), which was developmentally regulated in 11/8 species (FC>2/4). These genes were stipe upregulated (albeit weakly) in most species (*C. cinerea, P. ostreatus, M. kentingensis, A. ostoyae*), although their expression also increased in mature caps of *P. ostreatus, A. ostoyae, M. kentingensis* relative to younger cap stages and in mature fruiting bodies of *C. aegerita*, *A. ampla* and *S. commune* relative to earlier stages. The *C. cinerea* gene shows peak expression in meiotic gill tissues, whereas in *P. ostreatus* and *A. ostoyae*, where gills and caps were sampled separately like in *C. cinerea*, highest expression was observed in caps but not in gills.

**Table 34.** Summary of developmental expression dynamics of CDE orthogroups of terpene synthase genes across 12 species. Protein ID of a representative protein is given follows by the number of species in which the orthogroup is developmentally regulated at fold change 2 and 4 (FC>2 and FC>4, respectively). Putative function and ortholog in *S. cerevisiae* (if any) are also given. Abbreviations: 0-gene absent, 1-gene present but not developmentally regulated, 2 - developmentally regulated at fold change >2, 4- developmentally regulated at fold change >4. Species names are abbreviated as: Cc – *C. cinerea*, Aa – *A. ampla*, Ab – *A. bisporus*, Ao – *A. ostoyae*, Ca – *C. aegerita*, Lb – *L. bicolor*, Le – *L. edodes*, Lt – *L. tigrinus*, Mk – *M. kentingensis*, Pc – *Ph. chrysosporium*, Po – *P. ostreatus*, Sc – *S. commune*.

| ID | Fold change |  | Putative function | S.<br>*cerevisiae*<br>ortholog | Species |  |  |  |  |  |  |  |  |  |  |  |
| --- | --- | --- | --- | --- | --- | --- | --- | --- | --- | --- | --- | --- | --- | --- | --- | --- |
|  | >2 | >4 |  |  | Cc | Aa | Ab | Ao | Ca | Lb | Le | Lt | Mk | Pc | Po | Sc |
| 361541 | 8 | 5 | Phenylalanine ammonia-lyase | – | 4 | 1 | 4 | 2 | 4 | 4 | 0 | 0 | 2 | 0 | 4 | 2 |
| 386671 | 11 | 9 | Phenylalanine ammonia-lyase | – | 4 | 2 | 4 | 4 | 4 | 4 | 2 | 4 | 4 | 4 | 4 | 1 |
| 446258 | 11 | 8 | Putative polyketide synthase | YER061C | 4 | 4 | 4 | 4 | 4 | 1 | 2 | 2 | 2 | 4 | 4 | 4 |
| 394772 | 4 | 2 | Putative terpene synthase | – | 4 | 0 | 1 | 4 | 1 | 2 | 1 | 1 | 1 | 1 | 2 | 0 |
| 443805 | 3 | 2 | Putative terpene synthase | – | 4 | 0 | 2 | 0 | 0 | 0 | 0 | 1 | 0 | 0 | 4 | 0 |
| 495900 | 1 | 1 | Putative terpene synthase | – | 1 | 0 | 0 | 0 | 4 | 0 | 0 | 1 | 1 | 0 | 1 | 0 |
| 124670 | 5 | 2 | Putative terpene synthase | – | 4 | 0 | 1 | 4 | 1 | 2 | 0 | 2 | 1 | 0 | 2 | 0 |
| 30510 | 9 | 8 | Putative terpene synthase | – | 4 | 4 | 4 | 4 | 4 | 4 | 4 | 2 | 0 | 1 | 4 | 1 |
| 469738 | 10 | 8 | Putative terpene synthase | – | 1 | 1 | 4 | 4 | 4 | 4 | 4 | 2 | 4 | 4 | 4 | 2 |
| 201523 | 1 | 1 | Putative terpene synthase | – | 4 | 0 | 0 | 0 | 0 | 0 | 0 | 0 | 0 | 0 | 0 | 0 |

**Terpene synthases** have been a focus of secondary metabolism studies in mushroom-forming fungi. They encode highly promiscuous enzymes that can generate a large diversity of metabolites(Slot and Gluck-Thaler 2019) and thus provide the basis for fast adaptive evolution. Slot and Gluck-Thaler showed that morphologically complex fungi harbor more terpene synthases than simpler counterparts(Slot and Gluck-Thaler 2019). Similarly, terpene synthases were found to be strongly overrepresented in the genomes of Agaricomycetes compared to other fungi (FDR corrected p-value < 10^-125^)(Krizsán et al. 2019). Twelve sesquiterpene synthases (Fla1-12) were identified in *F. velutipes*(Tao et al. 2016), eleven in *C. aegerita* (Agr1-Agr11)(Zhang et al. 2020, 2021a) and in *Omphalotus olearius* (Omp1-10)(GT et al. 2012) and five (Cop1-Cop6) in *C. cinerea*(Agger et al. 2009), with information on the potential metabolite products in many cases (e.g. viridiflorene/viridiflorol, Δ^6-^protoilludene, γ-cadinene, germacrene).

Agger et al identified six genes, Cop1 to Cop6, related to terpene biosynthesis in *C. cinerea*(Agger et al. 2009). Of the genes reported by Agger et al, orthologs of Cop2 and Cop4 appear interesting from the perspective of mushroom development. Orthologs of Cop1 show an upregulation in primordia relative to vegetative mycelium in *C. aegerita, A. ampla* (only FC>2), *L. bicolor, Ph. chrysosporium* and were cap-upregulated in *A. ostoyae, P. ostreatus, L. edodes* and *M. kentingensis*. On the other hand, orthologs of Cop4 (*C. cinerea* protein ID: 30510) were FB-init in *C. cinerea, C. aegerita, A. ampla, L. bicolor, L. edodes* and *P. ostreatus* (Table 34). Cop2 (*C. cinerea* protein ID: 443805) is only present in *C. cinerea, A. bisporus* and *P. ostreatus* and is upregulated in primordia relative to vegetative mycelium in all species, suggesting it functions during fruiting body initiation. Products of these genes were reported to as δ-cadinene and germacrene A, but the functions of these natural products remain unknown.

We found 13-28 (63 in *A. ostoyae*) putative terpene synthases in the examined Agaricomycetes genomes. Of these, several were developmentally regulated, showing one of two most typical expression profiles: (i) highest expression in vegetative mycelium or (ii) a strong induction in primordia and other fruiting stages relative to vegetative mycelium. The latter expression profile is reminiscent of that of typical defense genes (e.g. pore-forming toxins, see above), but it is compatible with several other putative functions as well. Terpene synthases formed three CDE orthogroups (Table 34), two of these correspond to *C. cinerea* Cop1 (*C. cinerea* protein ID: 469738) and Cop4 (*C. cinerea* protein ID: 30510, contains *C. aegerita* Agr4 also), discussed above. The third CDE orthogroup (represented by *C. cinerea* 488324) is developmentally regulated in 8/6 species, but shows mixed expression profiles.

Taken together, terpene synthases might participate in important fruiting body functions, including events around the initiation of primordium development, yet their exact functions, whether the production of defense/repellent metabolites and/or compounds for communication, remain unknown for the moment.

**Phenylalanine ammonia-lyases** - We detected two highly conserved CDE orthogroups of phenylalanine ammonia-lyases (PAL), one of which (*C. cinerea* protein ID: 386671, ortholog of pal1 of *Pleurotus*) is developmentally regulated in 11/9 species, the other (*C. cinerea* protein ID: 361541, ortholog of pal2 of *Pleurotus*(Hou et al. 2019)) is in 8/5 (out of 9 species in which present) (Table 34). PAL enzymes are best known from plants, where they are part of the phenylpropanoid pathway, which is involved in the production of diverse secondary metabolites, associated with defense, lignin and flavonoid biosynthesis(Dixon et al. 2002). In fungi, phenylalanine ammonia-lyases are discussed mostly in the context of catabolism of phenylalanine to cinnamic acid(Hyun et al. 2011), although some reports of the production of diverse compounds have been published(Hyun et al. 2011). Recently, an association with fruiting body formation has been reported in *F. velutipes*(Yun et al. 2015) and *P. ostreatus* (Hou et al. 2019).

Our analyses revealed two PAL-encoding genes in most of the examined Agaricomycetes (up to 4 in *A. ostoyae*). Expression profiles of genes within the two CDE orthogroups were not conserved, although several genes showed an upregulation in primordia (e.g. both genes in *C. cinerea*). At the moment the physiological role of PALs is unknown, to our best knowledge, in any fungus. They could be involved in the production of defense-related metabolites, as in plants, or other fruiting body related functionalities.

### 4.12. Cytoskeleton

A plastic cytoskeleton is a key factor for eukaryotic cell shape regulation and has been named one of the prerequisites for diversification of size, form and function in multicellular eukaryotes(Knoll 2011). It is involved in the regulation of diverse morphogenetic processes in fungi, including hyphal growth, conidiation, biofilm formation, dimorphic switching, to name a few(Riquelme et al. 2018). In line with this, although at a somewhat limited scale, evidence is available for the role of cytoskeleton rearrangement in fruiting body development. Evidence from Gene Ontology analyses pointed at differential cytoskeletal gene expression in *P. microcarpus*(Pereira et al. 2017). Microscopy observations provided evidence for a key role of cytoskeletal elements during the development of mushroom-forming fungi. For example, longitudinally oriented microtubules were observed in the sterigmata during spore formation in *C. cinerea*, along which Golgi vesicles carry carbohydrates to the developing spore and spore wall(Kües and Liu 2000).

In terms of expression levels, we found only limited dynamics in cytoskeletal proteins, likely reflecting the need for continuous supply of its components for the cell. Cheng et al reported an upregulation of tubulin and tubulin-specific chaperone genes in primordium stages of *C. cinerea*(Cheng et al. 2013a) using 5’-SAGE technology. Although newer data did not confirm these observations(Muraguchi et al. 2015; Krizsán et al. 2019), both recent RNA-Seq studies reported upregulation of these genes in stipes of *C. cinerea*.

Following these lines of evidence, we annotated the main cytoskeletal protein groups in *C. cinerea* based on homology and domain composition and checked their orthologs’ expression patterns across mushroom-forming fungi. We grouped 67 cytoskeletal and associated protein coding genes into the following categories (Table 35): actin, actin nucleation, actin filament organization, actin-based motor protein, microtubule, microtubule heterodimerization, microtubule-based motor protein, regulation of microtubule-based motor proteins. Copy numbers in these categories agreed with previous annotations, including 2 copies of formins per species and a tubulin gene duplication in the common ancestor of Basidiomycota(Zhao et al. 2014). Of cytoskeletal genes, only kinesin genes and the Dynactin complex showed consistent expression dynamics. In both cases, expression peaks were observed in young fruiting body gills and stipes in *C. cinerea* (Fig. 14, Supplementary Fig. 10).

**Fig. 14.**
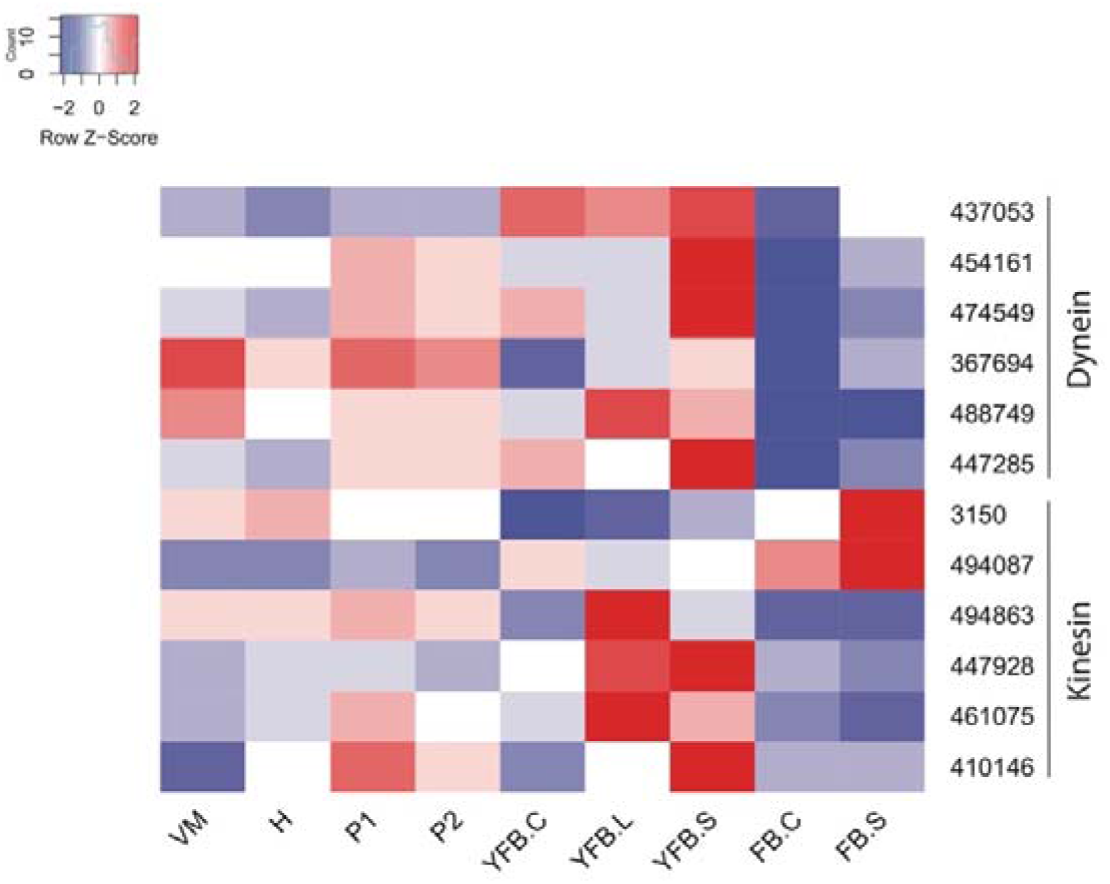
Expression heatmap of kinesin genes and members of the Dynactin complex in *C. cinerea*. Genes are denoted by Protein IDs. Blue and red colors represent low and high expression, respectively. Developmental stages are abbreviated as follows: VM – vegetative mycelium, H – hyphal knot, P1 – stage 1 primordium, P2 – stage 2 primordium, YFB.C – young fruiting body cap, YFB.L – young fruiting body gills, YFB.S – young fruiting body stipe, FB.C – mature fruiting body cap (including gills), FB.S – mature fruiting body stipe.

**Table 35.**
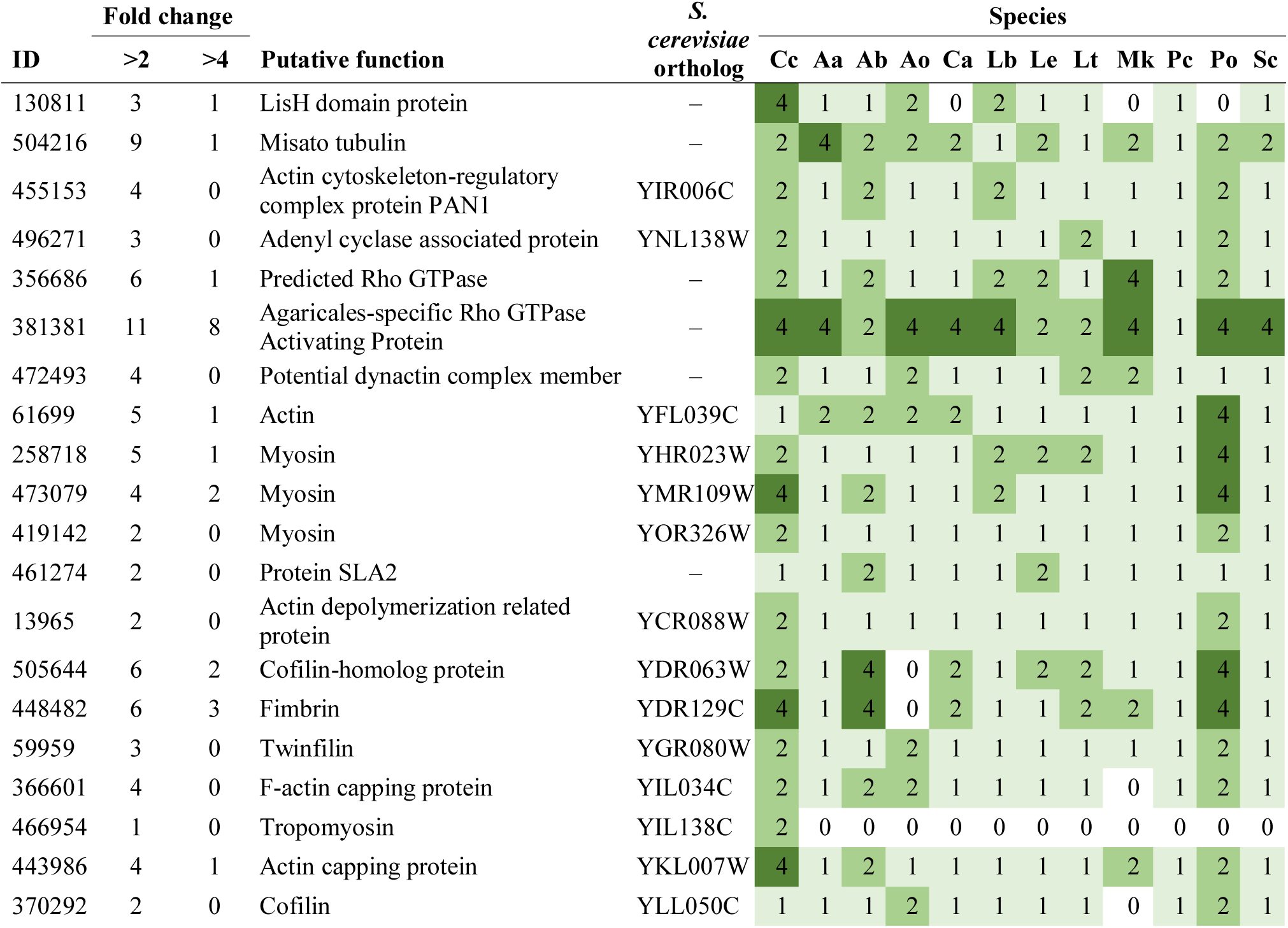
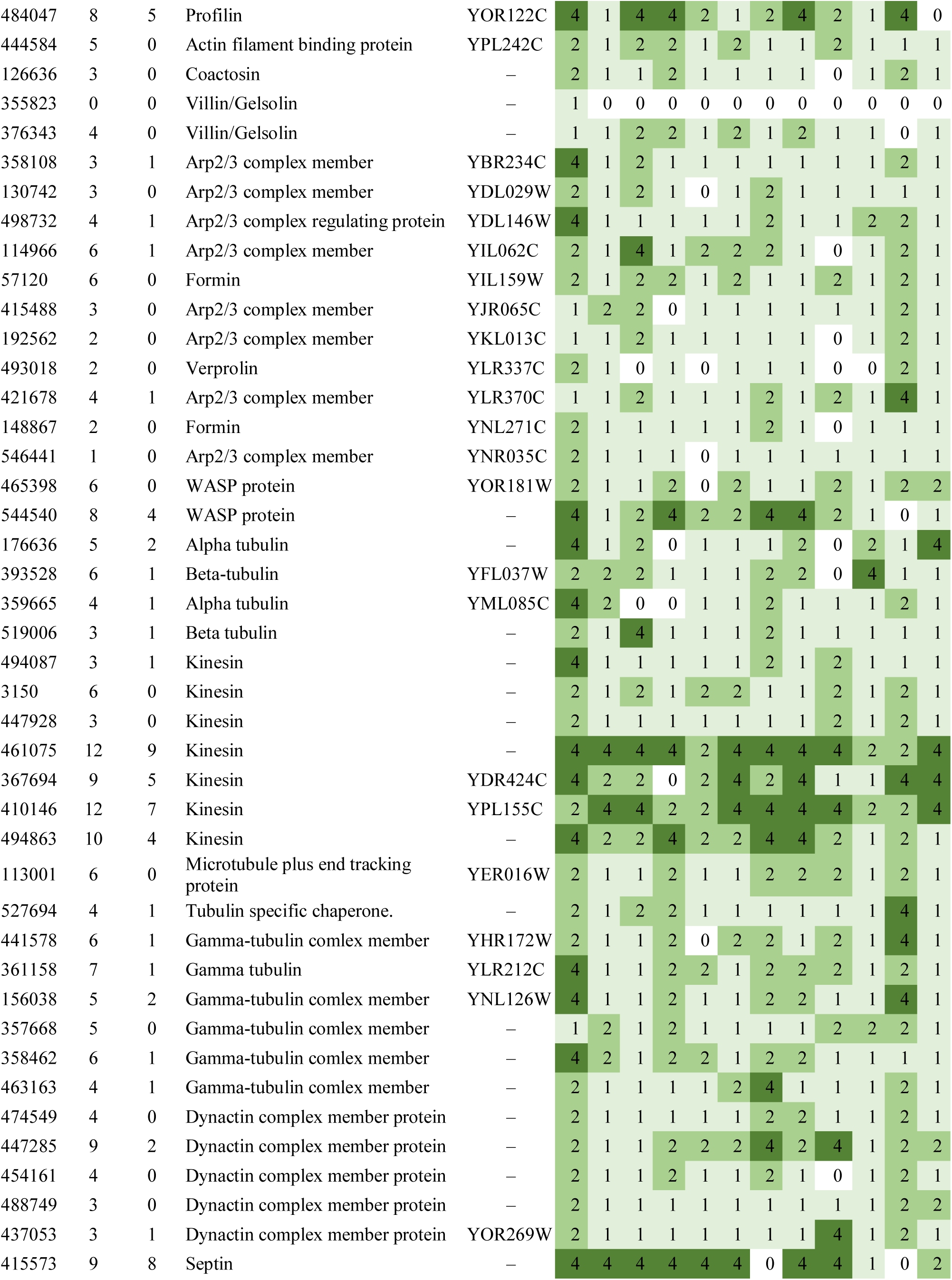
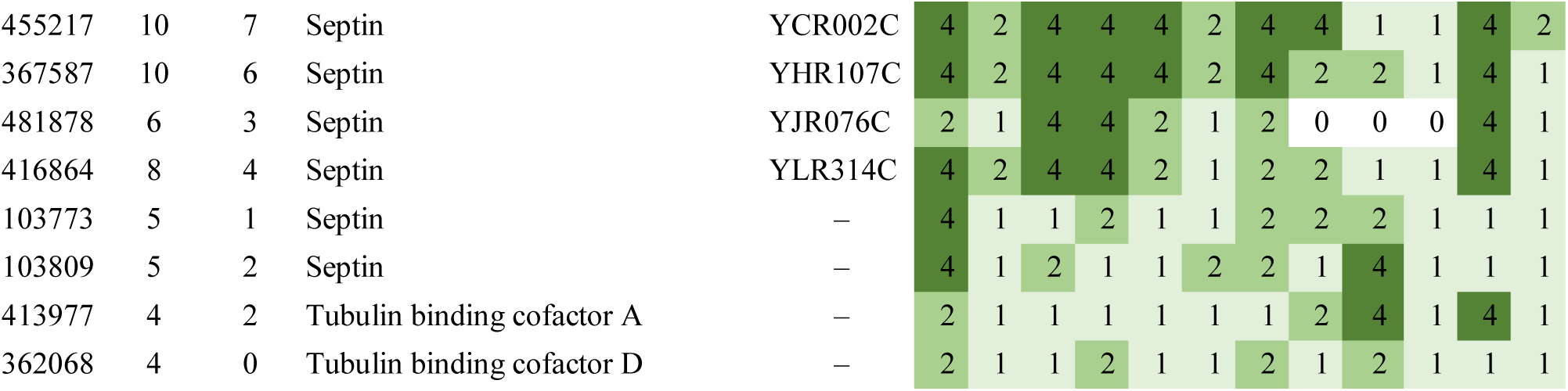
Summary of developmental expression dynamics of CDE orthogroups of cytoskeleton- related genes across 12 species. Protein ID of a representative protein is given follows by the number of species in which the orthogroup is developmentally regulated at fold change 2 and 4 (FC>2 and FC>4, respectively). Putative function and ortholog in *S. cerevisiae* (if any) are also given. Abbreviations: 0-gene absent, 1-gene present but not developmentally regulated, 2 - developmentally regulated at fold change >2, 4- developmentally regulated at fold change >4. Species names are abbreviated as: Cc – *C. cinerea*, Aa – *A. ampla*, Ab – *A. bisporus*, Ao – *A. ostoyae*, Ca – *C. aegerita*, Lb – *L. bicolor*, Le – *L. edodes*, Lt – *L. tigrinus*, Mk – *M. kentingensis*, Pc – *Ph. chrysosporium*, Po – *P. ostreatus*, Sc – *S. commune*.

Tubulin gene expression (alpha1-, alpha2-, beta-tubulin and tubulin chaperone) was previously reported to be regulated during early fruiting events of *C. cinerea*, with upregulation in stage 1 primordia based on 5’-SAGE data(Cheng et al. 2013a). Our data on the corresponding genes revealed that the three of these, alpha1- (*C. cinerea* protein ID: 176636), beta-tubulin (*C. cinerea* protein ID: 393528) and tubulin chaperone (*C. cinerea* protein ID: 413977) had constant expression in primordia, but show expression peak in young fruiting body stipes, whereas alpha2-tubulin (*C. cinerea* protein ID: 359665) was upregulated in young fruiting body caps. These expression patterns were not consistent across species (Table 35) and possibly reflect the fast stipe elongation of *C. cinerea*.

**Septins** - Septins form a highly conserved family of proteins involved in morphogenesis, development and in generating cell asymmetries in fungi and in eukaryotes in general(Gladfelter 2006; Khan et al. 2015). They have been shown to be responsible for morphogenesis in a number of model fungi, such as bud emergence in *S. cerevisiae*, septum formation in hyphae or appressorium formation in *Magnaporthe oryzae*, among others(Gladfelter 2006). Their broad involvement in morphogenesis and limited evidence from fruiting body forming fungi suggest that septins have important, although to date underappreciated, roles in fruiting body development. For example, they have been reported in stipe elongation in *C. cinerea*(Shioya et al. 2013).

We annotated septins based on their domain composition: the combination of two IPR domain terms (IPR030379, IPR016491) are diagnostic for fungal septins. Most mushroom-forming fungi have 7 septin genes (*C. cinerea* protein ID: 367587, 416864 [*cdc3*, = *A. nidulans* AspE, =AspB], 103773, 415573 [top hit of *A. nidulans* AspE], 103809, 455217, 481878), with some having only 6 while a few having 8, which is similar to the septin repertoires seen in filamentous fungi and yeasts(Douglas et al. 2005).

Septins are highly variable genes, and did not form conserved orthogroups, but nevertheless maintain some functional conservation in mushroom-forming fungi. Two orthogroups of septins showed consistent upregulation at the initiation of fruiting body development, in the very first primordium stages in all species in which they are present(Krizsán et al. 2019; Merényi et al. 2021). One of these is the top hit of AspE of *A. nidulans*, which is involved in conidiation and spore production(Hernández-Rodríguez et al. 2014). Septin expression seems to be highly induced in stipes in *C. cinerea, M. kentingensis, A. ostoyae*, but following a different expression tendency in *P. ostreatus* (mostly in primordia) and *A. bisporus* (mostly in young fruiting bodies). The stipe induced septins included *Cc.cdc3* of *C. cinerea*, which has been shown to be involved in stipe elongation(Shioya et al. 2013). The ortholog of *C. cinerea cdc3* was also mostly stipe-induced in *A. bisporus*(卢园萍 et al. 2019). Muraguchi et al reported a downregulation of septin expression from the 4_0hrPri primordia (approximately coinciding with stage 2 primordia of Krizsan et al and Kues 2000(Kües 2000; Krizsán et al. 2019)) to the 5_12hrPri stage of *C. cinerea*(Muraguchi et al. 2015).

The limited diversity of septins raises the question how they can fulfill the diverse roles they have in fungal morphogenesis. Although most septins had peak expression in stipes of mushroom-forming fungi (except in certain species) their expression level was dynamically changing in other tissues and from one developmental stage to the other as well, suggesting that there is still a long way to uncover their roles in fruiting body development. Understanding their protein-protein interaction networks is a promising future avenue of research.

### 4.13. Ubiquitin-proteasome system

#### 4.13.1. Cop9 signalosome

The Cop9 signalosome is a protein complex conserved across all eukaryotes, which catalyzes the deactivation of cullin-RING ubiquitin ligase (CRL) complexes by removing the Nedd8 protein from the cullin subunit of the complex (deneddylation reaction). As such, it is an important component of various signal transduction pathways and is involved primarily in multicellular developmental processes. In fungi, it was found dispensable for vegetative growth, but necessary for sexual fruiting body formation(Busch et al. 2003; Braus et al. 2010), secondary metabolism and it was reported to interact with the Velvet complex(Kress et al. 2012; Meister et al. 2019). Recognition of the COP9 signalosome (CSN) as an important player in fruiting body formation started with genetic screens that identified CsnE and CsnD, which displayed identical defects in fruiting body initiation in *A. nidulans*(Busch et al. 2003, 2007). The *A. nidulans* Cop9 complex represents an 8-subunit, stereotypical complex, whereas that of other model fungi (*N. crassa, S. cerevisiae* or *S. pombe*) possess fewer subunits and are not typical(Braus et al. 2010). Agaricomycetes genomes appear to encode a typical 8-subunit CSN (Table 36). Neither we nor previous studies found significant expression dynamics of any subunit of the CSN in mushroom-forming fungi. It is not known whether the complex is involved in fruiting body development as it is in Ascomycota, nevertheless, it will be an interesting target for future experiments, especially because it is involved in light regulation in *N. crassa* and can interact with the Velvet complex, two key factors that regulate Basidiomycota development also.

**Table 36.** Summary of developmental expression dynamics of CDE orthogroups of Cop9 signalosome genes across 12 species. Protein ID of a representative protein is given follows by the number of species in which the orthogroup is developmentally regulated at fold change 2 and 4 (FC>2 and FC>4, respectively). Putative function and ortholog in *S. cerevisiae* (if any) are also given. Abbreviations: 0-gene absent, 1-gene present but not developmentally regulated, 2 - developmentally regulated at fold change >2, 4- developmentally regulated at fold change >4. Species names are abbreviated as: Cc – *C. cinerea*, Aa – *A. ampla*, Ab – *A. bisporus*, Ao – *A. ostoyae*, Ca – *C. aegerita*, Lb – *L. bicolor*, Le – *L. edodes*, Lt – *L. tigrinus*, Mk – *M. kentingensis*, Pc – *Ph. chrysosporium*, Po – *P. ostreatus*, Sc – *S. commune*.

| ID | Fold change |  | Putative function | S.<br>*cerevisiae*<br>ortholog | Species |  |  |  |  |  |  |  |  |  |  |  |
| --- | --- | --- | --- | --- | --- | --- | --- | --- | --- | --- | --- | --- | --- | --- | --- | --- |
|  | >2 | >4 |  |  | Cc | Aa | Ab | Ao | Ca | Lb | Le | Lt | Mk | Pc | Po | Sc |
| 357329 | 9 | 5 | Proteasome component | – | 4 | 2 | 4 | 2 | 1 | 2 | 4 | 4 | 1 | 1 | 4 | 2 |
| 502637 | 6 | 0 | Defective-in-cullin neddylation protein | – | 1 | 0 | 1 | 2 | 2 | 2 | 1 | 1 | 2 | 1 | 2 | 2 |
| 496080 | 4 | 2 | Nedd8-like ubiquitin | – | 2 | 1 | 4 | 1 | 0 | 2 | 1 | 1 | 0 | 1 | 4 | 1 |
| 438017 | 7 | 1 | COP9 signalosome complex subunit 5 | YDL216C | 4 | 1 | 2 | 2 | 1 | 2 | 2 | 2 | 2 | 1 | 1 | 1 |
| 497773 | 3 | 0 | Ubiquitin carboxyl-terminal hydrolase 12 | YJL197W | 1 | 1 | 1 | 1 | 0 | 2 | 2 | 1 | 1 | 1 | 2 | 1 |
| 494197 | 4 | 0 | Cop9 signalosome complex subunit 12 | YJR084W | 2 | 2 | 1 | 1 | 1 | 1 | 1 | 1 | 1 | 1 | 2 | 2 |
| 450685 | 3 | 0 | NEDD8-conjugating enzyme UBC12 | YLR306W | 2 | 1 | 1 | 1 | 1 | 2 | 1 | 1 | 2 | 1 | 1 | 1 |
| 503638 | 7 | 0 | NEDD8-activating enzyme E1 regulatory subunit | YPL003W | 2 | 2 | 2 | 2 | 1 | 1 | 1 | 2 | 2 | 1 | 2 | 1 |
| 485679 | 8 | 3 | NEDD8-activating enzyme E1 catalytic subunit | YPR066W | 4 | 2 | 4 | 2 | 1 | 1 | 2 | 2 | 4 | 1 | 2 | 1 |
| 237692 | 9 | 4 | Ubiquitin carboxyl-terminal hydrolase | – | 4 | 1 | 4 | 2 | 1 | 2 | 4 | 2 | 2 | 2 | 4 | 1 |
| 439373 | 5 | 0 | COP9 signalosome complex subunit 1 | – | 2 | 1 | 1 | 2 | 1 | 1 | 1 | 2 | 2 | 1 | 2 | 1 |
| 204652 | 3 | 0 | COP9 signalosome complex subunit 3 | – | 2 | 1 | 1 | 1 | 1 | 1 | 2 | 1 | 2 | 1 | 1 | 1 |
| 444512 | 3 | 0 | COP9 signalosome complex subunit 4 | – | 2 | 1 | 2 | 1 | 1 | 1 | 1 | 2 | 1 | 1 | 0 | 1 |
| 390754 | 4 | 0 | COP9 signalosome protein | – | 1 | 1 | 2 | 2 | 1 | 1 | 2 | 1 | 1 | 1 | 2 | 1 |
| 35248 | 4 | 0 | COP9 signalosome subunit 6 protein | – | 1 | 1 | 1 | 2 | 1 | 1 | 2 | 2 | 1 | 1 | 2 | 1 |
| 437024 | 4 | 1 | Csn8 subunit protein | – | 2 | 1 | 1 | 1 | 1 | 1 | 2 | 1 | 2 | 1 | 4 | 1 |
| 442909 | 2 | 0 | Proteasome component | – | 1 | 1 | 1 | 1 | 2 | 1 | 2 | 1 | 1 | 1 | 1 | 1 |
| 255029 | 2 | 0 | COP9 signalosome complex subunit 2 | – | 2 | 0 | 1 | 1 | 1 | 1 | 1 | 1 | 2 | 1 | 1 | 1 |

#### 4.13.2. Protein ubiquitination

Covalent protein modification by the ubiquitin-proteasome system is a widely conserved regulatory machinery that functions in several essential processes in eukaryotes and was recently detected as a potentially very interesting function during fruiting body development(Krizsán et al. 2019). The regulatory potential of the ubiquitin-proteasome system stems from its ability to attach the 76-amino acid ubiquitin protein to lysines of target proteins, which either destines them for degradation by the proteasome or modifies their activity, cellular localization or DNA-binding ability(LC et al. 2007; Liu and Xue 2011; Finley et al. 2012). Ubiquitination can modify the activity of transcription factors, which provides an additional layer of transcriptional regulation(Freiman and Tjian 2003).

The protein ubiquitination machinery is composed of E1 ubiquitin-activating, E2 ubiquitin-conjugating enzymes and E3 ubiquitin ligases, which attach the ubiquitin to target proteins. While E1 and E2 enzymes are generic components of the system and are present in limited numbers in eukaryotic genomes (Table 37), ubiquitin ligases provide the substrate specificity and come in many subtypes and display considerable diversity due to interchangeable protein components, of which hundreds may exist in the genomes of some species(Hutchins et al. 2013; Zheng and Shabek 2017). The most diverse such parts are F-box and BTB/POZ domain proteins, both of which can confer substrate-specificity to the E3 complexes(R et al. 2003) and RING-type zinc finger proteins, which are scaffold proteins linking target proteins with E2 enzymes(Metzger et al. 2012). These three gene families were found to be strongly expanded in mushroom-forming fungi, with frequent tissue-specific developmental regulation seen in RNA-Seq data(Almási et al. 2019; Krizsán et al. 2019) (see also Table 37). The expansion stems from Agaricomycetes-specific duplication events and a correlation between fruiting body complexity and F-Box, BTB/POZ and RING-type zinc finger copy number and expression has been suggested(Krizsán et al. 2019), which could explain the evolution of increasingly complex mushroom morphologies in the class.

**Table 37.** Summary of developmental expression dynamics of CDE orthogroups of protein ubiquitination and the proteasome related genes across 12 species. Protein ID of a representative protein is given follows by the number of species in which the orthogroup is developmentally regulated at fold change 2 and 4 (FC>2 and FC>4, respectively). Putative function and ortholog in *S. cerevisiae* (if any) are also given. Abbreviations: 0-gene absent, 1-gene present but not developmentally regulated, 2 - developmentally regulated at fold change >2, 4- developmentally regulated at fold change >4. Species names are abbreviated as: Cc – *C. cinerea*, Aa – *A. ampla*, Ab – *A. bisporus*, Ao – *A. ostoyae*, Ca – *C. aegerita*, Lb – *L. bicolor*, Le – *L. edodes*, Lt – *L. tigrinus*, Mk – *M. kentingensis*, Pc – *Ph. chrysosporium*, Po – *P. ostreatus*, Sc – *S. commune*.

| ID | Fold change |  | Putative function | *S. cerevisiae* ortholog | Species |  |  |  |  |  |  |  |  |  |  |  |
| --- | --- | --- | --- | --- | --- | --- | --- | --- | --- | --- | --- | --- | --- | --- | --- | --- |
|  | >2 | >4 |  |  | Cc | Aa | Ab | Ao | Ca | Lb | Le | Lt | Mk | Pc | Po | Sc |
| 354939 | 11 | 5 | BTB/POZ domain containing protein | – | 4 | 1 | 2 | 4 | 4 | 2 | 2 | 4 | 2 | 2 | 4 | 2 |
| 379603 | 9 | 8 | BTB/POZ domain containing protein | – | 4 | 4 | 1 | 4 | 0 | 4 | 4 | 2 | 0 | 4 | 4 | 4 |
| 383774 | 7 | 5 | BTB/POZ domain containing protein | – | 4 | 2 | 0 | 4 | 4 | 2 | 0 | 0 | 0 | 0 | 4 | 4 |
| 405851 | 7 | 5 | BTB/POZ domain containing protein | – | 4 | 0 | 0 | 2 | 2 | 4 | 4 | 0 | 4 | 0 | 4 | 0 |
| 446662 | 5 | 5 | BTB/POZ domain containing protein | – | 4 | 0 | 4 | 1 | 4 | 4 | 0 | 1 | 1 | 1 | 4 | 0 |
| 449824 | 8 | 8 | BTB/POZ domain containing protein | – | 4 | 4 | 1 | 4 | 0 | 4 | 4 | 1 | 4 | 1 | 4 | 4 |
| 451372 | 9 | 4 | BTB/POZ domain containing protein | – | 2 | 4 | 4 | 2 | 1 | 1 | 2 | 2 | 4 | 1 | 2 | 4 |
| 459680 | 11 | 8 | BTB/POZ domain containing protein | – | 4 | 2 | 4 | 4 | 4 | 4 | 4 | 4 | 4 | 1 | 2 | 2 |
| 374447 | 10 | 7 | BTB/POZ domain containing protein | – | 4 | 4 | 4 | 4 | 2 | 1 | 0 | 2 | 2 | 4 | 4 | 4 |
| 545078 | 5 | 1 | BTB/POZ domain containing protein | – | 2 | 2 | 1 | 2 | 0 | 2 | 1 | 1 | 4 | 1 | 1 | 1 |
| 358351 | 5 | 0 | Orthologs of *A. nidulans* GrrA | – | 2 | 1 | 1 | 2 | 2 | 1 | 1 | 1 | 2 | 1 | 2 | 1 |
| 447658 | 10 | 5 | F-box protein | – | 4 | 4 | 1 | 4 | 1 | 2 | 4 | 2 | 4 | 2 | 2 | 2 |
| 503496 | 9 | 5 | F-box protein | – | 4 | 1 | 4 | 4 | 2 | 2 | 4 | 4 | 2 | 1 | 2 | 1 |
| 145676 | 10 | 6 | F-box protein | – | 4 | 2 | 4 | 4 | 2 | 4 | 4 | 2 | 4 | 1 | 2 | 1 |
| 359613 | 10 | 6 | F-box protein | – | 2 | 2 | 4 | 4 | 4 | 2 | 4 | 4 | 4 | 1 | 2 | 1 |
| 470790 | 10 | 6 | F-box protein | – | 4 | 2 | 4 | 4 | 2 | 4 | 4 | 2 | 4 | 1 | 1 | 2 |
| 166096 | 9 | 7 | F-box protein | – | 4 | 0 | 0 | 4 | 2 | 4 | 4 | 4 | 4 | 2 | 4 | 0 |
| 356986 | 9 | 6 | F-box protein | – | 4 | 1 | 4 | 4 | 2 | 4 | 2 | 2 | 4 | 1 | 4 | 1 |
| 377171 | 7 | 2 | F-box protein | – | 2 | 2 | 2 | 4 | 1 | 1 | 1 | 2 | 4 | 1 | 1 | 2 |
| 382134 | 7 | 4 | F-box protein | – | 4 | 1 | 1 | 4 | 2 | 1 | 4 | 2 | 0 | 1 | 4 | 2 |
| 438595 | 7 | 7 | F-box protein | – | 4 | 1 | 0 | 4 | 1 | 4 | 4 | 0 | 4 | 1 | 4 | 4 |
| 445665 | 8 | 7 | F-box protein | – | 4 | 1 | 4 | 4 | 4 | 4 | 4 | 2 | 4 | 1 | 0 | 1 |
| 451935 | 7 | 6 | F-box protein | – | 1 | 4 | 1 | 4 | 0 | 4 | 0 | 4 | 0 | 4 | 4 | 2 |
| 471789 | 6 | 6 | F-box protein | – | 4 | 1 | 4 | 4 | 4 | 4 | 4 | 0 | 1 | 0 | 0 | 1 |
| 490594 | 7 | 3 | F-box protein | – | 4 | 1 | 1 | 2 | 2 | 1 | 4 | 1 | 2 | 1 | 4 | 2 |
| 501200 | 12 | 9 | F-box protein | – | 4 | 4 | 4 | 4 | 4 | 4 | 4 | 2 | 2 | 2 | 4 | 4 |
| 440772 | 11 | 7 | RING-type zinc finger domain containing protein | – | 4 | 4 | 2 | 4 | 2 | 2 | 2 | 0 | 4 | 4 | 4 | 4 |
| 364816 | 10 | 3 | RING-type zinc finger domain containing protein | — | 2 | 1 | 4 | 2 | 2 | 2 | 4 | 2 | 2 | 4 | 2 | 1 |
| 378120 | 8 | 8 | RING-type zinc finger domain containing protein | — | 4 | 4 | 0 | 4 | 4 | 1 | 4 | 1 | 4 | 0 | 4 | 4 |
| 455937 | 9 | 2 | RING-type zinc finger domain containing protein | — | 4 | 2 | 2 | 2 | 0 | 4 | 2 | 2 | 2 | 1 | 2 | 0 |
| 466739 | 10 | 7 | RING-type zinc finger domain containing protein | — | 4 | 2 | 2 | 4 | 4 | 1 | 4 | 4 | 4 | 2 | 4 | 1 |
| 467805 | 10 | 7 | RING-type zinc finger domain containing protein | — | 2 | 1 | 4 | 4 | 4 | 2 | 4 | 2 | 4 | 4 | 4 | 1 |
| 543850 | 11 | 9 | RING-type zinc finger domain containing protein | — | 4 | 4 | 2 | 4 | 4 | 2 | 4 | 4 | 0 | 4 | 4 | 4 |

The widespread cellular roles of ubiquitination suggests that it should be instrumental during fruiting body morphogenesis too. In line with this, the protein ubiquitination system was found differentially expressed in *F. velutipes* fruiting bodies(M et al. 2006; Liu et al. 2020b) and in caps of the same species subjected to differential CO_2_ treatments(Yan et al. 2019). In recent comparative transcriptomics analyses, we provided evidence for the developmental regulation of several components, mostly those that confer substrate specificity to E3 ligase complexes, in multiple Agaricomycetes species, including *C. cinerea, A. ostoyae, A. ampla, S. commune* or *P. ostreatus*, among others(Almási et al. 2019; Krizsán et al. 2019; Merényi et al. 2021). BTB domain-containing proteins, a family of E3 proteins, were reported to be upregulated in fruiting bodies in *Antrodia cinnamomea* (Lu et al. 2014). The combination of a remarkable gene family expansion with developmental regulation suggests that the ubiquitin-proteasome system could play a pivotal role in fruiting body development, although mechanistic details of the process remain to be understood. Ubiquitylation is key in developmental processes of the Ascomycota also, including fruiting body formation, and in some cases more mechanistic details have been uncovered (reviewed in(Riquelme et al. 2018)). However, a similar expansion of F-Box domains, as seen in Agaricomycetes, has not been reported.

Hereafter, we present an annotation of components of the protein ubiquitination machinery in selected mushroom-forming fungi, with particular emphasis on *C. cinerea* genes and comparisons across species. In this presentation, we follow the classification of E3 ubiquitin ligases used by Hutchins et al(Hutchins et al. 2013) and identify protein diversity of each protein family based on Interpro domain HMM-s. Results of the annotation along with orthologous genes and conservation levels are shown in Supplementary table 4. A limitation of this approach is that the presence of a defining domain in the protein does not in all the cases, indicate a function in the ubiquitination machinery. For example, BTB/POZ domains, which contribute to expanding the diversity of E3 ligase complexes(R et al. 2003) can, in rare cases, also occur in proteins not related to ubiquitination (e.g. cytoskeleton or transcription regulating proteins(Stogios et al. 2005)).

Several components of the ubiquitination machinery are very conserved in Agaricomycetes genomes, including E1 (on average 3 genes) and E2 enzymes (19-24 genes), Cullin genes (4-10 genes) and deubiquitinating enzyme-encoding genes (23-27 genes) (Supplementary table 4). Most of these genes show no or modest expression dynamics, which is not surprising given that the ubiquitination machinery is needed constantly in the cell. Of the E1 enzymes, only *S. cerevisiae* Uba3 orthologs (*C. cinerea* protein ID: 485679), which belong to the neddylation pathway, are developmentally regulated to some extent (8/3 species at FC=2/4). In *C. cinerea* 14 of the 23 E2 enzymes and half (13 of 26) of deubiquitinating enzymes are developmentally regulated, potentially indicating functional diversification of E2 enzymes and deubiquitinating enzymes during development (Supplementary table 5). We found that none of the cullin genes in *C. cinerea* are developmentally regulated. Significantly larger dynamics can be observed in E3 ligase components, which are discussed in detail below.

**RING-type Zinc finger proteins** - RING-domain containing proteins, named after RING (really interesting new gene), are E3 ubiquitin ligases that confer interchangeability to the ligase complexes(Deshaies and Joazeiro 2009). Eukaryotic genomes contain large numbers of RING-domain proteins, which, along with other E3 ligases, enable the ubiquitin system to selectively ubiquitinate proteins. RING-domain proteins are generally poorly known in filamentous fungi, though it was recently noticed that they are strongly overrepresented in Agaricomycetes(Krizsán et al. 2019). The *mfbAc* (=Le.MFB1) gene of *L. edodes* (Lenedo1_1172639) described as a protein with adhesive properties by Kondoh et al(Kondoh et al. 1995) also belongs to this gene family.

The examined Agaricomycetes possess 81-148 RING-domain containing proteins (Table 37), consistent with a previous broader sampling across 200 species. On average, 30-50 RING-domain proteins were developmentally regulated in the examined species(Almási et al. 2019; Krizsán et al. 2019), representing a significant pool of potentially important regulators. RING-domain proteins are highly variable, with most of them not showing clear orthology across species. Nevertheless, we found eight CDE orthogroups which displayed conserved developmental expression, mostly at FC>2 (Table 37). For example, one of these (represented by *C. cinerea* 378120) showed a very distinct peak in gills of *C. cinerea, A. ostoyae, P. ostreatus* and caps (containing gills) of *M. kentingensis, L. edodes* as well as mature fruiting bodies of *S. commune* and *A. ampla*. Another CDE orthogroup (represented by *C. cinerea* 440772) was upregulated in gill tissues of mature fruiting bodies in *P. ostreatus, A. ostoyae, M. kentingensis*, *L. bicolor* and mature fruiting bodies of *S. commune* and *A. ampla*, but not in *A. bisporus, L. edodes* or *C. cinerea*. These two orthogroups coincide (partly) with meiosis and sporulation, so they could be involved in these, or other gill-specific processes. Genes in another CDE orthogroup (C. cinerea protein ID: 543850) showed an abrupt, although modest upregulation at the initiation of fruiting body development and expression peak in mature fruiting bodies or gills therein in *C. cinerea, A. ostoyae, C. aegerita, A. ampla, L. bicolor, L. tigrinus, Ph. chrysosporium, P. ostreatus* and *S. commune* (not in *L. edodes*).

**F-box proteins -** F-box proteins were recently identified as a potentially key group of proteins in fruiting body development(Krizsán et al. 2019). They are generally responsible for providing target specificity to the ubiquitination machinery, by binding and docking substrate proteins to Cullin E3 ubiquitin ligase complexes. Some F-box proteins interact with multiple target proteins while others only with a single protein(Jonkers and Rep 2009). Similarly, some F-Box proteins function independently of the SCF complex(Jonkers and Rep 2009), allowing for additional, yet poorly known regulatory functions to emerge. For example, F-box proteins can act as transcriptional co-regulators in plants(Chae et al. 2008) and are present in large numbers in plant genomes (up to 700-1000 genes(Xu et al. 2009)), providing the possibility to assemble an enormous diversity of alternative ubiquitin ligase complexes.

The F-box family is highly expanded in mushroom-forming fungi and is composed of fast-evolving proteins with little similarity to each other. Agaricomycetes possess 69 to ∼1200 F-box proteins in their genomes, whereas other filamentous fungi and yeasts possess up to 90 and 20 such genes(Krizsán et al. 2019), respectively. This represents a strong enrichment in Agaricomycetes genomes (P<10^-140^), considering that the number of other proteins is not proportionately higher in this group than in other fungi. F-box protein encoding genes often showed tissue-specific expression and their numbers showed a positive correlation with the complexity level of the fruiting bodies(Krizsán et al. 2019).

F-Box proteins are highly variable and form few conserved orthogroups, compared to their diversity in fungal genomes. However, we found eleven orthogroups that showed some level of conservation and consistent developmental regulation across species (Table 37). These included orthologs of *A. nidulans* GrrA (*C. cinerea* protein ID: 358351), which is needed for meiosis and ascospore maturation(S et al. 2006a; Riquelme et al. 2018), are conserved across Agaricomycetes and developmentally regulated in five species (FC>2), indicating that some F-Box proteins are conserved despite the general volatility of the family. Due to this high variability, F-box proteins are hard to examine systematically using a comparative transcriptomics approach. Rather, they likely often fulfill species- or clade-specific roles, which could be studied either with more densely sampled transcriptomic data, or reverse genetics approaches.

**BTB/POZ domain proteins** - Proteins that contain the BTB/POZ motif, a protein-protein interaction (homodimerization) domain, can possess diverse cellular functions, the most frequent of which are regulation of chromatin structure (thus, transcription) and protein ubiquitination, as members of E3 ubiquitin ligase complexes(S et al. 1994; R et al. 2003). They are poorly known in fungi.

In animals, plants as well as fungi outside the Agaricomycetes, BTB domains are mostly fused to other conserved protein domains(Stogios et al. 2005) (in animals, mostly to zinc fingers(S et al. 1994)). In contrast, we observed that in the Agaricomycetes, less than 10% of BTB/POZ domain proteins associate with any other domains and virtually never with DNA binding transcription factor domains, indicating that the gene family expansion in mushroom-forming fungi is concerned with proteins that only contain the BTB/POZ domain. This domain composition suggests that the BTB/POZ family is primarily involved in E3 ligases in Agaricomycetes, therefore we discuss this family under the chapter on protein ubiquitination.

We found 8 CDE orthogroups with some level of conserved developmental expression in Agaricomycetes (Table 37). Members of two (*C. cinerea* protein ID: 379603, 449824) of these orthogroups are co-expressed in stages or tissues in which meiosis and sporulation occurs: gills of *A. ostoyae* and *P. ostreatus*, mature fruiting bodies of *Ph. chrysosporium, A. ampla* and *S. commune,* mature caps of *M. kentingensis, L. bicolor, L. edodes* (but not developmentally regulated in *C. cinerea*). Another CDE orthogroup (*C. cinerea* protein ID: 459680) was upregulated in stipes of *C. cinerea, A. ostoyae, M. kentingensis, L. edodes* (but not in *P. ostreatus* and *A. bisporus*), primordium stages of *A. ampla* and *S. commune* and during early development (primordia + young fruiting bodies) of *L. bicolor*. A fourth CDE orthogroup, which contains reciprocal best hits to the putative E3 ubiquitin ligase protein *btb1* of *Sch. pombe* (*C. cinerea* protein ID: 545078), but contains also domains that suggest it regulates chromatin condensation and thus gene expression. Members of this orthogroup were upregulated typically late during development.

### 4.14. Other gene groups

#### 4.14.1. Transporters

Among the conserved developmentally expressed genes (CDE orthogroups) we detected 49 putatively transport-related gene families. Many of these genes belong to the Major Facilitator Superfamily or ABC transporters, fast-evolving gene families that are represented by hundreds of genes in fungal genomes. Transporters have been reported in transcriptomic studies of several mushroom-forming species, both in fruiting body(Song et al. 2018b; Krizsán et al. 2019) and ectomycorrhiza formation(Xu et al. 2015). Due to the fast evolution of these families, orthology to genes of Ascomycota model systems and even within the Agaricomycetes is lacking in most cases, making it hard to infer their functions. Nevertheless, some conserved transporter genes were detected, including orthologs of *S. cerevisiae* Pns1 (choline transporter, *C. cinerea* 355745), *A. nidulans* MepA (ammonium permease, *C. cinerea* 538158), *Sch. pombe* Kha1 (plasma membrane potassium ion/proton symporter, *C. cinerea* 449705), *S. cerevisiae* Emp70 (endosomal transport protein, *C. cinerea* 495380), *S. cerevisiae* Opt1 (plasma membrane oligopeptide transporter, *C. cinerea* 496395), among others (see Table 38). In the absence of functional annotations for the vast majority of these genes, we list the genes in Table 38, but do not go into detailed analyses, because it is unlikely that we can arrive at any reasonable speculation about their function.

**Table 38.** Summary of developmental expression dynamics of CDE orthogroups of transporter encoding genes across 12 species. Protein ID of a representative protein is given follows by the number of species in which the orthogroup is developmentally regulated at fold change 2 and 4 (FC>2 and FC>4, respectively). Putative function and ortholog in *S. cerevisiae* (if any) are also given. Abbreviations: 0-gene absent, 1-gene present but not developmentally regulated, 2 - developmentally regulated at fold change >2, 4- developmentally regulated at fold change >4. Species names are abbreviated as: Cc – *C. cinerea*, Aa – *A. ampla*, Ab – *A. bisporus*, Ao – *A. ostoyae*, Ca – *C. aegerita*, Lb – *L. bicolor*, Le – *L. edodes*, Lt – *L. tigrinus*, Mk – *M. kentingensis*, Pc – *Ph. chrysosporium*, Po – *P. ostreatus*, Sc – *S. commune*.

| ID | Fold change |  | Putative function | S.<br>*cerevisiae*<br>ortholog | Species |  |  |  |  |  |  |  |  |  |  |  |
| --- | --- | --- | --- | --- | --- | --- | --- | --- | --- | --- | --- | --- | --- | --- | --- | --- |
|  | >2 | >4 |  |  | Cc | Aa | Ab | Ao | Ca | Lb | Le | Lt | Mk | Pc | Po | Sc |
| 538158 | 8 | 5 | Ammonium/urea transporter | — | 4 | 1 | 0 | 4 | 0 | 4 | 2 | 2 | 4 | 1 | 4 | 2 |
| 404532 | 9 | 5 | Major facilitator superfamily protein | — | 4 | 1 | 0 | 2 | 4 | 2 | 4 | 0 | 4 | 2 | 4 | 2 |
| 492885 | 9 | 5 | Major facilitator superfamily protein | — | 4 | 1 | 2 | 2 | 1 | 1 | 2 | 4 | 4 | 2 | 4 | 4 |
| S.c.<br>253716<br>3 | 9 | 5 | Multi antimicrobial extrusion protein | — | 0 | 2 | 0 | 4 | 4 | 2 | 2 | 4 | 4 | 1 | 4 | 2 |
| 233439 | 9 | 6 | Tetrapeptide transporter; OPT1/isp4 family | — | 2 | 4 | 2 | 4 | 4 | 1 | 4 | 4 | 1 | 2 | 4 | 1 |
| 30849 | 11 | 6 | 10TM phosphate transporter | — | 4 | 4 | 2 | 4 | 4 | 2 | 2 | 2 | 4 | 1 | 4 | 2 |
| 371400 | 11 | 9 | Major facilitator superfamily protein | — | 4 | 1 | 4 | 4 | 4 | 4 | 4 | 4 | 2 | 4 | 4 | 2 |
| 464027 | 7 | 6 | Major facilitator superfamily protein | — | 4 | 1 | 4 | 0 | 4 | 4 | 2 | 0 | 4 | 1 | 4 | 1 |
| 468563 | 6 | 6 | Major facilitator superfamily protein | — | 4 | 4 | 0 | 4 | 1 | 0 | 4 | 1 | 0 | 4 | 4 | 1 |
| 501434 | 9 | 9 | Major facilitator superfamily protein | — | 4 | 0 | 4 | 4 | 4 | 4 | 4 | 4 | 4 | 1 | 4 | 1 |
| 436779 | 10 | 0 | Nuclear pore protein 84/107 | — | 2 | 2 | 2 | 2 | 1 | 1 | 2 | 2 | 2 | 2 | 2 | 2 |
| S.c.<br>228395<br>7 | 8 | 6 | Sugar/inositol transporter | — | 0 | 2 | 0 | 4 | 4 | 4 | 4 | 4 | 0 | 1 | 4 | 2 |
| 355160 | 10 | 9 | Threonine/Serine exporter; ThrE | — | 4 | 4 | 1 | 4 | 4 | 4 | 4 | 2 | 4 | 1 | 4 | 4 |
| 432854 | 10 | 6 | Membrane transport protein family | YBR287W | 4 | 2 | 2 | 4 | 4 | 2 | 4 | 4 | 1 | 1 | 2 | 4 |
| 449861 | 10 | 4 | GABA-specific permease | YDL210W | 2 | 1 | 4 | 2 | 4 | 1 | 2 | 4 | 4 | 2 | 2 | 2 |
| 428934 | 10 | 6 | Sugar transporter STL1 | YDR536W | 4 | 2 | 4 | 4 | 4 | 4 | 4 | 1 | 2 | 1 | 2 | 2 |
| 422892 | 10 | 2 | Purine-cytosine permease FCY2 | YER056C | 2 | 2 | 4 | 2 | 4 | 2 | 1 | 2 | 2 | 2 | 2 | 1 |
| 455616 | 5 | 5 | Zinc-regulated transporter 1 | YGL255W | 4 | 1 | 4 | 4 | 0 | 4 | 4 | 1 | 1 | 1 | 1 | 1 |
| 465413 | 9 | 6 | Urea transporter | YHL016C | 4 | 1 | 0 | 4 | 4 | 4 | 4 | 2 | 4 | 1 | 2 | 2 |
| 496395 | 9 | 7 | Oligopeptide transporter 1 | YJL212C | 4 | 4 | 2 | 4 | 4 | 1 | 4 | 2 | 4 | 1 | 4 | 0 |
| 502591 | 9 | 6 | Polyamine transporter 1 | YLL028W | 4 | 2 | 4 | 1 | 1 | 2 | 4 | 4 | 4 | 0 | 2 | 4 |
| 360410 | 9 | 8 | Thiamine pathway transporter THI73 | YLR004C | 4 | 1 | 4 | 4 | 1 | 4 | 4 | 4 | 4 | 1 | 4 | 2 |
| 495380 | 7 | 1 | Transmembrane 9 superfamily member 1 | YLR083C | 4 | 2 | 2 | 2 | 1 | 1 | 2 | 1 | 0 | 1 | 2 | 2 |
| 223186 | 11 | 3 | Inorganic phosphate transporter PHO84 | YML123C | 4 | 1 | 2 | 2 | 2 | 4 | 2 | 2 | 2 | 2 | 4 | 2 |
| 393558 | 10 | 6 | Vacuolar basic amino acid transporter 1 | YMR088C | 4 | 4 | 2 | 4 | 2 | 1 | 4 | 2 | 1 | 4 | 2 | 4 |
| 502991 | 10 | 6 | Major facilitator superfamily protein | YMR279C | 4 | 4 | 4 | 4 | 2 | 2 | 4 | 2 | 0 | 1 | 4 | 2 |
| 421680 | 7 | 3 | Calcium channel YVC1 | YOR087W | 2 | 2 | 2 | 4 | 1 | 1 | 4 | 1 | 1 | 1 | 2 | 4 |
| 542873 | 10 | 5 | ATP-dependent permease PDR12 | YPL058C | 2 | 2 | 2 | 4 | 2 | 4 | 1 | 2 | 4 | 1 | 4 | 4 |
| 462297 | 10 | 2 | Sulfate transporter YPR003C | YPR003C | 2 | 1 | 2 | 4 | 2 | 2 | 2 | 2 | 4 | 1 | 2 | 2 |
| S.c.<br>250167<br>7 | 9 | 6 | Aquaporin | — | 0 | 4 | 4 | 4 | 2 | 1 | 4 | 2 | 4 | 1 | 4 | 2 |
| A.o.<br>253413 | 6 | 6 | Major facilitator superfamily protein | — | 0 | 0 | 4 | 4 | 0 | 4 | 4 | 4 | 0 | 0 | 4 | 0 |
| 493842 | 9 | 6 | Major facilitator superfamily protein | — | 4 | 1 | 4 | 4 | 4 | 2 | 2 | 1 | 4 | 1 | 4 | 2 |
| 495288 | 8 | 6 | Major facilitator superfamily protein | — | 4 | 4 | 2 | 4 | 4 | 4 | 0 | 4 | 1 | 1 | 2 | 1 |
| 496145 | 9 | 7 | Sugar/inositol transporter | — | 4 | 4 | 1 | 4 | 2 | 1 | 4 | 4 | 4 | 2 | 4 | 1 |
| 498282 | 10 | 6 | Major facilitator superfamily protein | — | 4 | 2 | 2 | 4 | 1 | 4 | 4 | 2 | 2 | 4 | 4 | 1 |
| 474705 | 10 | 6 | ABC transporter | — | 4 | 2 | 0 | 4 | 2 | 4 | 4 | 1 | 2 | 2 | 4 | 4 |
| 357414 | 6 | 5 | Major facilitator superfamily protein | — | 4 | 1 | 4 | 4 | 1 | 0 | 0 | 4 | 2 | 1 | 4 | 1 |
| 420930 | 8 | 7 | Major facilitator superfamily protein | — | 4 | 4 | 4 | 4 | 2 | 4 | 4 | 1 | 0 | 1 | 4 | 1 |
| 463045 | 9 | 6 | Major facilitator superfamily protein | — | 4 | 2 | 4 | 4 | 2 | 4 | 0 | 2 | 0 | 1 | 4 | 4 |
| 465489 | 9 | 5 | Major facilitator superfamily protein | — | 4 | 1 | 4 | 4 | 0 | 1 | 2 | 2 | 4 | 2 | 4 | 2 |
| 488859 | 9 | 5 | Mitochondrial substrate/solute carrier | — | 4 | 2 | 4 | 2 | 4 | 1 | 2 | 1 | 4 | 1 | 4 | 2 |
| 451236 | 10 | 2 | Secretory carrier membrane proteins (SCAMP) | — | 2 | 2 | 1 | 4 | 2 | 2 | 4 | 2 | 2 | 1 | 2 | 2 |
| 374240 | 10 | 8 | Sodium/calcium exchanger | — | 4 | 1 | 0 | 4 | 2 | 4 | 4 | 2 | 4 | 4 | 4 | 4 |
| 450674 | 9 | 7 | Sugar phosphate transporter | — | 4 | 2 | 4 | 4 | 2 | 1 | 4 | 4 | 4 | 1 | 4 | 1 |
| 537526 | 10 | 7 | Sugar phosphate transporter | — | 4 | 4 | 2 | 4 | 4 | 2 | 2 | 1 | 4 | 1 | 4 | 4 |
| 358584 | 7 | 5 | Sugar/inositol transporter | — | 4 | 4 | 2 | 4 | 1 | 0 | 1 | 4 | 1 | 1 | 4 | 2 |
| 487841 | 11 | 8 | Sugar/inositol transporter | — | 4 | 4 | 4 | 4 | 2 | 4 | 4 | 4 | 2 | 2 | 1 | 4 |
| 502428 | 7 | 5 | Sugar/inositol transporter | — | 4 | 2 | 2 | 4 | 1 | 1 | 1 | 4 | 4 | 1 | 4 | 1 |
| S.c.<br>262770<br>3 | 6 | 5 | Sugar/inositol transporter | — | 0 | 2 | 4 | 4 | 0 | 0 | 4 | 0 | 4 | 0 | 0 | 4 |
| 397645 | 9 | 6 | High glucose sensor RGT2 | YDL138W | 4 | 4 | 4 | 2 | 0 | 1 | 2 | 2 | 4 | 1 | 4 | 4 |
| 463015 | 8 | 7 | Hexose transporter HXT10 | YFL011W | 4 | 4 | 0 | 4 | 4 | 0 | 4 | 4 | 0 | 1 | 4 | 2 |

Notable CDE orthogroups include an MFS transporter *C. cinerea* 465489, which was developmentally regulated in 9/5 species (FC>2/4) and FB-init in *C. cinerea, A. ostoyae, P. ostreatus, S. commune* (in the latter at FC>2 only). A phosphate transporter CDE orthogroup (*C. cinerea* protein ID: 30849) was FB-init in *C. cinerea, C. aegerita, A. ostoyae, A. ampla, S. commune, M. kentingensis* and, at FC>2 in *L. bicolor (*not in *L. edodes*) (Table 38). A group of zinc-regulated transceptors (transporter + receptor, *C. cinerea* protein ID: 455616) orthologous to *S. cerevisiae* Zrt1 was also detected. In summary, transporters can be expected to fulfill key functions in fruiting bodies, in moving around goods, signal molecules, modulating osmolarity to name a few, however, information about these in fruiting bodies is still limited.

**Aquaporins** are water and solute-transporting membrane channels that are integral to several different aspects of the fungal lifestyle. They were reported to be overexpressed in primordia and stipes of *F. filiformis*, relative to other tissues and developmental stages(Liu et al. 2020b). They were also reported to be upregulated in *L. bicolor* fruiting bodies and ectomycorrhizae(Xu et al. 2015).

We found 2-11 aquaporin genes in the examined Agaricomycetes and observed conserved aquaporin expression in two aquaporin orthogroups. One of these (*C. cinerea* protein ID: 373141) contains putative orthologs of yeast Fps1, an aquaglyceroporin that has suggested roles in acetate uptake or glycerol transport(Nehls and Dietz 2014), not water uptake (see under chapter on Acetyl CoA). It is worth noting that aquaglyceroporins were also differentially regulated in *L. bicolor* fruiting bodies(Xu et al. 2015) and that these proteins were shown to have a capacity to transport various solutes, including NH_3_ and glycerol. This led Xu et al to hypothesize that these genes could be involved in osmosis regulation or signaling(Xu et al. 2015), which is different from the conclusion we make above on acetate transport.

The other orthogroup (represented by *S. commune* 2501677) comprises stipe-specific aquaporin genes, which show similar expression trajectories in *A. bisporus, A. ostoyae, M kentingensis* and *P. ostreatus*, as well as in an external *F. filiformis* dataset(Liu et al. 2020b). In all of these species, the genes showed expression peaks in stipes, mostly in young and mature fruiting bodies. It is notable that this pattern is seen in pileate-stipitate species and has been noted in a previous 3-species comparison of transcriptomes(Plaza et al. 2014b). Species without separate cap and stipe tissues either do not have this gene (*Pt. gracilis*) or they do, but the gene is not developmentally regulated (*S. commune, Ph. chrysosporium*). The current genome assembly of *C. cinerea* does not contain an orthologue in this group, which might be because it became dispensable for the unique developmental trajectory of this species or because the genome assembly is incomplete at this genomic region. The marked stipe expression peak suggests a widespread role of water transport during mushroom development. Further, the increasing expression parallel with developmental progression could indicate a role in water supply needed for cellular expansion. On a related note, *C. neoformans* also shows an upregulation of an aquaporin gene (Cryne_JEC21_1_3081) late in its development.

A consequence of growth by cell expansion is that growth rate is limited by the speed of water transport, not cell division and CW material deposition(Roper and Seminara 2019). Therefore, aquaporin-facilitated water transport might be a selected trait in many mushroom-forming fungi that undergo fast growth. Although unknown at this moment, if fruiting body expansion is mediated by locally produced osmolytes (e.g. by breaking down glycogen), then that could produce a gradient that draws water towards the expanding tissues. This may be further facilitated by water channels formed by aquaporins. Finally, it is possible that other physical solutions, e.g. inter-hyphal water passageways also help transporting (e.g. via surface tension) water in the fruiting body.

The shared stipe upregulation of aquaporins can shed light on the origins of the typical mushroom (pileate-stipitate) morphology. All examined stipitate Agaricales species showed a stipe-specific aquaporin expression, which supports a single origin of pileate-stipitate fruiting bodies in the Agaricales. On the other hand, *L. tigrinus,* which evolved pileate-stipitate fruiting bodies independently in the Polyporales, does not show stipe-specific aquaporin expression.

#### 4.14.2. Stress response genes

Diverse stress-related genes were detected among CDE orthogroups. Based on functional annotations and orthology relationships, many of these seem to be related to protection from oxidative stress, either through direct protection by their protein products (peroxiredoxins) or via the produced metabolites (ergothioneine) that protect from oxidative stress. We found evidence for all three pathways of reactive oxygen detoxification, the peroxiredoxin pathway, cytoplasmic catalases and glutathione peroxidases(Breitenbach et al. 2015), being active in fruiting bodies. The dynamic expression of these genes may point to the existence of an oxidative environment in fruiting bodies. However, it is currently unknown in which tissues/stages and in what cellular contexts this may arise. One possibility is that the action of cell wall modifying enzymes (e.g. peroxidases), which require an oxidative environment (H_2_O_2_), results in excess reactive oxygen species, though this has not been confirmed yet and remains a hypothesis for now. Below we discuss the main stress-related CDE orthogroups which we detected in the developmental transcriptomes of Agaricomycetes.

**Peroxiredoxins** - Peroxiredoxins are antioxidant proteins that can reduce diverse reactive oxygen species (ROS), including H_2_O_2_, and are one of the three cellular pathways of ROS detoxification in fungi. They have also been reported to be involved in signal transduction. The function of peroxiredoxins are not known in fruiting bodies and we identified a single report of peroxiredoxin upregulation in fruiting bodies, in *Termitomyces*(Rahmad et al. 2014). Another gene from *Antrodia camphorata* was cloned and biochemically characterized(Wen et al. 2007). We found orthologs of peroxiredoxin Tpx1 of *Sch. pombe* and Tsa1 of *S. cerevisiae* (*C. cinerea* protein ID: 468305) to be developmentally regulated in 12/8 species (FC>2/4) (Table 39). Tpx1 has been characterized as the H_2_O_2_ sensor that relays the redox signal to the transcription factor Pap1 by inducing the formation of intramolecular disulfide bonds in Pap1(Jara et al. 2007). Orthologs of Tpx1 are upregulated mostly in gill and cap tissues of mature fruiting bodies in the examined species. Orthologs of the Pap1 transcription factor (represented by *C. cinerea* 378471) are expressed constantly in mushroom-forming fungi. Orthologs of yeast Ahp1 peroxiredoxin (*C. cinerea* protein ID: 444300) were also developmentally regulated in the majority of species (10/4 species at FC=2/4, Table 39). This gene preferentially eliminates organic peroxides, rather than H_2_O_2_ and relays signal to the Cad1 transcription factor in *S. cerevisiae*(Iwai et al. 2010). In *C. cinerea* this gene shows a very strong induction in young fruiting body gills, whereas in other species no evidence for a transcript enrichment in gill tissues was found. Finally, we detected orthologs of *S. cerevisiae* Dot5 (*C. cinerea* protein ID: 379600) (Table 39), a thioredoxin (thiol-specific peroxidase) that is involved in the detoxification of peroxides(Cha et al. 2003; Izawa et al. 2003), but was also reported to be involved in the regulation of telomere length. Thioredoxins, together with peroxiredoxins (thioredoxin peroxidase) make up the thioredoxin system, which acts similarly to the glutaredoxin system in ROS scavenging. This orthogroup showed a characteristic upregulation in primordia relative to vegetative mycelium, then lower expression in late developmental stages in *C. cinerea, L. edodes, C. aegerita, A. ostoyae* and *A. ampla*, but not in other species.

**Table 39.**
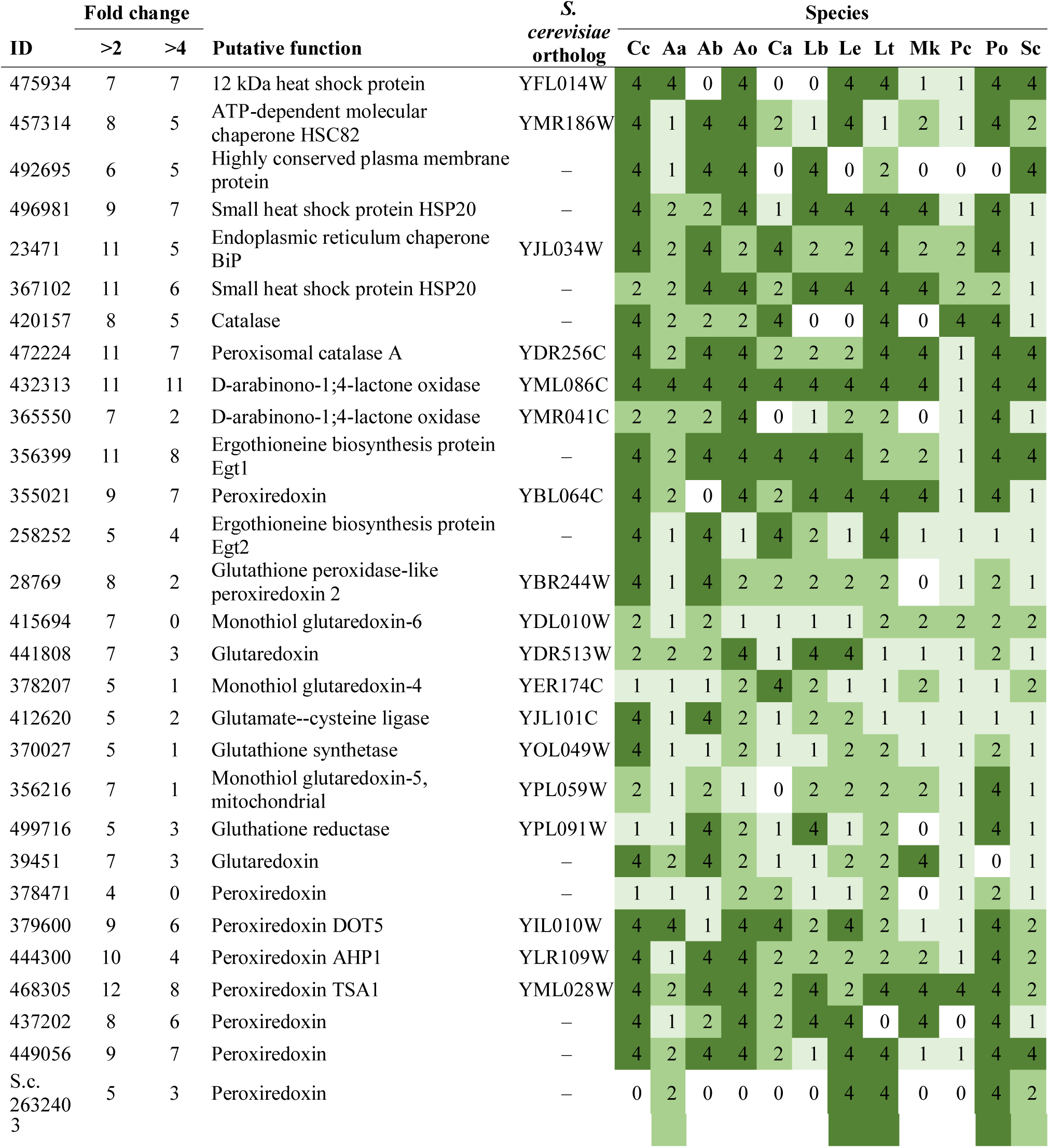
Summary of developmental expression dynamics of CDE orthogroups of stress response genes across 12 species. Protein ID of a representative protein is given follows by the number of species in which the orthogroup is developmentally regulated at fold change 2 and 4 (FC>2 and FC>4, respectively). Putative function and ortholog in *S. cerevisiae* (if any) are also given. Abbreviations: 0-gene absent, 1-gene present but not developmentally regulated, 2 - developmentally regulated at fold change >2, 4- developmentally regulated at fold change >4. Species names are abbreviated as: Cc – *C. cinerea*, Aa – *A. ampla*, Ab – *A. bisporus*, Ao – *A. ostoyae*, Ca – *C. aegerita*, Lb – *L. bicolor*, Le – *L. edodes*, Lt – *L. tigrinus*, Mk – *M. kentingensis*, Pc – *Ph. chrysosporium*, Po – *P. ostreatus*, Sc – *S. commune*.

**Ergothionine biosynthesis** - We also detected orthologs of the ergothioneine biosynthesis protein Egt1 of *S. pombe*. Ergothioneine is a histidine-derived antioxidant metabolite known to confer beneficial health effects to mushroom fruiting bodies(Weigand-Heller et al. 2012; Kalaras et al. 2017). For eukaryotic cells, it might be needed for growth and survival under oxidative stress conditions, for example during conidial survival and germination(MH et al. 2012; Sheridan et al. 2016). Ergothioneine is produced via a two-step reaction pathway in *S. pombe*(Pluskal et al. 2014), comprising Egt1 then Egt2. Both are conserved in *C. cinerea* (*C. cinerea* protein ID: 356399 and 258252, respectively, Table 39), however, only the first gene is developmentally regulated in the majority of the species, the other is not. Expression patterns of the Egt1 orthologs varied, in certain species they are FB-init, while in others it showed stipe-upregulation (*A. ostoyae, L. bicolor, L. edodes, L. tigrinus, M. kentingensis*, but not *C. cinerea*). How this can be explained is not clear, nevertheless, our data provide some evidence for the upregulation of ergothioneine biosynthesis in fruiting bodies. Orthologs of *Sch. pombe* Egt2, which catalyzes the second step of ergothioneine production (*C. cinerea* protein ID: 258252) were developmentally regulated in a more restricted set of species.

**Dehydro-D-arabinono-1;4-lactone** - Two orthogroups that make up the biosynthesis pathway of dehydro-D-arabinono-1;4-lactone were developmentally expressed in several species. Dehydro-D-arabinono-1;4-lactone (previously known as D-erythroascorbate) is a protective metabolite against oxidative stress and is produced by a pathway comprising Ara1, Ara2 and Alo1 in *S. cerevisiae*(K et al. 2006). Of these, Ara2 and Alo1 have clear orthologs in Agaricomycetes (represented by *C. cinerea* 365550 and 432313, respectively)(Table 39), whereas Ara1 does not. Both Ara2 and Alo1 orthologs are strongly upregulated in fruiting bodies. ALO1 orthologs showed a clear upregulation in stipe tissues of *C. cinerea, A. ostoyae* and *P. ostreatus* (but not in *L. bicolor or L. edodes*), whereas in *S. commune, A. ampla and L. edodes* it is upregulated in primordia and stays high throughout fruiting body development. In *C. aegerita*, the gene is strongly upregulated in the first primordium stage, then expression drops to low levels.

**Catalases -** Two catalase orthogroups (represented by *C. cinerea* 472224 and 420157)(Table 39) were developmentally regulated in the majority of species. Catalases are involved in H_2_O_2_ scavenging and thus protection against reactive oxygen species, but also in other pathways (such as beta-oxidation, see above), so their assignment to oxidative stress response is less certain than that of the above-mentioned genes.

**Glutathione system -** The glutathione system is composed of (i) glutathione, a 3- amino acid redox-active metabolite(Pócsi et al. 2004; Sato et al. 2009); (ii) glutathione peroxidases, which reduce H_2_O_2_; (iii) glutathione reductases, which convert oxidized glutathione back to its non-oxidized form; as well as (iv) glutaredoxins, which repair oxidatively damaged proteins. In *S. cerevisiae* glutathione is produced by a two-step reaction from L-cysteine and L-glutamate by the enzymes GSH1 and GSH2(CM 2001). We annotated members of this pipeline in *C. cinerea* based on orthology information to components of the glutathione system in *S. cerevisiae*(Pócsi et al. 2004; Sato et al. 2009)( Table 39). We identified single-copy orthologs of two proteins in the glutathione synthesis pathway, glutathione reductase, glutathione peroxidase. Glutaredoxins are present, on average, as 5-6 distinct genes in Agaricomycetes genomes, except for *Pt. gracilis* and *M. kentingensis*, which harbor 9 and 11 glutaredoxin genes in their genomes, respectively. Unlike the other antioxidant systems, the expression patterns in the glutathione system show only moderate conservation across species, being developmentally regulated in 3-6 species (Table 39).

**Glutathione-S-transferases** are also involved in defense against oxidative damage. They represent a diverse gene family also involved in other cellular detoxification tasks, such as the neutralization of xenobiotics, drugs or pesticides(M et al. 2009b). Leon-Ramirez et al hypothesized that in *U. maydis* ‘basidiocarps’ glutathione-S-transferases are involved in the protection against oxidizing intracellular conditions generated by some enzymes, such as NAPDH oxidases(CG et al. 2017).Therefore, we only marginally treat them under this chapter, noting that any glutathione-S-transferase orthogroup, also by virtue of the lack of clear orthology to functionally characterized genes from model species, could be related to any of the possible functions of this family, not necessarily protection from oxidative damage. Glutathione S-transferases are a diverse gene family in fungi, with 20-49 genes (on average 32) in the genomes of Agaricomycetes. In each of the species, ∼50% of glutathione S-transferase encoding genes (up to 70% in *Coprinopsis*) were developmentally regulated, indicating that these play important roles in fruiting body development, however, due to the general lack of conservation of these genes, these did not form CDE orthogroups. Interestingly, each species had genes that showed a marked upregulation (FC>4) at the initiation of fruiting body development.

#### 4.14.3. Unannotated genes

Genes that encode proteins with no known conserved domain (Pfam or InterPro) signatures, referred to as unannotated genes here, are abundant in fungal genomes and represent some of the most exciting gene families. At the same time, they represent a challenge to link to fruiting body development, due to the complete lack of functional clues. In general, few literature reports are available on unannotated genes, possibly due to the difficulty of tackling their functions. An exception from this is a report of the enrichment of unannotated genes in mature peridioles of *Pisolithus microcarpus* fruiting bodies(Pereira et al. 2017). An example of unannotated genes that turn out to be developmentally relevant is the LPMO-like X325 gene family(Labourel et al. 2020), which was circumscribed just recently and appears to be related to the cell wall (see chapter 4.8.2.1).

We identified 176 CDE orthogroups of unannotated genes (Supplementary Table 6). These included, for example, *Spc14* and *Spc33* of *S. commune*, which are involved in the formation of the septal pore cap(Van Peer et al. 2010), are unannotated proteins but form conserved, single-copy gene families in the Agaricomycetes (represented by *C. cinerea* 419716 and 473658) that was developmentally regulated in 10/6 and 12/5 species, respectively. In *C. cinerea* alone, we identified 32 FB-init unannotated genes, several of which showed very significant fold changes (up to 150-200x).

This category of genes included *Ctg1* of *L. edodes* and *Cc.Ctg1* of *C. cinerea*. Ctg1 was originally identified in *L. edodes* as a target of the pre-mRNA splicing factor Cdc5 and was later shown to influence stipe elongation in *C. cinerea*(Nakazawa et al. 2008). In the latter study knockdown experiments were unsuccessful, whereas overexpression *C. cinerea* mutants showed accelerated stipe elongation(Nakazawa et al. 2009). In our data *Ctg1* orthologs formed a CDE orthogroup (represented by *C. cinerea* 528942) which was developmentally regulated in 10/8 species (FC>2/4)(Supplementary Table 6). Despite the conservation of developmental regulation, expression profiles were only partially conserved across species. For example, the gene was upregulated in mature fruiting bodies in *C. aegerita, A. ampla, C. cinerea, L. edodes, P. ostreatus*, whereas in other species (e.g. *A. ostoyae*), it showed upregulation in primordia relative to vegetative mycelium. Overall, CTG1 orthologs seem to be a very exciting group of developmental genes, with no conserved domains and no mechanistic information on how they might contribute to fruiting body formation. Its role in stipe elongation needs to be conserved in other species, if expression profile conservation correlates with phenotype, we may expect other phenotypic effects in other species.

In summary, unannotated genes may contain very interesting, novel gene families involved in fruiting body development, yet circumscribing their functions remains challenging at the moment and will require extensive genetic manipulation techniques.

#### 4.14.4. Functionally poorly known genes

In this group of genes we placed ones that had some (mostly automatic) annotations, but we could not clearly link them to cellular or fruiting-related processes. Thus, this category differs from unannotated genes in that unannotated genes are completely devoid of all (even automated) annotations. Altogether, we assigned 182 CDE orthogroups to this category (Supplementary Table 7). Some of these genes are discussed briefly below, but we note that many more interesting, fruiting-related genes are potentially hidden among the 182 orthogroups, however, clarifying their roles is left for future studies.

**Cytochrome p450 family** - We discuss cytochrome p450s in this category, because they participate in diverse functions, from secondary metabolite biosynthesis (as part of gene clusters, e.g.(Nofiani et al. 2018)) to unknown cellular pathways mediating stipe elongation(Muraguchi and Kamada 2000). We detected 8 CDE orthogroups of cytochrome p450s, most of which are functionally unknown. Among the cytochrome p450s, *C. cinerea* Eln2 should be mentioned. This gene was identified associated with a dominant mutation that confers an elongationless stipe phenotype(Muraguchi and Kamada 2000). Eln2 was reported to be a constitutively expressed cytochrome p450. In accordance with this, we did not find considerable expression dynamics on the orthogroup containing eln2 (represented by *C. cinerea* 381558). However, the developmentally regulated orthogroups may be worthy targets of functional analyses in the future.

**GT25 family** - This family includes enzymes with diverse activities (lipopolysaccharide β-1,4-galactosyltransferase, β-1,3- or β-1,2-glucosyltransferase). In bacteria, GT25 are responsible for the biosynthesis of the membrane component lipopolysaccharides(Bertani and Ruiz 2018) (‘endotoxins’). Most known GT25 enzymes are of bacterial origin, we found no functional study of fungal genes to date. The examined Agaricomycetes had mostly a single gene encoding GT25, which showed sporulation-associated expression peaks in most species: young fruiting body gills in *C. cinerea*, fruiting body gills in *P. ostreatus*, fruiting body cap in *L. edodes* and *C. aegerita*, mature fruiting bodies in *A. ampla* and *S. commune*, etc.. GT25 encoding genes showed very low expression in general, in many cases only detectable in the peak expressing stage/tissue. The broader orthogroup (*C. cinerea* protein ID: 501720) that contains the detected GT25 genes is specific to the Basidiomycota (result not shown), which could indicate bacterial origin into the phylum. The function of these proteins in fruiting bodies is currently unknown, however, expression peaks coincident with sporulation suggest they are related to late developmental events.

**UstYa-like mycotoxin biosynthesis proteins** - this gene family was first reported to be involved in the synthesis of the ribosomally synthesised peptide toxin ustiloxin in the ascomycete *Ustilaginoidea virens*(T et al. 2015). This family of genes was found to participate in the production of other toxic cyclic peptides, collectively named ‘dikaritins’, in plant pathogenic fungi (Ding et al. 2016; Vogt and Künzler 2019). Biochemically, they are oxidases of ribosomally produced peptides, including ustiloxins but also alpha-pheromones of *S. cerevisiae* (collectively Kex2-processed repeat proteins, KEPs)(Umemura 2020). Using a simple domain-based search, we detected 2-16 (28 in *A. ostoyae*) genes in the examined Agaricomycetes. Of these, each species had one to multiple genes that showed an induction (>4fold) in primordia relative to vegetative mycelia (except *C. cinerea*, in which two genes were induced in young fruiting body caps), and high expression in all subsequent fruiting body stages, suggesting that their protein product has an increased abundance in fruiting bodies. Although the role of these genes in the Basidiomycota remains unknown for now, we speculate they may be responsible for the production of toxic compounds, possibly with a role in defense.

**Carbonic anhydrases** (CA) - convert CO_2_ to HCO_3_^-^ which is then used in various ways by eukaryotic cells. CAs have been shown to be involved in fruiting body formation: CA knockout strains of the ascomycete *S. macrospora* showed slower vegetative growth and delayed fruiting body formation or defects in ascospore germination(Elleuche and Pöggeler 2009, 2010). The delay in fruiting body formation suggests a lower rate of HCO_3_^-^ buildup necessary for some as yet unknown downstream processes, that, however, can be made up in longer time. Similar roles of CAs have not yet been reported in Basidiomycota, however, they would be worth investigating as many CAs appeared developmentally regulated in fruiting bodies. We found 2-9 CA encoding gene in the examined Agaricomycetes and that each species had at least one developmentally regulated carbonic anhydrase gene, although clear patterns of expression conservation were not noticeable. We found one CDE orthogroup (*C. cinerea* protein ID: 450047) that was widely developmentally regulated in mushroom-forming fungi (8/5 species at FC2/4), showing diverse expression peaks in different species. It should be noted that CA expression levels potentially react also to ambient CO_2_ concentrations, so some of the expression variation we encountered may be due to the growth conditions used during fruiting.

**CipB gene family -** This category of genes included the *cipB* gene reported from *C. cinerea*(Nakazawa et al. 2009). The *cipB* gene was first identified as an interacting partner of the splicing factor cdc5 in *L. edodes*(Nakazawa et al. 2008). We found that the orthogroup that contained *cipB* orthologs (represented by *C. cinerea* 401749) was developmentally regulated in 10/2 species (FC>2/4), indicating that these genes have widespread although moderate expression dynamics (Supplementary Table 7). Based on conserved domain annotations, *cipB* genes are transmembrane ATP synthases located in the mitochondria; its contribution to fruiting body formation is not known at the moment.

#### 4.14.5. Ferric reductase superfamily proteins and iron metabolism

The Ferric Reductase Domain Superfamily comprises a conserved group of proteins that include ferric reductases and NAPDH oxidases, both of which are located in the plasma membrane(Zhang et al. 2013). Ferric reductases are part of a reductive iron uptake system which, in many model fungi, are involved in maintaining iron and potentially also copper homeostasis(Gaurav Bairwa et al. 2017). These genes have mostly been studied in the context of iron acquisition of pathogenic fungi from their hosts. Agaricomycetes don’t share clear orthology with experimentally characterized ferric reductases of the Ascomycota (reviewed in(Gaurav Bairwa et al. 2017))(cite Hegedus et al 2021). Nevertheless, we identified eight ferric reductase orthogroups, of which two were developmentally regulated in multiple species (Table 40), suggesting that ferric reductases have conserved roles in fruiting body development, although their function in the process is unknown. One of these was developmentally expressed in 9/5 species (represented by *C. cinerea* 354442, FC>2/4) contains genes orthologous to *S. cerevisiae* FRE7, which might be involved in copper homeostasis in *S. cerevisiae*(C et al. 2000). Another orthogroup, which contains orthologs of *A. nidulans NoxA* and *N. crassa* nox-1 (represented by *C. cinerea* 260659), was developmentally regulated in 5/1 species. This gene is a NADPH oxidase, which has been shown to be involved in developmental signaling and tissue differentiation(Nguyen et al. 2017).

**Table 40.** Summary of developmental expression dynamics of CDE orthogroups of ferric reductase genes across 12 species. Protein ID of a representative protein is given follows by the number of species in which the orthogroup is developmentally regulated at fold change 2 and 4 (FC>2 and FC>4, respectively). Putative function and ortholog in *S. cerevisiae* (if any) are also given. Abbreviations: 0-gene absent, 1-gene present but not developmentally regulated, 2 - developmentally regulated at fold change >2, 4- developmentally regulated at fold change >4. Species names are abbreviated as: Cc – *C. cinerea*, Aa – *A. ampla*, Ab – *A. bisporus*, Ao – *A. ostoyae*, Ca – *C. aegerita*, Lb – *L. bicolor*, Le – *L. edodes*, Lt – *L. tigrinus*, Mk – *M. kentingensis*, Pc – *Ph. chrysosporium*, Po – *P. ostreatus*, Sc – *S. commune*.

| ID | Fold change |  | Putative function | S.<br>cerevisiae<br>ortholog | Species |  |  |  |  |  |  |  |  |  |  |  |
| --- | --- | --- | --- | --- | --- | --- | --- | --- | --- | --- | --- | --- | --- | --- | --- | --- |
|  | >2 | >4 |  |  | Cc | Aa | Ab | Ao | Ca | Lb | Le | Lt | Mk | Pc | Po | Sc |
| 354442 | 9 | 5 | Ferric reductase | – | 4 | 0 | 4 | 4 | 0 | 4 | 4 | 2 | 2 | 2 | 2 | 1 |
| 152927 | 9 | 5 | Ferric reductase | YOR384W | 4 | 1 | 1 | 2 | 4 | 2 | 4 | 2 | 4 | 4 | 2 | 1 |
| 260659 | 5 | 1 | Ferric reductase | – | 1 | 1 | 4 | 2 | 2 | 1 | 1 | 1 | 2 | 1 | 2 | 1 |
| 448679 | 3 | 1 | Ferric reductase | – | 1 | 1 | 2 | 1 | 1 | 1 | 1 | 1 | 4 | 1 | 2 | 1 |
| 461481 | 4 | 0 | Ferric reductase | – | 2 | 1 | 2 | 2 | 2 | 1 | 1 | 1 | 0 | 1 | 1 | 1 |

The Ferric reductase superfamily also includes NADPH oxidases (NOXs)(Zhang et al. 2013), which are reactive oxygen-generating cell membrane proteins involved in differentiation-related signaling processes and tissue patterning(Blackstone). NOX genes and their regulator NoxR were reported to have been lost repeatedly in yeast lineages, underscoring their role in multicellularity(Nguyen et al. 2017). There is evidence for NOXs regulating sexual, but not asexual or vegetative development in *A. nidulans*(T et al. 2003) and *G. lucidum* and there is expression evidence in *H. marmoreus*(Zhang et al. 2015b). We identified two putative NOX orthogroups in Agaricomycetes (*C. cinerea* protein ID: 461481, 260659). A third (represented by *C. cinerea* 152927) was interpreted as a NOX by León-Ramírez et al(CG et al. 2017), but appears to be a ferric reductase. Despite clear reports of NOXs being induced in sexual structures of Ascomycota(T et al. 2003), we did not find evidence for consistent induction of these genes in fruiting bodies, rather, expression patterns vary among fruiting body forming fungi. NOX genes show modest expression dynamics in mushroom forming fungi (2<FC<4, Table 40). *C. cinerea* 448679 appears to be a single-copy homolog of noxR that is conserved among mushroom-forming fungi. Members of this orthogroup are not developmentally expressed.

## 5. A synoptic model of fruiting body development

This study aimed to provide a catalog of conserved fruiting body development genes based on a literature review and a meta-analysis of developmental transcriptomes of 12 mushroom-forming fungi. Based on sequence analyses of genes showing developmentally expression during fruiting body morphogenesis in these 12 species, we identified 921 conserved developmentally expressed orthogroups and several gene families. To more broadly explore pathways linked to these genes, as well as genes that are known to be involved in fruiting but do not show significant expression dynamics, we analyzed further 558 gene families. As a result, altogether 1479 genes of *C. cinerea*, and the orthologs of these in 11 other species are treated in this paper. These allowed us to identify developmental expression patterns that are shared across species. For most of the conserved developmental orthogroups, we provided broad functional classifications and speculations on their probable role in fruiting body development, although elucidating their exact function and molecular mechanisms of their action remains for future studies. In this chapter we synthesize these data into an updated, synoptic model of fruiting body development in the Agaricomycetes (Fig. 15). This model strongly builds on work by previous scholars, who attempted to provide overviews of developmental and/or molecular mechanisms of mushroom formation. In a series of papers Moore provided detailed descriptions of developmental events based on morphological and biochemical analyses(Moore 2005). Later, Kues summarized knowledge on the life history of *C. cinerea* and details of its developmental biology(Kües 2000; Kües and Liu 2000). More recently, de Plaza et al, Ohm et al and Muraguchi et al provided examples of transcription networks active during development and a molecular overview of *C. cinerea* development based on gene ontology, respectively(Ohm et al. 2011; Plaza et al. 2014b; Muraguchi et al. 2015). This paper further elaborates these models with more detail, and a selective presentation of only those processes that, based on gene expression patterns are conserved across twelve Agaricomycetes species. We hereafter discuss developmental events in three phases: early events (primordium formation and cell proliferation), growth by cell expansion and late events (sexual processes, sporogenesis).

**Fig. 15:**
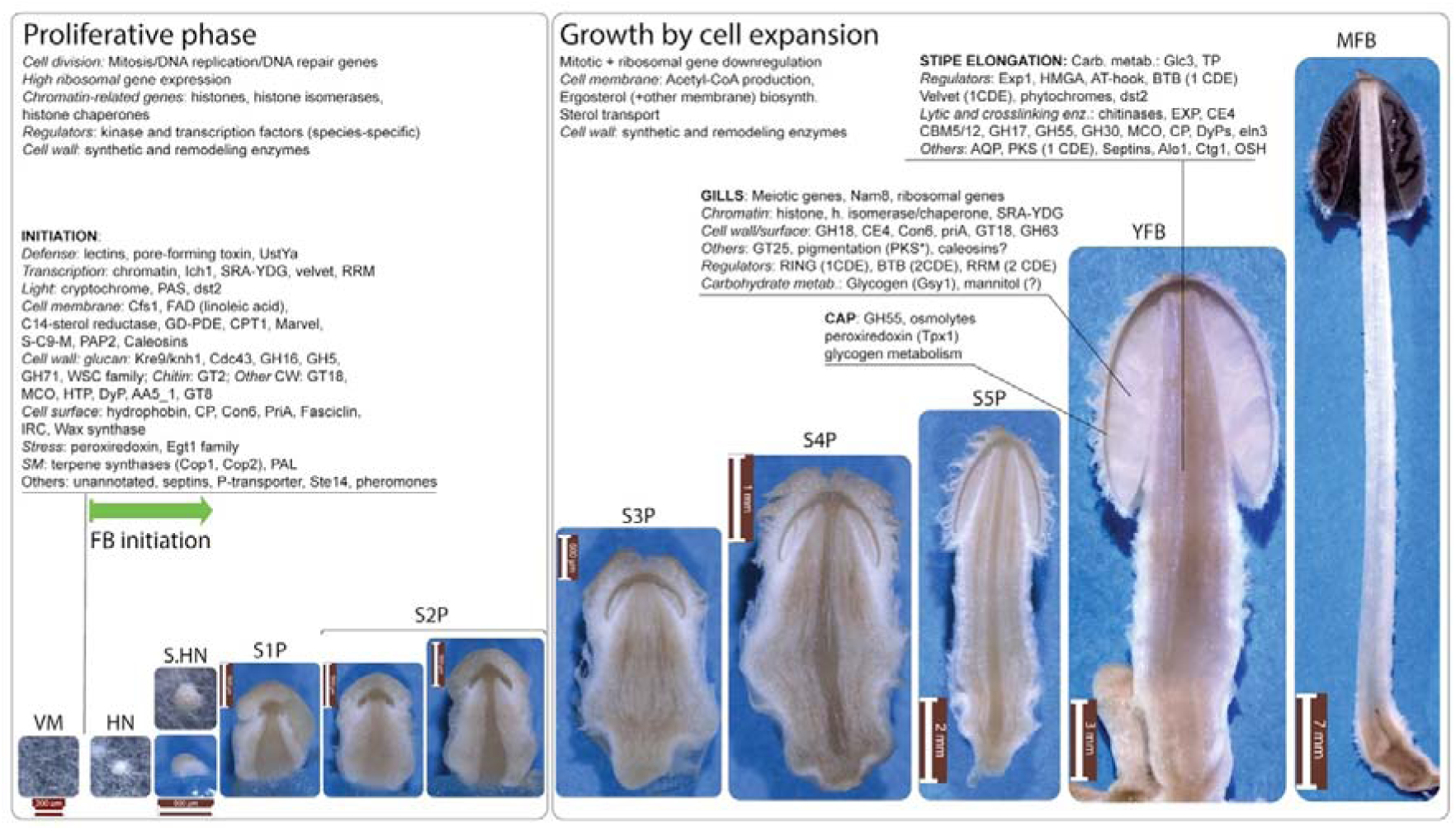
The most important, conserved genes belonging to each phase are shown - genes with limited expression conservation or species-specific expression are not depicted. [footnote: *pigment patterns vary by species] TP: trehalose phosphorylase, PKS: pyloketide synthase; OSH: Oxysterol Binding Protein Homologue; AQP: aquaporins; EXP: expansins; MCO: multicopper oxidase; RRM: RNA recognition motif family proteins; PAP2: diacylglycerol pyrophosphate phosphatases; FAD: fatty acid desaturase; GD-PDE: Glycerophosphodiester phosphodiesterases; S-C9-M: sphingolipid C9-methyltransferases; IRC: intradiol ring cleavage dioxygenase; PAL: phenylalanine ammonia-lyase; SM: secondary metabolism; HTP: heme-thiolate peroxidase; CP: cerato-platanin. genes associated with cap autolysis are not shown on this Figure, because it is a species-specific trait of *C. cinerea*. Note that light induction schemes are not shown on this Figure, as this varies from species to species.

### 5.1. Early events: primordium formation and cell proliferation

Expression patterns of developmentally regulated gene groups outline several distinct processes and a temporal sequence of cellular events in fruiting bodies. The first in this sequence and probably the most interesting is the initiation of fruiting body development, which involves the rearrangement of hyphal branching patterns, and the light-dependent formation of a primary, then a compact, secondary hyphal knot(Kües 2000; Liu 2005; Sakamoto et al. 2018). Hyphal knot formation represents a transition from loosely arranged hyphae with fractal dimensions to true complex multicellular structures(Nagy et al. 2018). Fruiting body initiation has been a subject of intense research. Several gene expression changes have been described in this transition, including light-regulated(Sakamoto et al. 2018), defense (e.g. galectins, pore-forming toxins(Plaza et al. 2014b), lectins(Boulianne et al. 2000; Walser et al. 2005)), fatty acid biosynthesis(Liu 2005), transcriptional regulation (TFs, velvet proteins(Ohm et al. 2011; Plaza et al. 2014b)), cell surface (hydrophobins, PriA, fasciclins(S et al. 1992; Y et al. 2007)), cell wall (Kre9/Knh1, laccases(Kües et al. 2011)) as well as others(Liu 2005; Hou et al. 2019). We update these observations by several functionalities in this work.

The largest gene group that shows an upregulation in primordium samples is related to mitotic cell division (See Fig. 3 in chapter 4.1.1 above), which coincides with tissue differentiation in the primordium and produces the final number of cells that makes up the fruiting body. Ribosomal and chromatin-remodeling related gene (e.g. histones) expression show very similar expression profiles in primordia (see Fig. 4 and 10 in chapter 4.1.2 and 4.5.4.1), suggesting that novel protein synthesis and chromatin remodeling are required cell proliferation/division in primordia. Chromatin remodeling might be required for mitotic chromosome condensation and/or for rearranging chromatin accessibility for fruiting-specific trans regulatory factors. We term this the proliferative phase of development, which, in *C. cinerea,* encompasses hyphal knots to stage 2 primordia (Fig. 15). This pattern of mitotic, histone and ribosomal gene expression has been observed in most species, suggesting that fruiting body development can, in general, be divided into proliferative phase and growth by cell division. However, some species (e.g. *A. bisporus*(Craig et al. 1977), see chapter 4.1.3) might not fully conform to this pattern and we expect non-Agaricales species (e.g. Polyporales, bracket fungi) to not follow this rule.

While the vegetative mycelium is entangled in the substrate and sensitive to bacterial and fungal competitors, fruiting bodies grow above ground and are, thus, more exposed to more extreme environmental conditions and biotic stress (animals grazers or infections). Accordingly, It has been reported that mycelial- and fruiting body-expressed defense gene repertoires show little overlap, indicating a dramatic switch in defense strategy upon fruiting(Plaza et al. 2014b). Defense genes generally show little conservation and high phylogenetic patchiness and did not form conserved orthogroups in our analyses. We, therefore, discussed them in the context of broader gene families (for detailed reviews see(Kunzler 2015; Künzler 2018)). Putative secondary metabolism and oxylipin-related genes might also serve defense and/or antimicrobial purposes.

Probably as a response to differences in environment compared to the substrate, such as higher temperature or humidity fluctuations, several cell surface protein-encoding and fatty acid metabolism genes showed induction in primordia. These included hydrophobins, cerato-platanins, wax synthases and the Con6 family, among others. Hydrophobins probably help retaining water or provide protection from excess water above-ground(Lugones et al. 1999b). Wax synthases and Con6 proteins, although haven’t been examined to date, might act in a similar way. We infer changes in cell membrane composition in primordia from expression changes in genes influencing membrane composition. For example, fatty acid desaturases related to linoleic acid biosynthesis are upregulated in fruiting bodies, probably contributing to the higher linoleic acid contents of fruiting bodies reported before(SHAW 1967; Song et al. 2018a), with potentially to oxylipin synthesis too(A et al. 2020)). Putative stress-related gene expression changes (peroxiredoxins and ergothioneine-biosynthesis genes) might also correspond to genetically encoded mechanisms for coping with adverse above-ground conditions.

Cell wall remodeling emerges as a potentially important process in fruiting body initiation both in this study and previous ones (e.g.(Liu et al. 2021)). Fruiting body-specific cell wall architectures exist and are probably produced by a diverse suite of lytic and cell wall remodeling enzymes that are induced in primordia (Fig. 15). From the array of conserved orthogroups, it appears that glucan remodeling is more important than chitin modification. Additionally, oxidative enzyme genes (HTPs, DyP, MCOs, AA5_1) seem to be important in primordium development, although their functions have not been elucidated, yet. Cell wall related gene families show conservation at the gene family level, not always at the level of strict CDE orthogroups. Therefore, the complete array of cell wall remodeling enzymes induced in primordia is much broader in any species than shown on Fig. 15. Several cell wall remodeling related gene families described from the Ascomycota were developmentally regulated in the Agaricomycetes as well (e.g. GH16/17, Kre9/Knh1), whereas some, that are central to Ascomycota cell wall remodeling showed constant expression (e.g. GH72) or are missing in the Agaricomycetes (e.g. GH76). Further, it appears that dynamic expression is mostly observed in cell wall-active enzyme genes, not in upstream (e.g. ER-localized) components of the cell wall assembly line.

Although the light-dependency of fruiting body initiation differs from species to species(Sakamoto 2018), we detected some orthogroups that probably correspond to light reception. These include phytochromes, the *C. cinerea* Dst2 family and potentially PAS domain containing proteins. The latter is particularly interesting, given that homologs in *N. crassa* regulate the white collar complex and mediate adaptation to changing light intensities during the day(Fuller et al. 2015).

The initiation of fruiting body development represents a transition from fractal-like to complex multicellularity. Accordingly, several complex multicellular processes, such as cell-to-cell communication, adhesion and the regulation of differentiation(Knoll 2011; Nagy et al. 2018) should turn on in primordia. Like for previous studies, decoding these aspects proved difficult for us too. We expect regulatory factors, such as transcription factors, kinases, phosphatases, potentially F-box proteins or non-proteinogenic factors (e.g. small RNA) to participate in cellular differentiation. We and previous work detected some transcription factor, kinase, NOX, pheromone, F-box orthogroups upregulated in primordia. However, these generally did not show significant conservation of expression, so the main regulatory mechanisms of cellular differentiation in fruiting bodies remain an enigma. It further remains to be established if differentiation mechanisms described from plants and animals, such as morphogens, hormone-like substances (e.g. oxylipins) or highly conserved gene regulatory circuits exist in fungi. Similarly, adhesion-related proteins are mostly unknown at the moment. Fasciclins and cell wall modification (e.g. by laccases) have been speculated to be related to adhesion(Nagy et al. 2018; Merényi et al. 2020b), though glucan remodeling enzymes, wax synthases, the carbohydrate-binding properties of lectins or species-specific mechanisms could also be key and more research is needed to elucidate what genes glue hyphae together in 3-dimension fruiting bodies.

### 5.2. Transition to growth by expansion

At the stage 2 primordium stage (following nomenclature in *C. cinerea*, Fig. 15), cell and tissue differentiation, cell proliferation has largely completed. This is reflected in the downregulation of mitotic, ribosomal and chromatin-remodeling genes. At this point, the fruiting body transitions to a different developmental mechanism, which we term ‘growth by cell expansion’. Growth by cell expansion is a well-known fungal trait and was already noted by de Bary in 1887(Bary et al. 1887), however, the molecular details have remained unknown. The phenomenon is marginally similar to fruit ripening in plants, but the mechanisms are probably completely different, though both involve cell wall loosening, remodeling and cell expansion.

Based on analyses of developmental transcriptomes, we detected what we believe is a plausible molecular mechanism for growth by expansion. This involves the upregulation of membrane biosynthesis genes and the rewiring of basic metabolic pathways for the production of excess Acetyl-CoA (see chapter 4.2), which is the starting point of the biosynthesis of membrane components (e.g. ergosterol). Upregulation of genes involved in sterol transport, ergosterol biosynthesis correlates with the rewiring of basic metabolism. At the same time, aquaporin gene expression increases in stipes, which probably supports water transport driving the expansion of cells via turgor. Turgor manipulation requires an osmolyte, however, despite speculations of trehalose or mannitol serving as osmolytes, its identity remain unresolved; the expression data analyzed here did not provide unequivocal evidence for any given osmotically active molecule in fruiting bodies. Trehalose, glycogen and mannitol metabolism genes are upregulated at this stage, but many of them in gills, which suggest an involvement in spore storage material production, rather than turgor manipulation.

For enabling cell volume increases, the fungus deploys a diverse and mostly species-specific array of cell wall remodeling enzymes. Cell expansion also probably follows different mechanisms in each tissue. For example, according to the model of stipe elongation in *C. cinerea* proposed by Liu et al(Liu et al. 2021), CE4 and GH30 genes may contribute to the rigidification of the cell wall in the stipe base, causing it to be more resistant to loosening by lytic enzymes than the stipe apex, thereby reducing its elongation potential. It is reasonable to assume that similar mechanisms underlie the differential expansion potential of different tissues within the fruiting body. We detected a large suite of conserved CAZyme orthogroups upregulated in elongating stipes (Fig. 15), whereas only one conserved orthogroup (GH55) was found in expanding caps. Since caps are composed of multiple tissues, it is possible that RNA-Seq techniques with higher resolution (e.g. single-cell RNA-Seq) are needed to uncover conserved tissue-specific cell wall remodeling gene expression.

What regulatory genes underlie growth by cell expansion will need to be determined in future studies. We detected conserved orthogroups of several developmentally expressed regulators (transcription factors, RING-type zinc fingers, F-box, BTB and RNA-binding proteins) but most showed tissue-specific expression patterns and no expression profile clearly suggested involvement in regulating cell expansion in general. Interestingly, the Exp1 transcription factor reported to be responsible for cap expansion in *C. cinerea*(Muraguchi et al. 2008b) was more frequently upregulated in stipes than in caps across the examined species. We detected a red light-sensing phytochrome orthogroup upregulated in stipes, which we hypothesize is responsible directing stipe growth away from closed spaces towards the open air.

### 5.3. Late events: meiosis and spore production

The young and mature fruiting body (in *C. cinerea* nomenclature) stages are the scene for most sexual events, including meiosis and spore production. Accordingly, meiotic genes as well as several genes involved in DNA replication/repair, chromatin remodeling, ribosome assembly, which showed a peak in the proliferative phase in primordia, show a very sharp expression peak associated with meiosis (see chapter 4.1.1, 4.1.2 and 4.5.4). We found several conserved orthogroups of BTB/POZ proteins and RING-type zinc fingers with expression peaks in gills; these might regulate key events associated with meiosis. A conserved transcriptional co-repressor, *Cag1* of *C. cinerea* should be mentioned here. Although this gene and its orthologs did not show significant expression dynamics, its deletion caused abnormalities in gill development(Masuda et al. 2016).

Genes associated with storage carbohydrate metabolism were upregulated in gills in most species, which is consistent with a rich body of literature reporting glycogen fluxes in fruiting bodies(Ji and Moore 1993; Kües 2000). These genes included ones related to glycogen mobilization, trehalose and mannitol metabolism. We found evidence for glycogen synthesis (*Gsy1* orthologs), suggesting that this carbohydrate is packaged into spores.

## 6. Outstanding questions related to fruiting body morphogenesis

Despite the advances in understanding cellular events of fruiting body morphogenesis, several significant questions remain open and await further research. We below outline eight key questions that we think are instrumental to develop a better understanding of fungal fruiting body formation and complex multicellularity. This represents a subjective assessment of significant challenges of fruiting body research, there may be other perspectives on remaining key challenges to be solved.

1. Although new information on fruiting body morphogenesis accumulates at a steady pace, several questions related to multicellular organization remain open. Fruiting bodies are complex multicellular structures and thus display complex developmental programs involving tissue differentiation and cell type specification. Besides a developmental program, adhesion and cell-to-cell communication are two key traits for complex multicellularity(Knoll 2011). While we are starting to understand transcriptional networks governing tissue differentiation (see chapter 4.5.1), we still know very little about the genes/proteins mediating hypha-hypha adhesion, communication or whether an extracellular matrix (other than the cell wall) exists in mushroom-forming fungi. Hypotheses have been put forth on adhesion (via laccases or fasciclins), however, these have not yet been tested experimentally. Tissue differentiation results in distinct cell types within the fruiting body, however, our current understanding of cell type diversity in mushroom-forming fungi is limited. All current estimates for cell type diversity are based on morphology, which probably represent underestimates; technologies like single-cell RNA-Seq might provide more insights into cell type classification.
2. Regarding regulatory genes, it is an open question whether ‘master’ regulators of fruiting body development or tissue differentiation exists, similarly to animal and plant developmental regulator systems(Meyerowitz 2002). Developmental processes are often organized modularly and hierarchically so that the resulting modules have a high level of autonomy. Such autonomy of mushroom development, to our best knowledge, has rarely been documented. An exception from this is monokaryotic fruiting, the process of forming basidiocarps on monokaryotic cultures without mating in certain species (e.g. (Herzog et al. 2016)). The fact that fruiting body initiation can bypass mating suggests that the process has a certain degree of autonomy, which might be reflected as modularity or hierarchy in the underlying regulatory networks. Moore et al characterized fruiting body morphogenesis as being made up of developmental subroutines that are somewhat independent of each other and that makes the whole process resilient to a certain degree of developmental imprecision(Loftus et al. 2020).
3. Transcriptional changes during the initiation of fruiting are significant. Most studies with a sufficient number of time points sampled (e.g.(Plaza et al. 2014b; Muraguchi et al. 2015; Sipos et al. 2017b; Almási et al. 2019; Krizsán et al. 2019; Merényi et al. 2021; Zhang et al. 2021d)) (Ruytinx et al in prep) found the largest number of differentially expressed genes at the transition from vegetative mycelium to the hyphal knot/primordium stage. However, whether this transcriptional rewiring event is purely a result of transcription factor activity or whether there are more general changes, for example, in chromatin accessibility or epigenetics, is unknown at the moment. Assays such as ATAC-Seq(Buenrostro et al. 2015) or ChIP-Seq (see e.g.(Vonk and Ohm 2021)) could provide insight into the magnitude of transcriptional versus chromatin-related rearrangement during fruiting body initiation. Similarly, posttranslational modification of proteins may be a significant regulatory mechanism in fruiting body morphogenesis(Pelkmans et al. 2017a). Gene methylation, an important developmental mechanism in animals, seems to be missing in fungi, on the other hand(Bewick et al. 2019).
4. Understanding tissue specificity of genes and encoded proteins is key to uncovering mechanisms of tissue differentiation and predicting gene function. The bulk expression studies and a few studies involving fluorescent protein tagging revealed numerous examples of tissue-specifically expressed genes; however, more systematic studies will be needed to understand tissue-specific expression and the mechanisms that generate it.
5. We identified altogether 176 conserved developmentally expressed orthogroups of unannotated genes. These genes encode proteins that contain no known protein domains, therefore, they lack even automated functional annotations. Unannotated genes are common in fungal genomes and represent an enigma for mycology. Their conservation implies that they have important functions these are, however, unknown in the vast majority of cases (see Spc genes of *S. commune* for an exception, chapter 4.14.3). Therefore, unannotated genes might represent a future treasure trove for developmental biology.
6. Growth by cell expansion is a widely conserved trait of mushroom-forming fungi. Above we provided evidence for this process involving excess Ac-CoA production, the upregulation of membrane biosynthetic processes and cell wall remodeling genes (chapter 4.2, 4.3). However, the identity of the osmolyte that drives turgor increase in fruiting bodies remains unknown and its identification represents a significant challenge for future research.
7. Although not strictly morphogenetic, how the fruiting body is supported by the mycelium is a question with important practical implications in mushroom industry. It is known that the directionality of the transport of goods in the mycelium switches as fruiting body development is initiated(Herman et al. 2020). Also, it has recently been suggested that secondary metabolite biosynthesis during fruiting body development depends also on the mycelium(Orban et al. 2021). Thus, transcriptional and metabolic reprogramming has to affect the entire fungal colony and not only the part where primordia emerge. The feeding and nutrient transport mechanisms as well as regulation of fruiting in relation to nutrient availability are exciting avenues of research, but beyond the primarily morphogenetic scope of the present paper.
8. While useful for discovering shared aspects of fruiting body morphogenesis, the comparative approach used by us, ignores species-specific aspects of morphology, ecological adaptation, gene repertoires and expression dynamics. Non-conserved aspects of development are certainly important, particularly for understanding the biology of individual cultivated species and will deserve further study in the future. However, given the available information, making functional inferences for taxonomically restricted or species-specific genes is even harder than for conserved genes, therefore, we here restrict ourselves to the discussion of the latter.

## 7. Conclusions

The development of mushroom fruiting bodies is one of the most complex morphogenetic processes in fungi. A significant body of previous research has focused on understanding the underlying genetics and has made significant progress in cataloging genes related to the initiation, morphogenesis, defense, pigmentation of fruiting bodies, among others. However, available knowledge is still patchy and lacks key details in multiple aspects, such as conserved events in fruiting body morphogenesis and corresponding genes or key regulators of major differentiation steps.

This study aimed at providing a catalog of conserved fruiting body development genes based on a literature review and a meta-analysis of developmental transcriptomes of 12 mushroom-forming fungi. We identified gene orthogroups that show conserved developmental expression across multiple species. Despite some limitations of the genomic and transcriptomic approach used here, this study provided novel insights (such as gene families, orthogroups, conserved expression profiles) into the morphogenesis of mushroom fruiting bodies. We hope these data reveal some aspects of the core genetic program of agaricomycete fruiting body morphogenesis and will provide a roadmap for functional analyses of fruiting body development. We are aware that the summary provided here is still an incomplete survey of developmental genes in fruiting body forming fungi. Comparisons at finer scales, such as within economically relevant genera, denser sampling of time points during development or at better spatial resolution (e.g. single-cell RNA-Seq) could yield additional insights. Joint analyses of gene expression patterns with gene phylogenies or with gene family copy number dynamics, functional genomics assays, the inference of gene co-expression networks or population genomics (e.g. GWAS analyses) among others, could all provide more detailed views on the synergy between evolutionary changes in gene content, expression regulation and morphogenesis. We anticipate that with advances in functional genomics of Agaricomycetes, it will become easier to conduct such studies and shed light on conserved and species-specific aspects of fruiting body morphogenesis.

## Acknowledgements

We are grateful to all friends and colleagues who, over the years, inspired our work on fruiting body development through collaborations or discussions. We appreciate the help of Levente Karaffa in making sense of Acetyl-CoA production and basic metabolism. We appreciate the permission of Francis Martin and Jason Tsai to use photographs of *L. bicolor* and *M. kentingensis* in Fig. 1, respectively. We are grateful to Annegret Kohler for providing pre-publication access to the transcriptome of *L. bicolor*. This work was supported by the European Research Council (grant no. 758161 to L.G.N.), the “Momentum” program of the Hungarian Academy of Sciences (contract No. LP2019-13/2019 to L.G.N.), the Hungarian National Research, Development, and Innovation Office (contract No. GINOP-2.3.2-15-2016-00052), the Szeged Scientists Academy under the sponsorship of the Hungarian Ministry of Innovation and Technology (FEIF/646-4/2021-ITM_SZERZ, to B.Sz., A. Cs. And L.G.N.) and with the professional support of the Doctoral Student Scholarship Program of the Co-operative Doctoral Program of the Ministry of Innovation and Technology financed from the National Research, Development and Innovation Fund (KDP-17-4/PALY-2021 , to Cs.F.). T.V. was supported by the National Talent Programme (NTP-NFTÖ-21-B-0074) of the Hungarian Government. FH gratefully acknowledges funding from the German Research Foundation (Deutsche Forschungsgemeinschaft, DFG) under grant HE 7849/3-1.

## Table Legends

Supplementary Table 1. Table summarizing ortholog protein ID for all discussed orthogroups in twelve Agaricomycetes.

Supplementary Table 2. Summary of read numbers and alignments in the analyzed fruiting body transcriptomes.

Supplementary Table 3. Summary of developmental expression dynamics of CDE orthogroups of genes related to alternative splicing across 12 species. Protein ID of a representative protein is given follows by the number of species in which the orthogroup is developmentally regulated at fold change 2 and 4 (FC>2 and FC>4, respectively). Putative function and ortholog in *S. cerevisiae* (if any) are also given. Abbreviations: 0-gene absent, 1-gene present but not developmentally regulated, 2 - developmentally regulated at fold change >2, 4- developmentally regulated at fold change >4.

Supplementary table 4. Copy number distributions of genes related to protein ubiquitination and the proteasome in twelve examined Agaricomycetes.

Supplementary table 5. Genes related to protein ubiquitination and the proteasome in *C. cinerea*.

Supplementary Table 6. Summary of developmental expression dynamics of CDE orthogroups of unannotated genes across 12 species. Protein ID of a representative protein is given follows by the number of species in which the orthogroup is developmentally regulated at fold change 2 and 4 (FC>2 and FC>4, respectively). Putative function and ortholog in *S. cerevisiae* (if any) are also given. Abbreviations: 0-gene absent, 1-gene present but not developmentally regulated, 2 - developmentally regulated at fold change >2, 4 - developmentally regulated at fold change >4.

Supplementary Table 7. Summary of developmental expression dynamics of CDE orthogroups of functionally poorly known genes across 12 species. Protein ID of a representative protein is given follows by the number of species in which the orthogroup is developmentally regulated at fold change 2 and 4 (FC>2 and FC>4, respectively). Putative function and ortholog in *S. cerevisiae* (if any) are also given. Abbreviations: 0-gene absent, 1-gene present but not developmentally regulated, 2 - developmentally regulated at fold change >2, 4- developmentally regulated at fold change >4.

## Footnotes

1 We deviaed from this rule where justified.

