## Supplement for "Lessons on fruiting body morphogenesis from genomes and transcriptomes of Agaricomycetes"

for

Published in XXX

2022.

Content:

Supplementary Figures 1.-10.

**Supplementary Fig. 1.** Expression heatmap of DNA replication, repair, mitosis and meiosis related genes in *A. ostoyae, M. kentingensis, P. ostreatus* and the simple *S. commune*. Well-delimited complexes mentioned in the paper are shown separately. Genes are denoted by Protein IDs. Blue and red colors represent low and high expression, respectively.

*Armillaria ostoyae:*

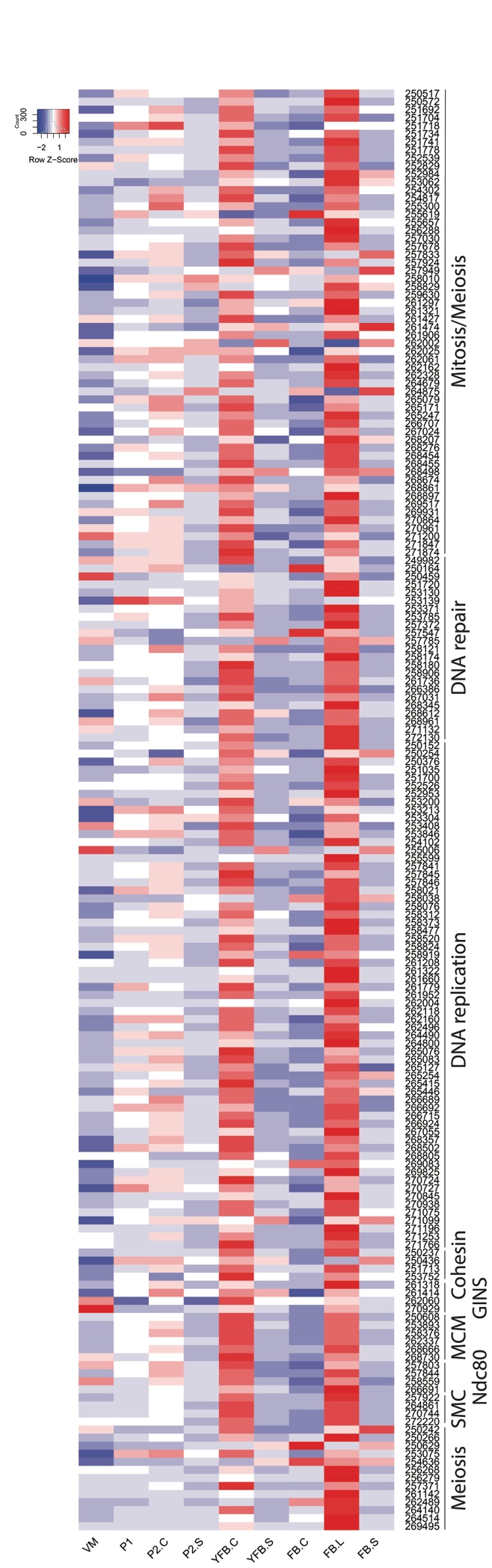

*Mycena kentingensis:*

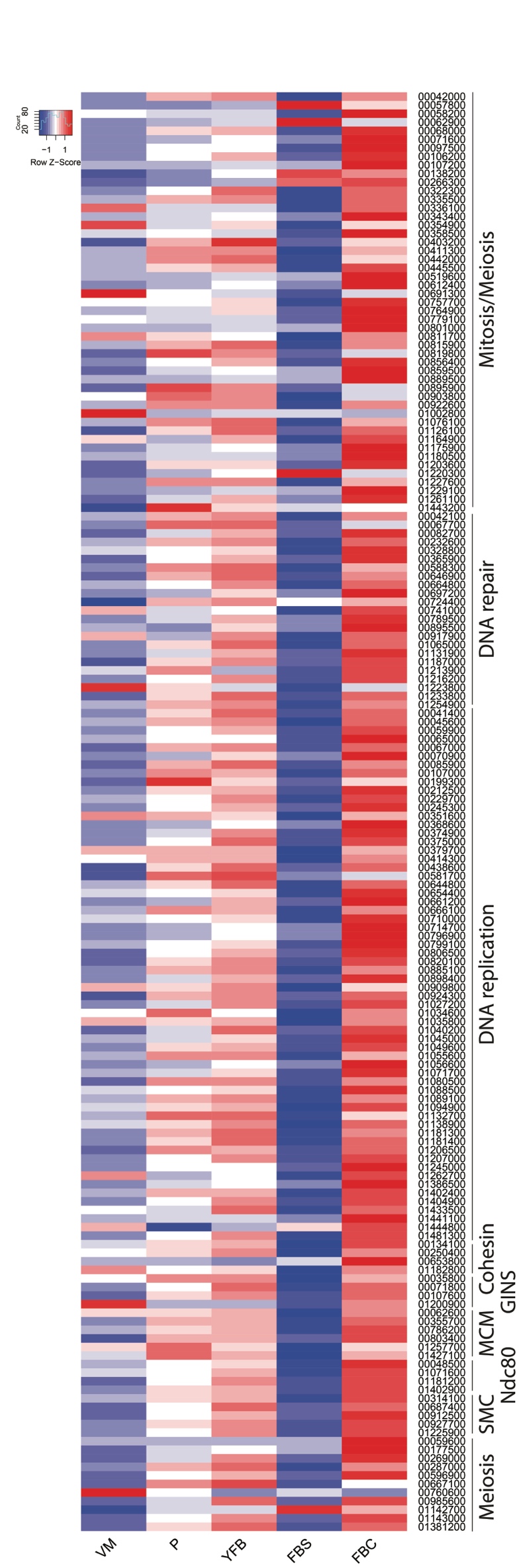

*Pleurotus ostreatus*

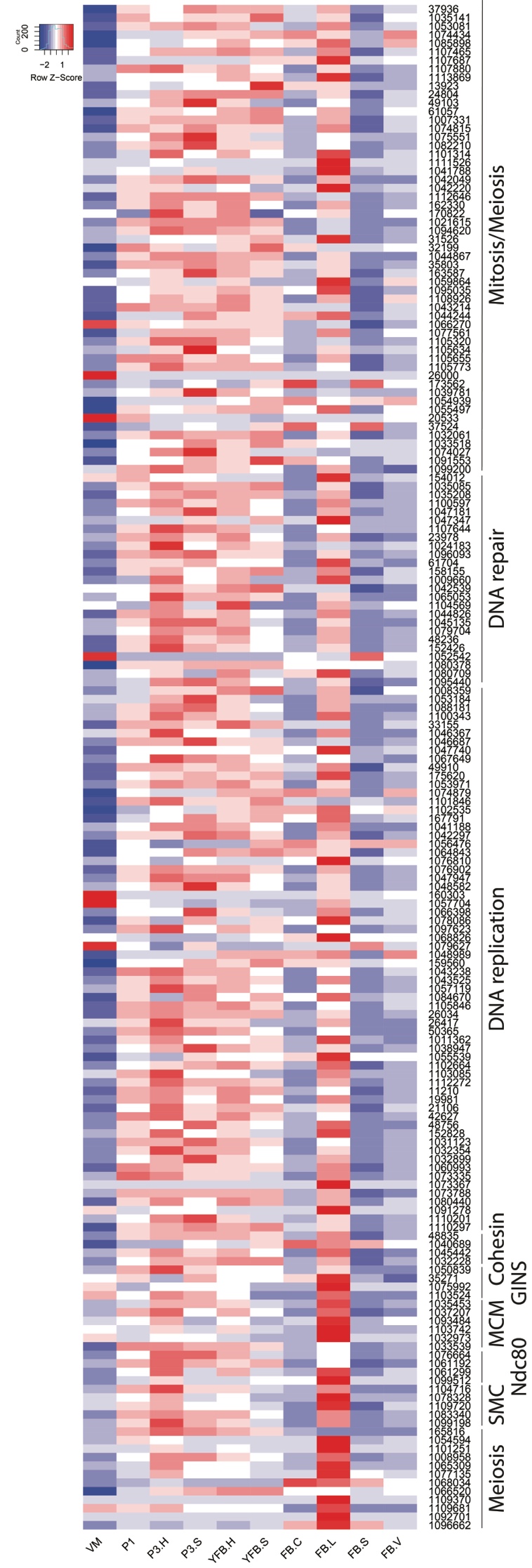

*Schizophyllum commune*

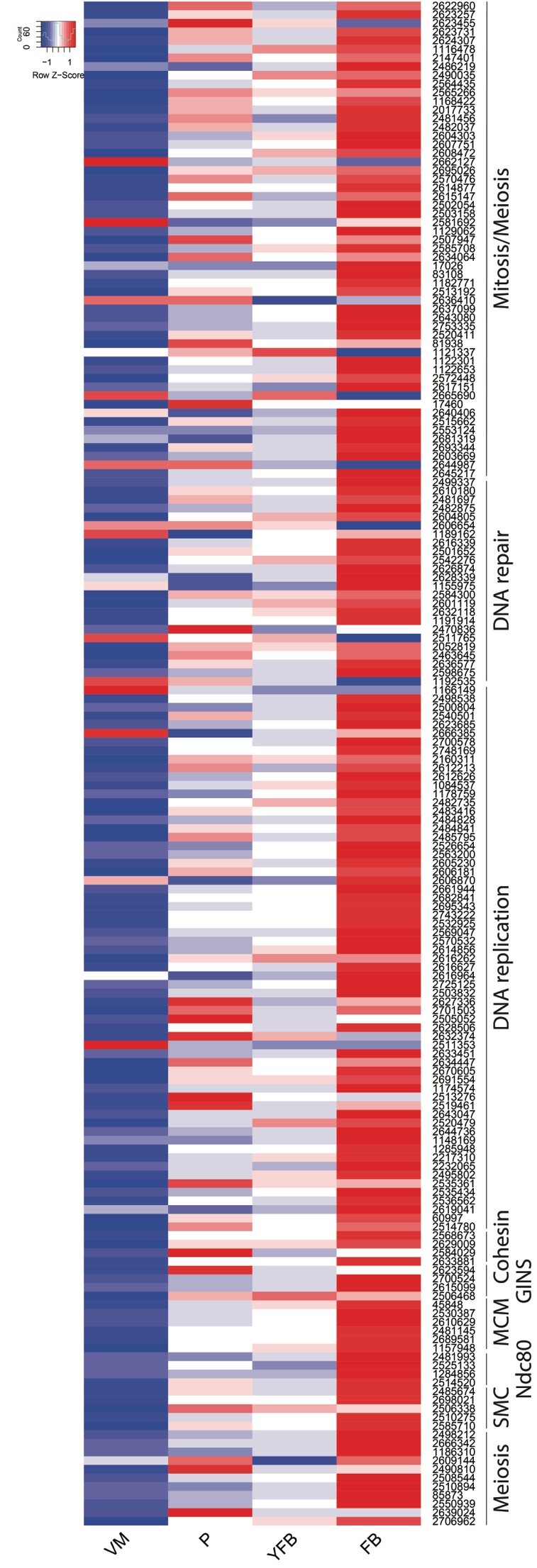

**Supplementary Fig. 2.** Expression heatmap of ribosomal protein encoding genes in *A. bisporus, A. ostoyae* and *P. ostreatus*. Genes are denoted by Protein IDs. Blue and red colors represent low and high expression, respectively.

*Agaricus bisporus*

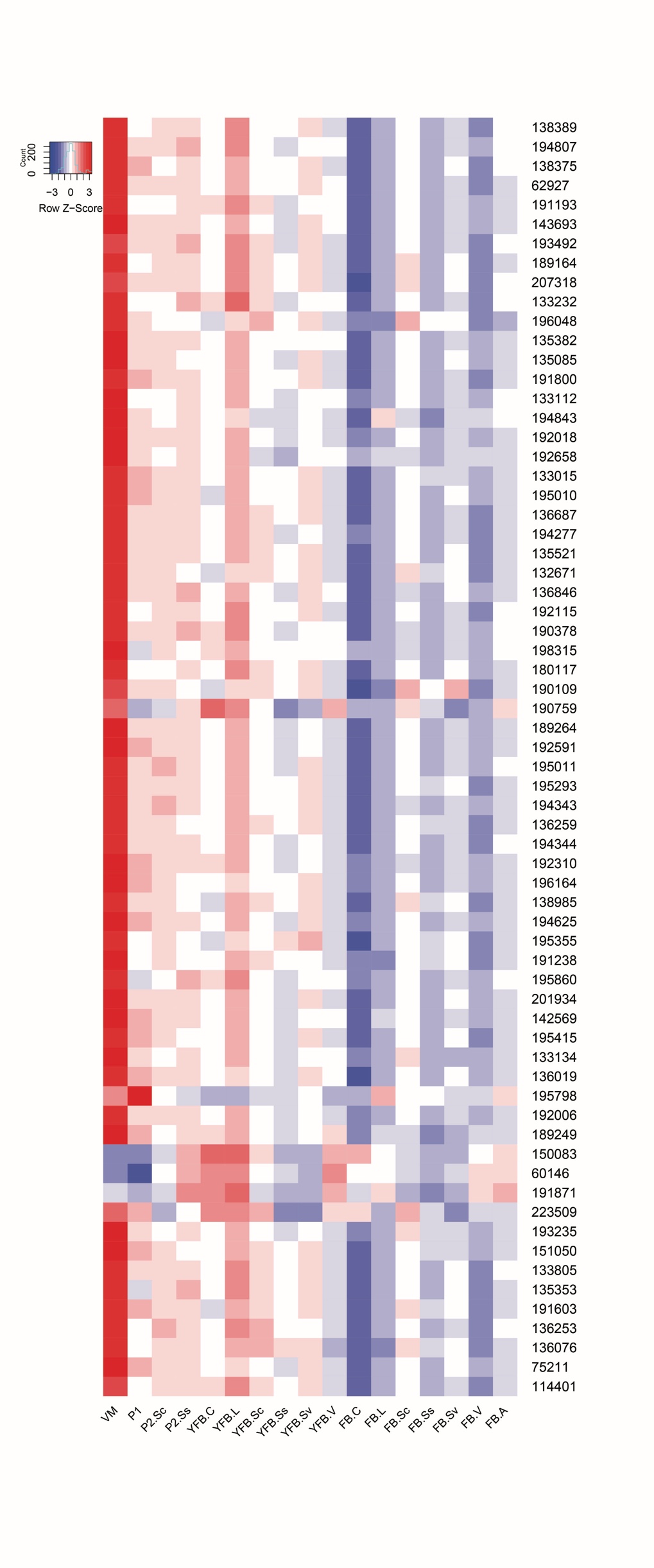

*Armillaria ostoyae*

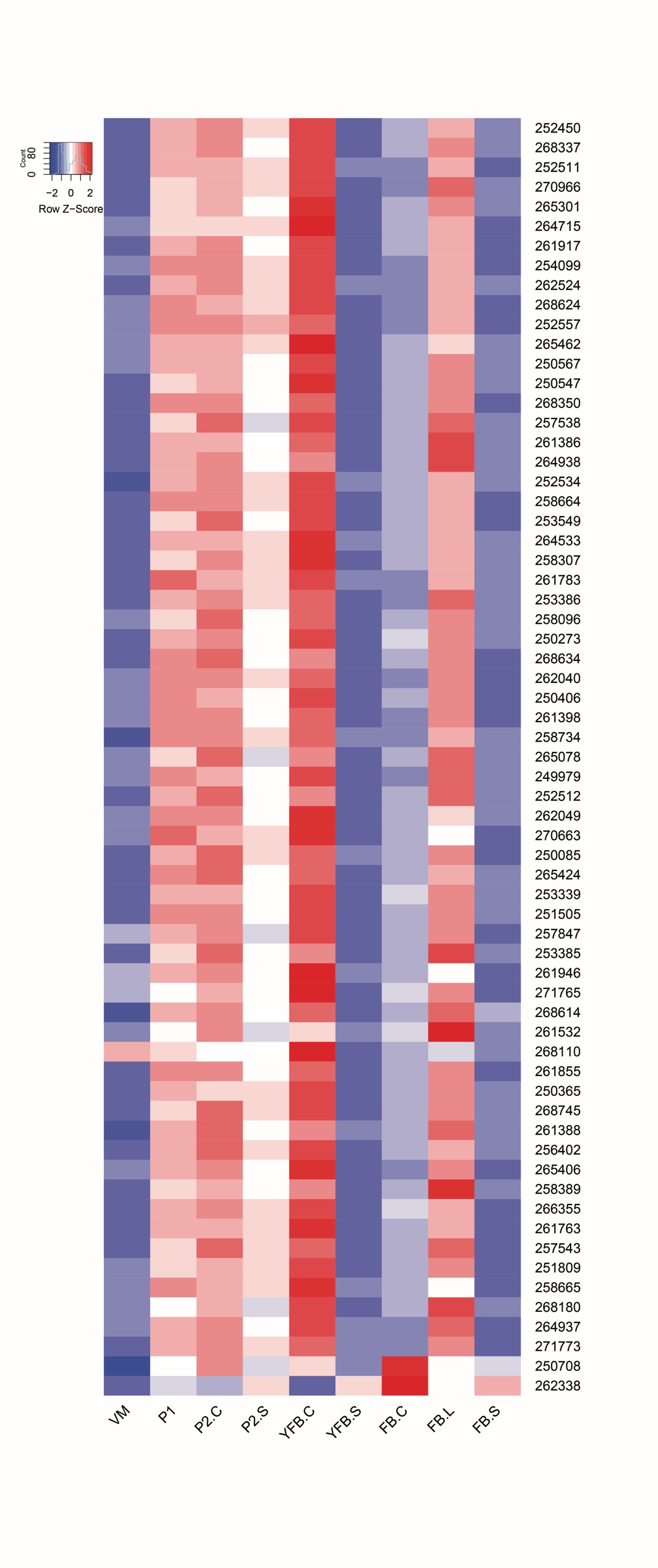

*Pleurotus ostreatus*

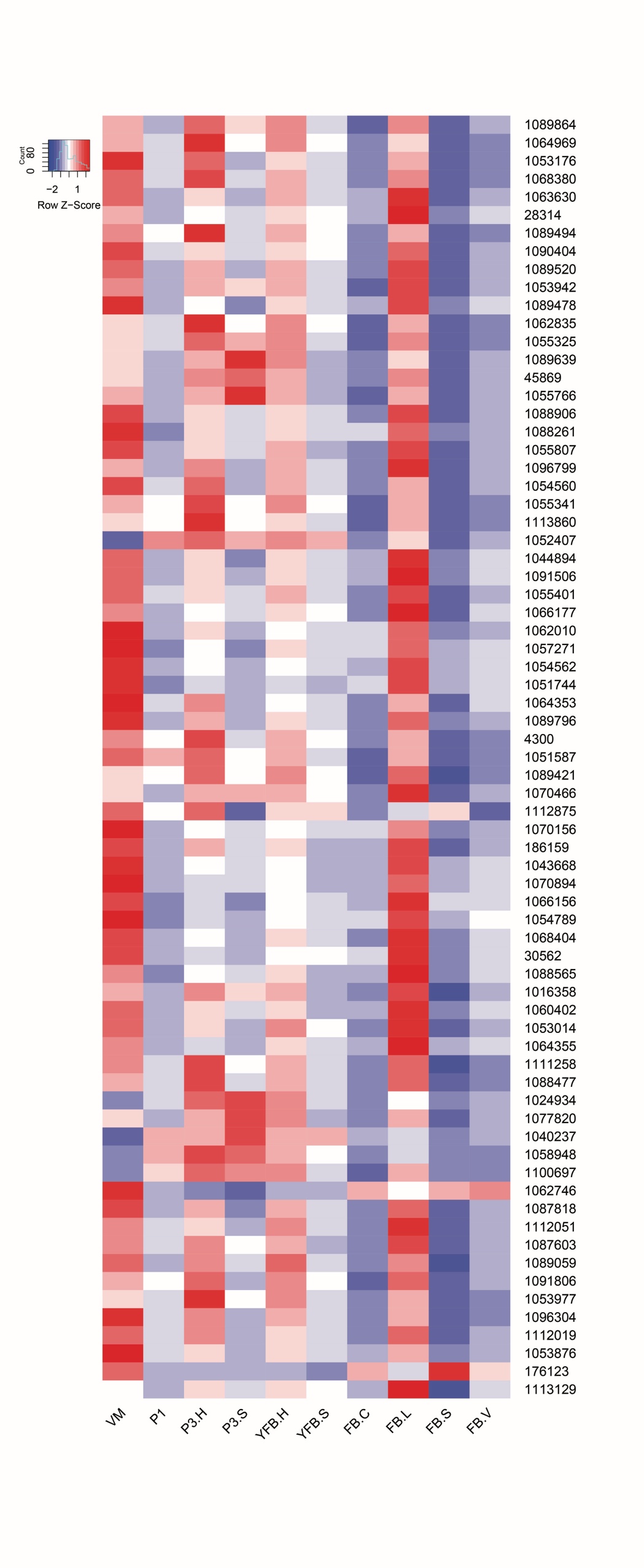

**Supplementary Fig. 3.** Annotated glycolysis pathway of *C. cinerea.* Genes in the pathway are denoted by the *S. cerevisiae* gene name, followed by the protein ID of the *C. cinerea* ortholog and, in parentheses, the number of species in which the gene was developmentally regulated at fold change >2/>4.

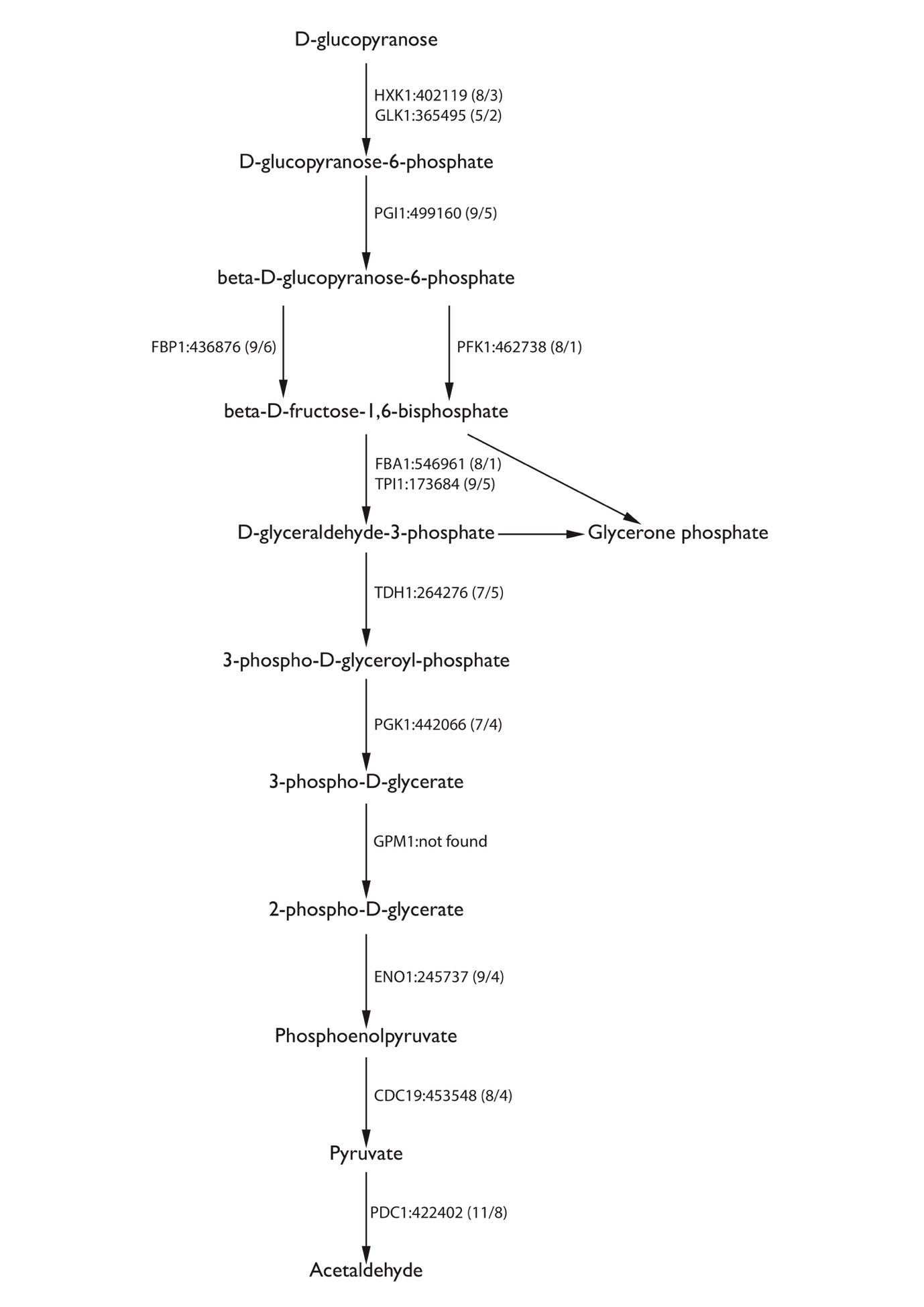

**Supplementary Fig. 4.** Expression heatmap of ergosterol and sphingolipid biosynthesis related genes in *A. ostoyae and P. ostreatus*. Genes are denoted by Protein IDs. Blue and red colors represent low and high expression, respectively.

*Armillaria ostoyae*

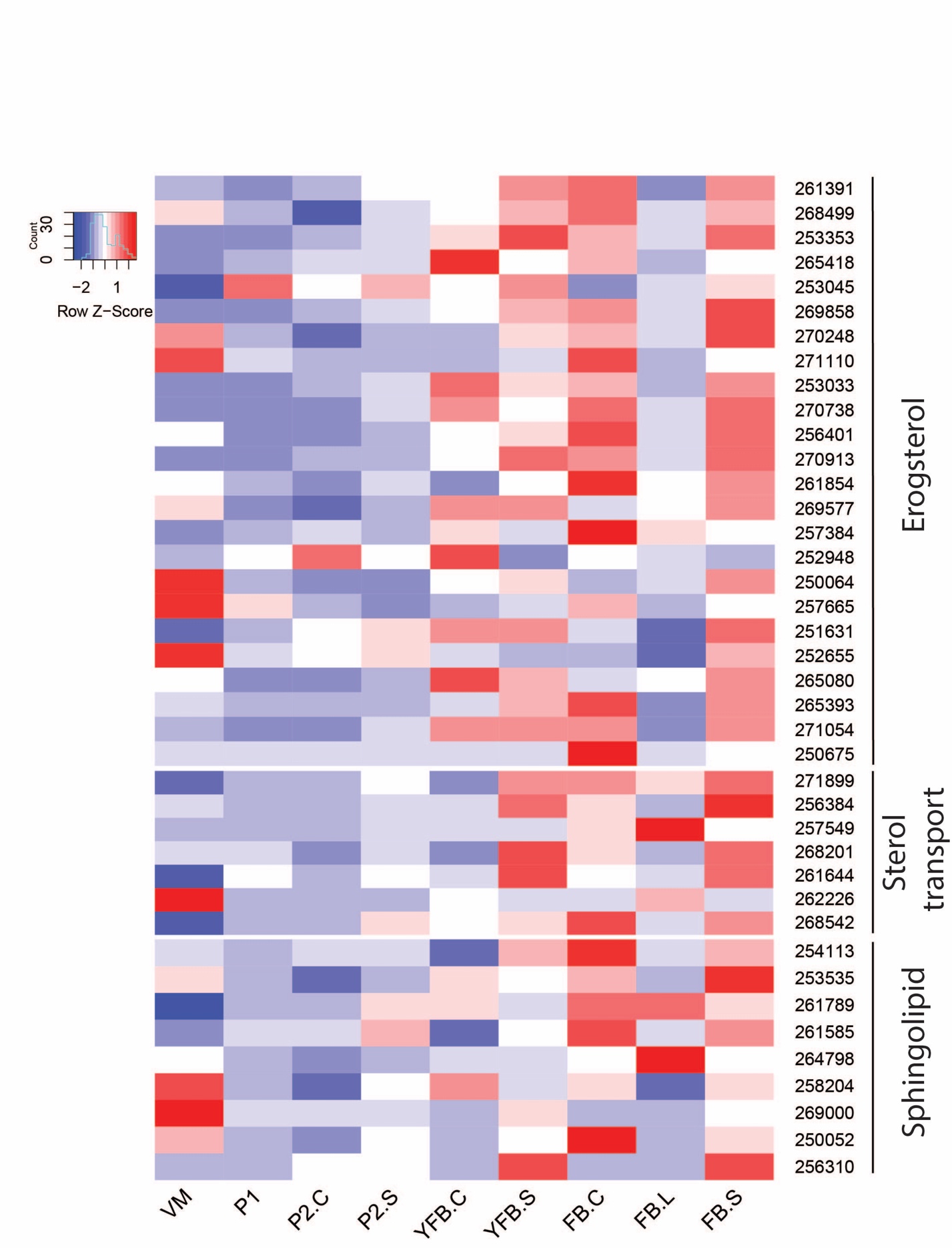

*Pleurotus ostreatus*

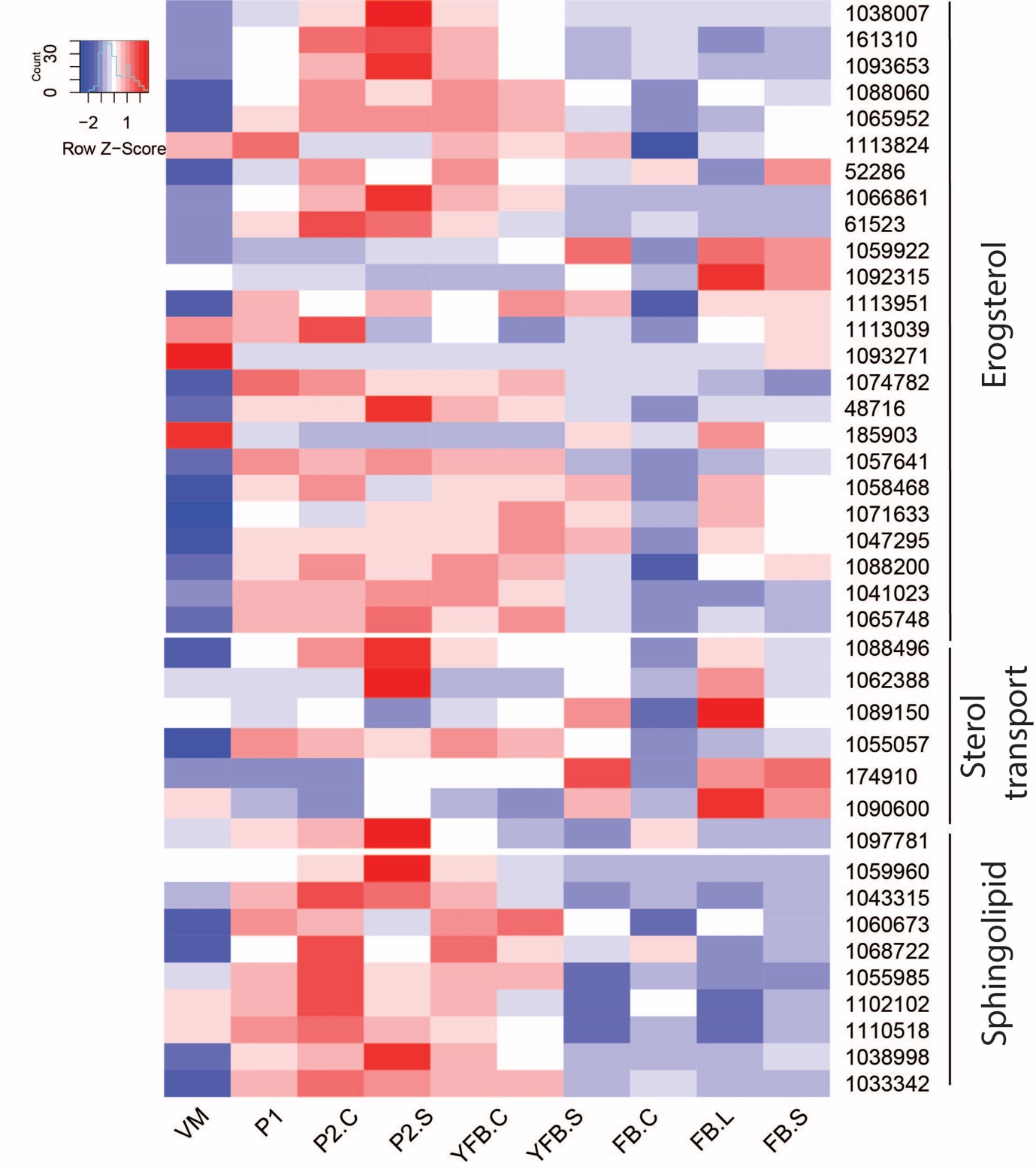

**Supplementary Fig. 5.** Expression heatmap of membrane phospholipid and fatty acid biosynthetic genes in *A. bisporus, A. ostoyae, M. kentingensis* and *P. ostreatus*. Genes are denoted by Protein IDs. Blue and red colors represent low and high expression, respectively.

*Agaricus bisporus*

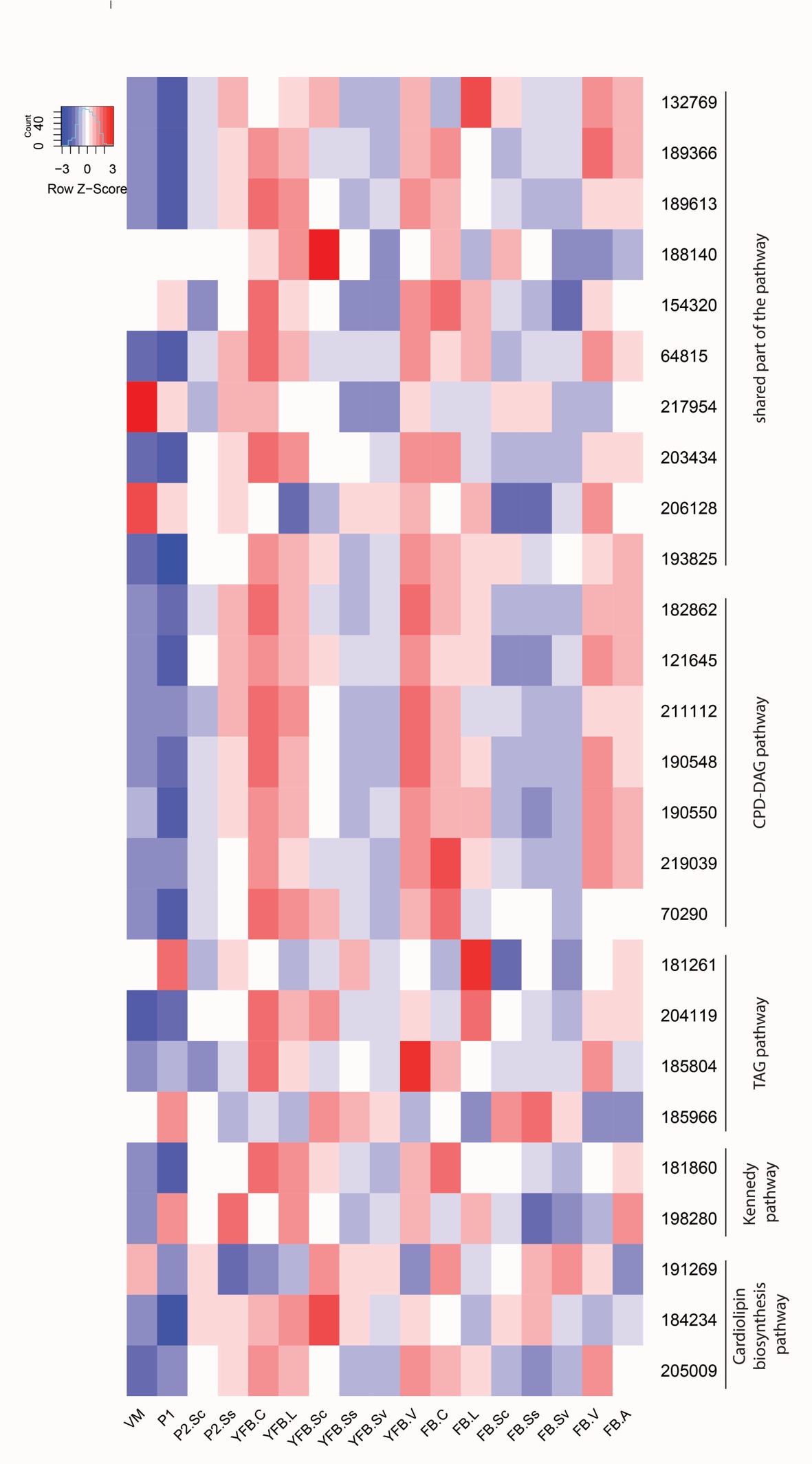

*Armillaria ostoyae*

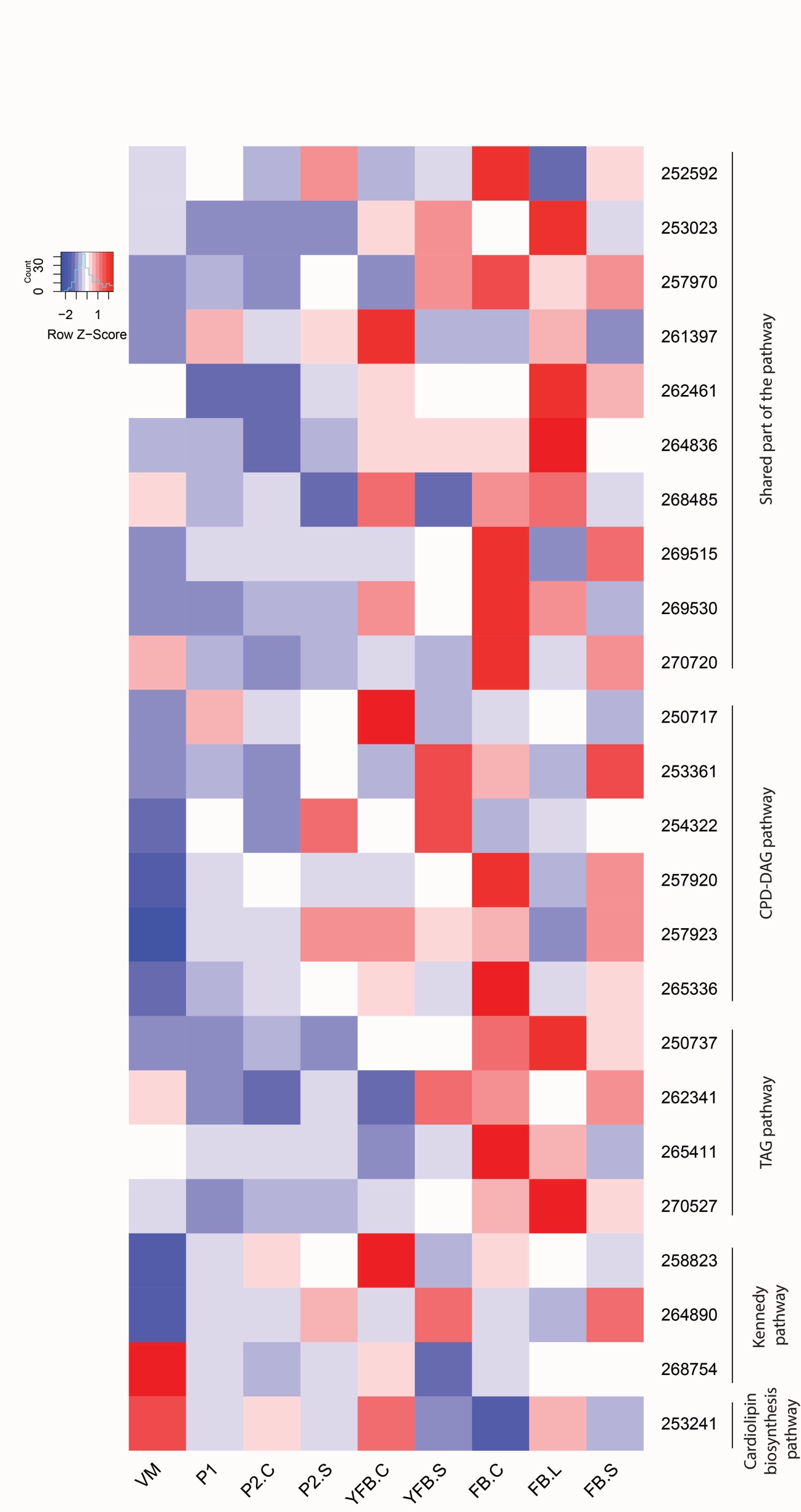

*Mycena kentingensis*

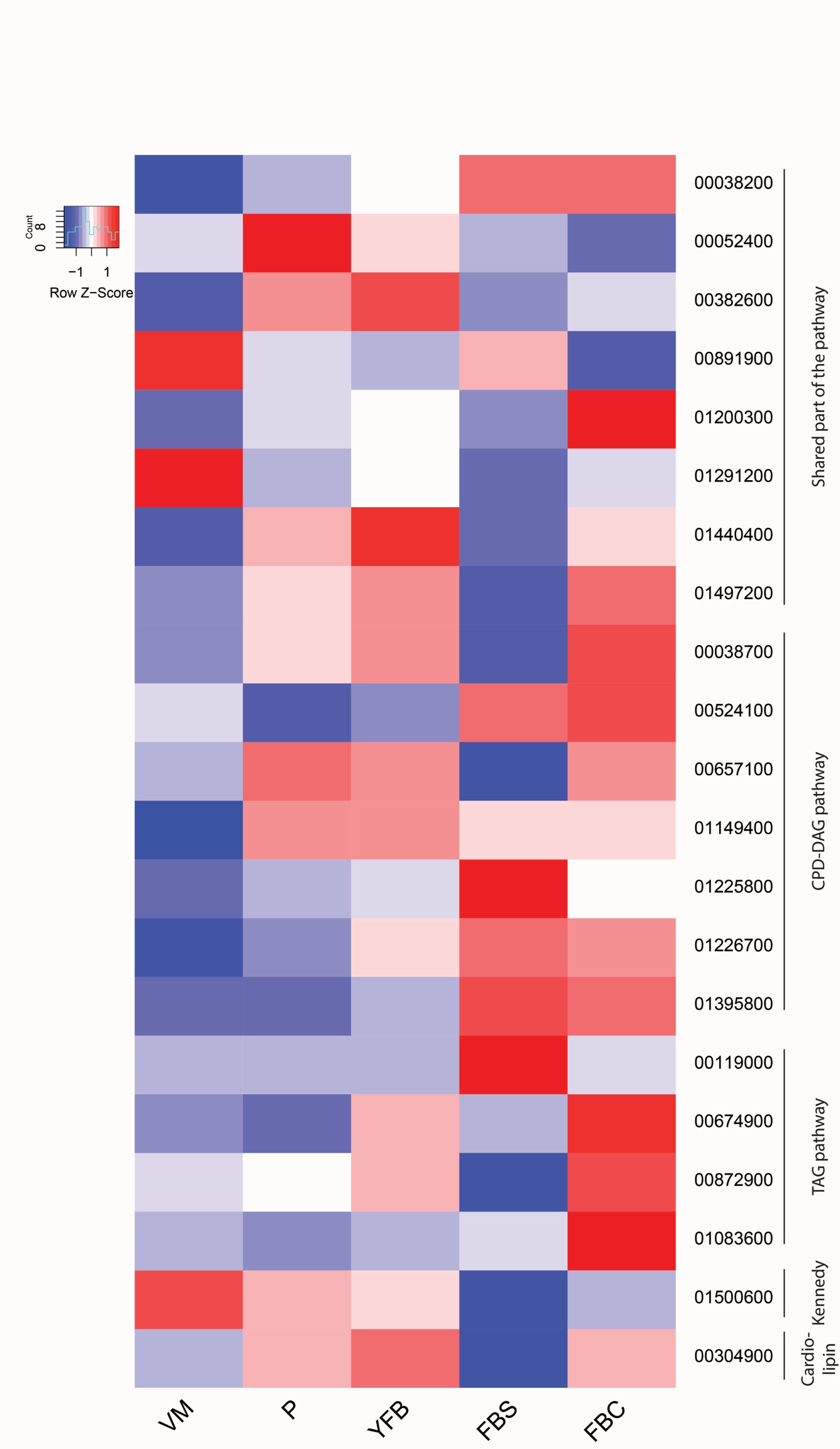

*Pleurotus ostreatus*

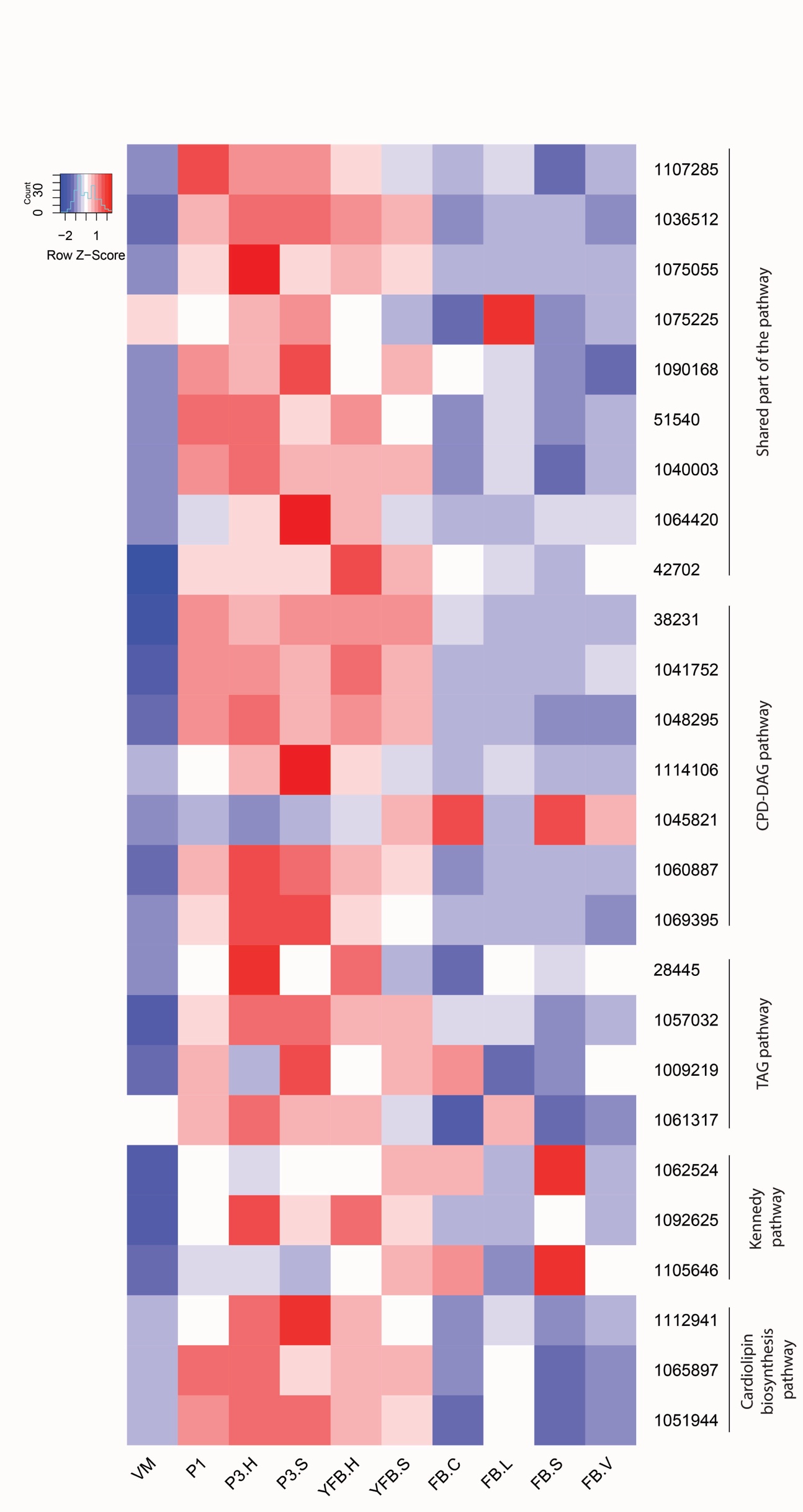

**Supplementary Fig. 6.** Expression heatmap of storage carbohydrate metabolism genes in *A. ostoyae, L. bicolor* and *P. ostreatus*. Genes are denoted by Protein IDs. Blue and red colors represent low and high expression, respectively.

*Armillaria ostoyae*

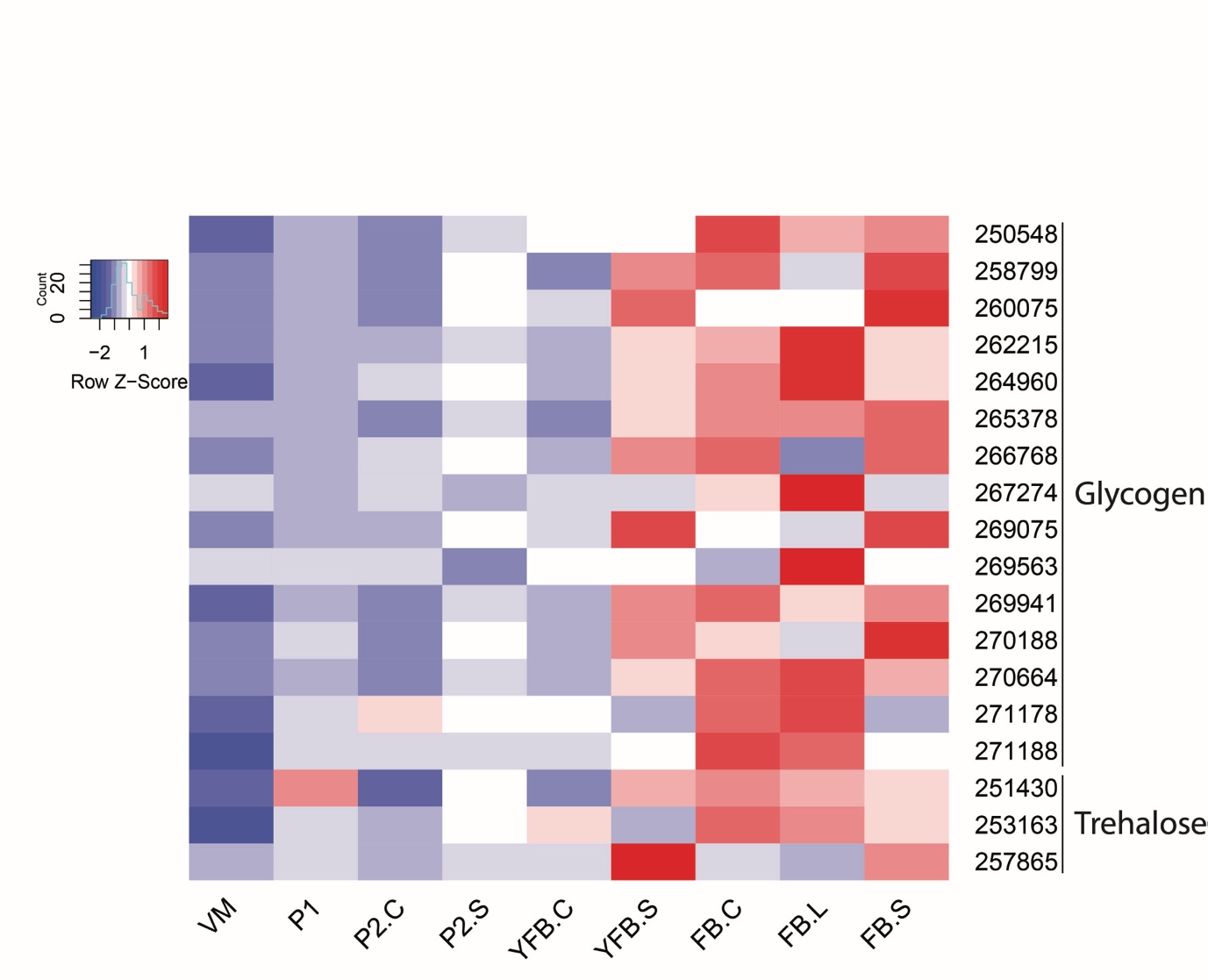

*Laccaria bicolor*

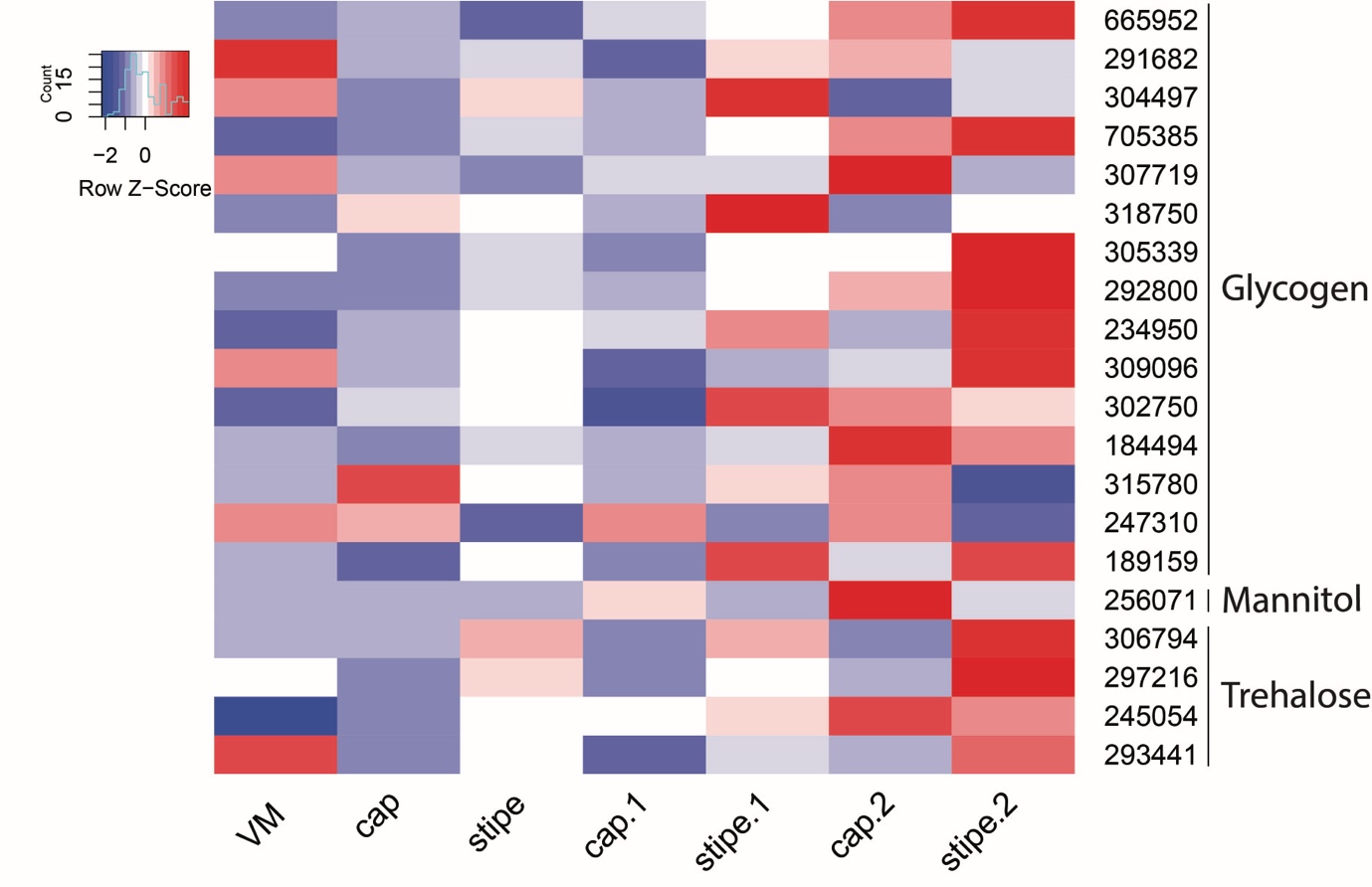

*Pleurotus ostreatus*

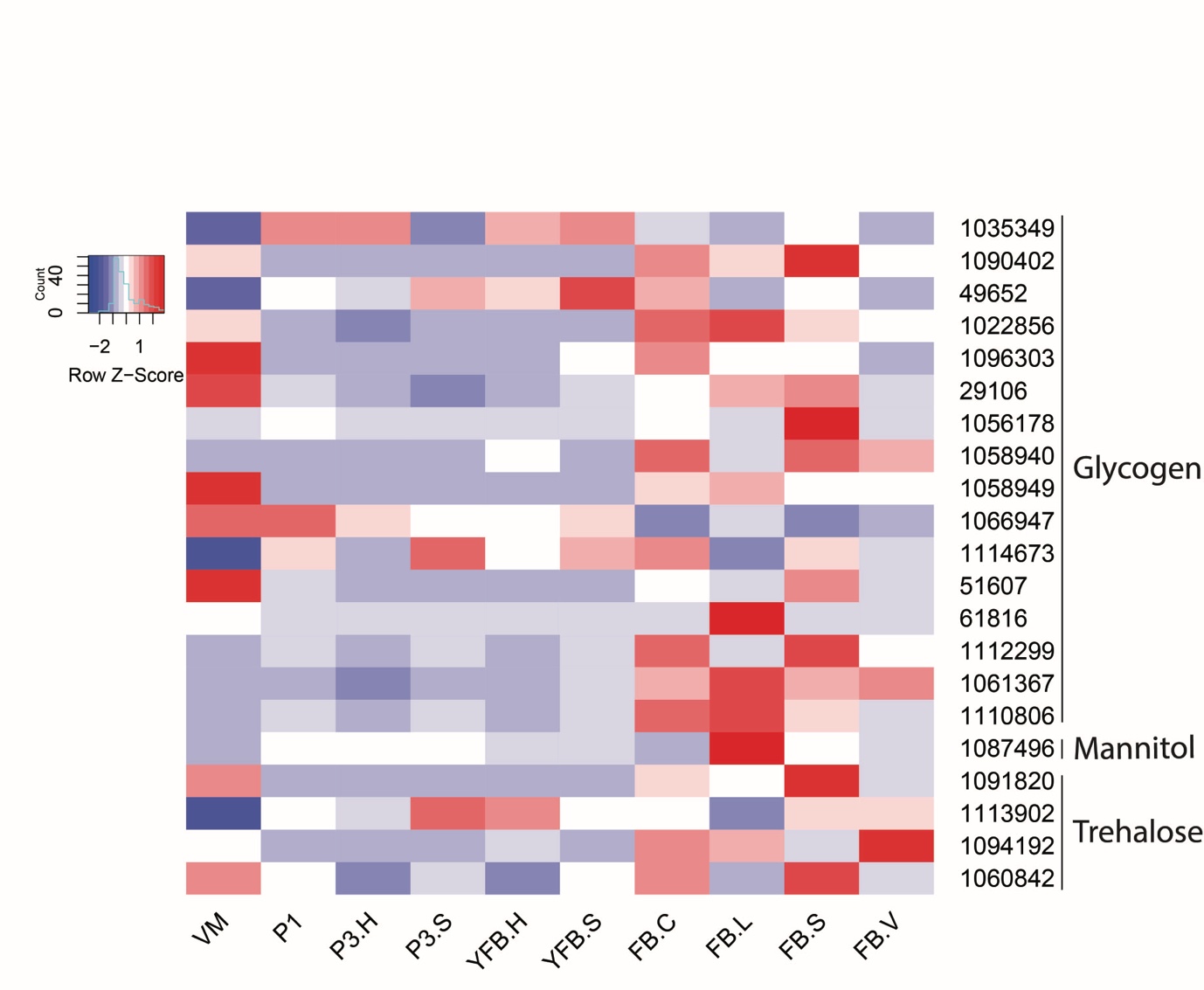

**Supplementary Fig. 7.** Expression heatmap of chromatin remodeling related genes (histones, histone chaperones, histone deacethylases) in *A. ampla, A. ostoyae, L. tigrinus, M. kentingensis* and *P. ostreatus*. Genes are denoted by Protein IDs. Blue and red colors represent low and high expression, respectively.

*Auriculariopsis ampla*

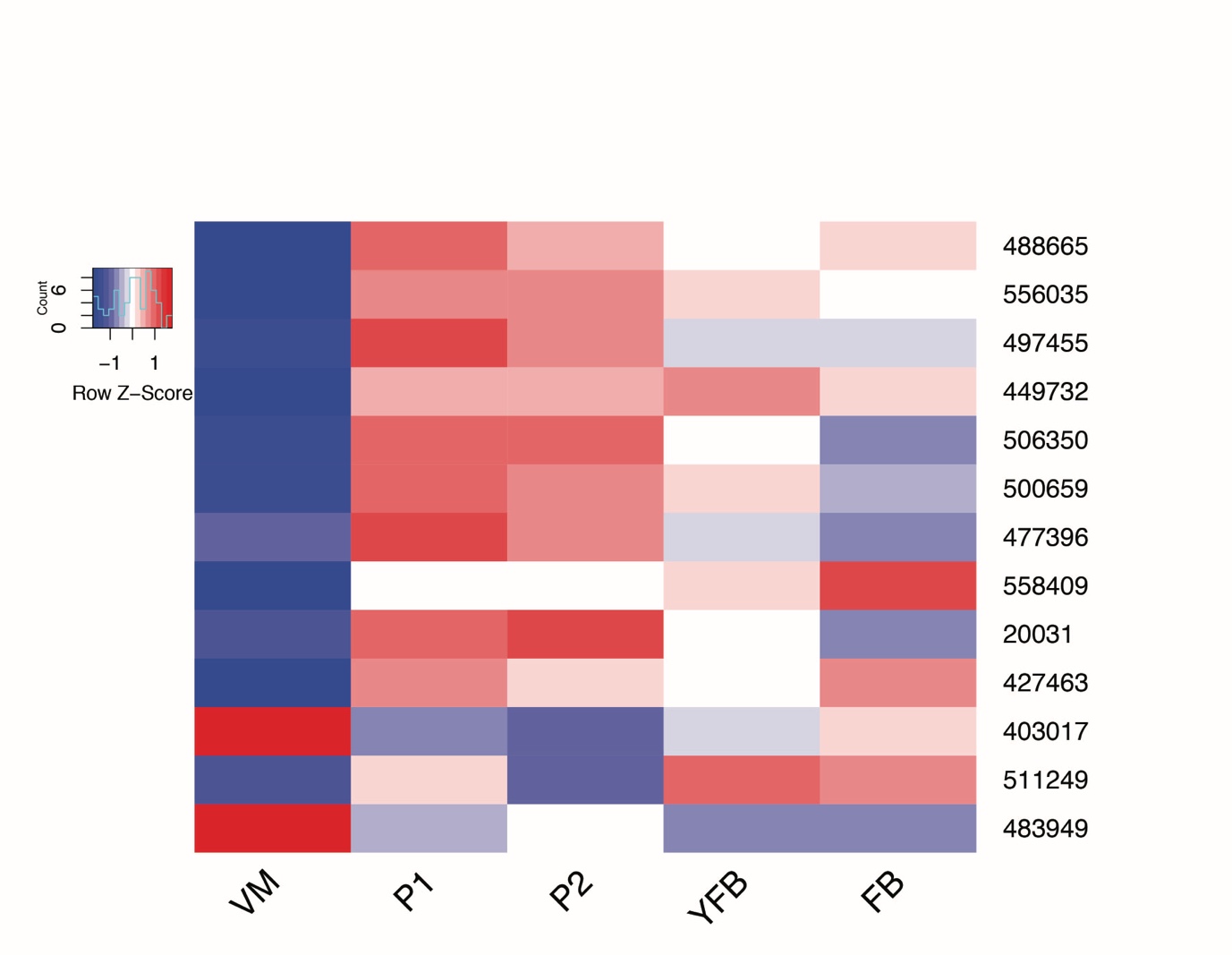

*Armillaria ostoyae*

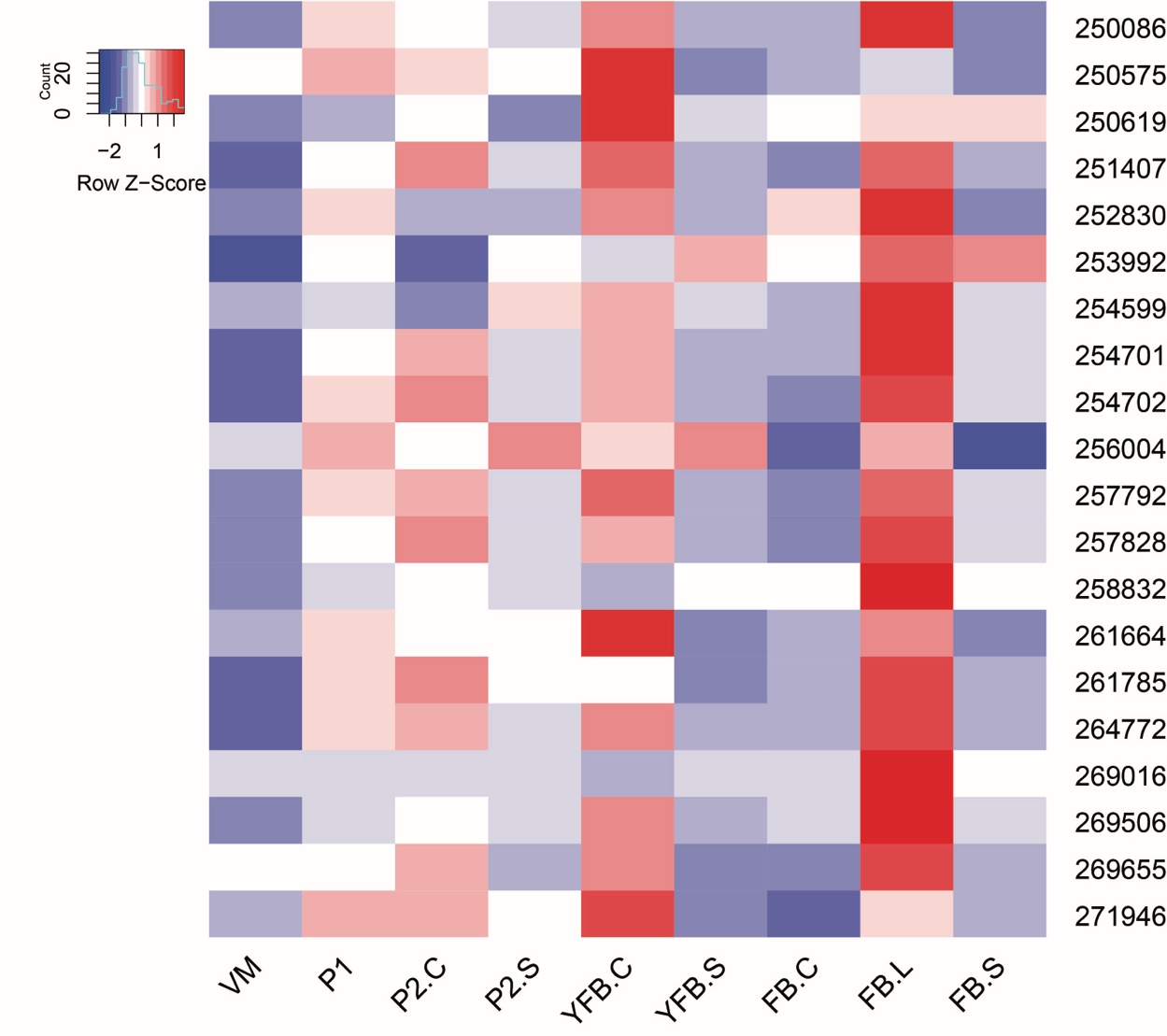

*Lentinus tigrinus*

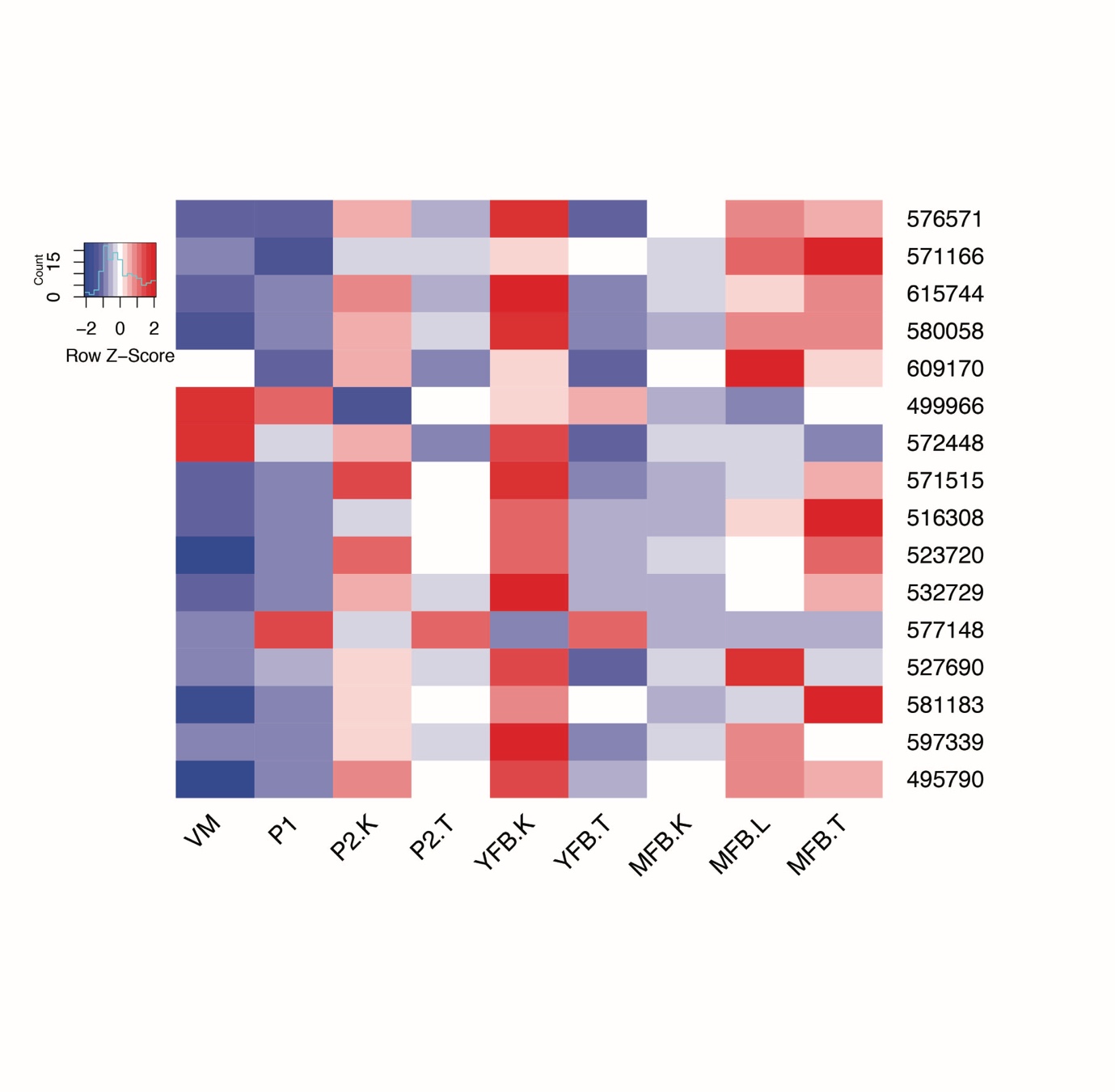

*Mycena kentingensis*

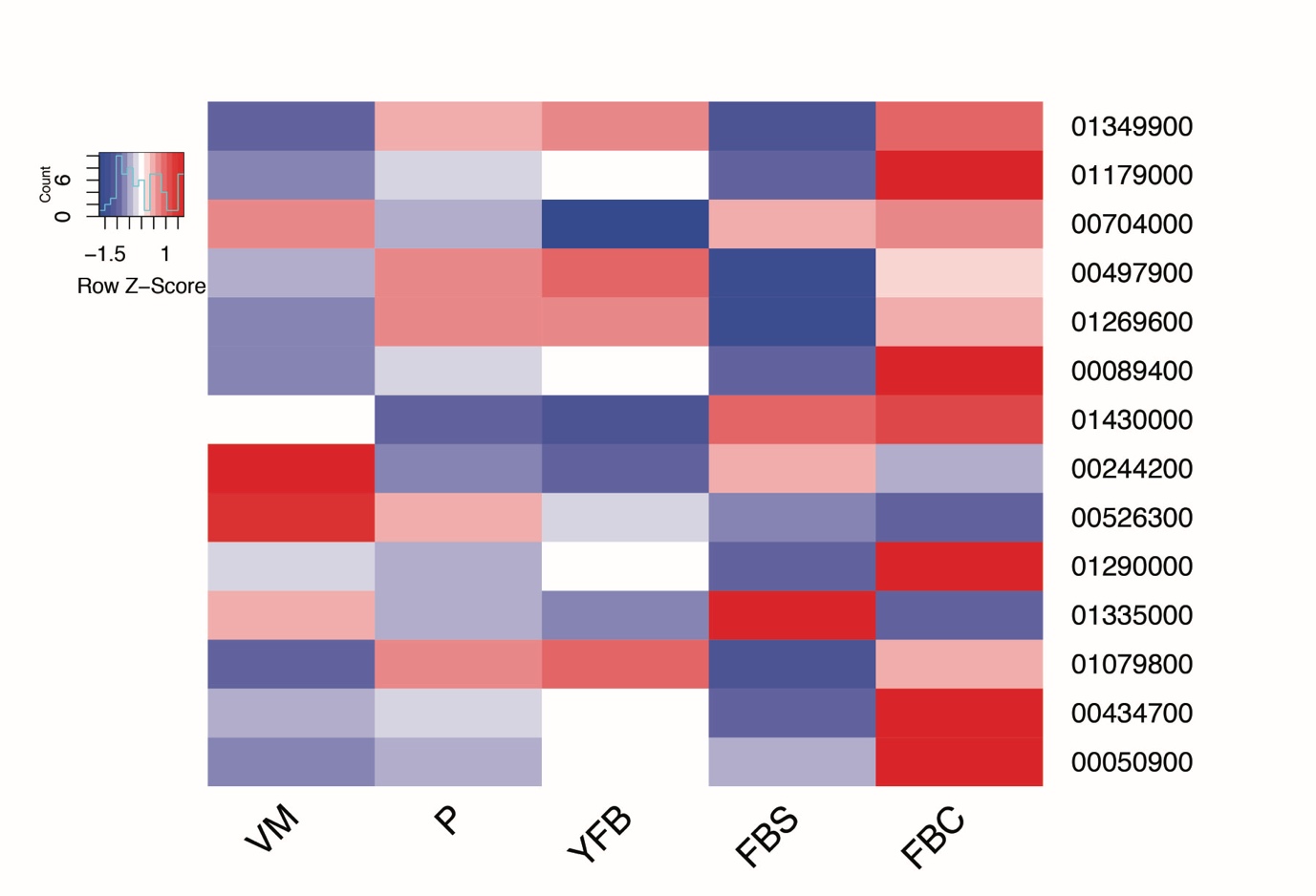

*Pleurotus ostreatus*

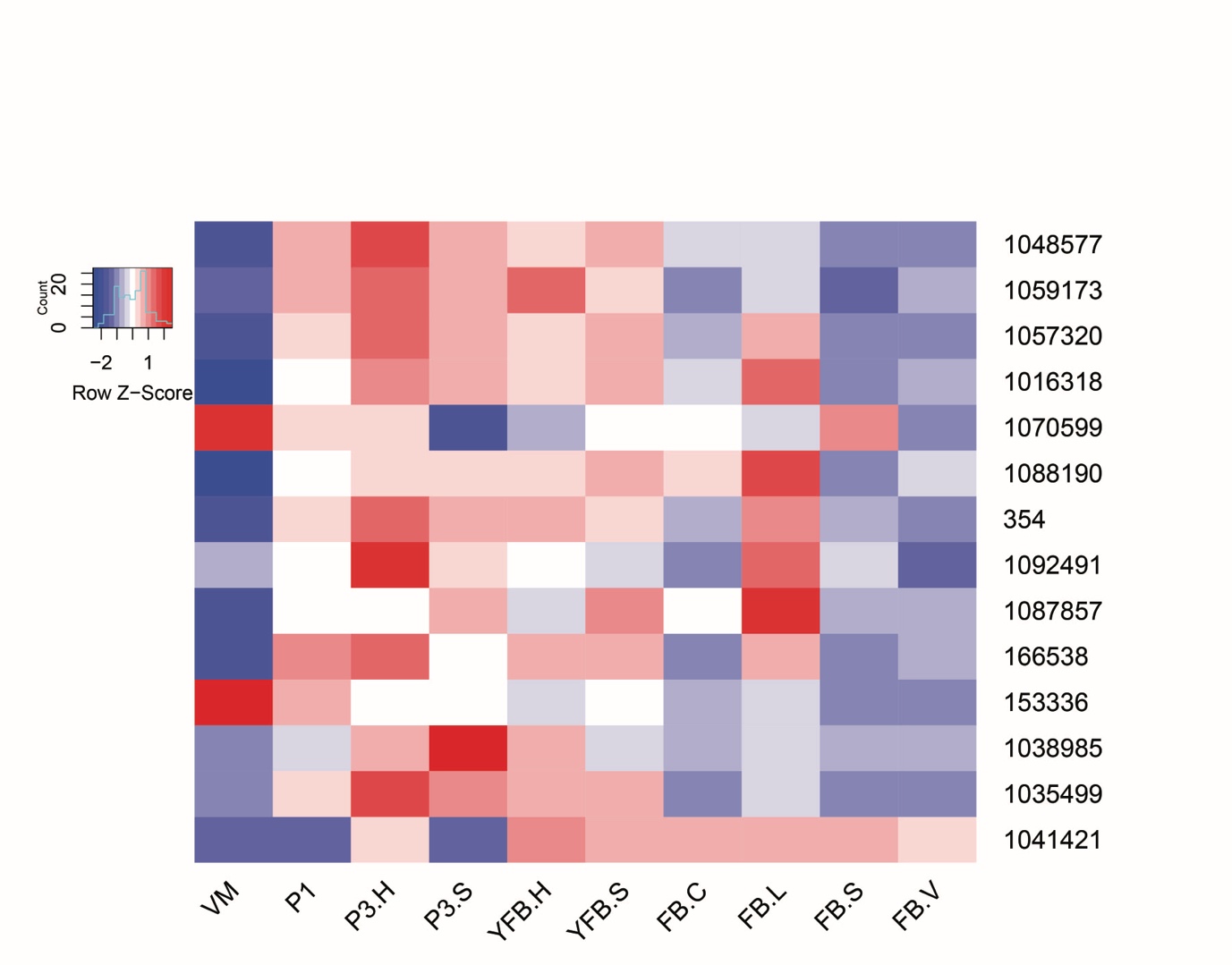

**Supplementary Fig. 8.** Expression heatmap of Con6 family cell surface protein-encoding genes in *A. ostoyae, A. ampla, L. tigrinus, M. kentingensis* and *P. ostreatus*. Genes are denoted by Protein IDs. Blue and red colors represent low and high expression, respectively.

*Auriculariopsis ampla*

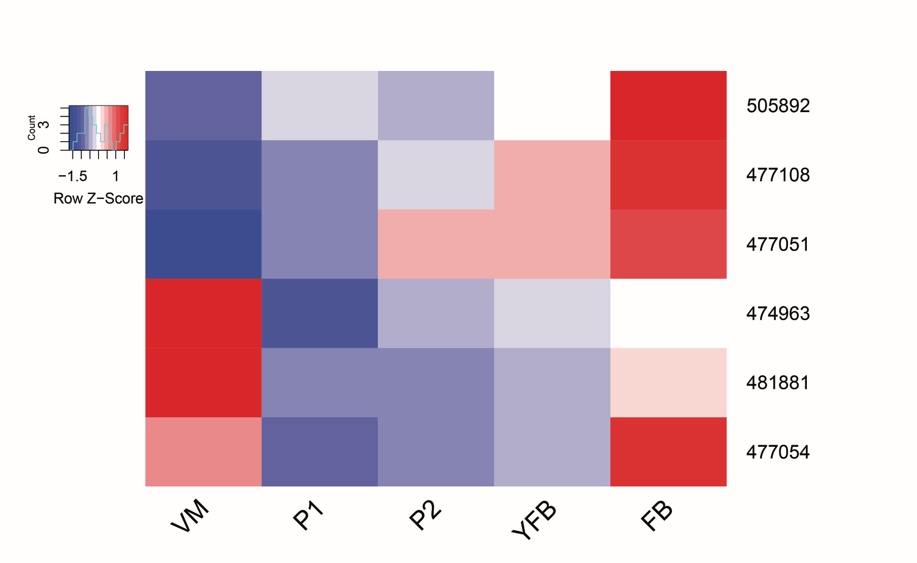

*Armillaria ostoyae*

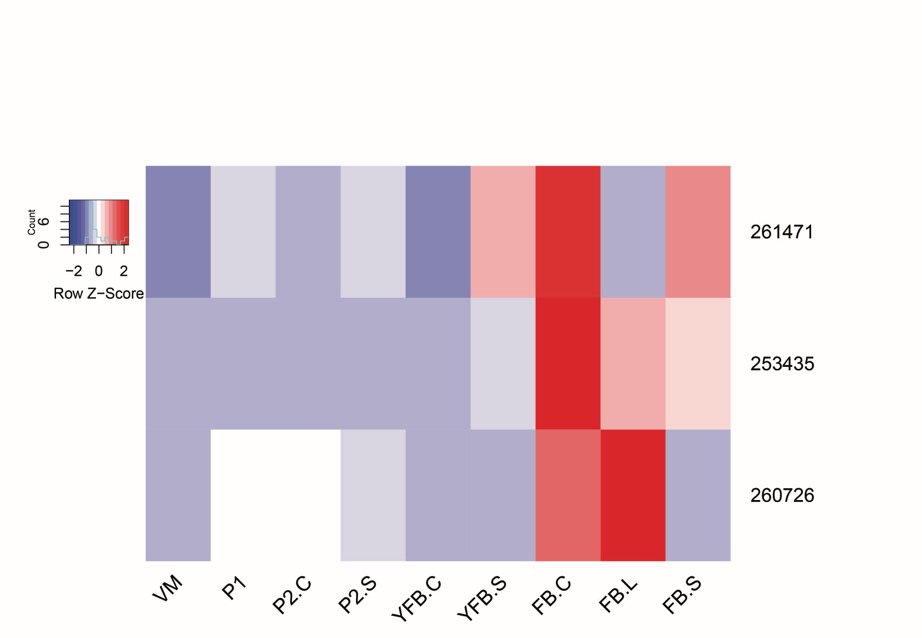

*Lentinus tigrinus*

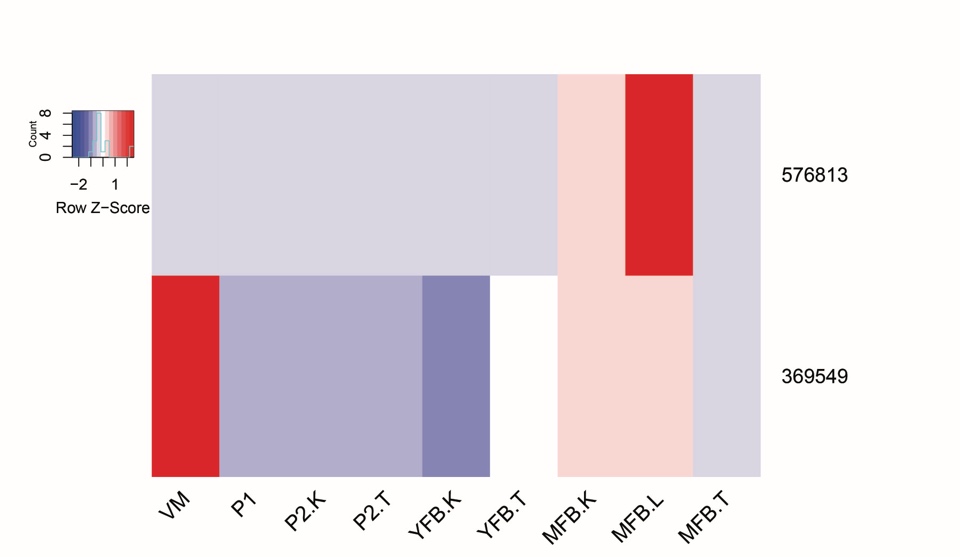

*Mycena kentingensis*

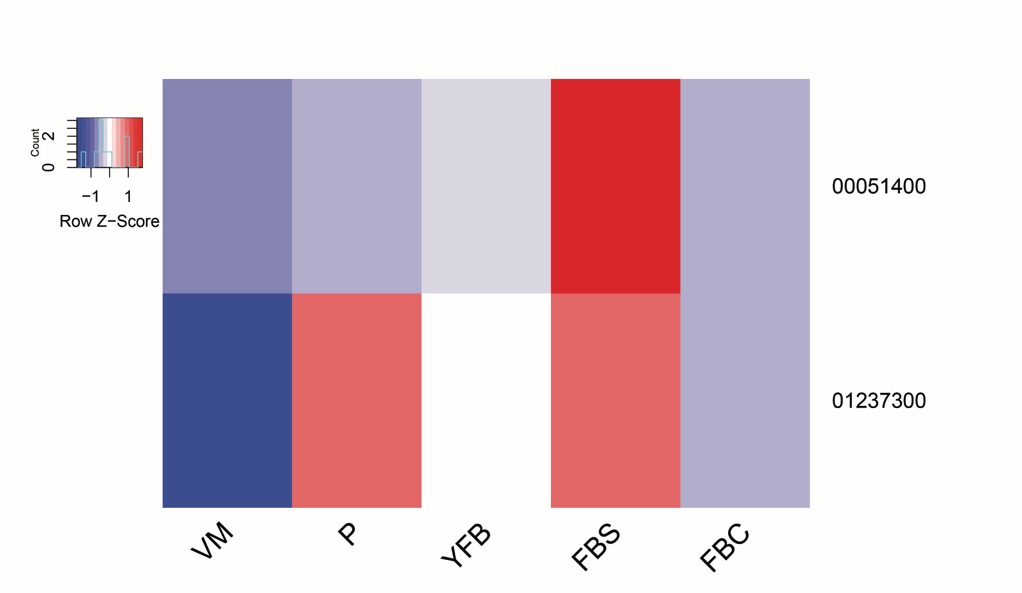

*Pleurotus ostreatus*

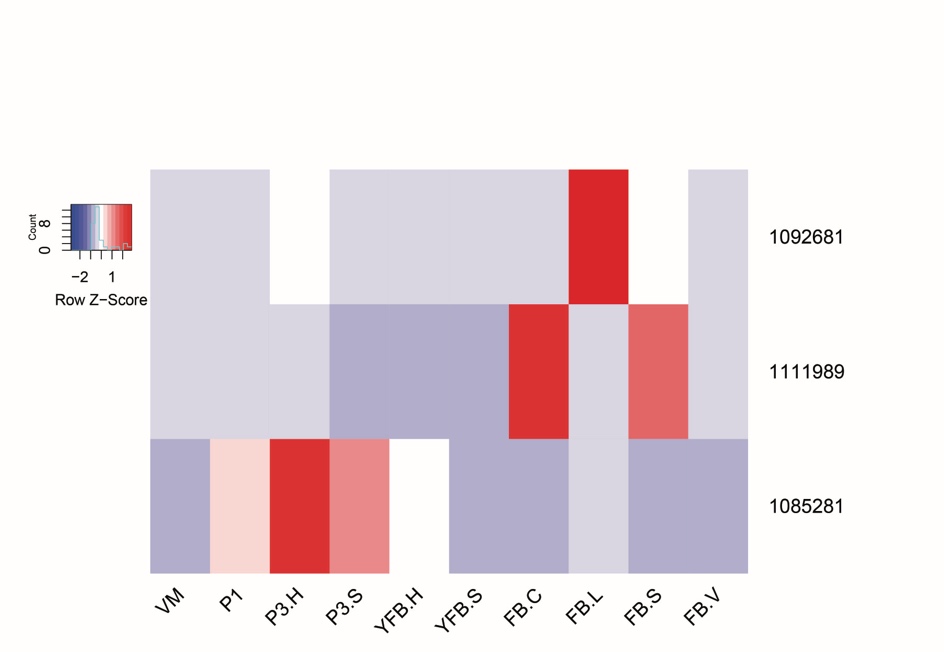

**Supplementary Fig. 9.** Expression heatmap of putative wax synthase encoding genes in *A. ostoyae* and *S. commune*. Genes are denoted by Protein IDs. Blue and red colors represent low and high expression, respectively.

*Armillaria ostoyae*

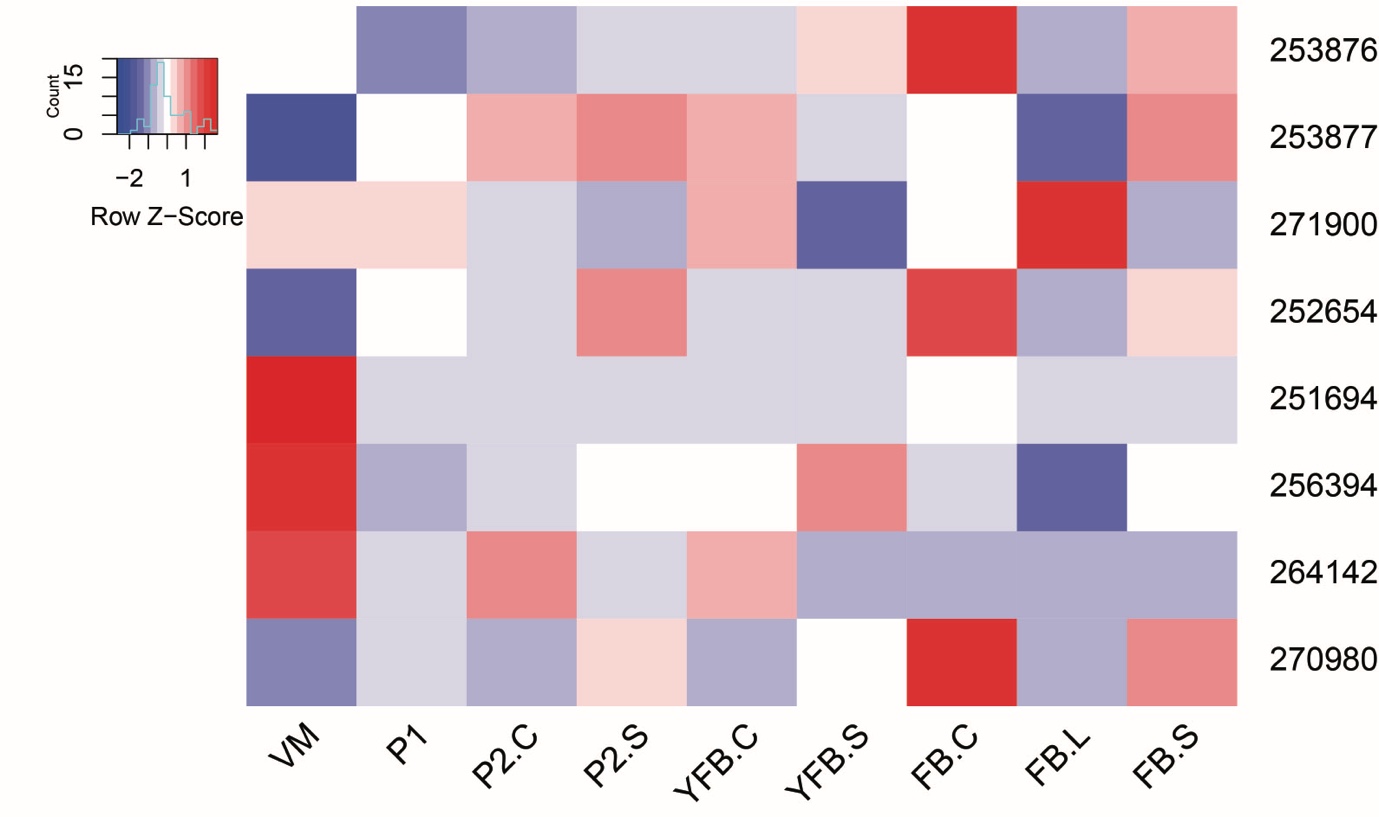

*Schizophyllum commune*

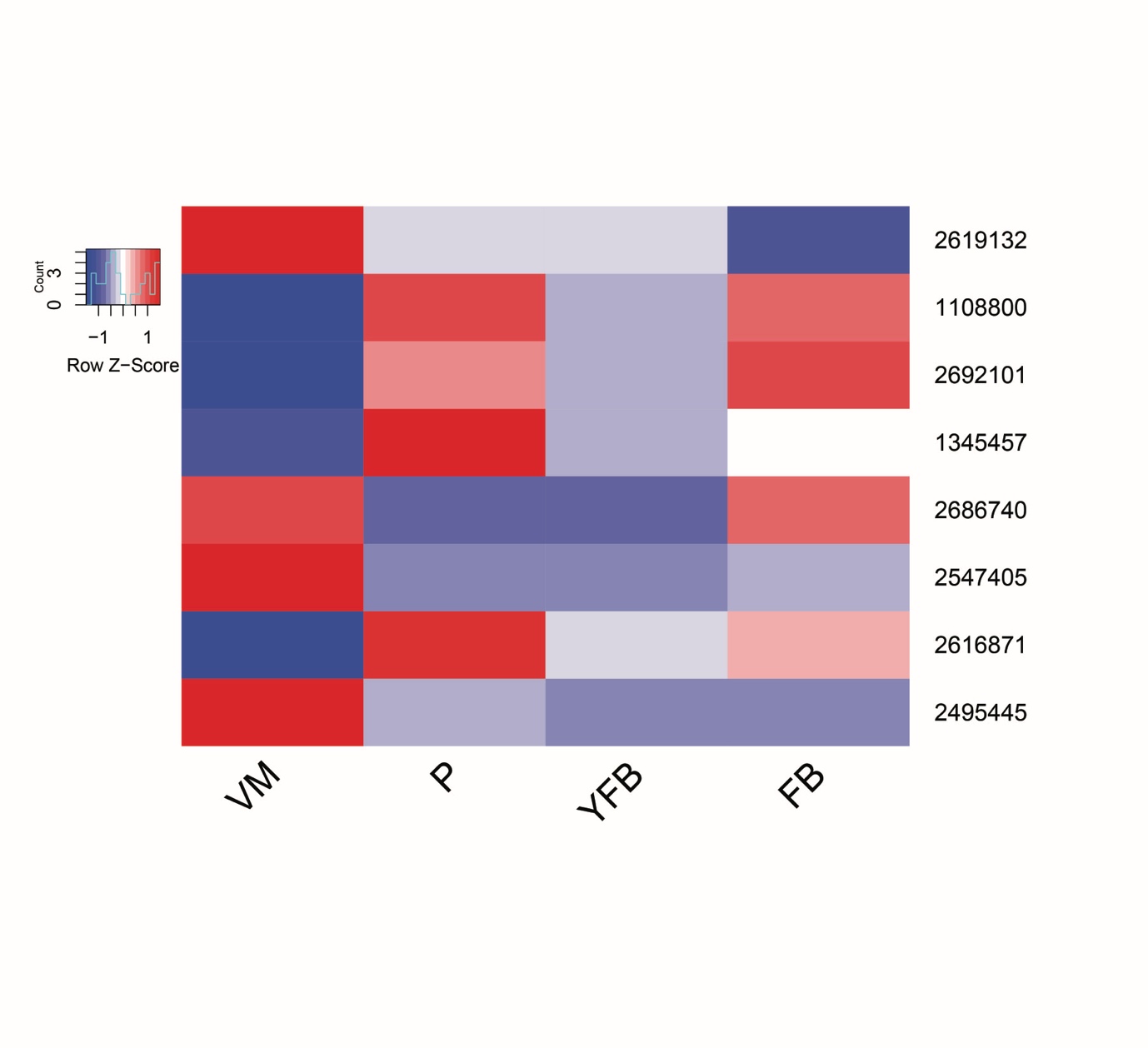

**Supplementary Fig. 10.** Expression heatmap of kinesin genes and members of the Dynactin complex in *A. ostoyae* and *P. ostreatus*. Genes are denoted by Protein IDs. Blue and red colors represent low and high expression, respectively.

*Armillaria ostoyae*

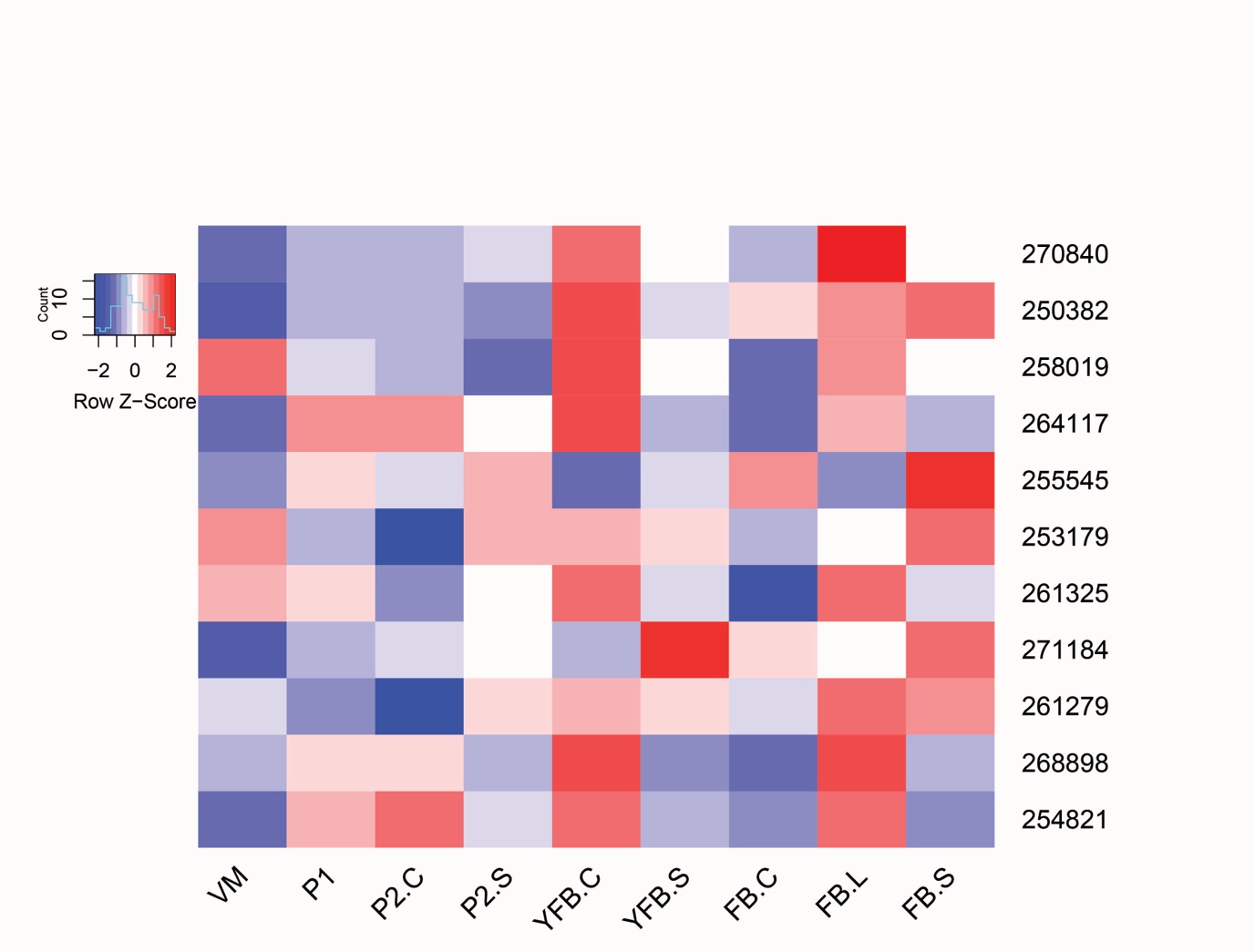

*Pleurotus ostreatus*

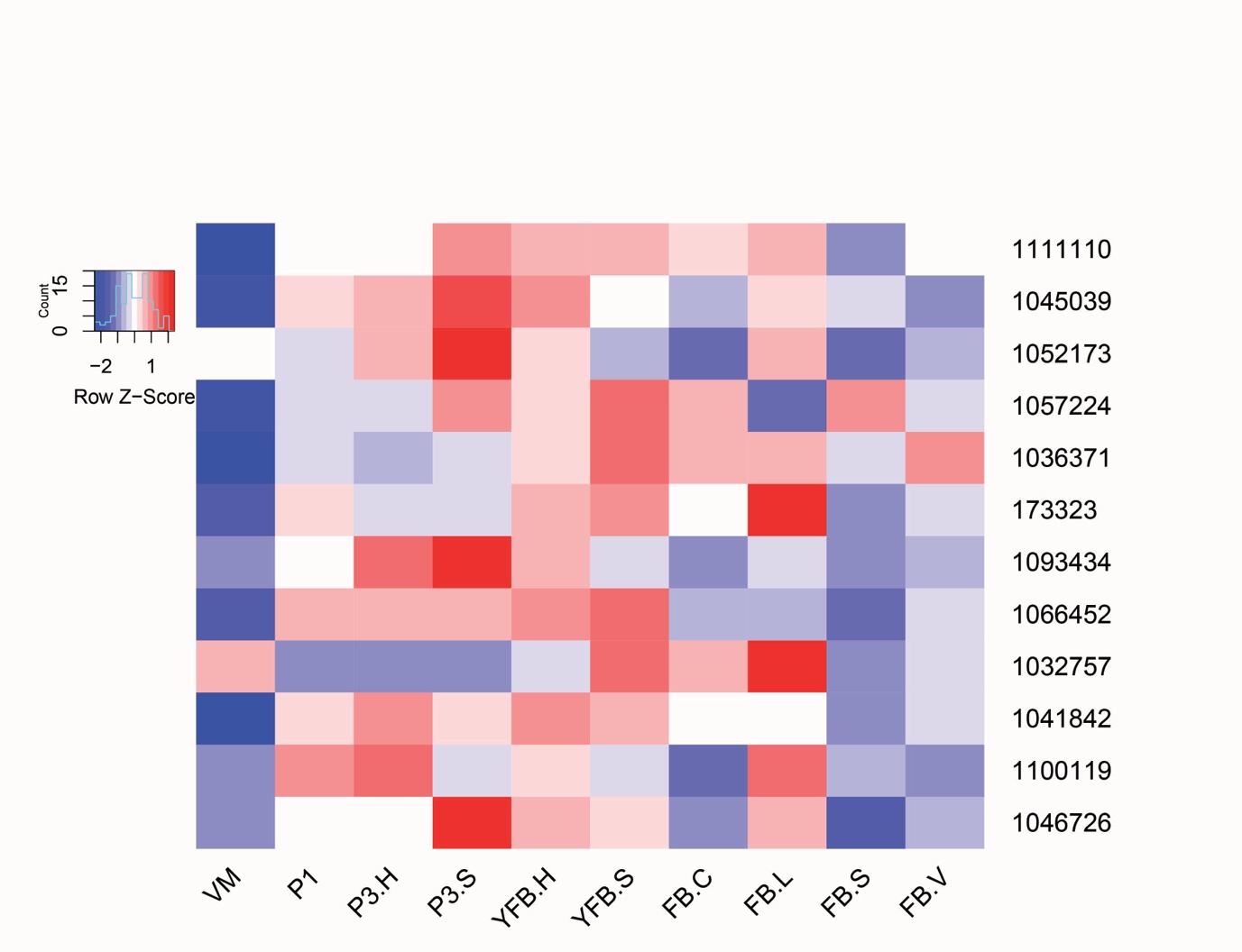
