## Supplementary Table 3 for "Lessons on fruiting body morphogenesis from genomes and transcriptomes of Agaricomycetes"

| **ID** | **FC>2** | **FC>4** | **putative function** | ***S. cerevisiae* ortholog** | **C.c.** | **A.a.** | **A.b.** | **A.o.** | **C.a.** | **L.b.** | **L.e.** | **L.t.** | **M.k.** | **P.c.** | **P.o.** | **S.c.** |
| --- | --- | --- | --- | --- | --- | --- | --- | --- | --- | --- | --- | --- | --- | --- | --- | --- |
| 16009 | 2 | 0 | putative splicing factor | YGR091W | 1 | 1 | 1 | 1 | 1 | 1 | 1 | 1 | 1 | 1 | 1 | 1 |
| 184391 | 2 | 0 | putative splicing factor | YER029C | 2 | 1 | 1 | 1 | 1 | 1 | 1 | 1 | 1 | 1 | 1 | 2 |
| 248479 | 5 | 1 | Pre-mRNA-splicing factor ATP-dependent RNA helicase PRP43 | YGL120C | 1 | 1 | 2 | 2 | 2 | 1 | 1 | 2 | 1 | 1 | 4 | 1 |
| 352445 | 2 | 0 | 5'-3' exoribonuclease 2 | YOR048C | 2 | 1 | 0 | 1 | 2 | 0 | 1 | 0 | 1 | 1 | 0 | 1 |
| 354362 | 12 | 4 | splicing factor |  | 2 | 2 | 4 | 2 | 2 | 2 | 2 | 2 | 4 | 2 | 4 | 4 |
| 358513 | 9 | 1 | U6 snRNA-associated Sm-like protein LSm3 | YLR438C-A | 2 | 1 | 1 | 2 | 1 | 2 | 2 | 2 | 2 | 2 | 4 | 2 |
| 359499 | 5 | 2 | Small nuclear ribonucleoprotein F | YPR182W | 4 | 1 | 0 | 2 | 0 | 2 | 1 | 1 | 0 | 2 | 4 | 1 |
| 360237 | 4 | 0 | putative splicing factor | YDR364C | 1 | 2 | 2 | 2 | 1 | 1 | 1 | 1 | 1 | 1 | 2 | 1 |
| 360732 | 2 | 0 | putative splicing factor | YDR473C | 1 | 1 | 2 | 1 | 1 | 1 | 1 | 1 | 1 | 1 | 2 | 1 |
| 369664 | 8 | 0 | Small nuclear ribonucleoprotein Sm D1 | YGR074W | 2 | 1 | 1 | 2 | 1 | 2 | 2 | 2 | 2 | 2 | 2 | 1 |
| 371517 | 5 | 0 | U6 snRNA-associated Sm-like protein LSm4 | YER112W | 1 | 1 | 2 | 1 | 1 | 2 | 1 | 2 | 2 | 2 | 0 | 1 |
| 387776 | 4 | 0 | putative splicing factor | YFR005C | 2 | 1 | 1 | 0 | 2 | 1 | 2 | 1 | 1 | 1 | 2 | 1 |
| 388237 | 1 | 0 | putative splicing factor | YER013W | 1 | 1 | 2 | 1 | 1 | 1 | 1 | 0 | 1 | 1 | 1 | 1 |
| 398525 | 7 | 1 | U6 snRNA-associated Sm-like protein LSm7 | YNL147W | 2 | 1 | 2 | 2 | 1 | 1 | 1 | 2 | 2 | 1 | 4 | 2 |
| 398634 | 5 | 1 | Small nuclear ribonucleoprotein E | YOR159C | 1 | 1 | 2 | 2 | 0 | 2 | 2 | 1 | 0 | 1 | 4 | 1 |
| 413309 | 8 | 2 | mRNA binding protein |  | 2 | 1 | 2 | 1 | 0 | 2 | 2 | 4 | 2 | 2 | 4 | 1 |
| 414944 | 7 | 2 | 13 kDa ribonucleoprotein-associated protein | YEL026W | 2 | 1 | 0 | 2 | 2 | 1 | 2 | 2 | 4 | 1 | 4 | 1 |
| 423725 | 2 | 0 | putative splicing factor | YPR178W | 1 | 1 | 1 | 2 | 0 | 1 | 1 | 1 | 1 | 1 | 2 | 1 |
| 428683 | 6 | 0 | Protein HSH49 | YOR319W | 2 | 2 | 2 | 1 | 2 | 1 | 1 | 1 | 1 | 2 | 2 | 1 |
| 436797 | 4 | 2 | putative splicing factor | YPR082C | 1 | 2 | 1 | 1 | 1 | 2 | 4 | 1 | 1 | 1 | 4 | 1 |
| 438732 | 5 | 1 | Ubiquitin-like modifier HUB1 | YNR032C-A | 1 | 1 | 2 | 1 | 0 | 2 | 2 | 1 | 2 | 1 | 4 | 1 |
| 439633 | 7 | 1 | Small nuclear ribonucleoprotein G | YFL017W-A | 2 | 1 | 1 | 2 | 0 | 1 | 2 | 2 | 2 | 2 | 4 | 1 |
| 442124 | 3 | 0 | putative splicing factor | YDR416W | 1 | 1 | 2 | 1 | 2 | 1 | 1 | 1 | 1 | 1 | 2 | 1 |
| 442716 | 3 | 2 | putative splicing factor | YPL184C | 1 | 1 | 1 | 1 | 4 | 1 | 1 | 2 | 4 | 1 | 1 | 1 |
| 445902 | 4 | 1 | putative splicing factor | YPR094W | 1 | 1 | 2 | 1 | 0 | 1 | 2 | 1 | 1 | 2 | 4 | 1 |
| 449065 | 2 | 1 | putative splicing factor | YPL151C | 1 | 1 | 2 | 1 | 1 | 1 | 1 | 1 | 1 | 1 | 4 | 1 |
| 452104 | 2 | 0 | putative splicing factor | YLR116W | 2 | 1 | 1 | 2 | 1 | 1 | 1 | 1 | 1 | 1 | 1 | 1 |
| 452642 | 4 | 0 | putative splicing factor | YGR278W | 2 | 1 | 2 | 1 | 1 | 2 | 1 | 1 | 1 | 1 | 2 | 1 |
| 454246 | 1 | 0 | putative splicing factor | YML010W | 1 | 1 | 2 | 1 | 1 | 1 | 1 | 1 | 1 | 1 | 1 | 1 |
| 455625 | 3 | 1 | putative splicing factor | YKL012W | 1 | 1 | 2 | 1 | 0 | 2 | 1 | 1 | 1 | 1 | 4 | 1 |
| 456807 | 3 | 0 | Lariat debranching enzyme | YKL149C | 1 | 1 | 1 | 2 | 2 | 1 | 1 | 2 | 1 | 1 | 1 | 1 |
| 459073 | 6 | 1 | U1 small nuclear ribonucleoprotein 70 kDa homolog | YIL061C | 2 | 1 | 2 | 2 | 2 | 1 | 2 | 1 | 1 | 1 | 4 | 1 |
| 463489 | 7 | 0 | Small nuclear ribonucleoprotein Sm D3 | YLR147C | 2 | 1 | 1 | 2 | 1 | 2 | 2 | 1 | 2 | 2 | 2 | 1 |
| 469346 | 7 | 2 | U6 snRNA-associated Sm-like protein LSm2 | YBL026W | 4 | 2 | 2 | 2 | 0 | 1 | 1 | 1 | 2 | 2 | 4 | 1 |
| 469749 | 0 | 0 | putative splicing factor | YKR086W | 1 | 1 | 1 | 1 | 1 | 1 | 1 | 1 | 1 | 1 | 0 | 1 |
| 471522 | 5 | 0 | Pre-mRNA leakage protein 1 | YLR016C | 2 | 1 | 1 | 1 | 1 | 1 | 2 | 1 | 2 | 1 | 2 | 2 |
| 473906 | 2 | 1 | putative splicing factor | YPR045C | 2 | 1 | 1 | 4 | 0 | 1 | 1 | 1 | 1 | 1 | 1 | 1 |
| 484439 | 3 | 0 | putative splicing factor | YNL286W | 2 | 1 | 1 | 1 | 2 | 1 | 2 | 1 | 1 | 1 | 1 | 1 |
| 494883 | 5 | 3 | U6 snRNA-associated Sm-like protein LSm5 | YER146W | 4 | 0 | 1 | 2 | 0 | 4 | 1 | 1 | 4 | 1 | 2 | 1 |
| 495814 | 5 | 1 | splicing factor C. cinerea Cdc5 |  | 2 | 1 | 2 | 1 | 2 | 1 | 1 | 2 | 1 | 1 | 4 | 1 |
| 499042 | 7 | 1 | Pre-mRNA-splicing factor CLF1 | YLR117C | 2 | 1 | 2 | 1 | 2 | 2 | 2 | 2 | 1 | 1 | 4 | 1 |
| 501262 | 7 | 1 | Serine/threonine-protein kinase SKY1 | YMR216C | 2 | 1 | 1 | 2 | 2 | 2 | 1 | 2 | 4 | 1 | 2 | 1 |
| 510352 | 3 | 1 | putative splicing factor | YDL030W | 1 | 2 | 2 | 1 | 1 | 1 | 1 | 1 | 1 | 1 | 4 | 1 |
| 520307 | 1 | 0 | putative splicing factor | YBR237W | 1 | 1 | 1 | 1 | 1 | 1 | 1 | 1 | 1 | 1 | 2 | 1 |
| 533202 | 3 | 1 | putative splicing factor | YDL043C | 1 | 1 | 2 | 0 | 1 | 1 | 2 | 1 | 1 | 1 | 4 | 1 |
| 538515 | 6 | 1 | ATP-dependent RNA helicase SUB2 | YDL084W | 2 | 1 | 2 | 0 | 1 | 1 | 2 | 2 | 2 | 1 | 4 | 1 |
| 539514 | 7 | 1 | U2 small nuclear ribonucleoprotein A' | YPL213W | 1 | 2 | 1 | 2 | 1 | 2 | 2 | 1 | 2 | 2 | 4 | 1 |
| 542441 | 7 | 1 | Pre-mRNA-splicing factor BUD31 | YCR063W | 1 | 2 | 1 | 0 | 1 | 2 | 2 | 2 | 2 | 2 | 4 | 1 |
| 543037 | 7 | 1 | Small nuclear ribonucleoprotein Sm D2 | YLR275W | 2 | 2 | 0 | 2 | 1 | 2 | 1 | 2 | 2 | 1 | 4 | 1 |
| 545156 | 5 | 0 | Pre-mRNA-splicing factor CWC2 | YDL209C | 1 | 2 | 2 | 1 | 2 | 1 | 1 | 1 | 1 | 1 | 2 | 2 |
